# *In vivo* commensal control of *Clostridioides difficile* virulence

**DOI:** 10.1101/2020.01.04.894915

**Authors:** Brintha P. Girinathan, Nicholas DiBenedetto, Jay N. Worley, Johann Peltier, Mario L. Arrieta-Ortiz, Rupa Immanuel, Richard Lavin, Mary L. Delaney, Christopher Cummins, Maria Hoffmann, Yan Luo, Narjol Gonzalez Escalona, Marc Allard, Andrew B. Onderdonk, Georg K. Gerber, Abraham L. Sonenshein, Nitin Baliga, Bruno Dupuy, Lynn Bry

## Abstract

We define multiple mechanisms by which commensals protect against or worsen *Clostridioides difficile* infection. Leveraging new systems-level models we show how metabolically distinct species of *Clostridia* modulate the pathogen’s colonization, growth, and virulence to impact host survival. Gnotobiotic mice colonized with the amino acid fermenter *Paraclostridium bifermentans* survived infection while mice colonized with the butyrate-producer, *Clostridium sardiniense,* more rapidly succumbed. Systematic *in vivo* analyses revealed how each commensal altered the gut nutrient environment, modulating the pathogen’s metabolism, regulatory networks, and toxin production. Oral administration of *P. bifermentans* rescued conventional mice from lethal *C. difficile* infection via mechanisms identified in specifically colonized mice. Our findings lay the foundation for mechanistically informed therapies to counter *C. difficile* infections using systems biologic approaches to define host-commensal-pathogen interactions *in vivo*.

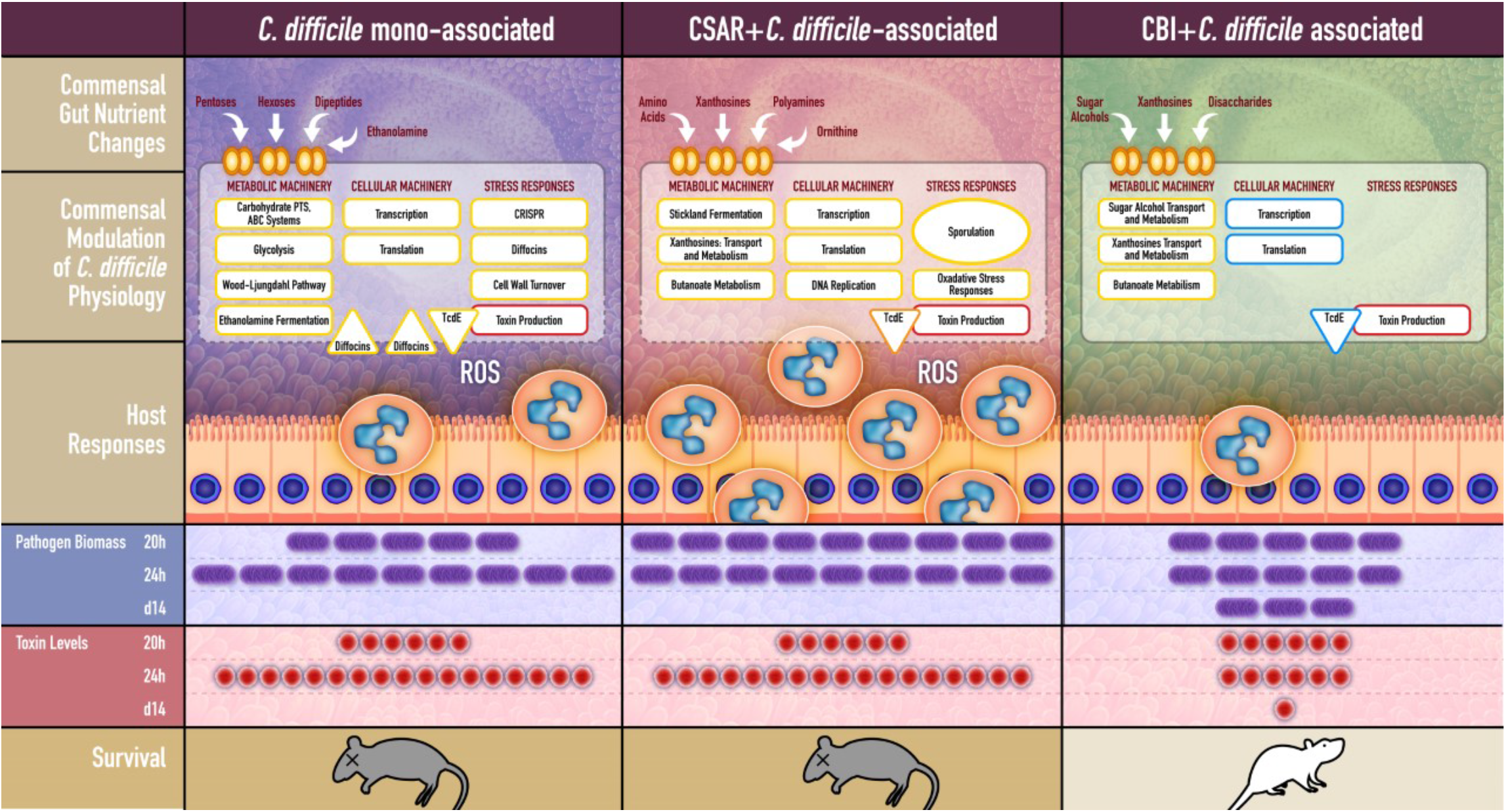
Graphical Abstract.

## Introduction

*Clostridioides difficile* infections cause substantial morbidity, mortality, and healthcare costs in the US (Allegretti et al., 2019; Worley et al., 2020). Fecal microbiota transplant (FMT) has become standard of care to treat recurrent infections. However, we know little about the molecular mechanisms by which specific microbes modulate the pathogen’s virulence *in vivo*. Given FMT-related deaths in immunocompromised patients, therapies informed by molecular mechanisms of action will enable options with improved safety and efficacy (Giles et al., 2019).

*C. difficile*’s pathogenicity locus (PaLoc) contains the TcdA and TcdB toxin genes and TcdE holin involved in toxin release. TcdR encodes a sigma factor specific for the toxin gene promoters (Bouillaut et al., 2015; Mani and Dupuy, 2001). *C. difficile*, like other cluster XI *Clostridia*, utilizes diverse carbon sources *in vivo*, including carbohydrates, amino acids fermented via Stickland reactions, and ethanolamine (Dubois et al., 2016). The Stickland pathways can drive rapid pathogen growth, particularly with abundant proline, glycine or leucine, which serve as electron acceptors for the *prd*, *grd* and *had* enzyme systems, respectively (Bouillaut et al., 2015; Hofmann et al., 2018; Neumann-Schaal et al., 2015). The pathogen can also fix carbon through the Wood-Ljungdhal pathway to generate acetate for metabolism (Nawrocki et al., 2018; Neumann-Schaal et al., 2019). *C. difficile’s* conditional regulatory networks closely integrate its cellular metabolism and virulence programs, particularly through the CodY and CcpA metabolic repressors that repress PaLoc expression under nutrient sufficient states (Martin-Verstraete et al., 2016) and promote toxin expression under starvation conditions to extract nutrients from the host. CodY and CcpA co-regulate additional gene systems involved in cellular metabolism, sporulation, and stress responses, by sensing factors released from the commensal microbiota and host-factors including inflammatory ROS and antimicrobial peptide responses (Kint et al., 2017; Neumann-Schaal et al., 2018).

Host and microbiota can thus influence the pathogen’s virulence through multiple mechanisms. Among Stickland-fermenting Cluster XI Clostridia, *Paraclostridium bifermentans* (PBI) preferentially uses Stickland fermentations for energy (Moore, 1993). In contrast, *Clostridium sardiniense* (CSAR), a strongly glycolytic Cluster I species, produces abundant butyrate through anaerobic carbohydrate fermentation (Moore, 1993). Both species colonize the human gut yet have very different metabolic capabilities.

Using defined colonization experiments in gnotobiotic mice with Environmental and Gene Regulatory Influence Network (EGRIN) and Phenotype of Regulatory Influences integrated with Metabolism and Environment (PRIME) models of *C. difficile’s* conditional regulatory networks (Arrieta-Ortiz ML, 2020), we show mechanisms by which individual *Clostridial* species affect host survival of *C. difficile* infection, to the level of the microbial pathways and small molecules that modulate *C. difficile’s* regulatory networks to drive pathogen growth and virulence. Findings promoted development of a defined bacteriotherapeutic able to rescue an infected host from lethal infection. By defining how individual commensals systematically modulate *C. difficile’s* physiology we enable mechanistically informed approaches for this disease.

## Results

### Single commensals dramatically alter host outcomes from *C. difficile* infection

*C. difficile* infection of 6-week old germfree mice caused rapid demise within 48h (Figs. 1A-B). Symptoms developed at 20h post-challenge, manifested by weight loss (Figs. S1A-B) and diarrhea. Animals demonstrated focal epithelial damage with neutrophilic infiltrates (Figs. 1C vs 1D) that by 24h had rapidly progressed to severe colitis with widespread erosions and worsening symptomatic disease (Fig. S1C).

**Figure 1:**
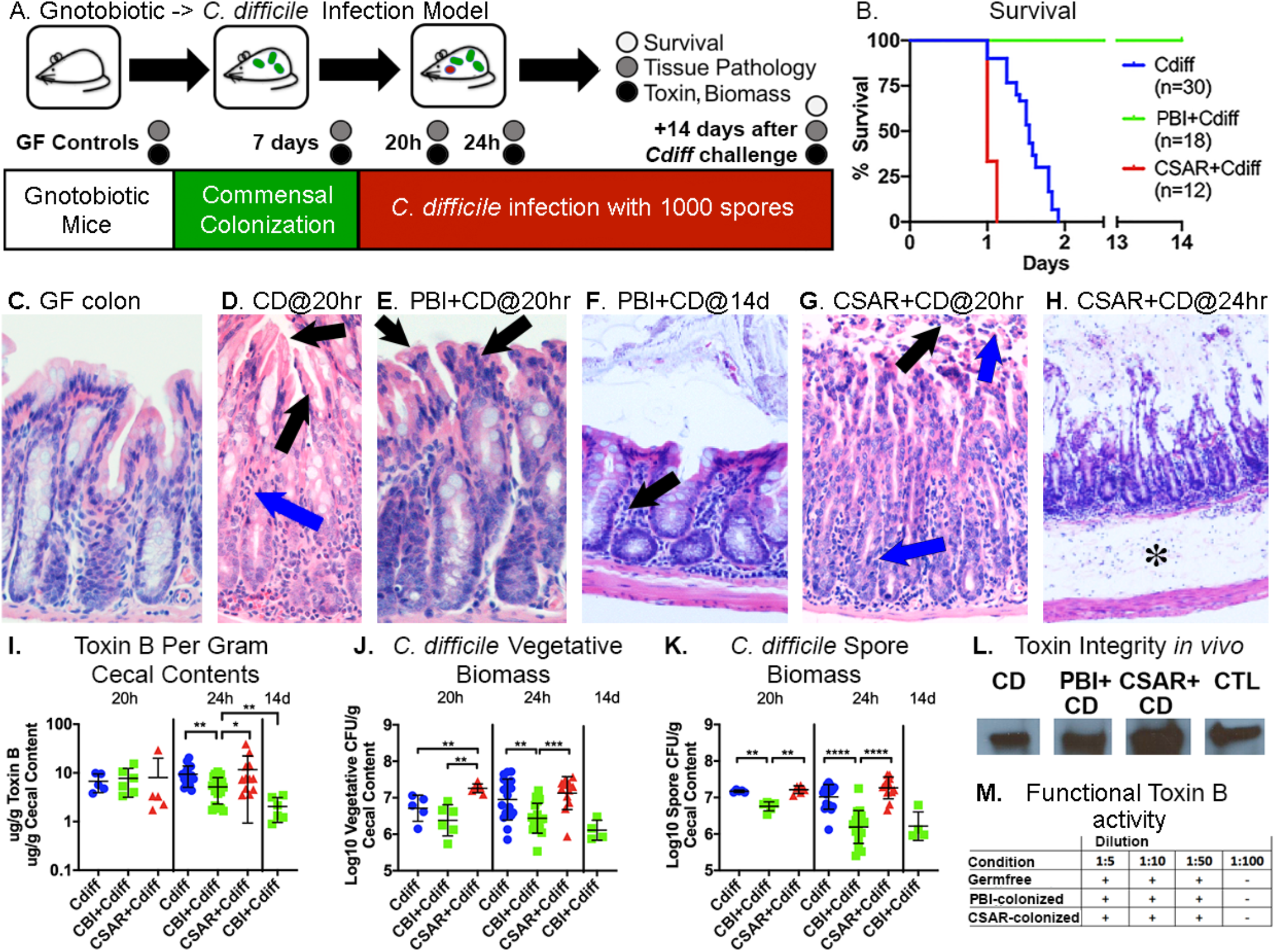
PBI protects germfree mice from lethal *C. difficile* infection while CSAR promotes more severe disease. **A**: Experimental overview. **B**: Survival curves. **C-H:** Colonic H&E stains; C-E: 200X, F-G: 100X and H: 40X magnification. **C:** Normal germfree mucosa. **D:** *C. difficile*- infected mice at 20h, showing epithelial stranding and vacuolation (black arrows) and neutrophilic infiltrates (blue arrow). **E:** PBI co-colonzied mice at 20h of infection showing vacuolization of apical colonocytes (black arrows) but nominal inflammation. **F:** PBI and *C.* difficile-infected mice at 14d showing intact epithelium and lymphocytic infiltrates (black arrow). **G:** CSAR and *C. difficile*-infected mice at 20h of infection showing surface epithelial loss with pseudomembrane formation (black arrow) and transmural neutrophilic infiltrates entering the lumen (blue arrows). **H:** CSAR co-colonized mice at 24h of infection showing complete epithelial loss and severe submucosal edema (asterisk). **I:** Log10 ug/g of extracellular cecal Toxin B. Significance values by Mann-Whitney test: *0.01<p*≤*0.05; **0.001<p*≤*0.01; ***0.0001<p*≤*0.001; ****p*≤*0.0001. **J.** Log10 *C. difficile* vegetative CFU, and **K:** spore biomass/gram of cecal contents. **L.** Western blot of cecal contents showing intact toxin B in *C. difficile* infected (CD), PBI (PBI+CD), or CSAR (CSAR+CD) co-colonized mice; CTL = control toxoid B. **M.** Effects of GF, PBI or CSAR monocolonized cecal contents on the toxicity of exogenously added toxin B, no differences were noted in toxin B cytopathic effect against human fibroblasts.

In contrast, mice pre-colonized with PBI prior to *C. difficile* challenge survived (Fig. 1B; p<0.0001) with milder symptoms and colonic damage (Figs. 1E vs S1D; Figs. S1A-B). Fourteen days post-infection animals had regained lost weight and demonstrated intact intestinal epithelium with a lymphocytic infiltrate having replaced acute neutrophilic infiltrates (Fig. 1F).

Mice pre-colonized with CSAR developed more rapidly lethal infection (Fig 1B; p<0.0001), with areas of epithelial sloughing and blood entering the lumen by 20h of infection (Figs. S1E vs. 1G), followed by widespread mucosal denudation and rapid demise (Fig. 1H).

Though toxin levels were comparable among mice at 20h of infection (Fig. 1I), pathogen vegetative and spore biomass in CSAR co-colonized mice were 3-fold higher (Fig. 1J-K). By 24h *C. difficile* monocolonized and CSAR co-colonized mice demonstrated 2-3-fold higher toxin levels and vegetative biomass than PBI co-colonized mice (Fig. 1I-J). After 14 days, toxin levels in surviving PBI co-colonized mice fell >80% from acute levels (Fig. 1I).

While CSAR biomass rose 10-fold in *C. difficile-*infected mice (Fig S1F), PBI biomass did not change over acute infection (Fig. S1G). Commensal colonization also did not alter toxin integrity or cytotoxic activity (Fig. 1L-M).

### Commensals condition the gut nutrient environment prior to *C. difficile’s* introduction

Given the effects of commensal colonization on pathogen biomass, carbon source enrichment analyses of the gut metabolomic environment evaluated the nutrient content available for *C. difficile* growth and metabolism (Figs. 2A-B; Supplemental Text, SDF_2-3).

**Figure 2:**
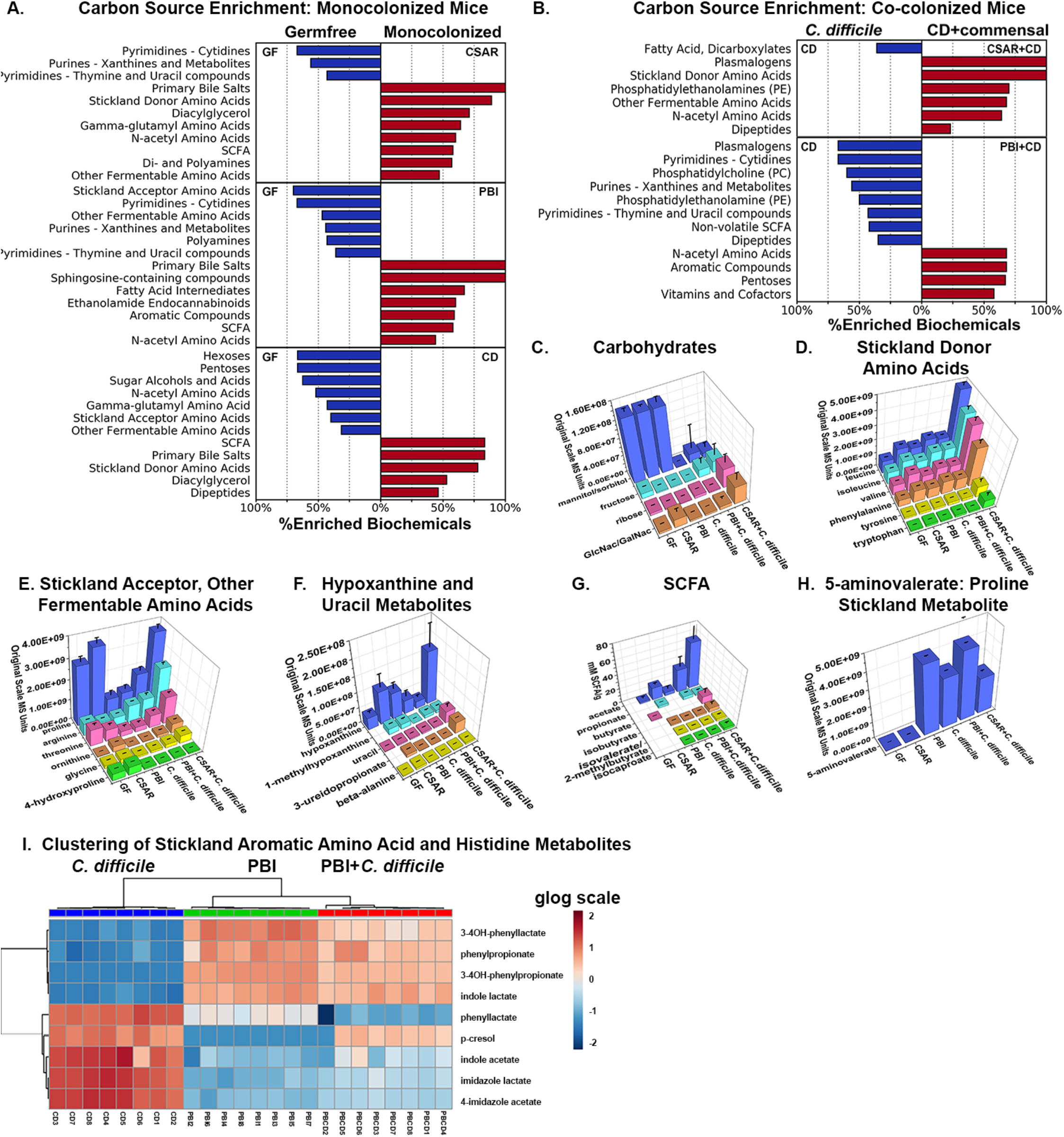
Commensal alteration of cecal carbon sources. **A:** Significantly enriched carbon source groups in germfree (left-hand side) versus monocolonized mice (right-hand side), showing groups with a Benjamini-Hochberg corrected p value≤0.05. Horizontal bars indicate the percent of biochemicals enriched in each carbon source group comparing germfree cecal contents with CSARmonocolonized mice at 7d (top), PBI monocolonized mice at 7d (middle), or *C. difficile* monocolonized for 20h (bottom). **B.** Significantly enriched carbon sources between *C. difficile* monocolonized mice at 20h of infection (right-hand side) vs CSAR monocolonized for 7d + 20h of *C. difficile* infection (top) or PBI monocolonized for 7d + 20h of *C. difficile* infection (bottom). **C-H:** Specifically enriched compounds (Y-axis) across colonization states (X-axis). Z-axis shows original scale mass spectrometry counts. Error bars indicate standard deviation. Values for a given compound are comparable across experimental groups. **C.** Carbohydrates. **D:** Stickland donor amino acids. **E.** Stickland acceptor amino acids, and other fermentable amino acids. **F:** Hypoxanthine and uracil metabolites. **G:** Cecal SCFA profiles. Z-axis shows mM of SCFA per gram of cecal contents**, H.** Proline Stickland metabolite 5-aminovalerate.. **I:** Heatmap of *C. difficile* and PBI-specific Stickland aromatic amino acid and histidine metabolites in cecal contents at 20h of infection. Color scale shows glog transformed MassSpec counts for metabolites; red–increased levels, blue–decreased levels. Hierarchical clustering by Pearson similarity and minimum-distance linkage are shown by sample along the top and by metabolite on the left Y axis.

Germfree cecal contents were enriched for multiple classes of carbohydrates (Fig 2A, 2C; SDF_3.8, 3.11, 3.25), including hexoses, pentoses, and sugar alcohols of dietary origin that have poor absorption from the gut (Theriot et al., 2014). Gut contents were also enriched for multiple fermentable amino acids (Figs. 2D-E; SDF_3.10, 3.23) including the Stickland acceptor amino acid proline (Fig. 2E), and multiple purines and pyrimidines (Fig. 2A, 2F, SDF_3.17-3.18). In the absence of colonizing microbiota SCFA were not detected (Fig. 2G).

Colonization with CSAR or with PBI markedly changed the luminal environment prior to *C. difficile’s* introduction. CSAR monocolonization enriched multiple amine-containing carbon sources (Fig 2A, top), including Stickland-fermentable amino acids (Figs. D-E; SDF_3.10, 3.24), *γ*-glutamyl-amino acids (SDF_3.7) (Griffith et al., 1979), and polyamines (SDF_3.15). Branched chain amino acids increased 2-3-fold (Fig. 2D) and ornithine >16-fold over germfree levels (Fig. 2E).

CSAR depleted luminal fructose, left mannitol/sorbitol unchanged, and enriched amino sugars, including ones originating from host glycoconjugates (Fig. 2C). CSAR monocolonization produced >5-fold increases in hypoxanthines (SDF_3.17), metabolites produced by other purinolytic *Clostridia* (Bradshaw, 1960) (Fig. 2F), and >10-fold increases in the uracil metabolites 3-ureidopropionate and beta-alanine (Vogels and Van der Drift, 1976) (Fig 2F; SDF_3.19). SCFA fermentation metabolites acetate and butyrate were produced (Fig. 2G).

In contrast, PBI monocolonization depleted polyamines, Stickland acceptor and other fermentable amino acids (Fig. 2A, middle), consuming >70% of proline, >50% of glycine, >50% of threonine, and 80% of 4-hydroxyproline, a collagen degradation product which many Stickland fermenters convert to proline (Fig. 2E; (Huang et al., 2018)). PBI produced abundant 5- aminovalerate (Fig. 2H) from proline, isocaproate from reductive leucine Stickland metabolism (Kim et al., 2006), and isobutyrate, isovalerate and 2-methylbutyrate from Stickland oxidative fermentation of branched-chain amino acids (Fig. 2G). Stickland aromatic amino acid metabolites including 3-phenylpropionate from phenylalanine, 3-(4-hydroxy-phenylpropionate) from tyrosine, and indole lactate from tryptophan were also produced (Fig. 2I, Supplemental Text).

PBI consumed >50% of fructose, left sugar alcohol levels unchanged (Fig 2C; SDF_3.25) and produced acetate and propionate (Fig 2G). PBI monocolonization enriched hypoxanthines (Fig 2F; SDF_3.17-3.19). 3-ureidopropionate increased to a lesser extent than in CSAR monocolonized mice, and without increased beta-alanine (Fig 2F; SDF_3.19).

*C. difficile* monocolonized mice demonstrated the broadest depletion of carbohydrate and amine-containing carbon sources (Fig 2A, bottom). The pathogen depleted Stickland acceptor amino acids consuming >50% of glycine, >70% of proline and >85% of 4-hydroxyproline (Fig. 2E), with concomitant increase in 5-aminovalerate (Fig. 2H). *γ*-glutamyl-amino acids (SDF_3.7), other fermentable amino acids including cysteine and threonine (SDF3.10), and N-acetyl conjugates of proline, branched-chain amino acids, and polyamines were also depleted (SDF_3.9, 3.10, 3.15). Hexoses, pentoses and sugar alcohols were depleted, including >99% of mannitol/sorbitol and >80% of fructose (Fig. 2C).

*C. difficile* monocolonization produced acetate (Fig. 2G) and Stickland branched-SCFA metabolites isobutyrate, isovalerate, 2-methylbutyrate and isocaproate (Fig. 2G). Aromatic amino acid metabolites specific to the pathogen’s Stickland metabolism were also produced, including phenylacetate and phenyllactate from phenylalanine, indole acetate from tryptophan, and *p*-cresol from the *p*-hydroxyphenylacetate metabolite of tyrosine metabolism (Passmore et al., 2018; Steglich et al., 2018), (Fig. 2I, Supplemental Text). *C. difficile* also has unique capacity to metabolize histidine in Stickland reactions, producing imidazole lactate and 4-imidazole acetate (Neumann-Schaal et al., 2019) (Fig. 2I).

By 24h of infection, with deteriorating mucosal conditions (Figs. 1G-H), CSAR and *C. difficile* co-colonized mice (Fig. 2B, top) demonstrated further enrichment of Stickland and other fermentable amino acids (Fig. 2D-E). Uracil levels increased >8-fold and 3-ureidopropionate increased >40-fold compared to *C. difficile* monocolonized mice (Fig. 2F; SDF_3.19). In contrast, PBI and *C. difficile-*co-colonized mice showed nominal differences in amine-containing carbon sources compared to *C. difficile*-monocolonized mice (Fig. 2D-E). PBI-specific Stickland aromatic amino acid metabolites predominated in the co-colonized state, in addition to reduced levels of *C. difficile-*specific histidine Stickland metabolites, suggesting a dominance of PBI’s Stickland metabolism (Figs. 2I, Supplemental Text). These findings illustrated CSAR’s capacity to create a nutrient enriched environment for *C. difficile* while PBI depleted preferred nutrients.

Microbial colonization altered additional host and microbial-origin metabolites. All three species enriched primary bile acids, including ones capable of inhibiting *C. difficile* germination, such as *β*-muricholate and chenodeoxycholate, and others with germination-stimulatory effects including cholate and taurocholate (Francis et al., 2013; Sorg and Sonenshein, 2009) (Fig 2A, right-hand side; SDF_3.17, Supplemental Text). PBI increased host-origin sphingosine-containing compounds (SDF_3.22), while PBI and CSAR each enriched compounds with host neuro-transmitter and anti-inflammatory properties such as ethanolamide endocannabinoids (Supplemental Text; SDF3.1, 3.4) (Lee et al., 2016). These molecules were further enriched with the severe mucosal damage from CSAR and *C. difficile* co-colonization.

### PBI and CSAR differentially modulate *C. difficile* gene expression *in vivo*

The commensal perturbations in luminal nutrients drove global perturbations in *C. difficile* gene expression (Figs. 3A-D, Fig. S2, SDF_4A-B). *C. difficile* in monocolonized mice enriched carbohydrate transport and metabolism systems for glucose, fructose, ribose and disaccharides (Figs. 3A, 3C, SDF_5.2, 5.5, 5.7-8, 5.12), dipeptides and oligopeptides (SDF_5.4), and Wood- Ljungdahl pathway genes for CO2 fixation to acetate (SDF_5.16). By 24h the pathogen up- regulated ethanolamine utilization genes (SDF_5.6), enabling capacity to use ethanolamine and amino-alcohol lipids from damaged mucosa (Nawrocki et al., 2018).

**Figure 3:**
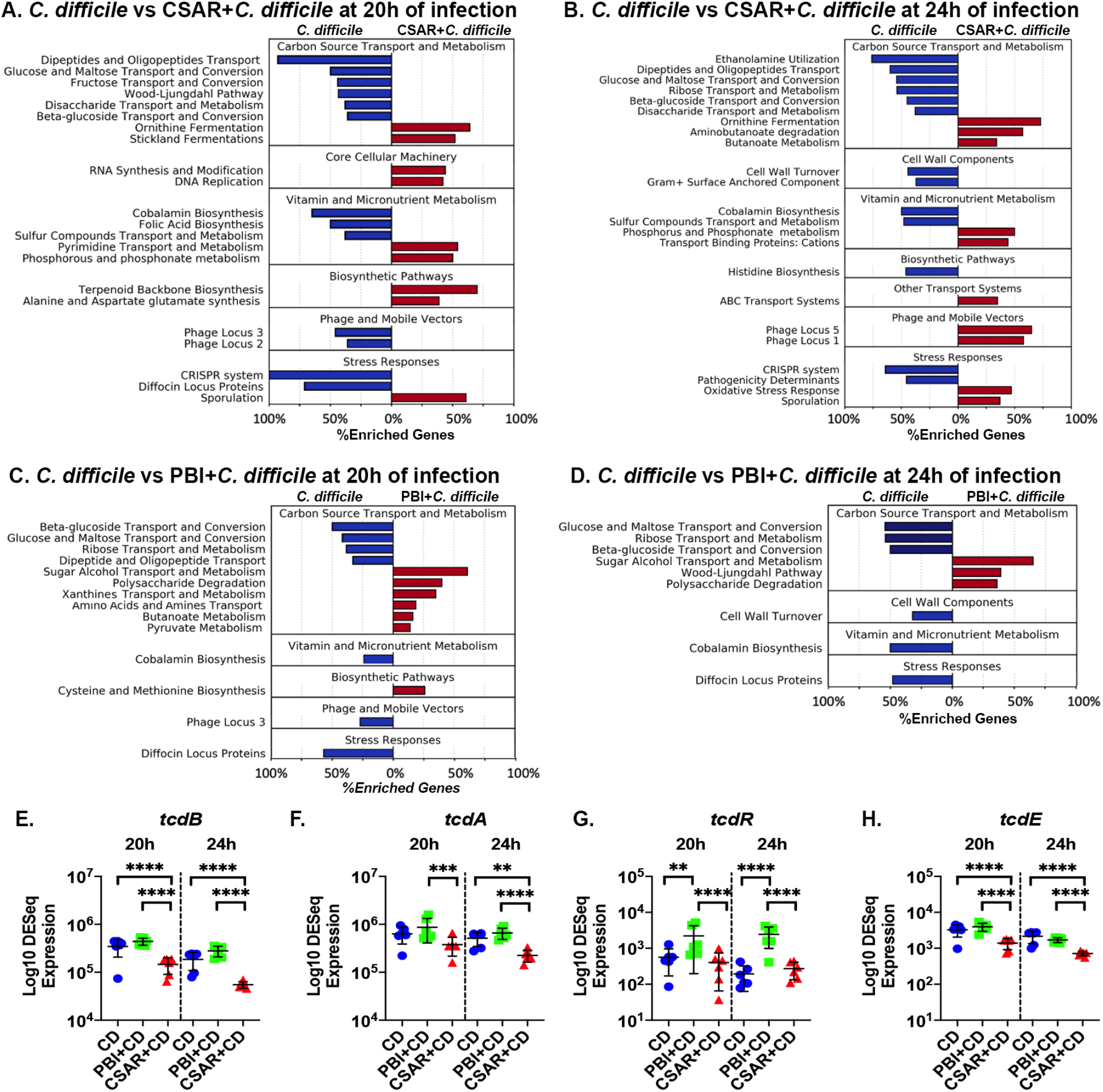
*C. difficile-*expressed pathways enriched *in vivo*. Horizontal bars indicate the percentage of genes within the pathway that were enriched in *C. difficile* monocolonized mice (LH side) or in mice co-colonized with CSAR or PBI prior to *C. difficile* infection (RH side). Pathways with a Benjamini-Hochberg adjusted p value≤0.05 are shown. **A:** *C. difficile* monocolonized mice at 20h of infection vs. CSAR and *C. difficile* co-colonized mice. **B:** Same comparison as panel A but at 24h of infection. **C.** Same comparison as in panel A at 20h of infection, comparing *C. difficile* monocolonized with PBI-co-colonized mice. **D:** Same comparison as panel D. but at 24h of infection. **Panels E-H:** DESeq-normalized reads for the PaLoc genes. X-axis indicates the colonization condition; Y-axis the Log10 DESeq normalized read counts that are normalized for biomass differences. Brackets indicate significant DESeq2 adjusted p values: *0.01<p*≤*0.05; **0.001<p*≤*0.01; ***0.0001<p*≤*0.001; ****p*≤*0.0001 **E.** *tcdA*, **F.** *tcdB*, **G.** *tcdR* and **H.** *tcdE*.

With the amino acid and polyamine enrichment from CSAR monocolonization, *C. difficile* up-regulated amino acid and polyamine transporters (SDF_5.15), pathways to convert CSAR- enriched ornithine to Stickland fermentable substrates (Fonknechten et al., 2009) (SDF_5.9), and its Stickland proline reductase and reductive leucine pathway genes (Fig. 4A-B; Supplemental Text; SDF_5.13).

**Figure 4:**
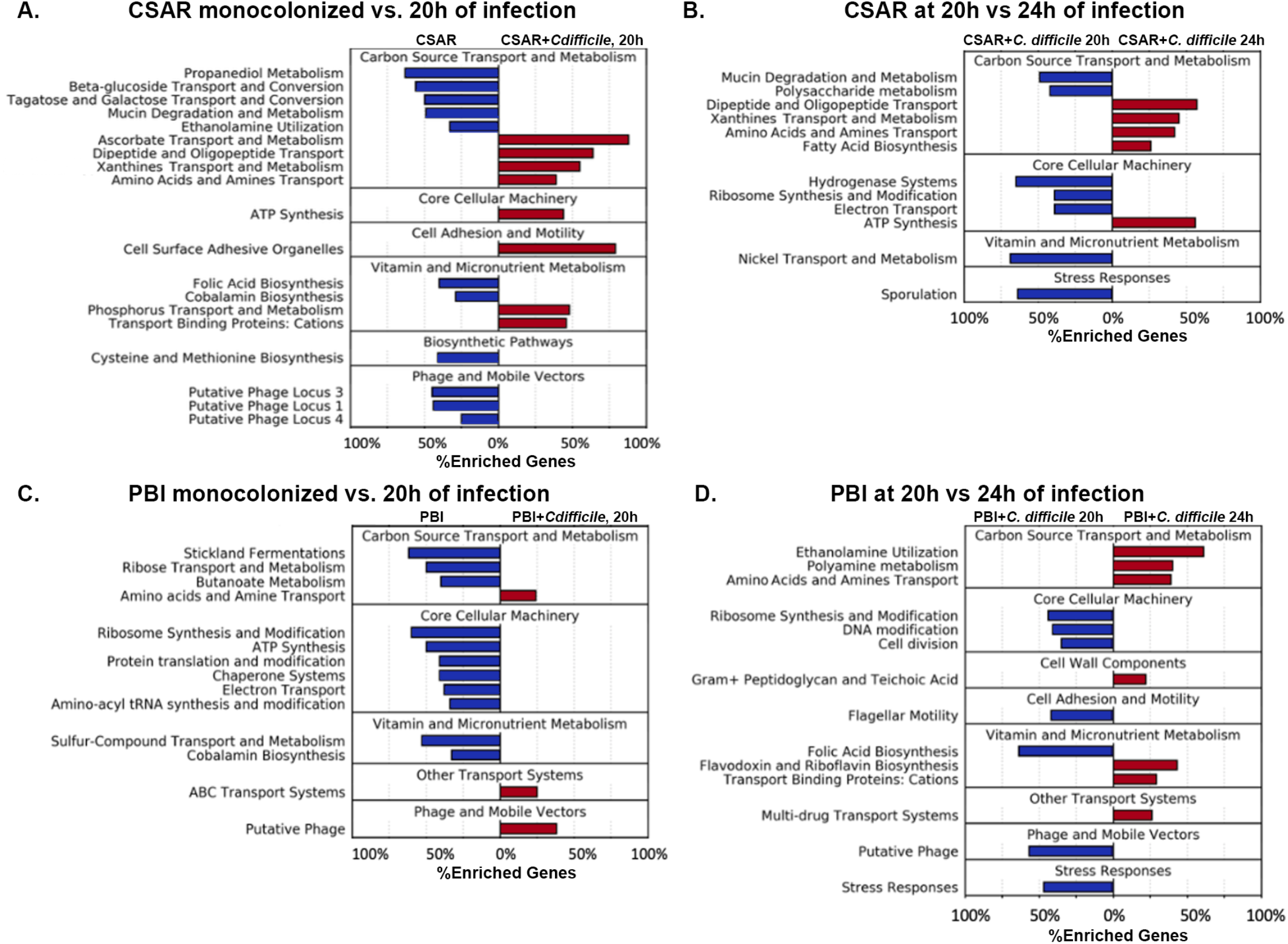
Commensal transcriptome analyses. Significantly enriched pathways in CSAR and PBI monocolonized mice and mice infected with *C. difficile*. Pathways with Benjamini-Hochberg adjusted p values ≤0.05 are shown. The left-hand Y-axis shows enriched pathways; X-axis indicates the percent of enriched genes per category. **A:** CSAR monocolonized mice at 7d prior to *C. difficile* infection vs. after 20h of *C. difficile* infection. **B:** CSAR and *C. difficile* co-colonized mice at 20h vs 24h of infection. **C:** PBI monocolonized mice at 7d vs after 20h of *C. difficile* infection. **D:** PBI and *C.* difficile co-colonized mice at 20h vs 24h of infection.

With PBI co-colonization (Fig. 3C-D), the pathogen adapted its metabolism to available nutrients, showing enrichment of genes to metabolize sugar alcohols including mannitol and sorbitol (SDF_5.14), disaccharides (SDF_5.5), and polysaccharides (SDF_5.10), carbon sources not utilized by PBI (Moore, 1993)*. C. difficile* also upregulated genes for the transport and metabolism of xanthines (Bradshaw, 1960) concomitant with enrichment of these compounds during PBI monocolonization (SDF_5.17).

In the presence of either commensal, *C. difficile* downregulated cobalamin (Figs. 3A-D; SDF_5.23), and folate biosynthesis genes when co-colonized with CSAR (Fig. 3A; SDF_5.24), suggesting cross-feeding of these nutrients with co-colonization.

Commensal colonization profoundly altered the pathogen’s cellular machinery. *C. difficile* genes associated with transcription and DNA replication were upregulated with CSAR co-colonization (Fig. 3A-B; SDF_5.18-5.19). In contrast, by 20h in PBI co-colonized mice, the pathogen significantly down-regulated translation, ribosome production, and ATP synthesis (Fig. 4C-D; SDF_5.20; 5.21).

With the host’s evolving inflammatory responses, commensal colonization altered *C. difficile’s* stress responses. By 20h of infection *C. difficile* monocolonized mice enriched expression of CRISPR genes (SDF_5.37) and two temperate bacteriophage loci (Arndt et al., 2019) with homology to phiMMP01 (locus 3; Fig. 3A; SDF_5.34), and phiCDHM19 (locus 2; Fig. 3A; SDF_5.33). Diffocin lytic genes (Gebhart et al., 2015), a phage-origin locus induced by quorum sensing that can lyse other *C. difficile* (Fig. 3A-D; SDF_5.37), and cell wall turnover enzymes (SDF_5.20) were also enriched. In CSAR-co-colonized mice, *C. difficile* sporulation pathways (SDF_5.40) and oxidative stress responses including nitroreductases and spore-associated superoxide dismutase (*sodA*), and catalase (*cotG)* were enriched (SDF_5.38), as were genes for terpenoid backbone synthesis (SDF_5.30), peptidoglycan and spore coat components with anti-oxidant activities (Bosak et al., 2008). Two additional, unlinked bacteriophage loci with homology to phiMMP03 (loci 1 and 5; Fig 3B; SDF_5.32, 5.35), separate from those enriched in monocolonized mice, were induced. In contrast, none of these systems were induced with PBI co-colonization (Fig. 3C-D).

Each commensal differentially affected *C. difficile’s* PaLoc expression which, in combination with alterations in pathogen biomass, impacted toxin levels and host disease (Fig 1I- J). The biomass-adjusted expression of *tcdB, tcdA* and *tcdE* in *C. difficile* monocolonized and PBI co-colonized mice remained comparable at 20 and 24h of infection (Figs. 3E-G), in spite of a 12-fold increase in *tcdR* expression in PBI-co-colonized mice (Fig. 3H). While CSAR co-colonized mice showed reduced *tcdB, tcdA* and *tcdE* expression at 20 and 24h of infection, these effects occurred in the context of higher pathogen vegetative biomass and released toxin in cecal contents (Fig. 1I-J).

### *C. difficile* infection profoundly alters commensal gene expression

*C. difficile* infection also systematically altered each commensal’s gene expression, metabolism and stress responses (Figs. 4A-D; SDF_6-7). In the monocolonized state, CSAR expressed systems involved in mucin degradation and metabolism (Fig 4A; SDF_6.5, Supplemental Text) and the biosynthesis of cobalamin and folate (SDF_6.16-17). Three phage loci, two with homology to *Clostridial* bacteriophage vB_CpeS_CP51 (loci 3 and 1; SDF_6.23-24), and one with homology to phiCT19406B (locus 4; SDF_6.25), were also enriched (Arndt et al., 2019).

By 20h of infection, concomitant with the mucosal damage inflicted by infection, CSAR profoundly downregulated its mucin degradation machinery (Fig 4A; SDF_6.5), while enriching expression of genes for ascorbate transport and metabolism (Fig 4A; SDF_6.1), systems that provide protection against oxidative stress. Multiple peptide and amino acid transport systems (SDF_6.3, 6.9), and genes involved in the transport and metabolism of xanthines, were enriched (SDF_6.10). By 24h CSAR had profoundly down-regulated ribosome production (SDF_6.14) and sporulation pathways (SDF_6.26). Energy generation and transport were also significantly affected as infection progressed, showing depletion by 24h of multiple electron transport systems (SDF_6.12) and CSAR’s nickel-based hydrogenase system (Fig 4B; SDFs_6.18, 6.13) while enriching expression of F-type ATP synthesis genes (Fig 4B; SDF_6.11).

In monocolonized mice PBI showed strong enrichment of its Stickland reductase systems (Fig 4C; SDF_7.5). Multiple systems involved in protein translation and export (SDFs_7.7, 7.10, 7.13-14), and energy generation (SDF_7.12) were also enriched. As with CSAR, PBI showed enrichment of its cobalamin synthesis genes (SDF_7.17). By 20h of infection, as mucosal damage developed, these processes showed reduced gene expression as compared to the monocolonized state (Fig 4C), with enrichment of multiple amino acid transport systems (SDF_7.6, 7.23) and a putative bacteriophage with homology to phiCT19406A (SDF_7.25) (Arndt et al., 2019). By 24h of infection, PBI enriched expression of genes for ethanolamine utilization (Fig 4D; SDF_ 7.2), suggesting a re-tooling of its metabolism for newly available carbon sources from damaged mucosa, in addition to genes for riboflavin transport and biosynthesis (SDF_7.19). Multiple stress response systems including *lexA, relA* and CSP-family cold shock proteins showed depletion by 24h of infection (SDF_7.26), in addition to genes involved in ribosome production, cell division and flagellar motility (Fig 4D; SDFs_7.14, 7.9, 7.15).

### Differential contributions of commensal arginine fermentation on *C. difficile* virulence

CSAR and PBI can each ferment arginine to ornithine via the arginine deiminase (ADI) pathway, a system that *C. difficile* lacks (Pols et al., 2020). However, *C. difficile* and other Stickland fermenters can convert ornithine to Stickland-fermentable substrates (Fonknechten et al., 2009). The species-level context of the ADI pathway demonstrated very different outcomes on *C. difficile* growth and host survival (Fig 5).

**Figure 5:**
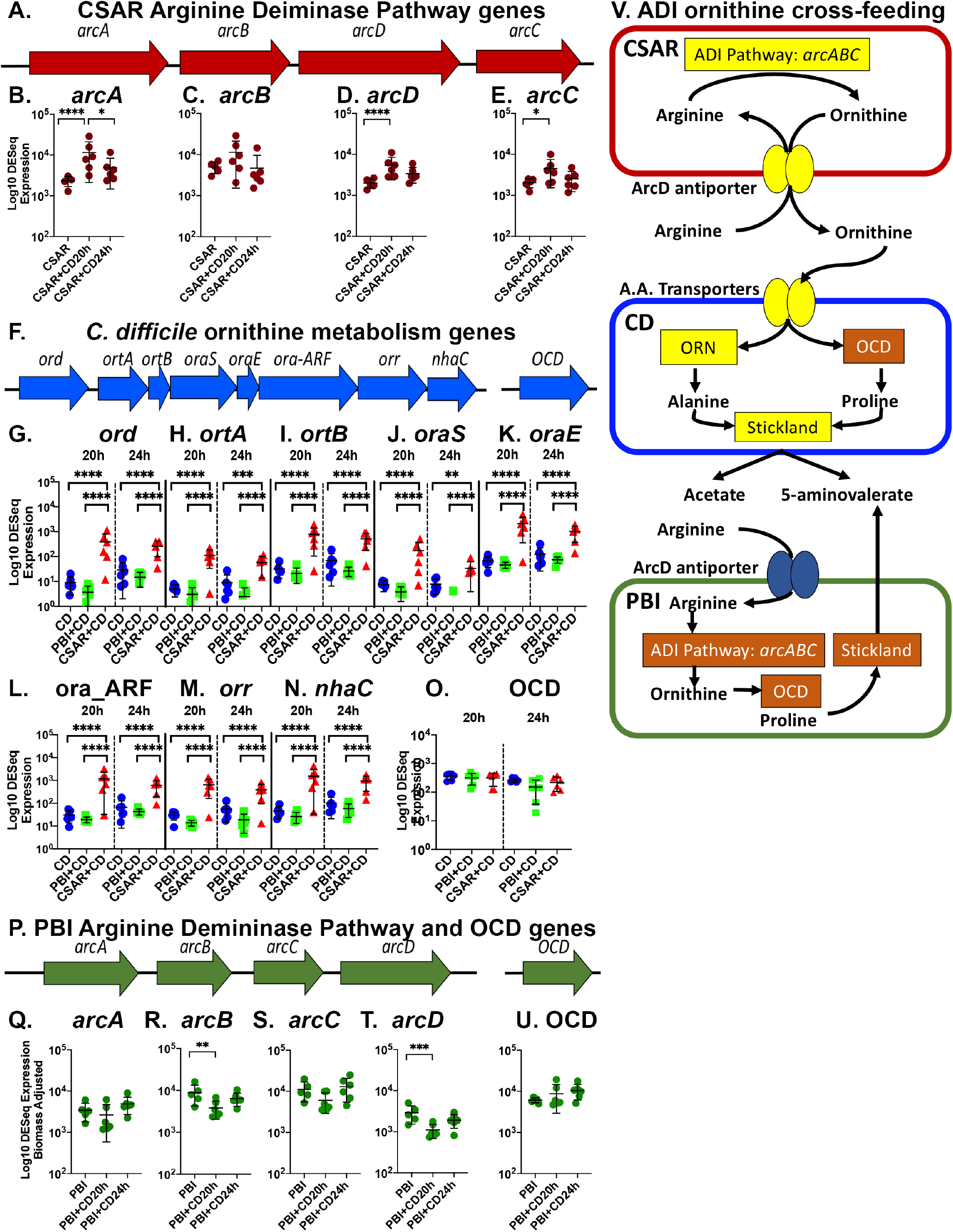
Commensal-pathogen ornithine cross-feeding. **A:** Schematic of CSAR’s arginine deiminase pathway genes. **Panels B-E:** Log10 DESeq2 normalized ADI gene expression. **B:** *arcA,* arginine deiminase gene (****p=0.0001; *p=0.0387), **C:** *arcB,* ornithine carbamoyltransferase, **D:** *arcD,* arginine:ornithine antiporter (****p=0.0008), and **E:** *arcC*, carbamate kinase gene (*p=0.0259). **F:** *C. difficile* ornithine to alanine conversion genes (operon_0246) and unlinked ornithine cyclodeaminase gene (GeneIDs: fig|1496.5150.peg.3209; UAB_RS0203800). **Panels G-O:** DESeq2 normalized expression for operon_0246 genes,*0.01≤p<0.05; **0.001≤p<0.01; ***0.0001<p*≤*0.001; ****p*≤*0.0001. **G:** *ord*, 2,4-diaminopentanoate dehydrogenase, **H:** *ortA,* 2-amino-4-ketopentanoate thiolase subunit alpha, **I:** *ortB,* 2-amino-4- ketopentanoate thiolase subunit beta, **J:** *oraS,* D-ornithine aminomutase S component, **K:** *oraE,* D-ornithine aminomutase E component, **L:** ora_ARF, reactivating factor for adenosylcobalamine- dependent D-ornithine aminomutase, **M:** *orr*, ornithine racemace, **N:** *nhaC,* Na+/H+ antiporter. **O:** ornithine cyclodeaminase (OCD) expression. **P:** Schematic of PBI arginine deiminase pathway and OCD genes. **Q:** *arcA, arginine deiminase,* **R:** *arcB,* ornithine carbamoyltransferase, **S:** *arcC*, carbamate kinase **T:** *arcD,* arginine:ornithine antiporter (*p=0.0227), and **U:** OCD. **V:** Schematic showing differing effects of the ADI arginine fermentation in CSAR and CBI on *C. difficile* metabolism.

By 20h of infection, CSAR upregulated expression of its a*rcA* arginine deiminase (Fig. 5B), *arcC* carbamate kinase (Fig. 2E), and *arcD* arginine:ornithine antiporter (Fig. 2D) which imports arginine and exports the ornithine product, identifying a cause for the elevated ornithine in CSAR monocolonized mice (Fig. 2E). In *C. difficile* the most strongly enriched genes with CSAR co-colonization converted ornithine to alanine (Fig 5F-N), a product able to support oxidative Stickland metabolism, cell wall synthesis, and other growth-promoting pathways (Peltier et al., 2011; Shrestha et al., 2017). Notably, *C. difficile’s* OraSE D-ornithine aminomutase (Fig. 5J-K) requires cobalamin (Chen et al., 2001), a factor which also demonstrated enriched gene expression for its biosynthesis in CSAR monocolonized mice (Fig. 4A). The pathogen’s ornithine cyclodeaminase (Fig. 5F) which converts ornithine to the Stickland acceptor proline remained constitutively expressed (Fig. 5O).

In contrast to CSAR, at 20h of infection PBI down-regulated its *arcD* arginine:ornithine antiporter (Fig. 5P, 5T) while maintaining constitutive expression of its ornithine cyclodeaminase (Fig. 5U) conserving a proline-convertible carbon source for its Stickland metabolism (Figs. 5D; SDF_7.18).

Figure 5V illustrates the differing inter-species effects of ADI fermentation on *C. difficile in vivo*. While the non-Stickland fermenter CSAR enriched luminal ornithine for *C. difficile’s* use, PBI’s combined ADI and Stickland pathways support its own metabolism and growth, depriving *C. difficile* of these growth-promoting carbon sources.

Systems biologic models predict epistatic effects *of C. difficile’s* CodY and CcpA PaLoc metabolic repressors on pathogen phenotypes *in vivo*.

CcpA and CodY repress *C. difficile* toxin expression when sufficient GTP, Stickland- fermentable substrates and carbohydrates support metabolism (Dubois et al., 2016). Their absence contributes to toxin gene de-repression. PRIME model predictions inferred additional epistatic effects of *codY* and *ccpA* on the pathogen’s ability to colonize and grow *in vivo* (Arrieta-Ortiz ML, 2020), factors required for infection. Using new genetic manipulation systems supporting the creation of serial deletion mutants in *C. difficile* (Star Methods) we validated effects of these regulators in the gnotobiotic and PBI-co-colonized states (Fig. 6)

**Figure 6:**
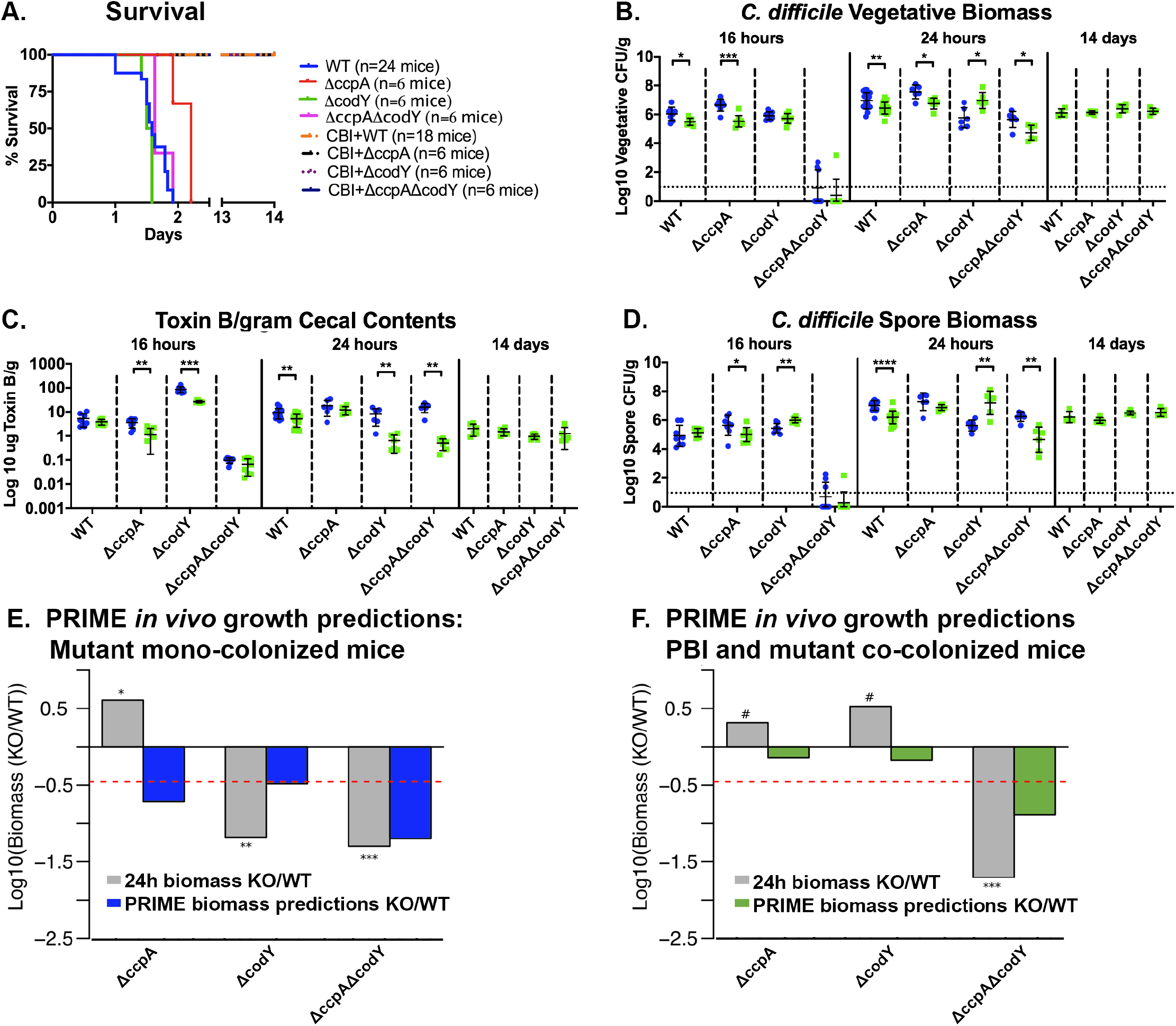
Combinatorial effects of *C. difficile’s* CodY and CcpA PaLoc metabolic repressors on the *in vivo* dynamics of infection in mono- and PBI-co-colonzied mice. **A.** Survival curves of GF and PBI co-colonized mice infected with *C. difficile* wild-type (WT) or mutant strains; n=at least 6 mice/group. The curves for PBI co-colonized mice overlap and were significantly different from monocolonized controls (p<0.0001). *ΔcodY-*infected mice declined more rapidly than WT-infected mice (p=0.01), while lethality in *ΔccpA*-infected mice was delayed (p=0.0002). **B-D:** Cecal biomass and extracellular toxin B levels. Blue: *C. difficile-*associated mice; Green: mice monocolonized with PBI and infected with *C. difficile*. Asterisks indicate significance values by Mann-Whitney log rank test: *0.01<p≤0.05; **0.001<p≤0.01; *** 0.0001<p≤0.001; ****p≤0.0001. **B:** Log10 of ug toxin B levels per gram of cecal contents. **C:** Cecal Log10 of *C. difficile* vegetative cells and **D:** spores at 16h, 24h, and at 14d in surviving PBI co-colonized mice. **E-F:** PRIME model predictions of pathogen growth in monocolonized (blue bars; E) and PBI co-colonized mice (green bars; F). Dashed red lines indicates threshold for regulator essentiality in supporting *in vivo* growth relative to the wild-type strain. # symbol indicates non-significant t-test p values ≥ 0.05. The ‘*’, ‘**’ and ‘***’ symbols indicate p values < 0.05, < 0.01 and < 0.001, respectively.

*ΔcodY*, *ΔccpA,* and double *ΔcodYΔccpA C. difficile* mutants were each lethal in monocolonized mice while PBI-co-colonized mice survived (Figs. 6A, S2A-C). Pathogen biomass and toxin studies in cecal contents identified PBI’s effects on toxin production among knockout strains (Figs. 6B-D, S2D-F). PBI-co-colonized mice infected with the *ΔccpA* mutant demonstrated reduced *C. difficile* vegetative biomass and toxin levels at 16h (Figs. 6B-C, S2D-E). At 24h of infection, toxin levels were comparable between colonization states with the mutant and fell >80% by 14 days in surviving PBI-co-colonized mice (Fig. 6C).

In contrast, *ΔcodY*-infected mice demonstrated better growth with PBI-co-colonization, while the double mutant grew poorly (Figs 6B, S2D). At 16h of infection the *ΔcodY* mutant showed comparable vegetative biomass in monocolonized and co-colonized states, and elevated spore biomass in PBI-co-colonized mice (Fig. 6D). Toxin levels with the *ΔcodY* mutant were elevated at 16h as compared to wild-type controls, though levels in PBI-co-colonized mice were 70% lower than in *ΔcodY* monocolonized controls (Figs. 6C, S2E). However, from 16h to 24h, toxin levels fell >40-fold in PBI-co-colonized mice (Figs. 6D, S2E), in spite of the *ΔcodY* mutant’s higher vegetative biomass at 24h in co-colonized mice (Figs. 6B, S2D).

PRIME model predictions inferred dysregulation of multiple gene networks controlling central carbon metabolism, lipid and nucleotide biosynthesis in *C. difficile* mono-colonized mice (Arrieta-Ortiz ML, 2020). In contrast, in the nutrient-depleted conditions in PBI co-colonized mice (Fig 6F), PRIME predicted that single deletion of CodY and CcpA were non-essential in promoting growth, while lack of both regulators severely limited growth. In the PBI-co-colonized mice, growth limitation of the double knockout strain was driven by combinatorial effects of both regulators on essential genes in lipid, cell wall and nucleotide biosynthesis, and dysregulation of genes supporting the metabolism of gut-available carbon sources in the PBI co-colonized state, namely sugar alcohols, polysaccharides, and supporting electron transport systems (Arrieta-Ortiz ML, 2020). These findings illustrated PBI’s continued protective effects in the absence of the PaLoc CodY and CcpA metabolic repressors per the double-knockout’s inability to metabolize available carbon sources and adapt its cellular systems to support active growth.

### PBI bacteriotherapy rescues infected conventional mice

To assess PBI’s use as a therapeutic, clindamycin-treated conventional mice were orally challenged with 1,000 *C. difficile* spores. At the onset of symptomatic infection mice received 10^8^ CFU of PBI or vehicle-only control by gavage (Fig. 7A). All PBI-treated mice survived while control-treated mice demonstrated 40% lethality (Figs. 7B, p=0.0061; S3A-C). At 30h post-*C. difficile* challenge, at the height of symptomatic infection, PBI-treated mice demonstrated reduced toxin levels and pathogen vegetative and spore biomass (Fig. 7C-E). By 14 days surviving mice had low to undetectable toxin (Fig 7C) and had largely cleared both species (Figs. 7D-E; S3C).

**Figure 7:**
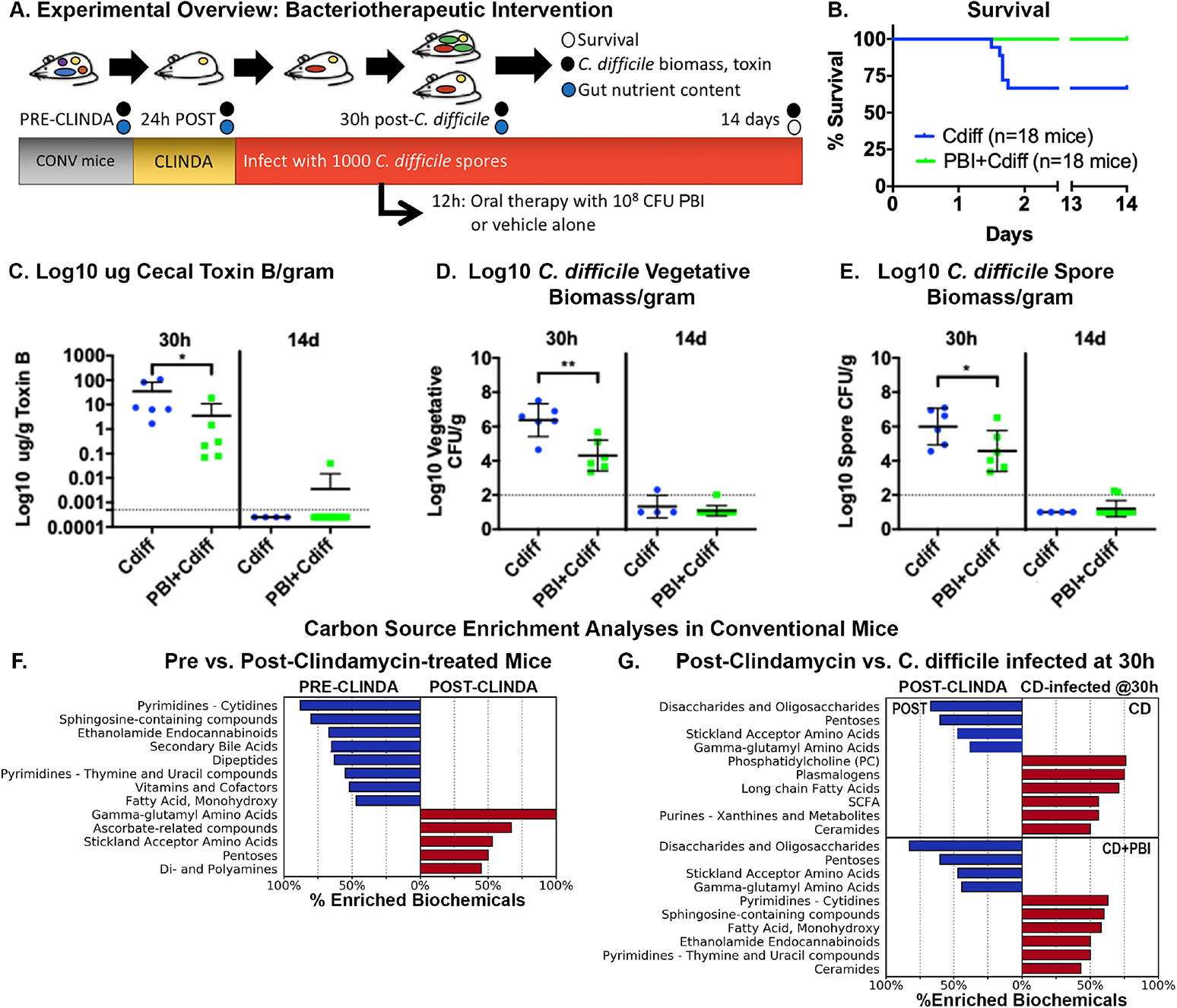
PBI oral bacteriotherapy protects *C. difficile* infected conventional mice. **A:** Experimental overview. Samples for timed analyses (circles) were taken before and after clindamycin, at 30h post-infection (12h post-treatment), and at 14d in surviving mice. **B:** Survival curve; blue: *C. difficile*-infected and vehicle control treated; green: PBI treated mice showed improved survival (p=0.0081). **C-E:** Cecal toxin and *C. difficile* biomass. Horizontal dotted line shows thresholds of detection. **C:** Log10 ug Toxin B per gram of cecal contents (*p=0.026). **D:** Log10 *C. difficile* vegetative (**p=0.0087) and **E:** spore biomass in cecal contents (p=*0.0411). **F:** Carbon source enrichment analyses in mice pre- and post-clindamycin treatment showing enriched groups with a Benjamini-Hochberg corrected p value ≤0.05. **G:** Enriched carbon source groups between post-clindamycin treated and mice at 30 hours of infection with *C. difficile* (top) or with PBI treatment (bottom).

Clindamycin treatment enriched multiple carbon sources capable of supporting *C. difficile* and PBI growth including Stickland acceptor amino acids, polyamines, and *γ*-glutamyl-amino acids (Fig. 7F-G, SDF_10.4, 10.19, 10.8), nutrients also enriched in CSAR-monocolonized mice (Fig. 2A). In control infected mice, the pathogen depleted oligosaccharides, pentoses, Stickland acceptor- and *γ*-glutamyl-amino acids (Fig. 7G, top; SDF_10.4, 10.9, 10.19, 10.7). With PBI- treatment, these and additional compounds were depleted (Fig. 7G, bottom; SDF_10.4, 10.7). Clindamycin treatment also depleted host ethanolamine endocannabinoids and sphingosines, compounds with improved recovery in PBI-treated mice (Fig. 7G; SDF_10.5, 10.18).

These findings validated PBI’s therapeutic efficacy in an infected conventional host with specific reductions in pathogen biomass and toxin levels.

## Discussion

We demonstrate the application of systems biologic approaches to define how individual commensals modulate *C. difficile’s* virulence *in vivo*. Using a tractable gnotobiotic infection model we identified the remarkably protective effects of a single commensal species, *P. bifermentans,* against *C. difficile,* and capacity for another *Clostridial* species, *C. sardiniense,* to promote more severe disease. Findings informed interventional studies in conventional, antibiotic-treated mice where PBI administration rescued infected mice from lethal infection. These findings have important implications to treat and prevent *C. difficile* infections, including the fact that residual microbiota and those in uncharacterized FMT preparations can contain both protective and disease-exacerbating species that may exhibit different behaviors near- and longer-term in antibiotic-depleted versus intact microbiota.

Development and severity of *C. difficile* infection occurs as a function of the pathogen’s biomass, toxin production, and duration to which host tissues are exposed to toxin. Each commensal modulated *C. difficile*’s virulence through multiple mechanisms (graphical abstract). Commensal colonization altered the gut nutrient environment per enrichment or depletion of *C. difficile-*preferred carbon sources and required micronutrients, including cobalamin and folate. These factors modulated the pathogen’s growth, stress responses, and toxin production.

Commensal colonization also affected pathways impacting *C. difficile’s* cellular integrity and toxin release. *C. difficile* monocolonized mice up-regulated diffocins, a locus induced through quorum-sensing mechanisms that lyses a portion of the population, releasing toxin to promote host damage and nutrient release to support surviving populations (Gebhart et al., 2015). Multiple temperate bacteriophage harboring lytic peptidoglycan hydrolases were induced in monocolonized and CSAR co-colonized mice (Garneau et al., 2018). Sporulation, strongly enriched in CSAR co-colonization, also induces mother cell lysis. These mechanisms may enable abrupt toxin release through cell wall disruption, processes known to occur in other toxigenic species including *C. perfringens, Shigella* and ETEC *(Bielaszewska et al., 2012; Duncan, 1973; El Meouche and Peltier, 2018)*.

In response to CSAR’s enrichment of amine-containing carbon sources *C. difficile* up- regulated multiple amino acid transporters, Stickland fermentation pathways, and genes to convert CSAR enriched ornithine to Stickland metabolizable substrates. Resulting biomass of *C. difficile* and CSAR expanded and, with nutrient release from damaged tissues, further stimulated microbial growth and toxin production, resulting in a rapidly lethal infection.

Notably, CSAR and PBI possess arginine deiminase fermentation pathways, a system that modulated very different effects *in vivo* with *C. difficile* infection (Fig. 5E). With CSAR, the commensal’s export of ornithine provided a new nutrient source for *C. difficile* metabolism and growth. In contrast, PBI’s capacity to convert ornithine to proline supported its own Stickland metabolism, depriving the pathogen of this nutrient source, while potentially enhancing its competitive advantage over *C. difficile*. This example highlights the importance of the underlying genomic and metabolomic context of pathways shared among commensal species when considering their effects on other microbes and on host phenotypes.

PBI limited *C. difficile’s* growth and toxin production through multiple mechanisms including in the absence of *C. difficile’s* CodY and CcpA PaLoc repressors. As an active Stickland fermenter, PBI depleted fructose and amino acids preferred by *C. difficile,* leaving sugar alcohols, complex polysaccharides, and PBI-enriched hypoxanthines available. *C. difficile* adjusted its metabolism for these carbon sources but by 24h of infection showed reduced biomass and toxin as compared with monocolonized and CSAR-co-colonized mice. PBI colonization also down- regulated genes supporting translation and ribosome production in *C. difficile* in addition to the pathogen’s cellular lytic systems and oxidative stress pathways, providing additional mechanisms by which PBI reduced the host’s exposure to toxin (graphical abstract). PRIME model predictions inferred additional combinatorial effects of CodY and CcpA on multiple cellular systems involving metabolism, electron transport, and biosynthetic reactions needed to support growth.

All three microbial species induced oxidative stress systems with symptomatic infection. The up-regulation of sporulation responses by *C. difficile* and CSAR when co-colonized, including the spore coat proteins superoxide dismutase (*sodA*) and manganese catalase (*cotG*), illustrated how sporulation in the confined space of the gut lumen may benefit vegetative cell populations of the same and other species by detoxifying host-produced ROS. Notably, PBI is a microaerophilic species, able to tolerate up to 8% O2, concentrations at which CSAR and *C. difficile* cannot survive (Katarzyna Leja, 2014), a factor enabling its competition against *C. difficile* during host ROS responses. PBI colonization also reduced the severity of the host’s acute inflammatory responses (Fig. 1E, graphical abstract), in part from reduced host exposure to high levels of *C. difficile* toxin (Fig 1I).

Interventional studies in antibiotic-treated conventional mice illustrated PBI’s efficacy as an oral bacteriotherapeutic. Clindamycin treatment enriched multiple amino acid and carbohydrate sources, including ones enriched in GF and CSAR monocolonized mice, supporting relevance of findings from germfree infection studies to complex microbiota. These findings support a broader systems-level view for perturbations that create nutrient states conducive to *C. difficile* colonization and rapid growth *in vivo*, particularly given the pathogen’s diverse carbon source metabolism and adaption to different gut environments. Notably, nutrient conditions enhancing *C. difficile* growth can also enhance the growth of commensal Stickland fermenters. However, disease-triggering antimicrobials to which *C. difficile* often harbors innate resistance, such as clindamycin, beta-lactam and fluoroquinolones, rapidly ablate commensal Stickland fermenters, opening new nutrient conditions in the absence of key competitors (Gao and Huang, 2015).

Within the colon, unabsorbed dietary components and host factors, mucins in particular, provide fermentable carbohydrates and amino acids, including the Stickland acceptors proline, glycine and leucine (Wesley et al., 1985). In combination, these amino acids can enable rapid pathogen growth (Battaglioli et al., 2018), with increased risks for population crashes that lead to abrupt spikes in toxin release.

Stickland fermentation is a hallmark of Cluster XI *Clostridial* metabolism, and occurs in other *Clostridial* species, notably *Clostridium scindens* (Cluster XIVa) which carries the proline and glycine reductases but not the reductive leucine pathway (Fig. S4). While conversion of progermination primary bile acids to germination-inhibitory secondary bile acids by *C. scindens* has been hypothesized to mediate *in vivo* protection against *C. difficile (Thanissery et al., 2017)* we show more fundamental capacity for commensals, singly or in aggregate, to rapidly change the gut nutrient environment to modulate *C. difficile’s* virulence. Notably, the PBI and CSAR strains used did not demonstrate 7*α*-hydroxysterol-dehydrogenase activity in mice (Supplemental Text), further emphasizing their modulation of disease outcomes independently of microbial bile salt transformations.

Stickland fermenting species represent <1% of the human gut microbiota. Our findings highlight the importance of these low-abundance members to consume growth-promoting nutrients for *C. difficile* and identify conditions that could be created by other commensals, singly or in aggregate, to modulate *C. difficile’s* virulence. These conditions act in concert with the host’s digestive and immune functioning. Host and commensal effects may also explain why *C. difficile* phenotypes identified *in vitro* do not necessarily reflect behaviors seen *in vivo.* Armed with refined mechanistic knowledge, findings establish a robust framework in which to develop therapeutics with enhanced efficacy and improved safety for this disease.

## Supporting information

Supplemental Text File (SDF1)

## Acknowledgments

We thank Andrea Dubois, Cameron Friedman, Vladimir Yeliseyev, Qing Liu, Rebecca Krinzman and Teri Bowman for technical support, and Peter Sewell for graphic design support. RNA sequencing was performed by the Harvard Medical School Biopolymers Core. Partners Healthcare’s ERIS team supported the Massachusetts Host-Microbiome Center’s high performance computing resources. We would also like to thank Jessica Allegretti, Laurent Bouillaut, Laurie Comstock and Aimee Shen for critical reading of the manuscript and helpful comments. **Funding:** This work was supported by the BWH Precision Medicine Institute, Harvard Digestive Diseases Center grant P30 DK034854, and R01 AI153605 (LB). BP was supported by T32 HL007627; the work of JW was supported by the Intramural Research Program of the National Library of Medicine, National Institutes of Health. JP received support from the Institut Pasteur (Bourse ROUX).

## Author Contributions

The following author contributions were made. Conceptualization: LB, AS, GG. Methodology: BP, ND, JW, JP, MLAO, RI, MD, MH, YL, NG, MA, AO, CC, GG, AS, NB, BD, LB. Software: MLAO, RI, NB, JW, LB. Formal Analysis: BP, ND, JW, MLAO, RI, RL, MD, MH, YL, NG, MA, NB, LB. Investigation: BP, ND, JW, JP, MLAO, RI, RL, MD, AS, NB, BD, LB. Resources: BP, ND, JP, MLAO, RI, RL, MD, MH, YL, NG, MA, AO, BD, AS, NB, LB. Data Curation: JW, MLAO, RI, NB, LB. Writing: LB, BP, ND, JW, JP, MD, MH, NG. Review and Editing: BP, ND, JW, MLAO, RI, JP, MD, MH, MA, AO, GG, AS, NB, BD, LB. Visualization: BP, ND, JW, MLAO, RI, NB, LB. Supervision: LB, BD, NB, MA

## Competing interests

LB and GG are co-inventors on patents for *C. difficile* microbiota therapeutics. LB, GG and AS are SAB members and hold stock in ParetoBio. GG is an SAB member and holds stock in Kaleido, Inc. AS is a co-owner of ExArca Pharmaceuticals, LLC. Remaining authors declare no competing interests.

### Data and materials availability

Metabolomic and transcriptomic datasets will be available from NCBI Bioproject PRJNA278886, and from the supplementary materials.

## STAR METHODS

### Bacterial strains and culture conditions

Table S1 shows the bacterial strains and *in vitro* culture conditions. For quantitation of *C. difficile* and commensal biomass, mouse cecal contents were collected into pre-weighed Eppendorf tubes with 0.5mL of pre-reduced PBS with 40mM cysteine (Sigma Chemical, St. Louis, MO) as a reducing agent. Tubes were weighed after adding material and transferred into a Coy anaerobic chamber (Coy Labs, Grass Lake, MI) at 37°C for serial dilutions with plating to selective *C. difficile* CHROMID agar (Biomérieux, Durham, NC) or Brucella agar (Becton Dickinson, Canaan, CT) for commensal quantitation*. C. difficile* colonies were counted at 48 hours of incubation and identified as large black colonies. For the *Δ*codY *Δ*ccpA double mutant, colonies were quantitated at 72 hours of incubation. Commensal colonies were counted after 24h of incubation. CSAR were identified as small, round beta-hemolytic colonies, and PBI as opaque and larger round colonies. Representative colonies were species-confirmed by rapid ANA (Remel, Lenexa, KS). For studies in conventional mice, pre-infection and post-clindamycin fecal pellets showed no positive colonies on CHROMID agar.

*C. difficile* spore preparations and counts were defined by exposing pre-weighed material to 50% ethanol for 60 minutes followed by serial dilution and plating to *C. difficile* CHROMID agar, as described (Bucci et al., 2016). Vegetative cell biomass was calculated by subtracting the spore biomass from the total biomass and normalizing to the cecal mass. Data were evaluated in Prism 8.0 (GraphPad, San Diego, CA) for visualization and log-rank tests of significance among groups. A p value <0.05 was considered significant.

### Construction of *C. difficile codY, ccpA* and double *codY ccpA* mutant strains

Table S2 indicates plasmid vectors and primer sequences used to generate gene-deleted mutants in ATCC43255. Mutants were created using the toxin-mediated Allele-Coupled Exchange (ACE) vector (Girinathan BP, 2020). For deletions, allelic exchange cassettes were designed to have approximately 900 bp of homology to the chromosomal sequence in both up- and downstream locations of the sequence to be altered. The homology arms were amplified by PCR from *C. difficile* strain ATCC43255 genomic DNA (Table S1) and purified PCR products were cloned into the PmeI site of pMSR0 using NEBuilder’s HiFi DNA Assembly. pMSR0-derived plasmids were transformed into *E. coli* strain NEB10β and inserts verified by sequencing. Plasmids were then transformed into *E. coli* HB101 (RP4) and transferred by conjugation into *C. difficile* ATCC43255 after a brief period of heat shock as described (Kirk and Fagan, 2016).

### Mouse Studies

All animal studies were conducted under an approved institutional IACUC protocol. Defined- colonization experiments were conducted in negative pressure BL-2 gnotobiotic isolators (Class Biologically Clean, Madison, WI). Conventional studies were conducted in OptiMice containment cages (Animal Care Systems, Centennial, CO). Mice were singly housed for all studies.

### Gnotobiotic Mouse Colonization and Infection Studies

One week prior to infection with *C. difficile* equal ratios of 6-7 week old male and female gnotobiotic mice were gavaged with 1x10^8^ CFU of *P. bifermentans* (PBI), *C. sardiniense* (CSAR), or sterile vehicle control, and allowed to colonize for 7 days prior to challenge with 1x10^3^ of wild- type or mutant *C. difficile* spores. Fecal pellets from mice were cultured prior to infection to confirm association with the defined species, or maintenance of the GF state. Progression of disease was assessed via body condition scoring (Fekete et al., 1996) and body mass measurements taken by ethylene-oxide sterilized, battery powered OHAUS scales (Thermo-Fisher, Waltham, MA). Mice were sacrificed at a BCS of 2-, or at defined timepoints at 7 days of commensal monocolonization or GF controls, and at 20, 24h or 14 days post*-C. difficile* challenge. For *C. difficile* mutant infection studies, timepoints at 16h and 24h post-challenge was collected. Cecal contents were collected for functional studies. The GI tract and internal organs were fixed in zinc-buffered formalin (Z-FIX, Thermo-Fisher, Waltham, MA) for histopathologic assessment.

### Conventional Mouse Infection Studies

5-week old conventional mice (Taconic Farms, Inc., Taconic, NY) were singly housed and acclimated for a week prior to treatment with USP-grade clindamycin phosphate (10mg/kg; Sigma Chemical, St. Louis, MO) via intraperitoneal (IP) injection. 24 hours post-clindamycin treatment, mice were challenged with 1x10^3^ wild-type *C. difficile* spores via oral gavage and treated with 1x10^8^ CFU of *P. bifermentans* (PBI) or vehicle control at 12h post *C. difficile* challenge, the earliest point of symptomatic diarrhea in conventional mice. Progression of disease was assessed via BCS and body mass measurements. Survival studies were followed to 14 days post *C. difficile* challenge. For *C. difficile* biomass, toxin B levels and cecal metabolomic studies, 12 mice per group were also sacrificed and cecal contents collected at pre-clindamycin treatment, post-clindamycin treatment just prior to *C. difficile* challenge, 30 hours *post C. difficile* challenge, and at 14 days following control or PBI treatment.

### Histopathologic Analyses

Formalin-fixed gut segments from GF or specifically-associated mice were paraffin embedded and 5μm sections cut for staining with hematoxylin and eosin (H&E; Thermo-Fisher, Waltham, MA) as described (Bry and Brenner, 2004). Slides were visualized under a Nikon Eclipse E600 microscope (Nikon, Melville, NY) to assess epithelial damage, per cellular stranding and vacuolation, the nature of Inflammatory infiltrates, mucosal erosions, and tissue edema.

### Toxin B ELISA

Cecal toxin B levels were quantified as described (Zarandi et al., 2017). Briefly, microtiter plates were coated with 5 μg/mL of anti-TcdB capture antibody (BBI solutions, Madison, WI). Supernatants of spun cecal contents and standard curve controls of toxin B (Campbell, CA) were assayed in triplicate. After incubation and washing with anti-toxin B biotinylated antibody (mouse- anti-*C.difficile* TcdB; (BBI solutions, Madison, WI) followed by high Sensitivity Streptavidin-HRP conjugate (Thermo-Fisher, Waltham, MA), signal was detected with TMB substrate (Thermo- Fisher, Waltham, MA) at 450nm using a BioTek Synergy H1 plate reader (Biotek Instruments Inc, Winoski, VT). Values were analyzed in Prism 8.0 (GraphPad, San Diego, CA) to calculate μg of toxin B/gram of cecal contents. Significant differences among groups were evaluated by non-parametric Kruskal-Wallis ANOVA and Dunn’s post-test. A p value ≤0.05 was considered significant.

### Effects of Commensal Colonization on Toxin Function

The Quidel *C. difficile* cell culture functional toxin assay (Beck et al., 2014) was used to evaluate if commensal colonization altered the functional toxicity of *C. difficile* toxin. Cecal contents were collected from germfree mice or from mice monocolonized for 7 days with *P. bifermentans, or C. sardiniense*. 100μL of purified toxin B control solution (Quidel Inc., San Diego, CA) was added to 1 gram of cecal contents and incubated for 30 minutes prior to making 1:10 to 1:500 serial dilutions in the Quidel-provided dilution buffer and adding materials to confluent cultures of human MRC- 5 fibroblasts. Fibroblast cells were incubated at 37°C for 48 hours and checked daily by compound microscope for signs of cytopathic effect (CPE) indicated by balling up of cells and loss of adhesion. Additional control samples included cecal contents incubated with toxin B for 30 minutes followed by addition of neutralizing antibody to confirm specificity of CPE by toxin B. Cells where CPE occurred in the presence of toxin B, but not with cecal contents alone or with neutralizing antibody were called positive. All conditions were repeated in triplicate. The highest dilution at which CPE occurred was identified for each condition.

### Western blot for Toxin B Integrity

Cecal supernatants from mice at 20h of infection were subjected to SDS-PAGE and transferred to PVDF membrane (PerkinElmer, Waltham, MA) as described (Girinathan et al., 2018). Toxin B was detected by using Sheep Primary Antibody, Donkey Secondary Antibody (R&D Systems, Minneapolis MN), diluted 1:1000 in 5% nonfat dry milk blotting buffer (25mM Tris, pH 7.4, 0.15M NaCl, 0.1% Tween 20), and by chemiluminescence using the SuperSignal West Pico Plus Western Blotting Substrate (part# 34577; Thermo-Scientific, Waltham, MA).

### Metabolomic studies

For GF colonization studies cecal contents from 8 mice per group across 2 experimental replicates were harvested from GF mice at baseline, after 7 days of monocolonization with PBI or CSAR, and at 20h post-infection with *C. difficile* alone or with each commensal (6 groups, 48 mice total). For conventional studies, cecal contents were collected from 12 mice per group prior to clindamycin treatment, 24h post-clindamycin treatment, and at 30h post *C. difficile* challenge, at the height of symptomatic infection. Materials were snap frozen into pre-weighed tubes and weighed to determine mass of cecal contents. Global metabolomic screen of samples was performed by Metabolon (Raleigh, NC) with sample extraction and MALDI-TOF analyses as described (Fletcher et al., 2018; Ryals et al., 2007). Results were obtained as Original Scale mass spectrometry counts.

### Cecal short chain fatty acid (SCFA) measurements

Volatile short chain fatty acids from specifically-associated mice were quantified as described (Moore, 1993). In brief, acidified internal standards with 100 µL of ethyl ether anhydrous or boron trifluoride-methanol was added to 100μl of supernatant from homogenized cecal contents. Chromatographic analyses were carried out on an Agilent 7890B system with flame ionization detector (FID). Chromatogram and data integration were carried out using the OpenLab ChemStation software (Agilent Technologies, Santa Clara, CA). SCFA in samples were identified by comparing their specific retention times relative to the retention time in the standard mix. Concentrations were determined and expressed as mM of each SCFA per gram of sample for the raw cecal/fecal material. The Agilent platform cannot discriminate between isovalerate and 2- methylbutyrate and thus reports these compounds out as a single peak and interpolated value.

### Carbon source enrichment analyses

A variation of pathway enrichment analysis (Marco-Ramell et al., 2018) was used to evaluate carbon source availability and consumption *in vivo*. Curated carbon source groups, optimized to reflect carbon source metabolism of gut commensal species, were developed with review of primarily literature regarding anaerobic metabolism of carbohydrate, amino acid and other amine-containing compounds, lipids, aromatic compounds, purines and pyrimidines, vitamins, micronutrients and other input sources for microbial metabolism and growth. Additional sources of reviewed information included published maps of *C. difficile’s* biochemical pathways (Janoir et al., 2013; Pettit et al., 2014) and BioCyc and MetaCyc content for *C. difficile* strain CD630 (Marco-Ramell et al., 2018). A carbon source group required a minimum of 6 biochemicals for evaluation. For studies in GF mice, 506 biochemicals, of 787 identified by the Metabolon panel, 64.3% of the dataset, were curated into carbon source groups (SDF 1.1-1.2). For studies in conventional mice, 667 biochemicals of 858, 77.8% of the dataset, were curated into carbon source groups (SDF 9.1-9.2).

Mass spectrometry datasets were filtered to remove biochemicals with values <50,000 counts across all samples (<3% of biochemicals). Remaining zero-value data points were assigned a value of 25,000 to support calculation of Log2 fold-change between comparisons. Datasets were Log2 transformed for significance testing of each biochemical by Welch’s T test and Benjamini-Hochberg multi-hypothesis correction (Benjamini, 1995; van den Berg et al., 2006). Thresholds for enrichment used a Log2 fold-change of ≥0.32192809 (1.25X), and a Log2 fold-change ≤ -0.32192809 (-1.25X) for depletion, and per-biochemical adjusted p value ≤0.05. Biochemicals in pairwise comparisons were ranked by adjusted p value and up to the top 40% of significantly changing biochemicals were used in analyses.

The number of enriched and depleted biochemicals per carbon source group, and total number of enriched and depleted biochemicals in datasets were calculated. Carbon source groups with ≥4 enriched or ≥4 depleted biochemicals underwent hypothesis testing by hypergeometric test, followed by Benjamini-Hochberg multi-hypothesis correction (Benjamini, 1995). An adjusted p value ≤0.05 for enriched or depleted carbon source groups was considered significant. Significantly enriched or depleted groups were plotted using the Python library Matplotlib (Hunter, 2007). Results for biochemicals within enriched groups were plotted in OriginLab (OriginLab, Wellesley Hills, MA) using the 3D XYY function, or with the Metaboanalyst 4.0 visualization tools (Chong et al., 2018).

### Cluster analyses of Stickland aromatic and histidine metabolites

Stickland aromatic amino acid and histidine metabolites with known specificity for *C. difficile* or PBI (Mead, 1971; Neumann-Schaal et al., 2019) were clustered by mouse sample using the Metaboanalyst 4.0 clustering tools and Pearson’s correlation matrices (Chong et al., 2018). Similarities among samples were evaluated by amova (Schloss, 2019).

### *C. difficile* ATCC43255 and CSAR genome sequencing and annotation

Given discrepancies for multiple genes and bacteriophage loci in the RefSeq genome for *C. difficile* ATCC43255, a closed, reference genome was generated using Oxford Nanopore GridIon sequencing (Oxford Nanopore Technologies, Oxford, UK) according to the manufacturer’s instructions. The prepared library was sequenced on a MIN106D flow cell (R9.4.1) for 72 hours using fast calling model. Reads were base called in real-time and demultiplexed using Minknow v3.6.5. The genome was *de novo* assembled using reads >5 kb in size with Flye v2.4.1 (Kolmogorov et al., 2019) to produce one circular contig with 400X coverage and size of 4,313,281 bp. A second hybrid assembly using the nanopore long reads with short reads generated by Miseq sequencing (NCBI SRA ID: SRS5656519) was generated using Unicycler v.0.4.8 (Wick et al., 2017). Mauve (Darling et al., 2011) was used to align and determine the synteny of both assemblies.

To correct potential stop codon and frameshift sequencing errors in coding regions, the hybrid assembly was compared to 26 pre-existing *de novo* SPAdes assemblies for ATCC43255 generated using Illumina MiSeq data from Worley, et al. (Worley et al., 2020). In total, 65 corrections were made and the genomic data loaded to NCBI RefSeq (Accession pending).

The updated reference genome was annotated using the NCBI Prokaryotic Genome Automatic Annotation Pipeline (Sayers et al., 2020), PATRIC (Wattam et al., 2017), and PROKKA (Seemann, 2014) to extract gene features for support of transcriptome pathway enrichment analyses. Bacteriophage loci and genes were identified using PHASTER (Arndt et al., 2019).

A genome for CSAR was generated using the methods as described in Nudel, *et al* (Nudel et al., 2018) by Illumina MiSeq, and was annotated as described. To assess presence of a putative ornithine cyclodeaminase gene, the CSAR genome was subjected to nucleotide and protein BLAST using the ornithine cyclodeaminase nucleotide coding regions and amino acid sequences from *C. difficile* ATCC43255 (geneIDs: fig|1496.5150.peg.3476, UAB_RS0202485), PBI (geneID: fig|1490.7.peg.2150), the cluster I *Clostridium, Clostridium botulinum* (NCBI GeneID: WP_072587252), and *Enterococcus faecalis* (NCBI GeneID; WP_002358614.1). No homologous genes in CSAR were identified.

### *In vivo* bacterial RNA sequencing

RNA was extracted from 15-20mg of flash frozen cecal contents (n=6 mice per group) using the Zymo Direct-zol RNA purification kit (R2081; Zymo, Irvine, CA). The quality of extracted RNA was assessed using an Agilent 2100 Bioanalyzer (Agilent Technologies, Lexington, MA) and samples with RNA Integrity Number (RIN) >= 8.0 were processed through Ribo-Zero Gold rRNA removal kit (MRZH116; Illumina, San Diego, CA) or NEBNext bacterial and rRNA Depletion Kit to deplete prokaryotic and eukaryotic rRNAs, and eukaryotic poly-A mRNAs (New England Biolabs, Ipswich, MA). The transcriptome sequencing libraries were constructed using the Illumina TruSeq mRNA Library Prep kit (20020594, 20020493; Illumina, San Diego, CA) or NEBNext Ultra II Directional RNA library prep kit (New England Biochemical, Ipswich, MA), per the manufacturer’s specifications. Library sizes were checked using a Bioanalyzer DNA High Sensitivity chip and TapeStation and quantified using Qubit dsDNA HS Assay Kit (Q32854; Thermo-Fisher, Waltham, MA). For sequencing runs, 12 libraries were pooled and sequenced on an Illumina Nextseq500 (Illumina, San Diego, CA) in paired-end 150 (PE150) nucleotide runs.

### Transcriptome Data Processing

To map reads to gene features, the *P. bifermentans* ATCC638 (DSM14991) reference genome, NZ_AVNC01000001.1, was obtained from PATRIC (Antonopoulos et al., 2017), and *Mus musculus* C57BL6/J genome, GCF_000001635.26, from NCBI. The genomes for *C. difficile* ATCC43255 and *C. sardiniense* DSM599 were generated as described.

Paired-end reads were quality filtered and trimmed then mapped to mouse and microbial genomes using Bowtie2 (Langmead B, 2012) using strict requirements for read orientation. The “--no-mixed” and “--no-discordant” flags were used to ensure that paired reads aligned to the same section of the genome in the expected orientation, respectively. Read pairs with a mapping quality <10, a measure of alignment uniqueness, were filtered. Reads aligning to >1 genome was flagged for subsequent analysis to identify potential sites of homology among genomes.

Mapped reads were assigned to gene features using HTSeq (Anders S, 2015) with flags “--nonunique all” to allow reads mapping to multiple features to be called to account for polycistronic RNAs, and “-a 10” to set the minimum mapping quality score at 10, a measure of alignment uniqueness. The identity of unaligned reads was analyzed with Kraken2 (Neves et al., 2017) to confirm association of mice with the expected species. Supplemental Table 3 shows the total and average read counts mapped to *C. difficile*, CSAR, PBI, or mouse across experimental conditions and replicates.

HTSeq results from each experimental replicate were binned by species and formatted for DESeq2 analyses (Guo et al., 2013). Gene features where no set of experimental replicates averaged more than 10 reads per replicate were filtered from further analysis (<3% of genes). A widely used DESeq2 analysis template was modified for differential expression analysis (https://gist.github.com/stephenturner/f60c1934405c127f09a6). Read data from all experimental replicates of a given organism were included for pairwise DESeq2 analyses to insure the same adjusted read counts and estimates of dispersion across pairwise comparisons.

### Transcriptome Pathway Enrichment Analyses

For *C. difficile,* mappings leveraged multiple previously published gene-level and pathway annotations for CD630 (Dembek et al., 2015; Hofmann et al., 2018; Janoir et al., 2013), assignments of gene function in the *C. difficile* EGRIN model (Arrieta-Ortiz ML, 2020), and the bacterial-based Riley schema (M, 1993) to define microbial pathways and super-pathways, with addition of pathways such as “Mucin Degradation” to describe anaerobe-host categories, or ones that were missing or incompletely annotated in public resources. An operon map of *C. difficile* genes was created from the BioCyc content for CD630 (Marco-Ramell et al., 2018). Genes present in ATCC43255, but not CD630, were treated as single-cistron operons (SDF 3.1-3.2). PATRIC and PROKKA (Seemann, 2014) annotation of the CSAR and PBI genomes were used to develop pathway maps for these species. Gene features in the commensals were also subjected to BLAST against the CD630 reference genome to provide additional annotation information. The annotated microbial gene features are shown in SDF 3.2-3.4 for *C. difficile, C. sardiniense* and *P. bifermentans,* respectively.

A minimum of 8 genes across at least 2 putative operon structures were required to define a pathway category. Thresholds for gene enrichment or depletion were set at +/-1.5X fold-change (Log2 fold-change of +/- 0.584962501) and with a DESeq2 per-gene adjusted p value ≤0.05. Up to the top 40% of significantly changing genes, ranked by the per-gene adjusted p value, were analyzed in each pairwise comparison. Pathways with a minimum of 5 enriched or of 5 depleted genes underwent hypothesis testing by hypergeometric test. Multi-hypothesis adjusted p values were calculated using the Benjamini-Hochberg method (Benjamini, 1995). Pathways with an adjusted p value ≤0.05 were considered significant. Enriched pathways were plotted using Python library Matplotlib (Hunter, 2007). Heatmaps of all genes in enriched or depleted pathway categories were visualized using the Metaboanalyst 4.0 microbiome tools (Chong et al., 2018), with hierarchical gene-level clustering by Pearson similarity and minimum-distance linkage.

### Genomic DNA extraction and qPCR

Genomic DNA was extracted from cecal contents using the Zymo Quick-DNA Fecal/Soil Microbe Miniprep Kit (kit# 11-322; Zymo, Irvine, CA) and qPCR was performed using Taqman primers and probes specific for *P. bifermentans, C. sardiniense* and *C. difficile* with the conditions as described (Abdel-Gadir et al., 2019; Bucci et al., 2016) on a QuantStudio 12K Flex Real time PCR system (Applied Biosystems, Beverly, MA). Samples were run in triplicate and compared against standard curves of known biomass of each organism spiked into germfree cecal contents and then extracted to provide normalized CFU counts per gram of cecal contents.

## Supplementary information title and legends

**Figure S1:**
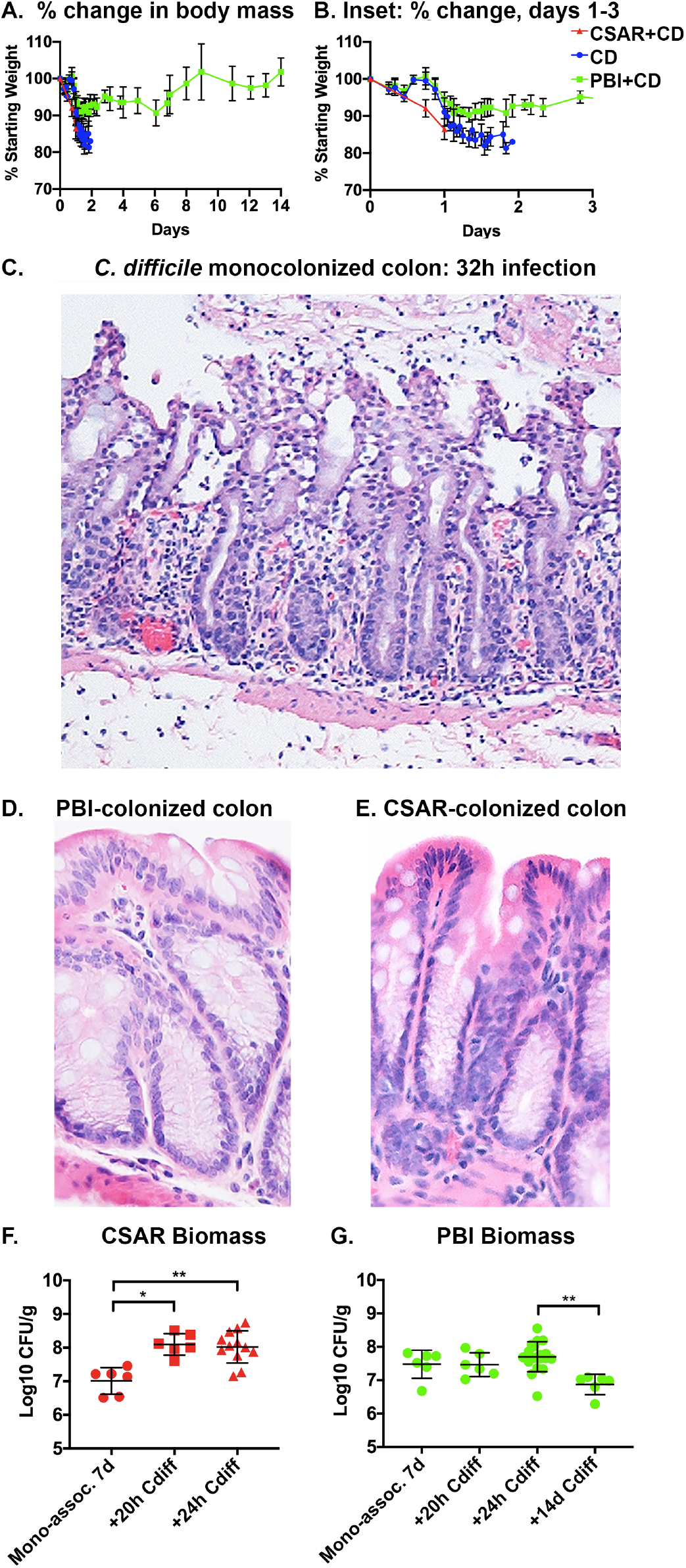
PBI protects germfree mice from lethal *C. difficile* infection while CSAR promotes worse disease. **A:** Percent change in starting mouse body mass change over infection, prior to *C. difficile* challenge with 1000 spores. Blue: *C. difficile* monocolonized mice; Green: PBI- monocolonized for 7 days prior to infection with *C. difficile;* Red: CSAR-monocolonized for 7 days prior to infection with *C. difficile.* Bars show mean and standard deviation for each group. Only PBI-associated mice survive beyond 48h. **B:** Inset of panel A showing changes over the first 72h. **C:** *C. difficile* monocolonized colon (100X) at 32h of infection showing epithelial sloughing, transmural neutrophilic infiltrates and blood entering the lumen. **D:** H&E-stained colon (200X) from PBI-monocolonized mice and **E:** CSAR-monocolonized mice after 7 days of colonization, showing intact epithelium and normal mucosa. **F:** Log10 CFU/gram of CSAR biomass in cecal contents. X-axis shows the timepoint; *p=0.0128; **p=0.0065. **G:** Log10 CFU/gram of PBI biomass in cecal contents. Timepoints shown on the X- axis; **p=0.0026.

**Figure S2:**
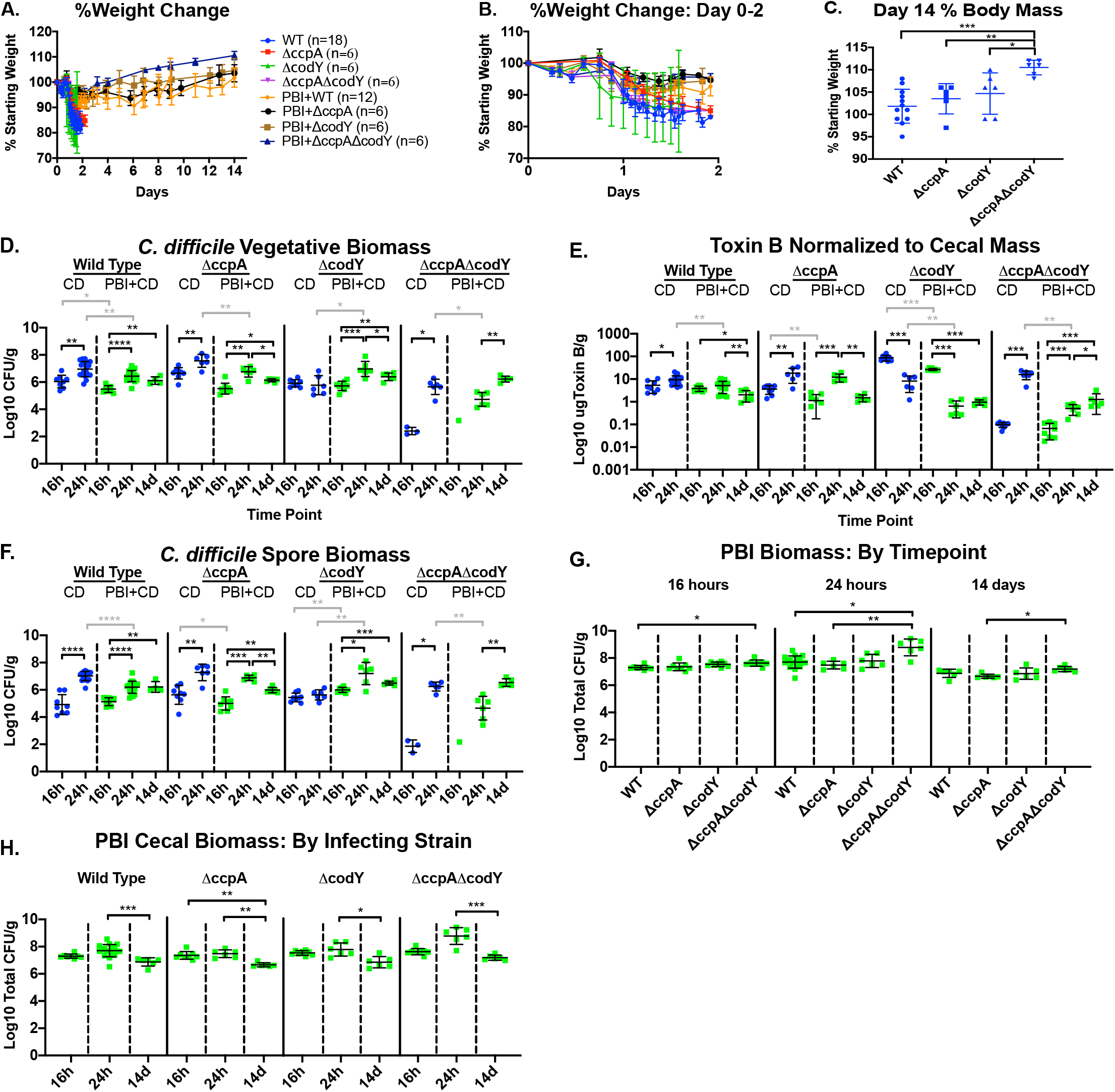
Infection with ΔcodY, ΔccpA, and ΔcodY ΔccpA mutant strains. **A:** Percent change in starting body mass in infected mice over 14 days. Color key shows colonization conditions and infecting mutant strain. **B:** Inset of panel A over the first 48h. **C:** Day 14 body mass, normalized to body mass prior to infection, in surviving PBI co-colonized mice; *0.05≥p>0.01; **0.01≥p>0.001; ***0.001≥p>0.0001; ****p>0.0001. **D-G:** Results as in main figure 6B-E shown by strain and timepoint. Blue: *C. difficile* monocolonized mice; Green: PBI monocolonzied mice. Black bars indicate significant differences by Mann-Whitney log rank test within a group, and grey horizontal bars between colonization states for a given strain. **D:** Log10 CFU/g of *C. difficile* vegetative cells or **E:** ug Toxin B per gram of cecal contents, **F:** Log 10 *C. difficile* spores per gram in cecal contents. **G:** Log10 PBI CFU per gram of cecal contents (Y-axis) by infecting mutant strain of *C. difficile* or **H:** timepoint.

**Figure S3:**
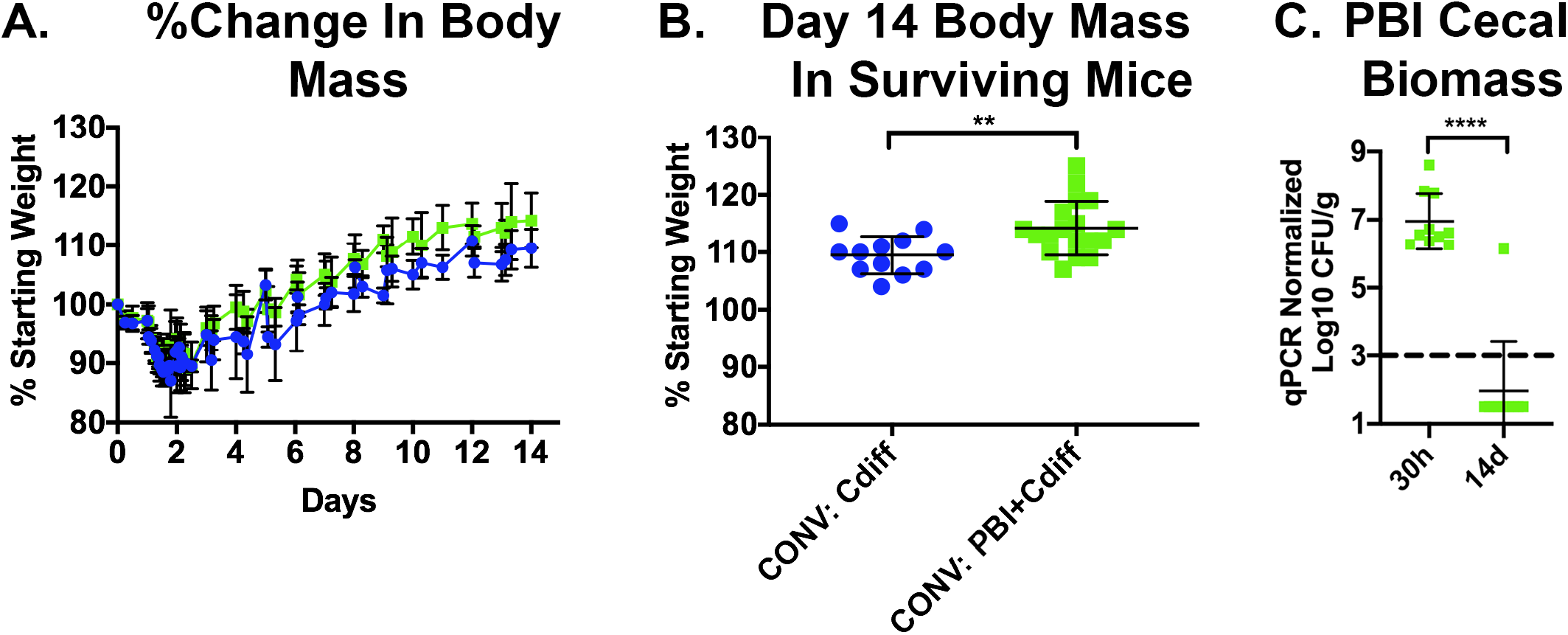
PBI administered as an oral bacteriotherapeutic protects *C. difficile* infected conventional mice. **A:** Percent change in body mass of conventional mice. Blue: *C. difficile*- infected mice receiving vehicle-alone at 12h of infection; Green: infected mice receiving 10^8^ CFU of PBI. **B.** Day 14 body mass in surviving mice normalized to starting weight prior to infection. Error bars indicate mean and standard deviation; **p=0.0062. **C.** Cecal PBI biomass by Taqman qPCR at 30h of infection (18h after oral administration of PBI), and at 14 days after challenge with *C. difficile* (****p<0.001). Horizontal dotted line shows threshold of detection at 1,000 CFU/g. Error bars indicate mean and standard deviation.

**Figure S4:**
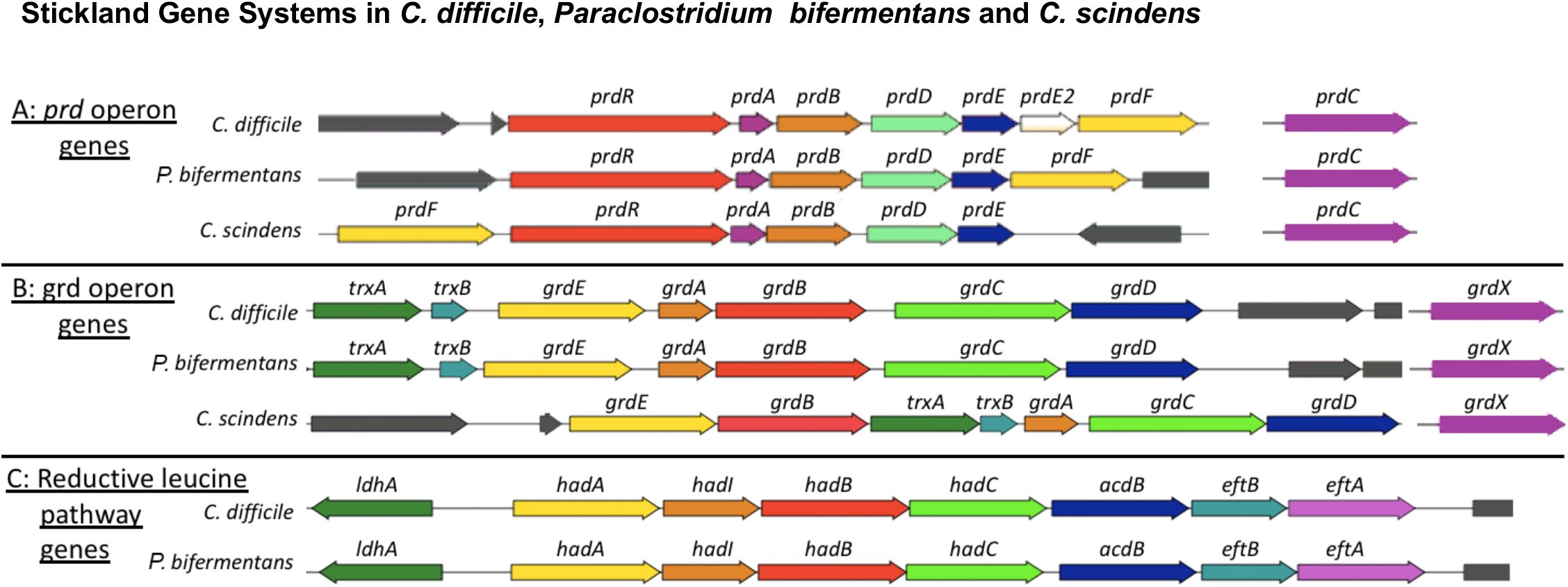
Stickland gene systems in *C. difficile, C. bifermentans* and *C. scindens*. **A:** *prd* proline-reductase genes in *C. difficile* ATCC43255*, C. bifermentans* 638 and *C. scindens* ATCC37504, i.e. the *prd* operon and the unlinked *prdC* gene. **B:** Glycine reductase (*grd*) genes. **C:** Reductive leucine pathway in *C. difficile* and *C. bifermentans* that were absent in *C. scindens.* External, non-Stickland genes are shown in grey.

**Table S1.**
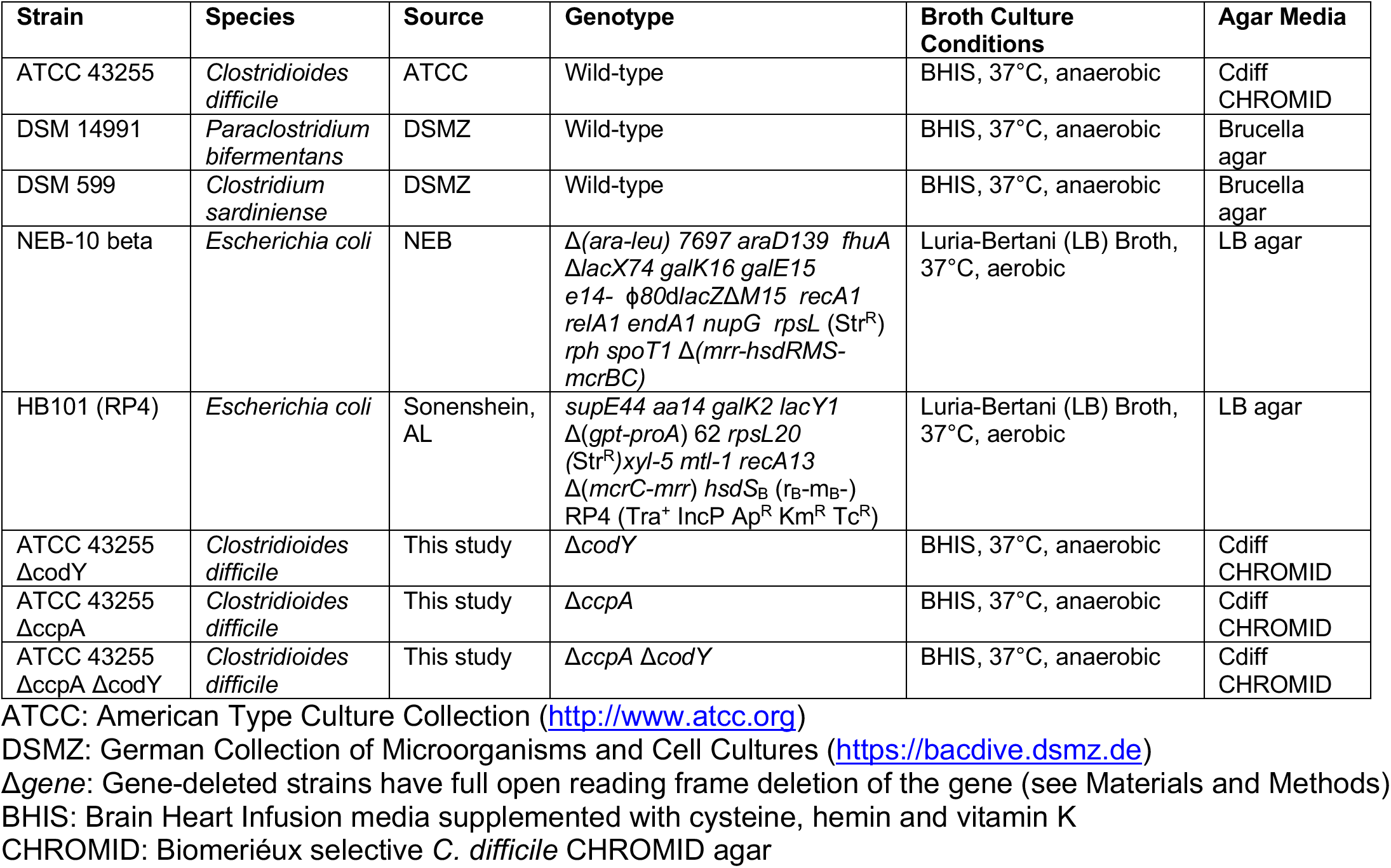
Strains and Culture Conditions. Different strains used for this study and their culture conditions are shown.

**Table S2.**
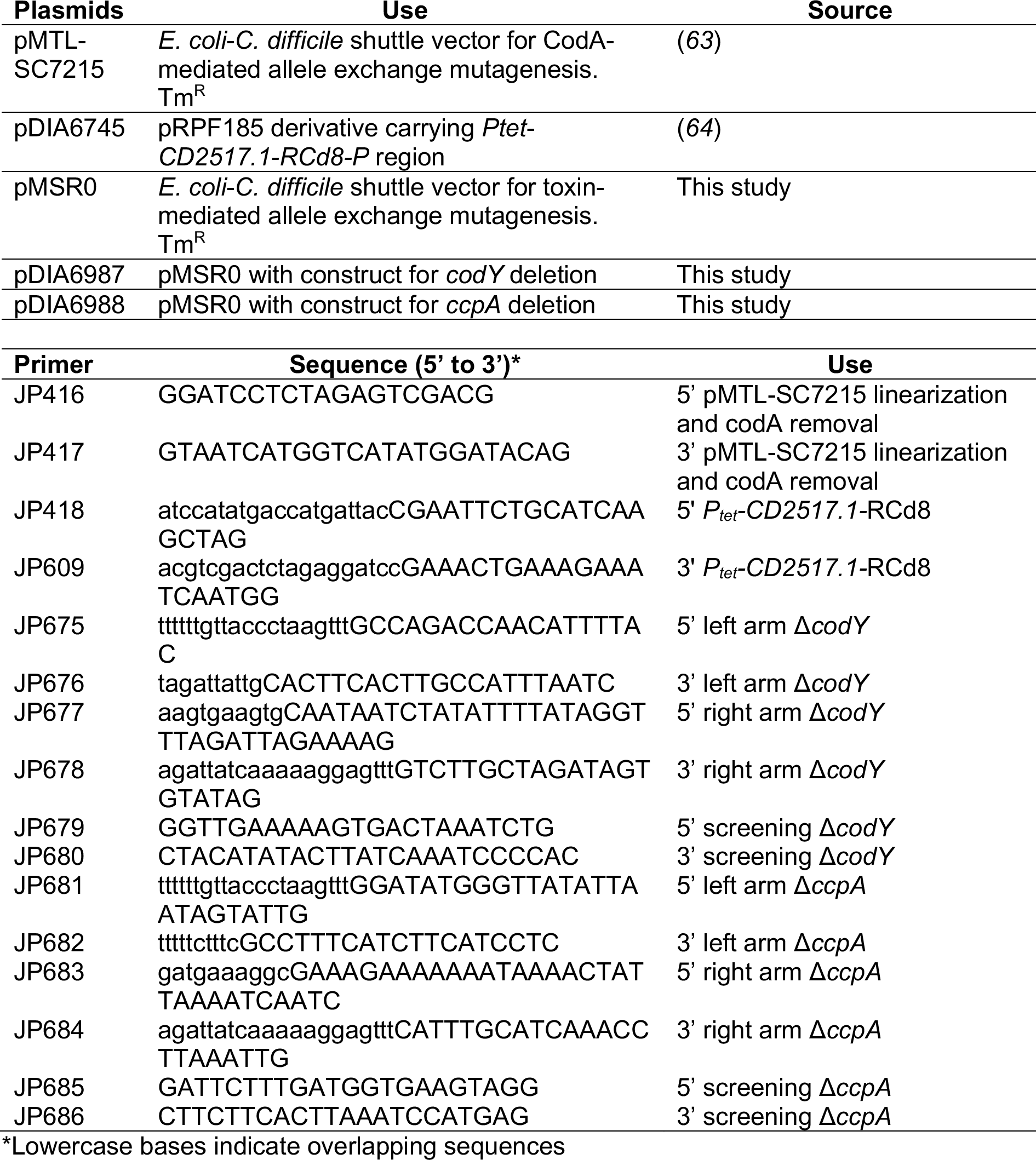
Plasmids and Oligonucleotides used in the present study.

**Table S3.** SuppelementalTable3_ReadCounts.xlsx. Table S3 shows the total and percentage of reads passing quality filtering that were assigned to each microbe, or to mouse, in addition to ambiguous reads and reads not mapping to any genome.

| Supplemental Data Files | Format |
| --- | --- |
| SDF_01 – Supplemental Text | PDF |
| SDF_02 - GF carbon source and KEGG sub-pathway maps; tabular enrichment results | Excel |
| SDF_03 - Heatmaps of biochemicals in enriched carbon source groups across *in vivo* conditions in germfree and specifically-associated mice | PDF |
| SDF_04 - *C. difficile*, CSAR and PBI gene content for transcriptome enrichment analyses and tabular results of *C. difficile* and commensal transcriptome enrichment analyses | Excel |
| SDF_05 - Heatmaps of gene expression in *C. difficile* significantly enriched pathways | PDF |
| SDF_06 - Heatmaps of gene expression in CSAR significantly enriched pathways | PDF |
| SDF_07 - Heatmaps of gene expression in PBI significantly enriched pathways | PDF |
| SDF_08 - Enriched and depleted genes not included in enrichment analyses | Excel |
| SDF_09 - CONV carbon source and KEGG sub-pathway maps; tabular enrichment results | Excel |
| SDF_10 - Biochemicals in enriched carbon source groups across *in vivo* conditions in conventional mice | PDF |

## Notes

### Summary of Updates

This manuscript version has updated analyses from use of new C. difficile EGRIN and PRIME models used to enhance gene feature annotation in C. difficile and also support in vivo predictions of metabolic and cellular pathways driving the pathogen's growth in vivo.

