## Supplemental Text File (SDF1) for "*In vivo* commensal control of *Clostridioides difficile* virulence"

**Survival studies in specifically-colonized mice:** Mice co-colonized with PBI and *C. difficile* showed reduced weight loss (Fig. S1A-B). *C. difficile*-monocolonized mice at 32h (Fig S1C) showed severe colonic pathology with complete sloughing of the epithelial surface, massive neutrophilic inflammatory infiltrates and blood entering the gut lumen. In contrast, mice monocolonized with PBI (Fig. S1D) or CSAR (Fig. S1E) showed no signs of epithelial damage or of untoward immune responses in the gut mucosa. While the cecal biomass of CSAR rose >10-fold over the course of infection, as compared to biomass in the mono-associated state (Fig. S1F), PBI's biomass did not change during acute infection (Fig. S1G). At 14 days post-infection, PBI's biomass decreased ~60% from that seen at 24h of infection (p=0.0022) though differences were not significantly different from the monocolonzied state or at 20h of infection (Fig S1G).

**Metabolomic studies in specifically-colonized mice:**

**Stickland metabolites in PBI and *C. difficile* co-colonized mice.** *C. difficile* and PBI produce unique metabolites from Stickland fermentation of aromatic amino acids (*1, 2*). Specific metabolites produced by *C. difficile* include phenyllactate from phenylalanine, indoleacetate from tryptophan, and *p*-cresol from Stickland oxidative fermentation of tyrosine and subsequent action of glycyl radical enzymes upon the 3(4-hydroxyphenyl)acetate metabolite (*3, 4*), while PBI preferentially produces 3(4-hydroxyphenyl)lactate and 3(4-hydroxyphenyl_propionate from Stickland tyrosine fermentation, 3-phenylpropionate from phenylalanine and indolelactate from tryptophan fermentation.  In contrast to many other cluster XI species, including PBI, *C. difficile* can use histidine as a Stickland donor amino acid (*5*), producing 4-imidazoleacetate and imidazole lactate from the Stickland oxidative reactions. Cluster analyses of cecal aromatic and histidine Stickland metabolites (Fig. S2A) showed that PBI and *C. difficile* co-colonized mice clustered with PBI-monocolonzied mice (Pearson's correlation 0.522; Fig. S2B), while profiles were negatively correlated with *C. difficile*-monocolonzied mice (Pearson's correlation = -0.78) even though both organisms had comparable vegetative biomass (Fig. 1C, S1G).

PBI-monocolonization enhanced levels of other aromatic compounds, including ones with hormonal and neurotransmitter effects in the gut and systemically, including serotonin, tyrosol and tyramine (SDF_2.1). In contrast GABA-eric compounds and metabolites, including g-aminobutyric acid (GABA) levels were increased in *C. sardiniense* monocolonization, a potential metabolite from anaerobic threonine metabolism. *C. difficile*-monoclonization increased luminal levels of dimethylglycine 13-fold over levels occurring in cecal contents of germfree mice (SDF_2.24), an NMDA-agonist, and compound that can arise from host as well as microbial metabolism (*6*). The functions of these compounds at concentrations occurring in the gut lumen are ill-defined, but have identified capacity for specific *Clostridial* species to enrich levels per their own metabolism and/or with host contributions.

**Carbon source group enrichment analyses**

Supplemental data file 1 shows the carbon source group map (SDF1.1-1.2) and % of biochemicals in enriched carbon source groups with a Benjamini-Hochberg adjusted p value ≤0.05 (SDF1.3-1.4). SDF_2 shows heatmaps for the component biochemicals in significantly enriched carbon source groups in specifically-colonized mice.

**SDF 2.1. Aromatic compounds** were enriched in PBI-monocolonized mice as compared to GF controls as well as in PBI and *C. difficile* co-colonized mice. These compounds include host and microbial metabolites of amino acid metabolism and host-glucuronidated derivates. As noted with PBI's enrichment of aromatic compounds, biochemicals in this group included serotonin and tryptamine. In contrast, tyrosol was singly enriched in CSAR-monoassociated and CSAR and *C. difficile*-co-colonized mice.

**SDF2.2 Diacylgrlycerols** were enriched in CSAR-monocolonized and *C. difficile*-monocolonized mice as compared to GF controls. These compounds can originate from the host and can also be generated by the action of *Clostridial* phospholipases on phosphatidylcholine and other host-origin lipids (*7*).

**SDF 2.3. Di- and Polyamines** were enriched in CSAR-monocolonized mice as compared to GF controls and depleted in PBI-monocolonzied mice. These carbon sources were further enriched in CSAR and *C. difficile* co-colonized mice as compared to *C. difficile* monocolonized mice. All compounds in this category were significantly elevated in CSAR-associated mice as compared to GF controls and contributed to luminal pools of amine compounds including ornithine and citrulline which *C. difficile* can convert to proline.

` **SDF_2.4 Dipeptides** were enriched in *C. difficile*-infected mice compared to GF controls. *C. difficile* and other proteolytic *Clostridia* express multiple excreted proteases and transport systems to import peptide fragments for growth and metabolism (*8*). Many enriched dipeptides included preferred Stickland donor or acceptor amino acids, or hydroxyproline from host or dietary collagen breakdown which Stickland fermenters can convert to proline (*9*).

**SDF 2.5. Ethanolamide Endocannabinoids** were enriched in PBI-monocolonized mice as compared to GF controls. These compounds have neurotransmitter and anti-inflammatory effects. In the gut environment, they have been shown to modulate aspects of digestive functioning, gut motility, and severity of acute inflammatory responses (*10*). PBI can actively ferment ethanolamine, of which ethanolamine head-group lipids provide a potential carbon source. Though *C. difficile* and CSAR both possess genomic machinery for ethanolamine fermentation, these effects were seen specifically with PBI colonization, and in the setting of an intact normal gut mucosa (Fig S1D). Enriched compounds in this group included behenoyl-, palmitoleoyl-, lignoceroyl-, palmitoyl-, and oleyl ethanolamides. Levels also rose substantively in cecal contents with active *C. difficile* infection potentially from damaged mucosa.

**SDF 2.6. Fatty Acid - Dicarboxylates** were depleted in mice co-colonized with PBI or with CSAR and *C. difficile,* as compared to *C. difficile* mono-associated mice. The depletion suggested microbial consumption of pre-exiting pools of host- or dietary-origin compounds. Some compounds, including sebacate and suberate, seen in CSAR and *C. difficile*-co-colonized mice may have been released with severe host cellular and mitochondrial membrane damage from infection.

**SDF_2.7. Fatty Acid intermediates** were enriched in PBI-monocolonized mice as compared to germfree controls. Malonate and multiple choline- and carnitine-conjugates were enriched with monocolonization.

**SDF_2.8. Gamma-glutamyl Amino Acids** were depleted in *C. difficile*-monocolonzied mice as compared to GF controls and were enriched with CSAR-monocolonization. γ-glutamyl amino acids can originate from host and microbial sources (*11*). γ-glutamyl conjugates of Stickland fermentable amino acids have previously been shown to be depleted with *C. difficile* infection, suggesting their potential consumption by the pathogen *(12).* In *C. difficile*-monocolonized mice, γ-glutamylglycine, γ-glutamylisoleucine, γ-glutamylvaline and others were significantly depleted at 20hr of infection. Lesser effects of depletion were seen in PBI-monocolonized mice, while CSAR-monocolonized and co-colonized mice enriched these and additional conjugates.

**SDF_2.9. Hexoses** were significantly depleted in *C. difficile*-monocolonized mice as compared to GF controls. In particular *C. difficile* actively consumed fructose, glucose and multiple 6-carbon sugar alcohols and acids (SDF_2.19).

**SDF_2.10. N-Acetyl Amino Acids:** were depleted in *C. difficile* monocolonized mice and enriched in CSAR-monocolonzied mice as compared to GF controls. N-acetyl amino acids can originate from host, dietary and microbial sources (*13, 14*). Multiple N-acetyl amino acids were enriched or depleted across colonization states, including derivatives of Stickland donor or acceptor amino acids.

**SDF_2.11. Other Fermentable Amino acids** were significantly depleted in mice monocolonized with PBI or CSAR as compared to GF mice. Colonization with either commensal depleted multiple amino acids used in other fermentation pathways. Levels of cysteine, glutamate, aspartate, lanthionine and threonine were depleted in *C. difficile*-monocolonzied mice. PBI-monocolonzied mice also depleted these amino acids and asparagine. In contrast, CSAR monocolonized mice depleted threonine, arginine and glutamine while enriching levels of cysteine, methionine, taurine, citrulline, and others. Microbial colonization also enriched N-formylmethionine over GF controls, per prokaryotic *in vivo* protein synthesis.

**SDF 2.12. Pentoses** were significantly depleted in *C. difficile*-monocolonized mice as compared to GF controls, and were enriched in PBI and *C. difficile* co-colonzied mice as compared to *C. difficile*-monocolonized mice. This category includes ribose, arabinose and xylose which can support microbial growth. Many Gram positive species can also produce ribitol and derivatives, which can be incorporated into the bacterial cell wall or used in riboflavin biosynthesis (*15, 16*).

**SDF 2.13. Phosphatidylcholines (PC)** were significantly enriched in *C. difficile*-monocolonized mice as compared to PBI-co-colonized mice. These compounds are present in host cell membranes and likely represent cellular membrane damage from infection.

**SDF 2.14 Phosphatidylethanolamines (PE**) and **SDF 2.15 Plasmalogens** were significantly enriched in *C. difficile*-monocolonized mice as compared to PBI-co-colonized mice, and enriched in CSAR-co-colonized mice as compared to *C. difficile*-monocolonized mice. As with Phosphatidylcholines, these compounds are found in abundance in epithelial and other host cell membranes. Their enrichment in *C. difficile*-monocolonized and CSAR-co-colonized mice, but not PBI-co-colonized mice, likely reflects the altered severity of infection among these groups at 20h.

**SDF 2.16. Primary Bile Acids** All three *Clostridial* species enriched primary bile acids in monocolonized mice. PBI and CSAR each enriched levels of β-muricholate, chenodeoxycholate, cholate, and their taurine conjugates. *C. difficile*-monocolonization specifically enriched cholate and β-muricholate, and the taurine conjugates taurochenodeoxycholate, taurocholate and tauro-β-muricholate. Mouse-specific tauro-β-muricholate has been shown to have germination and growth inhibitory activities against *C. difficile in vitro* (*17*). Chenodeoxycholate is similarly a strong anti-germinant *in vitro (18)*. However, very different outcomes from infection resulted from PBI and CSAR co-colonization, in spite of their similar effects on primary bile acid enrichment, suggesting that the primary bile acids, in combination with other host and microbial effects may have nominal impact in the large intestine once spores have germinated.

**SDF 2.17. Purines - Xanthines and Metabolites** were depleted in PBI and CSAR mono-associated mice as compared to GF controls, and in co-colonized CSAR and *C. difficile* mice as compared to *C. difficile* mono-associated mice. Xanthine is abundant in gut secretions, being produced as part of host nucleic acid catabolism, epithelial cell turnover, and as a substrate for the generation of ROS by neutrophils and other inflammatory cells (*19, 20*). While PBI depleted many compounds in this category it enriched levels of hypoxanthine, and 1- methylhypoxanthine potential metabolites of host and/or microbial adenine metabolism (*19, 21*).

**SDF 2.18. Pyrimidines - Cytosines** were significantly depleted in mice monocolonized with PBI or CSAR and in mice co-colonize with CSAR and *C. difficile* as compared to *C. difficile* monocolonized mice. Both PBI and CSAR enriched levels of 2'-O-methylcytidine, 5-methyl-2'-deoxycytidine, 2'-deoxycytidine and cytidine. While pyrimidine-degrading species of *Clostridia* have been identified (*22*), these activities have not been described in Cluster I or XI species.

**SDF 2.19. Pyrimidines - Thymine and Uracil compounds** were significantly enriched in *C. difficile*-monocolonized mice and depleted in PBI- and CSAR-monocolonzied mice. In infected mice, this carbon source group was also depleted in PBI-co-colonized mice as compared to *C. difficile*-monocolonzied mice. All three *Clostridial* species produced 3-ureidopropionate from uracil, with highest levels in CSAR-monocolonized and co-colonized mice (Fig 3E). Beta-alanine, however, demonstrated the most elevated levels with CSAR-monocolonization and co-colonization with *C. difficile.* This metabolite is produced through further reductive metabolism of uracil and serves as an important source for nitrogen assimilation, including from nitrogen bases.

**SDF 2.20 SCFA (volatile intermediates)** were significantly enriched in all monocolonized mice as compared to germfree controls, per microbial fermentative metabolism, including Stickland and glycolytic intermediates.

**SDF_2.21 SCFA (non-volatile)** were significantly enriched in *C. difficile*-monocolonzied vs. PBI-co-colonized mice, and included many compounds of potential host origin that could have originated from damaged mucosa.

**SDF 2.22. Sphingosine containing compounds** were significantly enriched in PBI mono-associated mice compared to GF controls. Sphingosines are amino alcohol lipids common in mammalian cell membranes, and that are also produced by commensal species of *Bacteroides (24)*. Their production has not been reported in commensal or pathogenic *Clostridia.* PBI-monocolonization enriched all sphingosine containing compounds. This enrichment co-occurred with that of ethanolamide endocannabinoids (SDF_2.5) and may suggest mechanisms by which PBI induces these host-origin compounds as potential carbon sources for its metabolism.

**SDF 2.23. Stickland Acceptor Amino Acids** were significantly depleted in mice monocolonzied with *C. difficile* or with PBI as compared to GF controls. This carbon source group includes the Stickland acceptor amino acids proline, leucine, and glycine and compounds convertible to these amino acids including hydroxyproline derivatives. These and additional potential Stickland acceptors were further enriched in CSAR-monocolonization and in CSAR and *C. difficile* co-colonized mice.

**SDF 2.24. Stickland Donor Amino Acids**: were enriched in CSAR-monocolonized and in *C. difficile*-monocolonized mice, and in *C. difficile* and CSAR-co-colonized mice.

**SDF 2.25. Sugar Alcohols and Acids** were significantly depleted in all *C. difficile*-infected mice. *C. difficile* metabolizes many sugar alcohols and acids of dietary origin, including sorbitol, mannitol, galactitol and others (*8*). Given the poor absorption of these compounds from the mouse and human gut, they provide a potentially available carbon source for metabolism and growth.

**SDF 2.26. Vitamin and Cofactors** were significantly enriched in PBI and *C. difficile*-co-colonzied mice, particularly for riboflavin, pyridoxal (vitamin B6) and nicotinamide.

**Sub-pathway enrichment analyses of specifically-colonized mice**

Metabolomic enrichment analyses were also performed using the Metabolon and KEGG-based Sub-Pathway mappings (SDF_1.5-1.7), information more attuned to host metabolic pathways (Figs. S3C-D). Findings for purines, pyrimidines and bile acids mirrored those found with the content for the anaerobe-focused carbon source groups. In comparisons between GF and mono-colonized mice, enriched categories included “Tryptophan Metabolism”, “Leucine, Isoleucine and Valine Metabolism”, and “Benzoate Metabolism” per the production of Stickland metabolites in PBI-monocolonized and *C. difficile*-monocolonized mice. The enrichment of “Urea Cycle: Arginine and Proline Metabolism” in germfree vs PBI-monocolonized mice reflected the commensal’s depletion of proline and proline-convertible compounds in Stickland metabolism in combination with arginine and related metabolites in its arginine deiminase fermentation pathway (Fig S3C).

In *C. difficile*-monocolonized versus PBI and *C. difficile*-co-colonized mice the enrichment of “Glutathione Metabolism, Ketone Bodies” in co-colonized mice captured enrichment of multiple SCFA and fatty acid biosynthetic intermediates from microbial and potential host origin.

***In vivo* transcriptomic analyses**

SDF3 show the gene content used in pathway enrichment analyses for *C. difficile*, CSAR, and PBI. SDF_4 has tabular results of enriched pathways shown in Figs. 3 and S3.

***C. difficile in vivo* enriched pathways**

SDF_5 shows heatmaps of *C. difficile* genes in enriched pathways. Alterations further highlighted *C. difficile’s* diverse metabolic machinery and capacity to adapt to changing conditions within the gut from commensal colonization and host responses to infection.

In monocolonized mice, the pathogen enriched genes for the transport and metabolism of beta-glucosides (SDF_5.2), PTS and ABC transport systems for glucose and maltose (SDF_5.8), fructose (SDF_5.7), ribose (SDF_5.13), and transport systems for dipeptides and oligopeptides (SDF_5.4). The *appABC* and *oppABCD* dipeptide transport systems, in particular, were enriched at 20 hours of infection in *C. difficile*-monocolonzied mice and were down-regulated with CSAR co-colonization, potentially from more readily available fermentable amino acids and polyamines. The pathogen up-regulated butanoate (SDF_5.3), and pyruvate metabolism genes (SDF_5.12) including a pyruvate formate lyase complex that couples acetogenesis in the also enriched Wood-Ljungdahl pathway (SDF_5.17) to other metabolic processes (*25*), and *aco* operon genes to convert acetoin to pyruvate (*26*). By 24h of infection, the pathogen up-regulated ethanolamine utilization genes, including those associated with the structural carboxysome and fermentative enzymes (*27, 28*) (SDF_5.6).

With CSAR-co-colonization, *C. difficile* upregulated multiple amino acid and amine transport systems (SDF_5.16), including ABC amino acid transport systems, the polyamine *potABCD* operon genes, and genes in operon_0246 to convert ornithine to alanine (SDF_5.10). With PBI co-colonization, *C. difficile* up-regulated the *brnQ1* and *brnQ2* branched-chain amino acid transporters, and genes associated with xanthine and hypoxanthine transport and metabolism (SDF_5.18), including the xanthine dehydrogenase enzyme complex (*29*). Fermentation of xanthines has been described in other *Clostridia*, particularly Cluster XII species (*19*) which metabolize many purines to glycine and to acetate, in species carrying Stickland glycine reductase genes. Similar studies have not been conducted in Cluster XI species to determine their capacity for comparable metabolism. Xanthines may also support functions to counter osmotic or oxidative stresses encountered *in vivo*.

With PBI co-colonization *C. difficile* enriched genes to metabolize polysaccharides (SDF_5.11) including starch, per $\alpha$-amylase (geneID UAB_RS0205550), and $\alpha$-glucoside PTS transport systems (operon_1762) (*22*). The pathogen also up-regulated multiple transport and conversion enzymes for sugar alcohols (SDF_5.15), including sorbitol (*srl*), and mannitol (*mtl*) operon genes.

Commensal colonization also profoundly altered *C. difficile's* cellular machinery. By 24h co-colonization with PBI, the pathogen profoundly down-regulated genes for ribosome synthesis (SDF_5.22) and protein translation (SDF_5.21). In contrast, at 20h of infection in CSAR-co-colonized mice, the pathogen showed gene enrichment of cellular machinery for transcription (SDF_5.23), translation (SDF_5.21), and DNA replication (7.20), while by 24h of co-colonization with CSAR, systems modifying cell surface components (SDF_5.25) were enriched. In *C. difficile*-monocolonized mice. Peptidoglycan hydrolases and other genes involved in cell wall turnover (SDF_5.24) were enriched at 20h.

Genes involved in cobalamin (SDF_5.26) biosynthesis were most highly expressed at 20h in *C. difficile*-monocolonized mice, and were down-regulated in co-colonzation with either commensal, suggesting commensal and/or host cross-feeding. Cobalamin is a required co-factor in carbon metabolism, and for methionine, purine, and pyrimidine biosynthesis. In carbon metabolism adenosylcobalamin provides a key cofactor for enzymes involved in ethanolamine utilization and ornithine metabolism to alanine. With CSAR-co-colonization, *C. difficile* also showed depletion of genes involved in folate biosynthesis, again suggesting cross-feeding of this vitamin from the commensal (5.28). The *ssuABC* and *ssu2ABC* operon systems for transport of sulfur and sulfonate-containing compounds (SDF_5.28) were also up-regulated at 20h in monocolonized mice, as were genes in the thiolation pathway of cysteine biosynthesis (*30*) a fermentable amino acid and one also critical for surviving oxidative stress. Nitrogen and nitrate transport and metabolism genes (SDF_5.31) were enriched in monocolonized mice at 20h of infection, and in PBI-co-colonized mice at 24h of infection. These genes modulate critical aspects of cellular nitrogen metabolism including detoxification of host-produced reactive nitrogen species. By 24h of infection in CSAR-co-colonized mice, *C. difficile* up-regulated multiple gene systems for cation transport (SDF_5.26) and for the transport of phosphorus and phosphonates (SDF_5.30).

Cysteine and methionine biosynthesis (SDF_5.32), particularly methionine synthesis genes in the *met* operon, were up-regulated by 24h in PBI-co-colonized mice. Multiple genes for fatty acid and phospholipid synthesis (SDF_5.33) were up-regulated in CSAR-co-colonized mice by 24h. Terpenoid synthesis genes (SDF_5.35), components of Gram positive spore coats and potential anti-oxidants (*31, 32*) were also up-regulated with CSAR-co-colonization, concomitant with up-regulation of sporulation programs and increased pathogen spore biomass (Fig 1K). Biosynthetic pathways for histidine production were also enriched (SDF_5.34).

*C. difficile* up-regulated multiple ABC-family transporter systems (SDF_5.36) of unknown specificities, particularly by 24h of infection with either commensal.

Stress responses were differentially expressed among colonization states. At 20h of infection, *C. difficile*-monocolonized mice expressed CRISPR (SDF_5.38), diffocin locus genes (SDF_5.39) (*33*) chemotaxis systems (SDF_5.37) and pathogenicity locus genes (SDF_5.41), including PaLoc genes and microbial toxin/anti-toxin systems. By 24h, sporulation programs (SDF_5.42) and multiple oxidative stress response genes (SDF_5.40) were highly induced in CSAR-co-colonized mice, including ruberythrins and the spore coat ROS detoxifying enzymes *sodA* and *cotG*.

**Additional pathogen and commensal alterations in gene expression *in vivo***

Supplemental Data File 10 (SDF_8.1) shows genes with significantly altered expression that were not incorporated in enrichment analyses, per non-specific gene identification, or insufficient underlying gene content to support placement into a usable pathway.

*C. difficile's* 7$\alpha$-hydroxysterol-dehydrogenase (geneID UAB_RS0201080; SDF_8.3) was up-regulated 2-fold at 24h in co-colonization with CSAR. Most other features were members of 2-component signal transduction systems, transcriptional regulators, membrane proteins, putative diguanylate-binding signaling proteins, or hypothetical proteins. While specific functions cannot at be assigned at this time, these alterations underscore the organismal-level changes occurring relative to the monocolonized and co-colonized states.

*C. sardininense* is a cluster I *Clostridium* phylogenetically related to *C. baratii* and *C. perfringens (34).* The organism has multiple gene systems involved in mucin degradation and metabolism, including exported glycosidases, proteases, and genes to ferment host-derived sugars including fucose, sialic acid, N-acetyl glucosamine and N-acetyl galactosamine (SDF_3.3). *In vitro,* CSAR is an abundant producer of hydrogen gas (*35*) and carries nickel transport systems and iron-nickel complexing machinery to form the core of its endogenous hydrogenase system. It also carries gene systems for the fermentation of carbon sources to propanediol, and utilization of ethanolamine, both of which likely occur in a carboxysome-like structure (SDF_3.3).

CSAR-monocolonized mice (SDF_8.6, 8.7) showed significantly increased expression of genes associated with queosine uptake and metabolism, a modified nucleoside found in prokaryotic tRNAs (fig|29369.6.peg.1854, fig|29369.6.peg.1863, fig|29369.6.peg.3177), and that co-occurred with enriched ribosome production and protein translation. At 20h of infection CSAR upregulated a hyaluronidase lyase (fig|29369.6.peg.1175), acetoin-specific 2,3-butanediol dehydrogenase (EC 1.1.1.4; fig|29369.6.peg.1605), two putative aspartate aminotransferases (fig|29369.6.peg.2510, fig|29369.6.peg.1356), and a putative malate transporter (fig|29369.6.peg.1426). At 24h of infection a DNA nucleoid binding protein (fig|29369.6.peg.3418), putative succinyl co-A synthase (fig|29369.6.peg.1628), and putative oxidoreductase family protein (fig|29369.6.peg.3013) were upregulated. This latter gene has general homology to oxo-acyl carrier protein reductases, and 71% amino acid identity to the *C. difficile* 7$\alpha$-HSDH, though alterations in secondary bile acid production were not detected.

*P. bifermentans*, like *C. difficile,* is a cluster XI *Clostridium,* and is phylogenetically closest to *C. sordelli (38).* As with *C. difficile,* PBI carries the proline, glycine and leucine Stickland pathways, operons supporting transport and fermentation of ethanolamine, arginine (Fig S3F), and carbohydrate transport and fermentation pathways for glucose, fructose and ribose. In contrast to *C. difficile,* PBI is more restricted in its carbohydrate use and does not utilize sugar alcohols, the disaccharides lactose and cellobiose, or starches (*35*).

PBI induced expression of its bile-acid deconjugating enzyme choloylglycine hydrolase (fig|1490.7.peg.621) at 20h of co-colonization with *C. difficile*, in addition to transporters for niacin (fig|1490.7.peg.220) and queosine (fig|1490.7.peg.1388; SDF_8.8). By 24 hours of infection, the niacin transporter was further upregulated (SDF_8.9).

As with *C. difficile* both commensals showed altered expression of multiple transcriptional regulators, 2-component sensors, membrane proteins and hypothetical proteins, from the monocolonized state versus at 20h and 24h of infection.

**PBI-mediated protection against *C. difficile* ΔcodY, ΔccpA, and ΔcodY ΔccpA mutant strains.** Figs. S4A-C show body mass changes in GF or PBI-precolonized mice infected with wild-type or mutant strains. Figs. S4D-F show the vegetative biomass, toxin B per gram of cecal contents, and spore biomass, respectively, by timepoint. Figs. S4G-H show PBI's biomass by timepoint and infecting mutant strain. At 24h of infection, PBI's biomass increased with the ΔcodY ΔccpA double mutant over the WT and ΔccpA strains, but not with the ΔcodY mutant, which may indicate PBI's capacity to better out-compete the double mutant and to respond to increased availability of nutrients toxin-induced host damage.

**PBI oral bacteriotherapeutic protects conventional mice infected with *C. difficile*.**

Figs. S5A-B show body mass changes in conventional mice infected with *C. difficile* after clindamycin treatment and administration of $10^8$ CFU of *P. bifermentans* or vehicle alone at 12 hours of infection. Days 2-14 in wild-type infected mice receiving vehicle alone represent 60% of as opposed to 100% of PBI-treated surviving mice. By 14 days post-infection, PBI-treated mice had improved weight gain (Fig. S5B). Neither *C. difficile,* nor PBI, persisted to 14 days in most mice (Fig. S5C). SDF_09 shows carbon source groups used in analyses of conventional mice and tabular results from enrichment analyses. SDF_10 shows changes in the component biochemicals in enriched and depleted carbon source groups across conditions.

**10.1 Ascorbate-related Compounds** were enriched in post-clindamycin-treated mice as compared to pre-treatment mice. Compounds included ascorbic acid, sulfated derivatives and gulonate. Many gut microbial species can metabolize these compounds. They also are potent anti-oxidants for host and microbial cells.

**10.2. Ceramides** were enriched in mice infected with *C. difficile* as compared to Clindamycin treated mice. These components of host cell membranes may represent release from damaged mucosa.

**10.3. Dipeptides** were depleted in mice treated with Clindamycin as compared to untreated mice. In addition to cluster XI *Clostridia*, many components of the colonic microbiota are proteolytic and contribute to the breakdown of host, microbial and residual dietary proteins. With antibiotic treatment these activities are known to be reduced (*44*).

**10.4. Di- and polyamines** were enriched in clindamycin-treated mice versus pre-treatment conventional mice. These compounds represent an additional potential metabolizable carbon source used by *C. difficile.*

**10.5. Disaccharides and Oligosaccharides** were enriched in post-clindamycin-treated mice and depleted in *C. difficile* infected mice treated with PBI or with vehicle-alone. Sugars depleted with *C. difficile* infection included many that the pathogen can metabolize, including galactinol, raffinose and sucrose (*35*).

**10.6. Ethanolamide Endocannabinoids** were depleted in mice 24 hours after clindamycin treatment but showed improved recovery in *C. difficile*-infected mice treated with PBI. The single dose of clindamycin depleted 67% of ethanolamide endocannabinoids, specifically arachidonyl-, behenoyl-, erucoyl-, lignoceroyl-, margaroyl-, oleoyl-, palmitoyl-, and steroyl-ethanolamides. C. *difficile* mice treated with PBI showed re-enrichment of these compounds at 30h post-infection (18h post-treatment) except for palmitoyl- and steroyl- ethanolamides. The enrichment with PBI bacteriotherapy post-*C. difficile* challenge, mirrored that observed in PBI-monocolonized mice. Of the listed endocannabinoids, oleoyl-ethanolamide, which showed recovery at 30h, and palmitoyl-ethanolamide, which did not, have known anti-inflammatory effects per their binding to host PPAR$\alpha$ and other G-protein receptors on host cells, including gut epithelium (*10, 45, 46*).

**10.7 Fatty Acids, Monohydroxy** were significantly enriched in pre- versus post-clindamycin treated mice, and in *C. difficile*-infected mice treated with PBI versus post-clindamycin mice. These compounds are commonly of host or and dietary origin but can undergo microbial biotransformation within the gut lumen, as potentially reflected by depletion of many biochemicals in this category after clindamycin treatment, and with some improved recovery in mice treated with PBI, but not vehicle-alone.

**10.8. Gamma-glutamyl Amino Acids:** were enriched in mice treated with clindamycin as compared to untreated mice, and were depleted in all *C. difficile*-infected mice. Clindamycin treatment enriched levels of all detected gamma-glutamyl amino acids, showing overlap with the enrichment in this category seen in CSAR-monoassociated mice (Fig. 2A). Levels showed depletion with active *C. difficile* infection at 30h post-challenge, with additional depletion seen in mice treated with PBI but not with the vehicle-control only.

**10.9. Long chain fatty acids** were enriched in *C. difficile*-infected mice as compared to post-clindamycin-treated mice. Enriched compounds in this class were primarily acyl-carnitines and may reflect enhanced damage to host mucosa.

**10.10. Pentoses** were enriched post-clindamycin treated mice as compared to mice pre-treatment, and were subsequently depleted in *C. difficile*-infected mice, whether treated with PBI or vehicle alone. As noted in germfree mice, *C. difficile* actively consumes available pentoses in the gut lumen, and showed overlap with monocolonized mice in consuming Arabitol/Xylitol, Arabonate/Xylonate and Ribitiol derivatives: Ribitol, Ribonate and Ribulonate/Xylulonate.

**10.11 Phosphatidylcholines** were enriched in *C. difficile*-infected mice as compared to post-clindamycin-treated mice, including oleoyl-, palmitoyl- and steroyl-lipids, and may reflect release from damaged host cell membranes.

**10.12. Plasmalogens** were enriched in mice infected with *C. difficile* as compared to Clindamycin treated mice. Plasmalogens are components of host cell membranes and are produced by some commensal species (*47-49*). Their enriched levels in *C. difficile* mice treated with vehicle alone, but not with PBI treatment, may reflect worse mucosal damage from infection.

**10.13. Pyrimidines-Xanthines, 10.14. Pyrimidines-Cytidines, and 10.15 Pyrimidines Thymine and Uracil Compounds** were depleted in post-clindamycin treated mice as compared to untreated mice and showed levels of recovery in *C. difficile*-infected mice. The single clindamycin dose substantively altered luminal pools of nitrogen bases. Many of the compounds rebounded by 30h of infection, potentially from mucosal damage, the host's inflammatory response, and/or growth of endogenous and exogenously administered *C. difficile* and PBI.

**10.16. Secondary Bile Acids** were depleted in clindamycin-treated mice treated as compared to untreated mice**,** per antibiotic ablation of the commensal microbiota. Levels had not recovered to pre-clindamycin levels by 30h of infection.

**10.17 SCFAs** showed enrichment in *C. difficile*-infected mice as compared to post-clindamycin treated mice. Antibiotic treatment depleted multiple branched-chain amino acid Stickland metabolites detected by the Metabolon panel, while alpha-hydroxy forms of these acids were enriched.

**10.18. Sphingosines** were significantly depleted in mice treated with clindamycin as compared to untreated mice. As in PBI-associated GF mice, components of a normal complex microbiota, induced higher levels of the compounds, some of which are likely of host origin but which may also come from microbial sources including species of the Bacteroidetes, as primary or precursors to microbial sphingolipid synthesis (*24, 50*).

**10.19. Stickland Acceptor Amino Acids** were enriched in mice after clindamycin treatment, and depleted in mice infected with *C. difficile*, whether treated with PBI or vehicle alone. The findings with Stickland acceptor amino acids mirror that seen in germfree and CSAR-monocolonized mice that had enriched levels, particularly of proline, glycine, leucine, and convertible derivatives including hydroxyproline and dipeptides containing these amino acid sources. Their enrichment after a single clindamycin dose illustrates a critical mechanism by which antibiotic ablation of the gut microbiota can rapidly create conditions conducive to the pathogen's colonization, rapid growth, and elaboration of toxin.

**10.20. Vitamins and Cofactors** were depleted in clindamycin-treatmed mice, and showed enrichment in mice receiving PBI as a therapeutic during *C. difficile* infection. Clindamycin depleted multiple vitamins, and associated metabolites and biosynthetic precursors, particularly among vitamin B6, nicotinamine, thiamin, riboflavin and pantothenic acid. The specificity of these changes from specific ablation of the microbiota suggests that many of these compounds in the gut lumen come from microbial sources. Interestinaly, levels of many of these compounds showed recovery with bacteriotherapy with PBI, but not with vehicle control, in infected mice.

### Sub-pathway enrichment analyses of conventional mice

Enrichment analyses were also performed using the Metabolon and KEGG-based Sub-Pathway mappings (SDF_11.5-11.7; Figs. S3C-D). The host-focused mappings identified multiple enriched lipid categories from potential host cell membrane damage from infection, including Plasmalogens, Sphingomyelins, and Phospatidylcholines. Notably, the single dose of clindamycin enriched many categories of host and dietary origin including Phosphatidylethanolamines, Lysophospholipids, Flavonoids, and Food/Plant Components, the latter of which included sugar alcohols and ferulates, compounds present in foodstuffs and from microbial metabolism of plant-based vanillins (*51*).
