## Supplemental Data File 2 for "*In vivo* commensal control of *Clostridioides difficile* virulence"

### Supplemental Data File 2: Carbon Source Enrichment Analyses in Specifically-associated Mice

Version 1.0; 01/03/2020

**Page 0:** Introduction, this page
**Page 1**: Sample Key


Page 2: 2.1 Aromatic compounds
3: 2.2 Diacylglycerols
4: 2.3 Di- and Polyamines
5: 2.4 Dipeptides
6: 2.5 Ethanolamide Endocannabinoids
7: 2.6 Fatty Acid Dicarboxylates
8: 2.7 Fatty Acid Intermediates
9: 2.8 Gamma-glutamyl Amino Acids
10: 2.9 Hexoses
11: 2.10 N-acetyl Amino Acids
12: 2.11 Other Fermentable Amino acids
13: 2.12 Pentoses
14: 2.13 Phosphatidylcholines (PC)
15: 2.14 Phosphatidylethanoloamines (PE)
16: 2.15 Plasmalogens
Page 17: 2.16 Primary Bile Acids
18: 2.17 Purines-Xanthines and Metabolites
19: 2.18 Pyrimidines-Cytidines
20: 2.19 Pyrimidines-Thymine and Uracil Compounds
21: 2.20 SCFA
22: 2.21 SCFA (non-volatile)
23: 2.22 Sphingosine Containing Compounds
24: 2.23 Stickland Acceptor Amino Acids
25: 2.24 Stickland Donor Amino Acids
26: 2.25 Sugar Alcohols and Acids
27: 2.26 Vitamins and Cofactors

### SDF2: Sample Key

#### 2.15 Pyrimidines: Cytidines

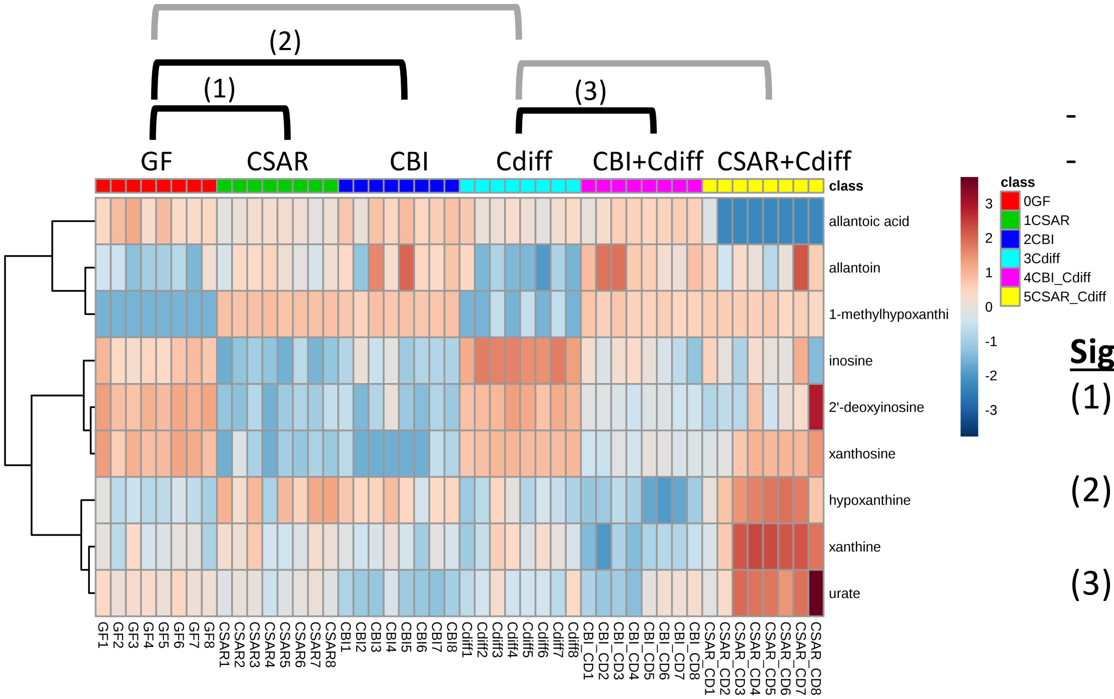


**Guide:**

- Group showing the enriched category is at the top ("2.15 Pyrimidines: Cytidines")
- Brackets indicate the 5 comparisons evaluated among groups for significant enrichment of the given carbon source group.
- Brackets shown in black for each group were significant.
- Significant comparisons and p values shown in the text box on the right-hand side

**Significantly Enriched Conditions:**

(1) Enriched in GF vs CSAR mono-associated mice (p=1.97E-03).
(2) Enriched in GF vs CBI mono-associated mice (p=2.04E-02).
(3) Enriched in Cdiff vs CBI+Cdiff infected mice (p=9.40E-03).

### 2.1 Aromatic Compounds

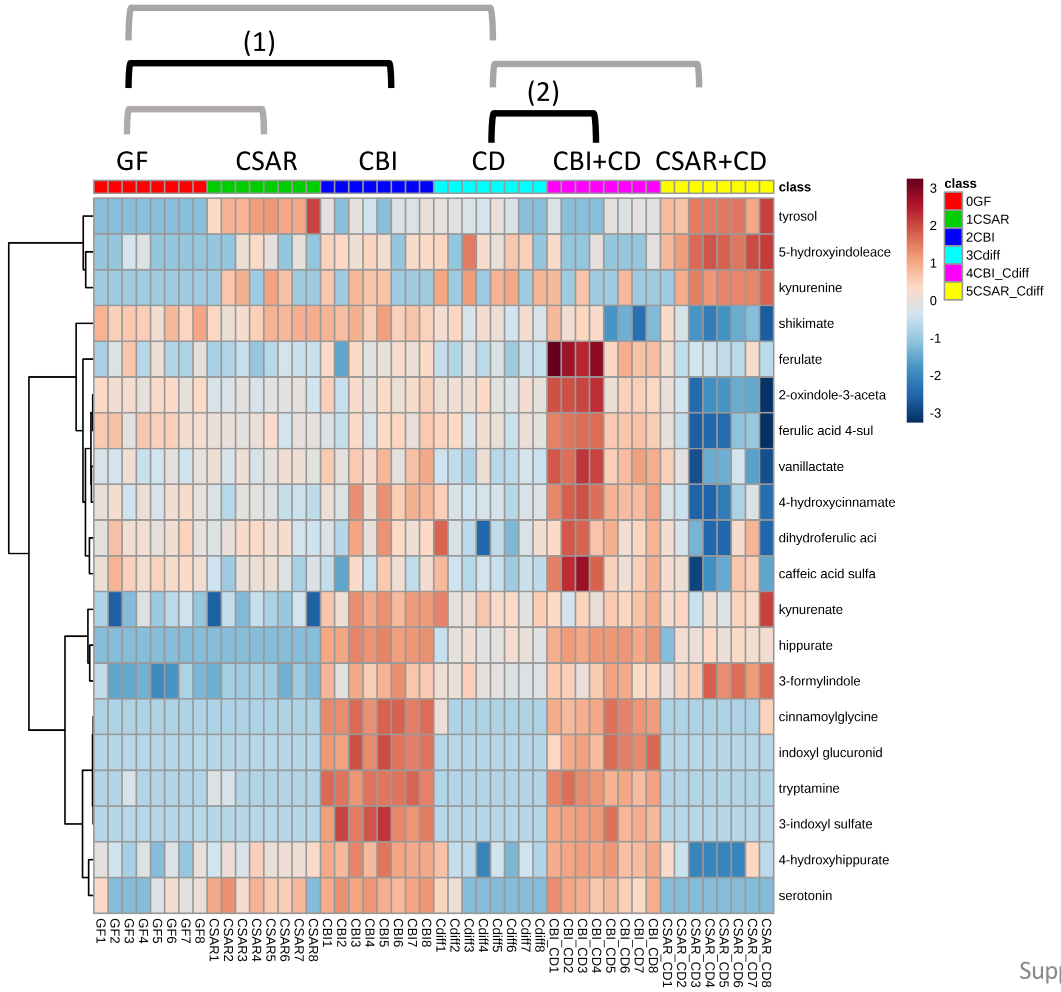


**<u>Significantly Enriched Conditions:</u>**

(1) Enriched in CBI mono-colonized vs GF mice at 7d (p=3.07E-03).

(2) Enriched in CBI+ *C. difficile* co-colonized vs *C. difficile* mono-colonized mice at 20h (p=3.95E-06).

### 2.2 Diacylglycerols

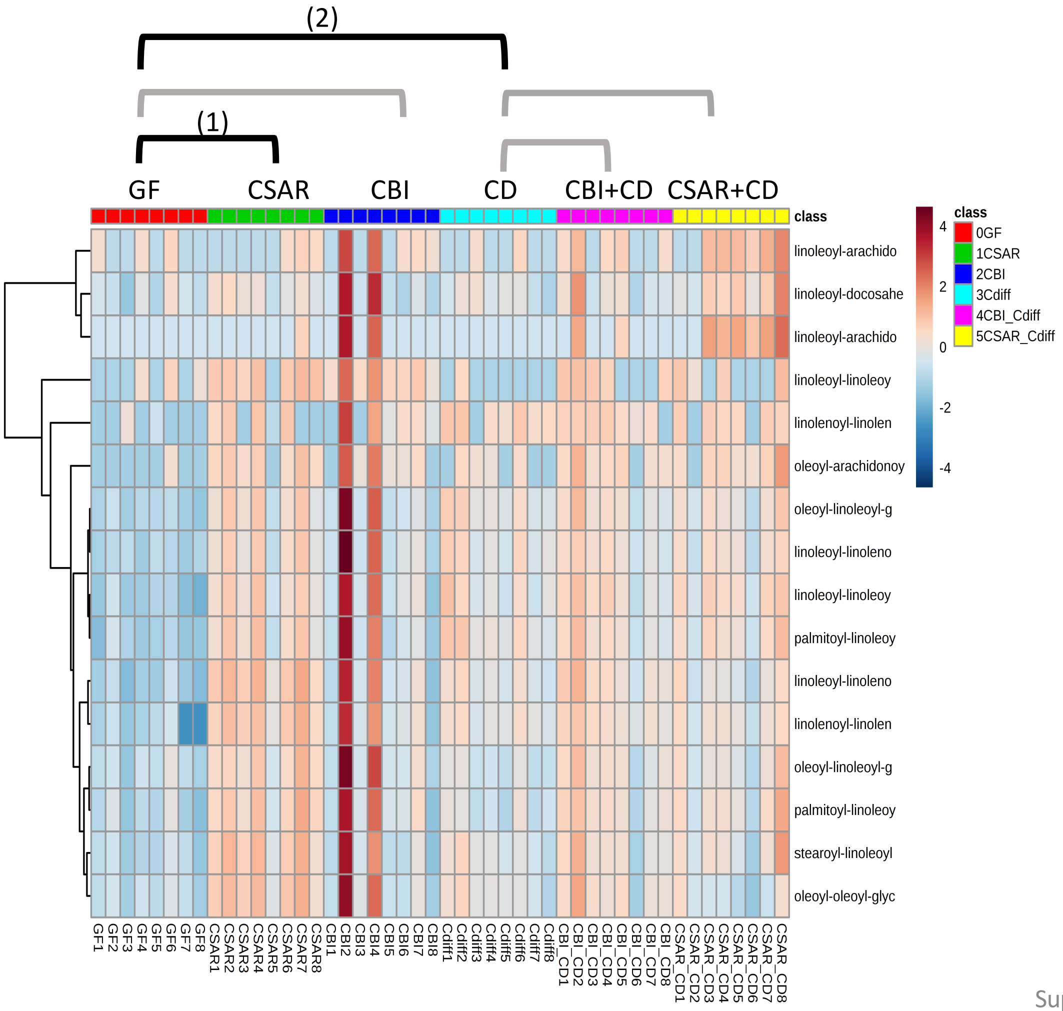


**Significantly Enriched Conditions:**

(1) Enriched in CSAR mono-colonized vs GF mice at 7d (p=9.22E-04).

(2) Enriched in *C. difficile* mono-colonized vs GF mice at 20h (p=1.30E-02).

#### 2.3 Di- and Polyamines

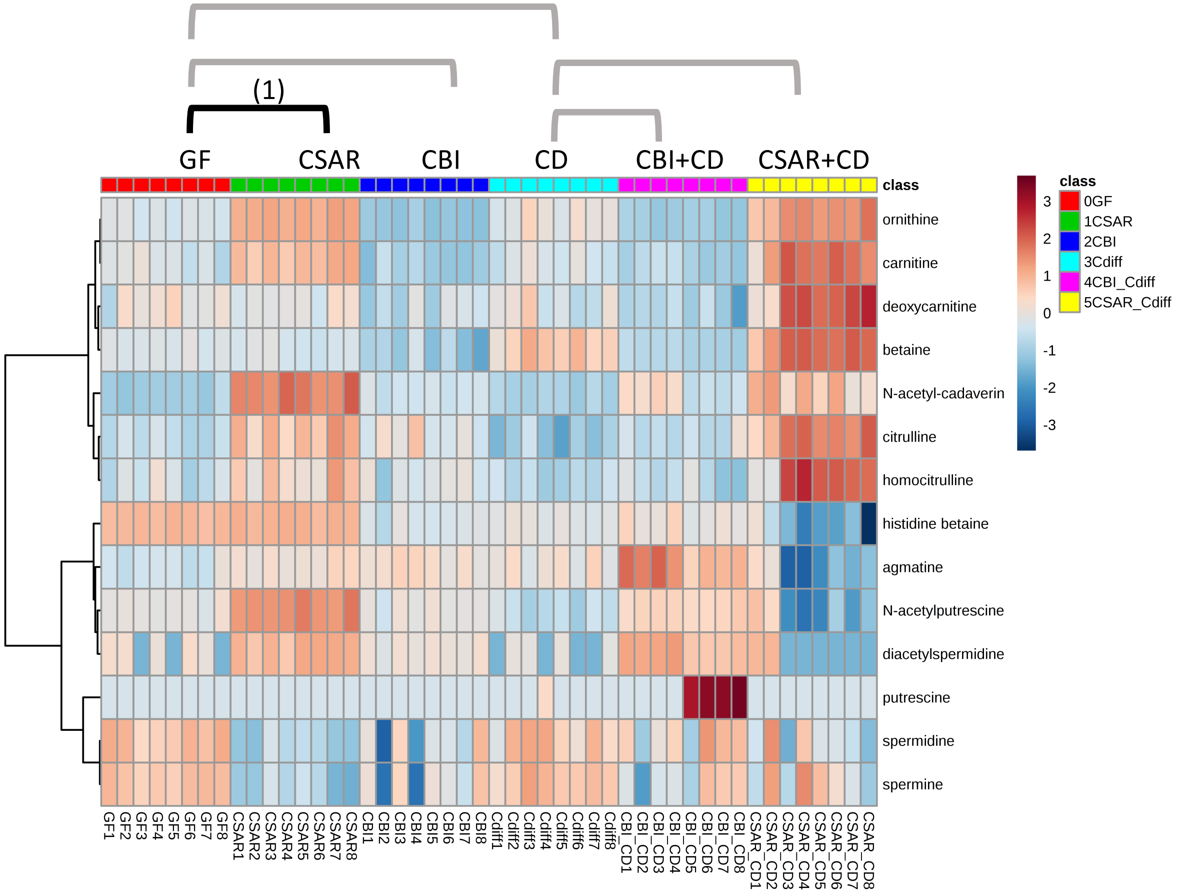


**Significantly Enriched Conditions:**

(1) Enriched in CSAR mono-colonized vs GF mice at 7d ($p=2.42E-02$).

### 2.4 Dipeptides

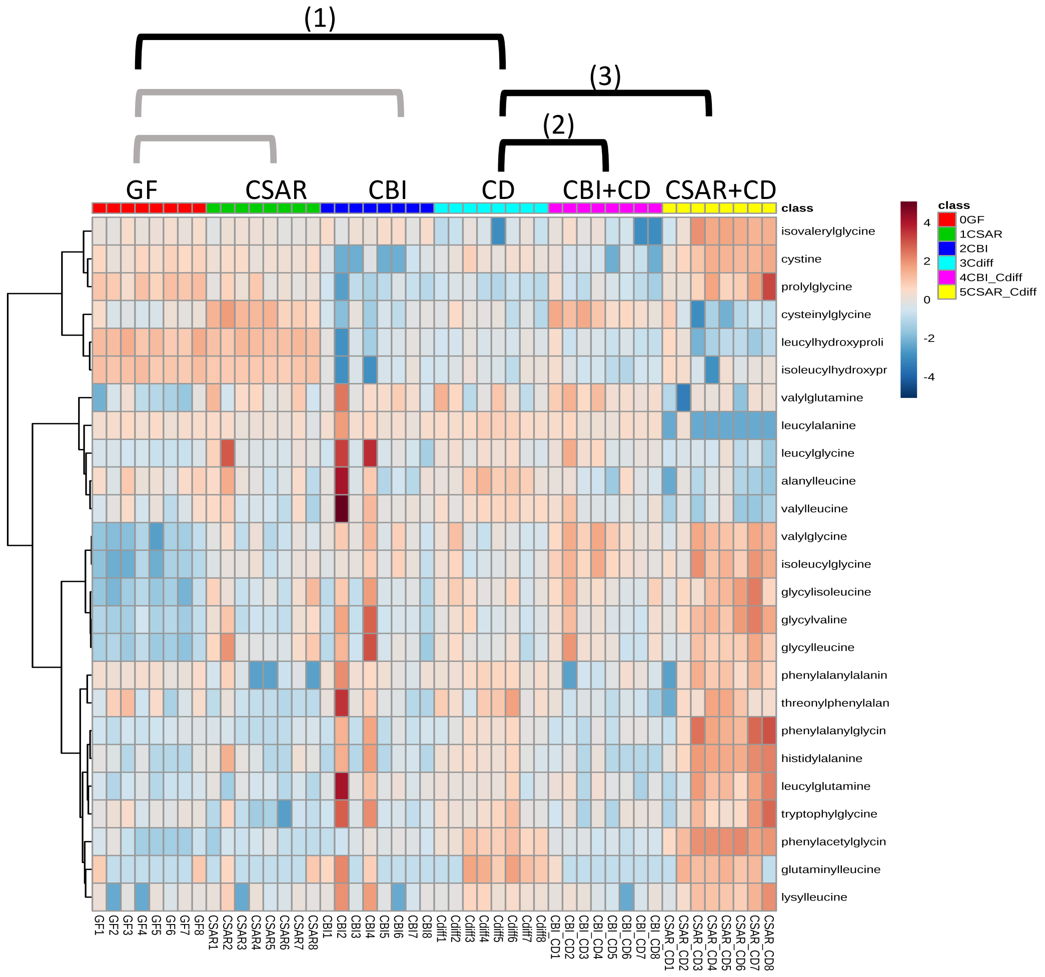


**Significantly Enriched Conditions:**

(1) Enriched in *C. difficile* mono-colonized vs GF mice at 20h (p=1.08E-02).

(2) Enriched in *C. difficile* mono-colonized vs CBI+*C. difficile* co-colonized mice at 20h (p=2.99E-02).

(3) Enriched in CSAR+*C. difficile* co-colonized mice vs *C. difficile* mono-colonized mice at 20h (p=4.19E-02).

### 2.5 Ethanolamide Endocannabinoids

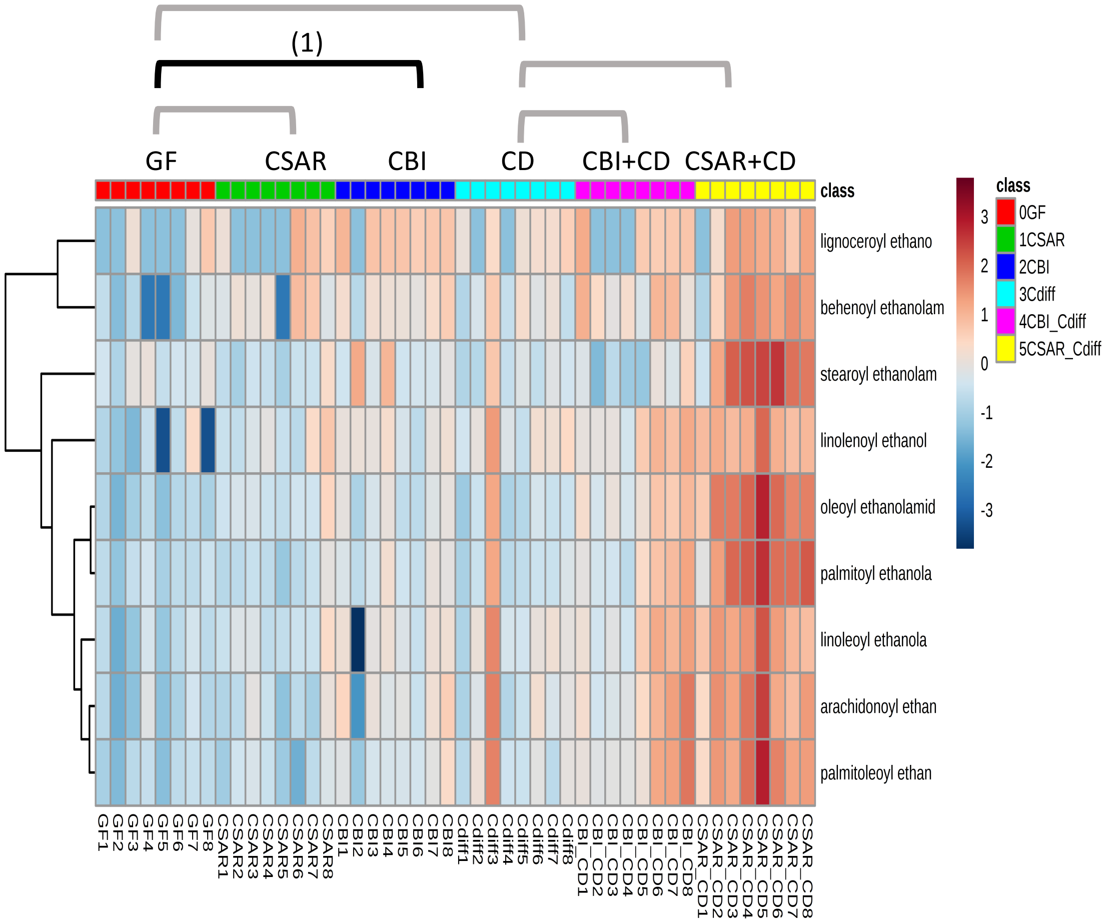


**Significantly Enriched Conditions:**

(1) Enriched in CBI mono-colonized vs GF mice at 7d (p=4.57E-02).

### 2.6 Fatty Acid Dicarboxylates

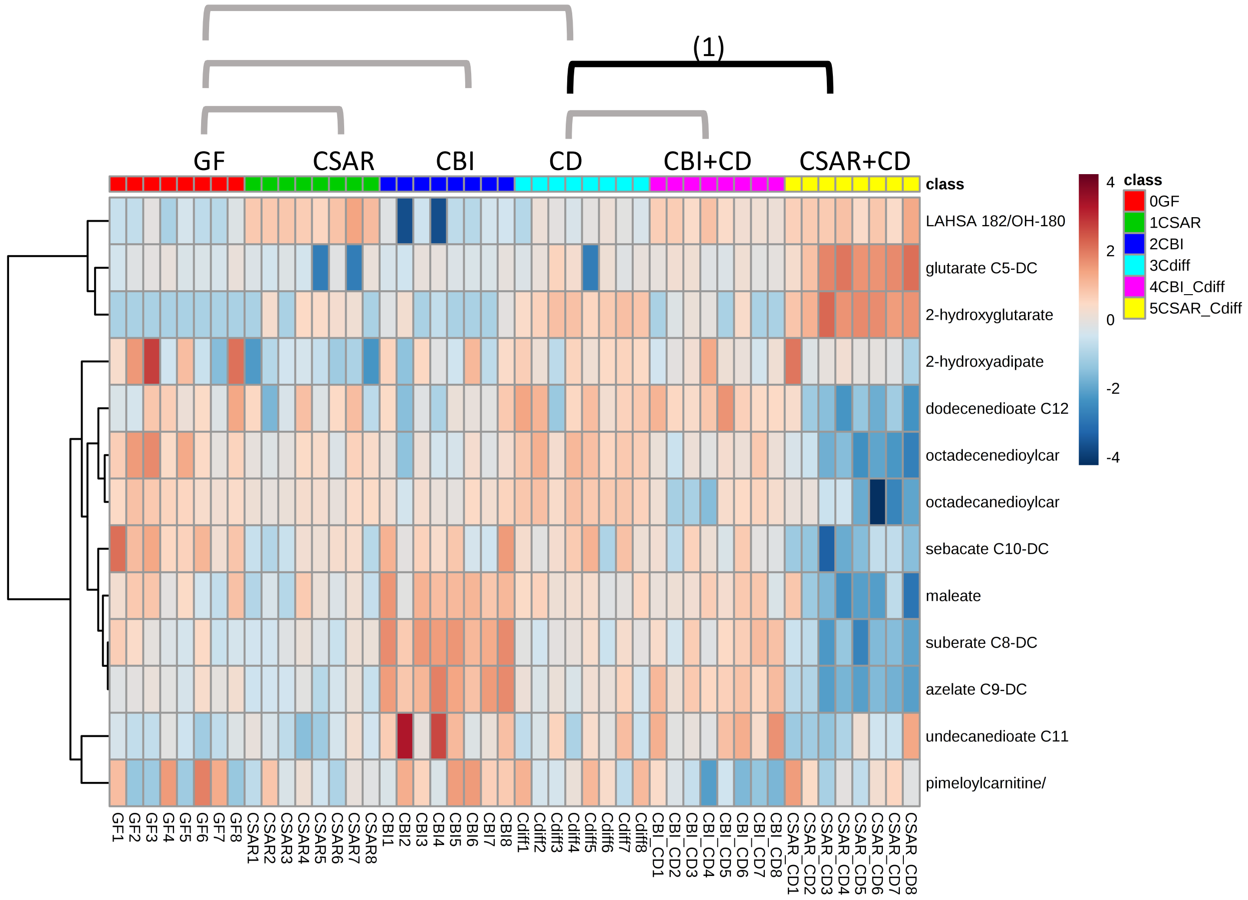


**Significantly Enriched Conditions:**

(1) Enriched in *C. difficile* mono-colonized vs CSAR+*C. difficile* co-colonized mice at 20h (p=1.25E-05).

### 2.7 Fatty Acid Intermediates

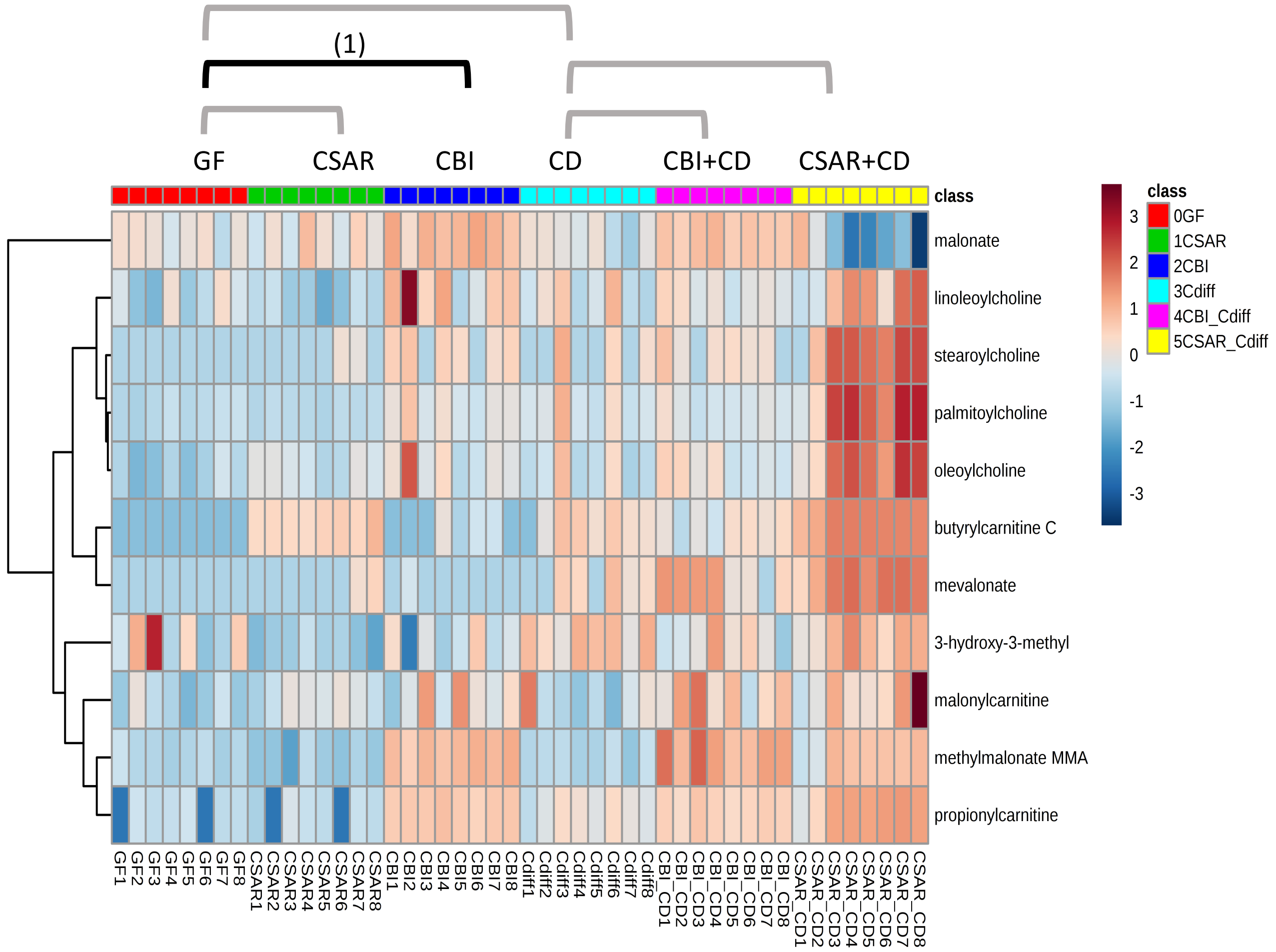


**Significantly Enriched Conditions:**

(1) Enriched in CBI mono-colonized vs GF mice at 20h (p=1.02E-02).

### 2.8 Gamma-glutamyl Amino Acids

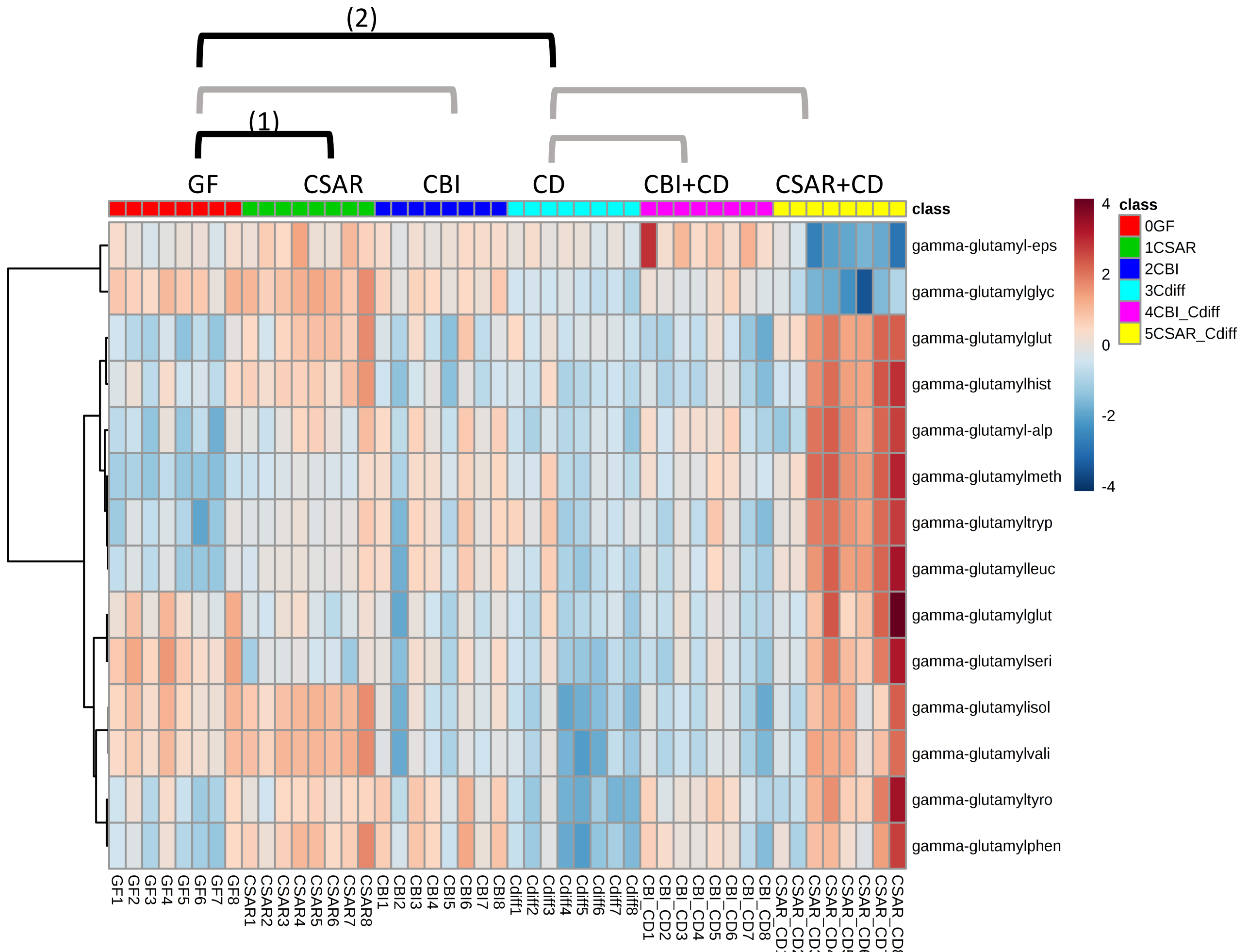


**<u>Significantly Enriched Conditions:</u>**

(1) Enriched in CSAR mono-colonized vs GF mice at 7d (p=3.61E-03).

(2) Enriched in GF vs *C. difficile* mono-colonized mice at 20h (p=8.99E-03).

### 2.9 Hexoses

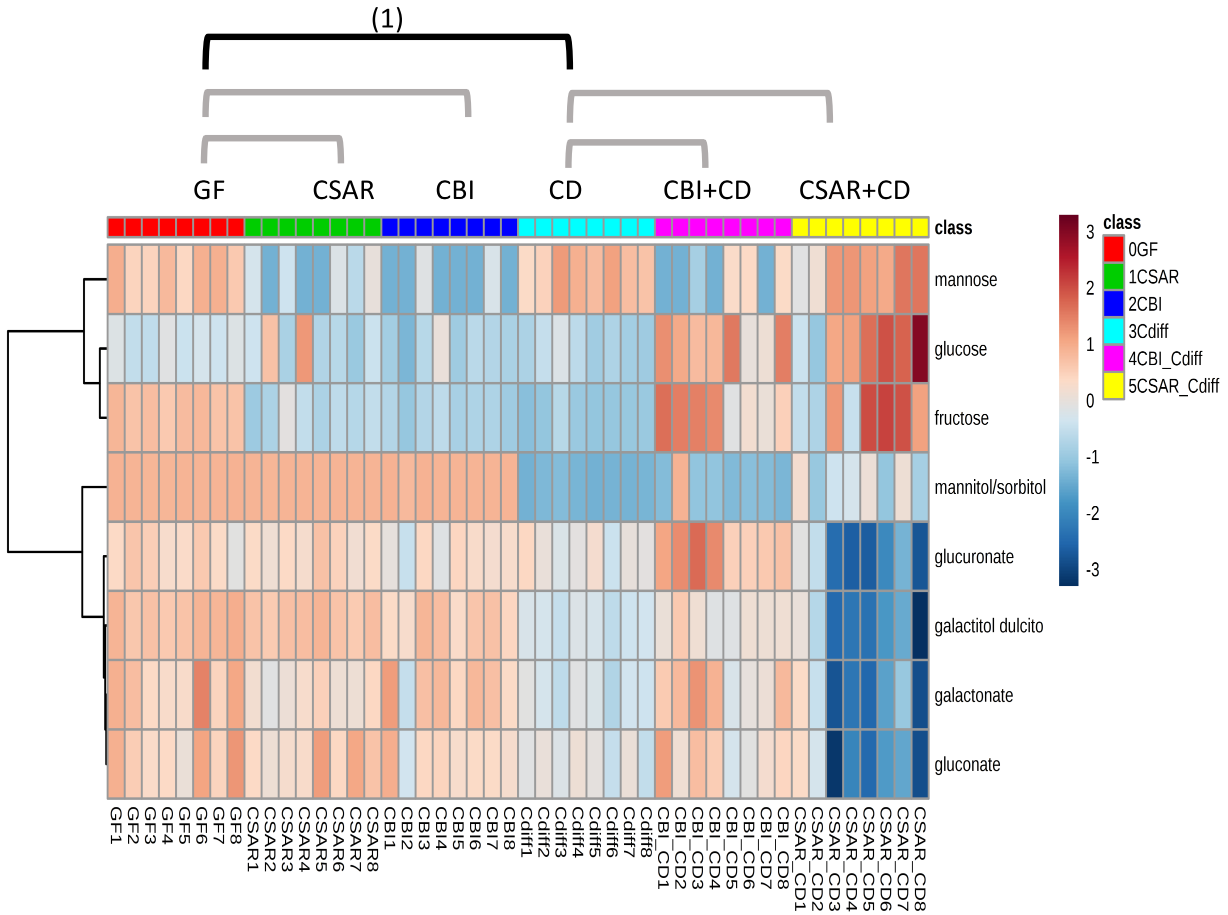


**<u>Significantly Enriched Conditions:</u>**

(1) Enriched in GF vs *C. difficile* mono-colonized mice at 20h (p=1.23E-03).

### 2.10 N-acetyl Amino Acids

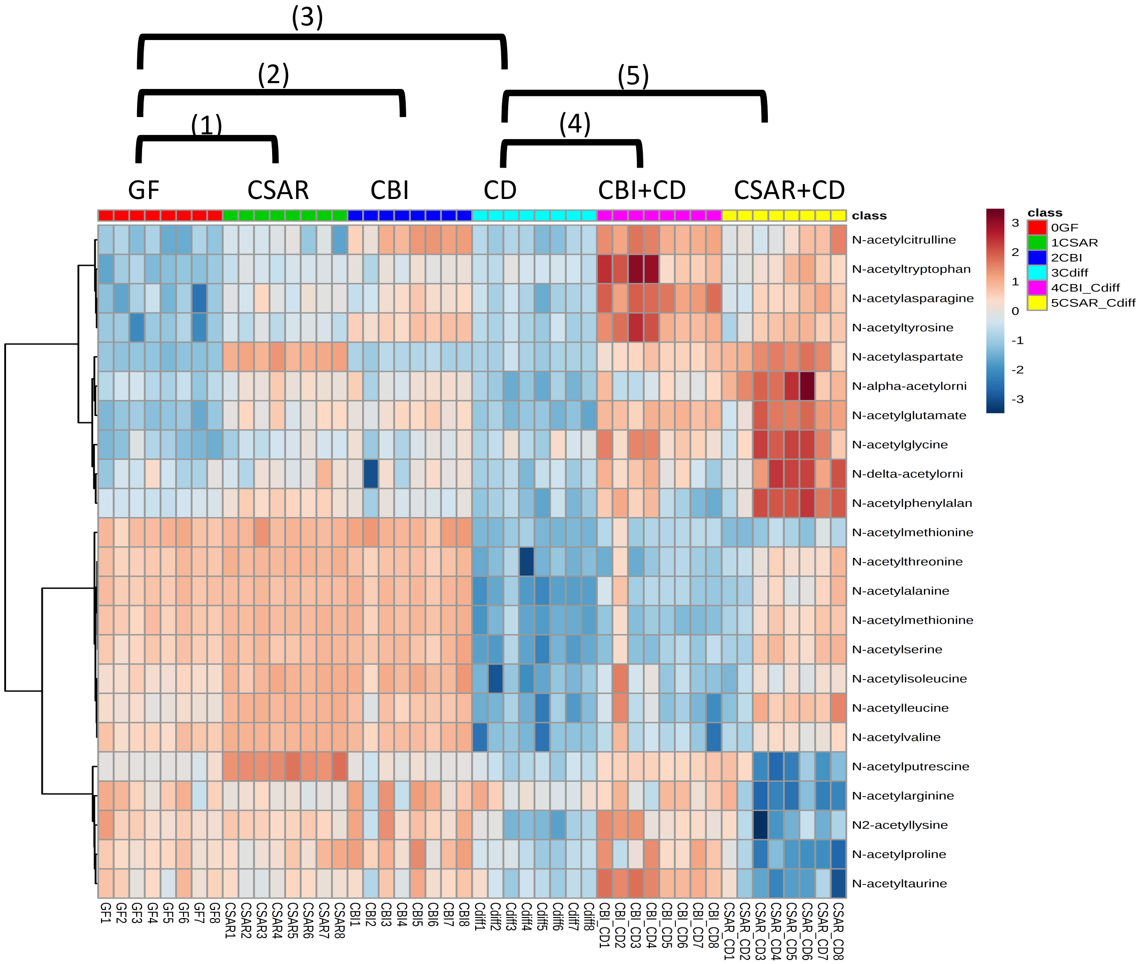


**<u>Significantly Enriched Conditions:</u>**

(1) Enriched in CSAR mono-colonized vs GF mice at 7d (p=6.10E-04).

(2) Enriched in CBI mono-colonized vs GF mice at 7d (p=2.18E-02).

(3) Enriched in GF vs *C. difficile* mono-colonized mice at 20h (p=1.78E-05).

(4) Enriched in CBI+ *C. difficile* co-colonized mice vs *C. difficile* mono-associated mice at 20h (p=1.38E-06).

(5) Enriched in CSAR+ *C. difficile* co-colonized mice vs *C. difficile* mono-associated mice at 20h (p=8.13E-03).

### 2.11 Other Fermentable Amino Acids

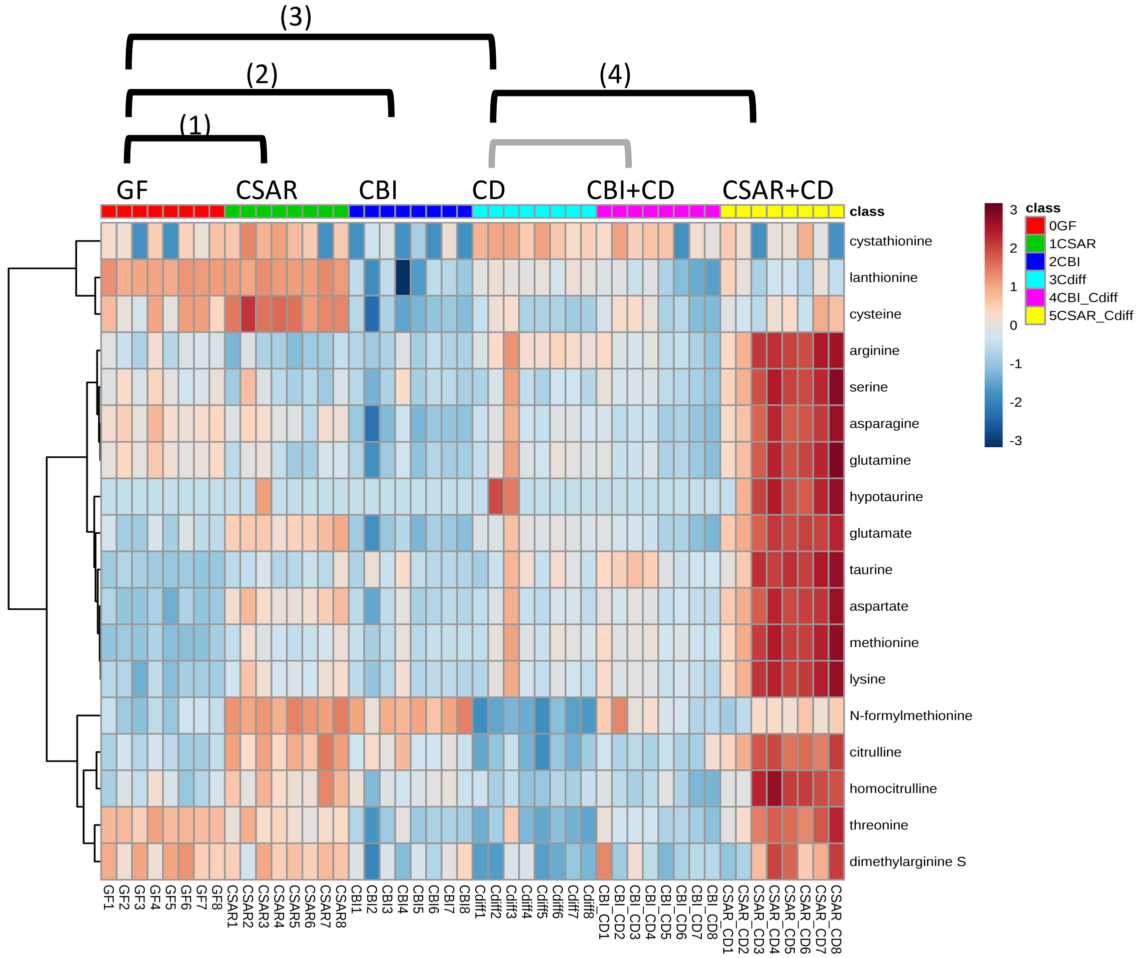


**Significantly Enriched Conditions:**

(1) Enriched in CSAR mono-colonized vs GF mice at 7d (p=2.36E-02).

(2) Enriched in GF vs CBI mono-colonized mice at 7d (p=4.30E-04).

(3) Enriched in GF vs *C. difficile* mono-colonized mice at 20h (p=3.44E-02).

(4) Enriched in CSAR+ *C. difficile* co-colonized mice vs *C. difficile* mono-associated mice at 20h (p=6.61E-03).

### 2.12 Pentoses

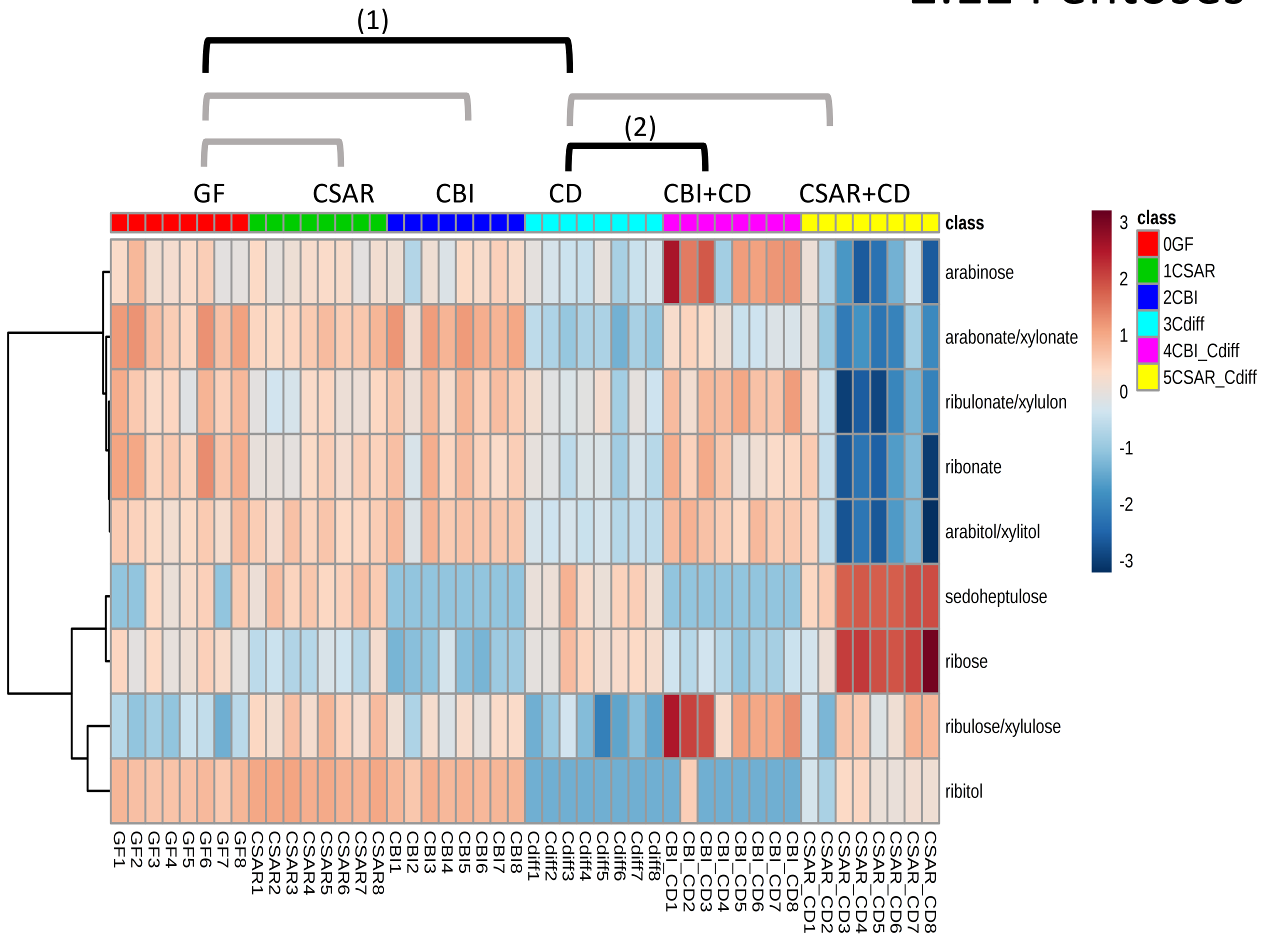


**Significantly Enriched Conditions:**

(1) Enriched in GF vs *C. difficile* mono-colonized mice at 20h (p=1.23E-03).

(2) Enriched in CBI+ *C. difficile* co-colonized mice vs *C. difficile* mono-associated mice at 20h (p=6.82E-03).

### 2.13 Phosphatidylcholines (PC)

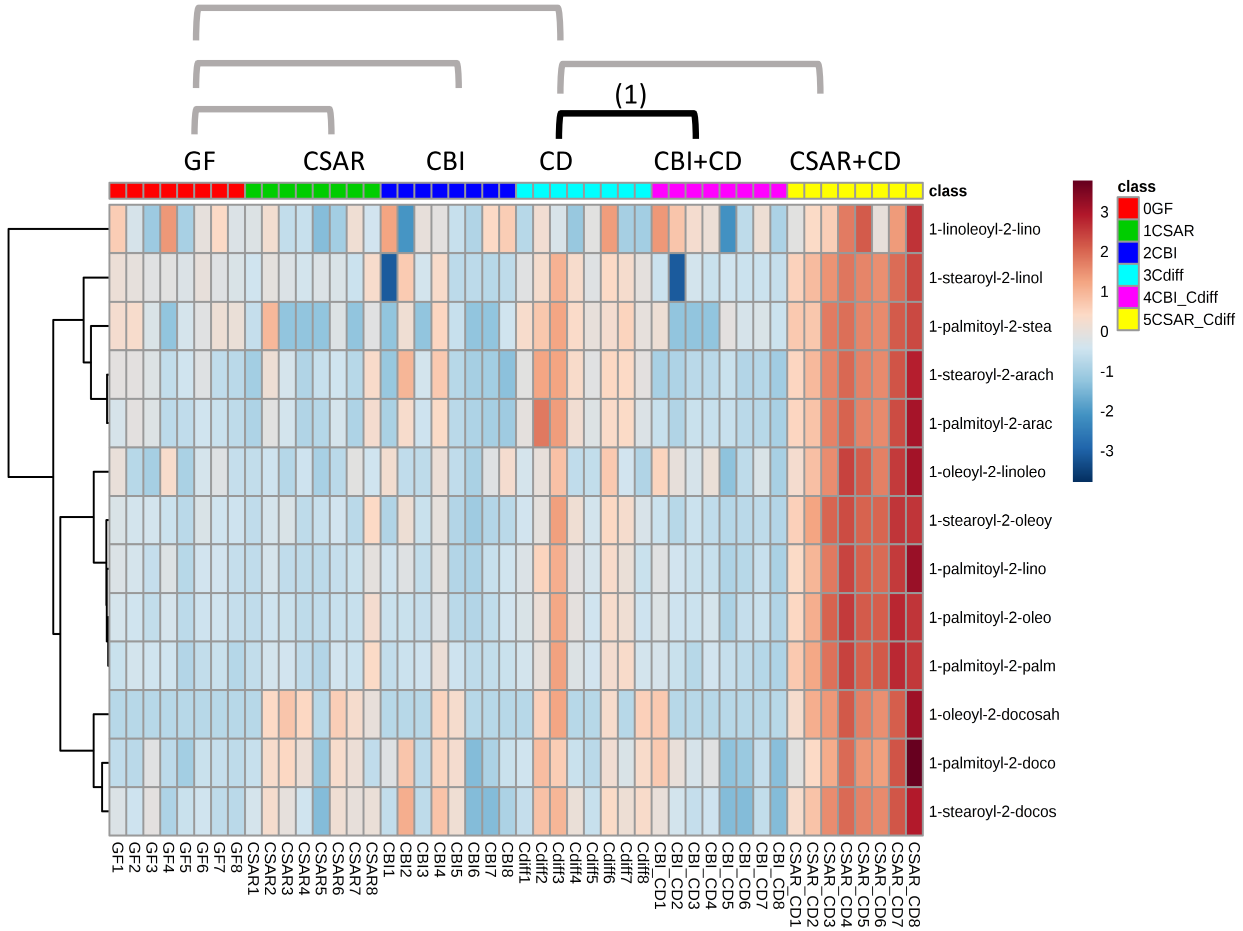


**Significantly Enriched Conditions:**

(1) Enriched in *C. difficile* mono-colonized mice vs CBI+ *C. difficile* co-colonized mice at 20h (p=9.86E-04).

Levels of enrichment at the threshold cutoffs did not attain significance between *C. difficile*-monocolonized and CSAR-co-colonized mice

### 2.14 Phosphatidylethanolamines (PE)

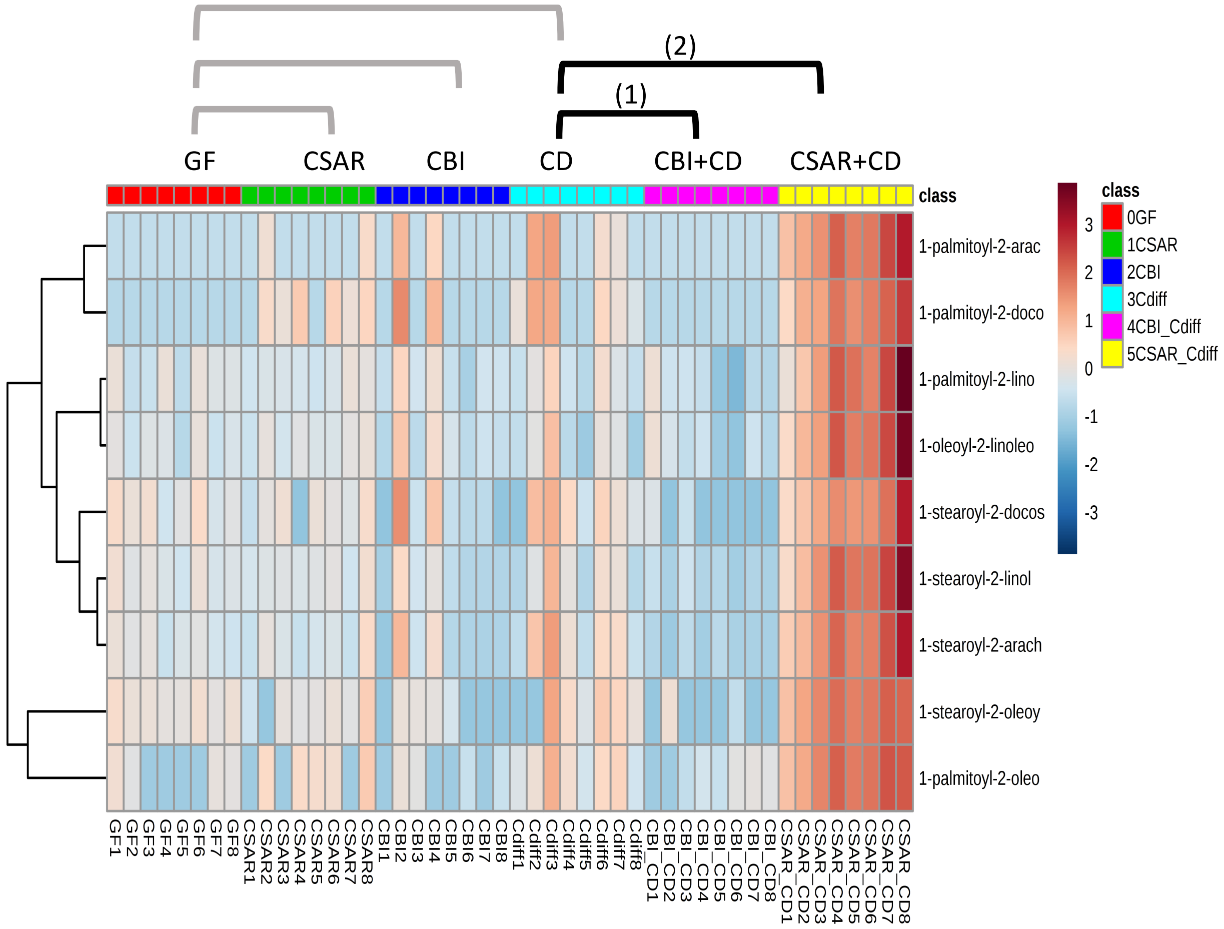


**<u>Significantly Enriched Conditions:</u>**

(1) Enriched in *C. difficile* mono-associated mice vs CBI+ *C. difficile* co-colonized mice at 20h (p=3.36E-02).

(2) Enriched in CSAR+ *C. difficile* co-colonized mice vs *C. difficile* mono-associated mice at 20h (p=3.29E-02).

### 2.15 Plasmalogens

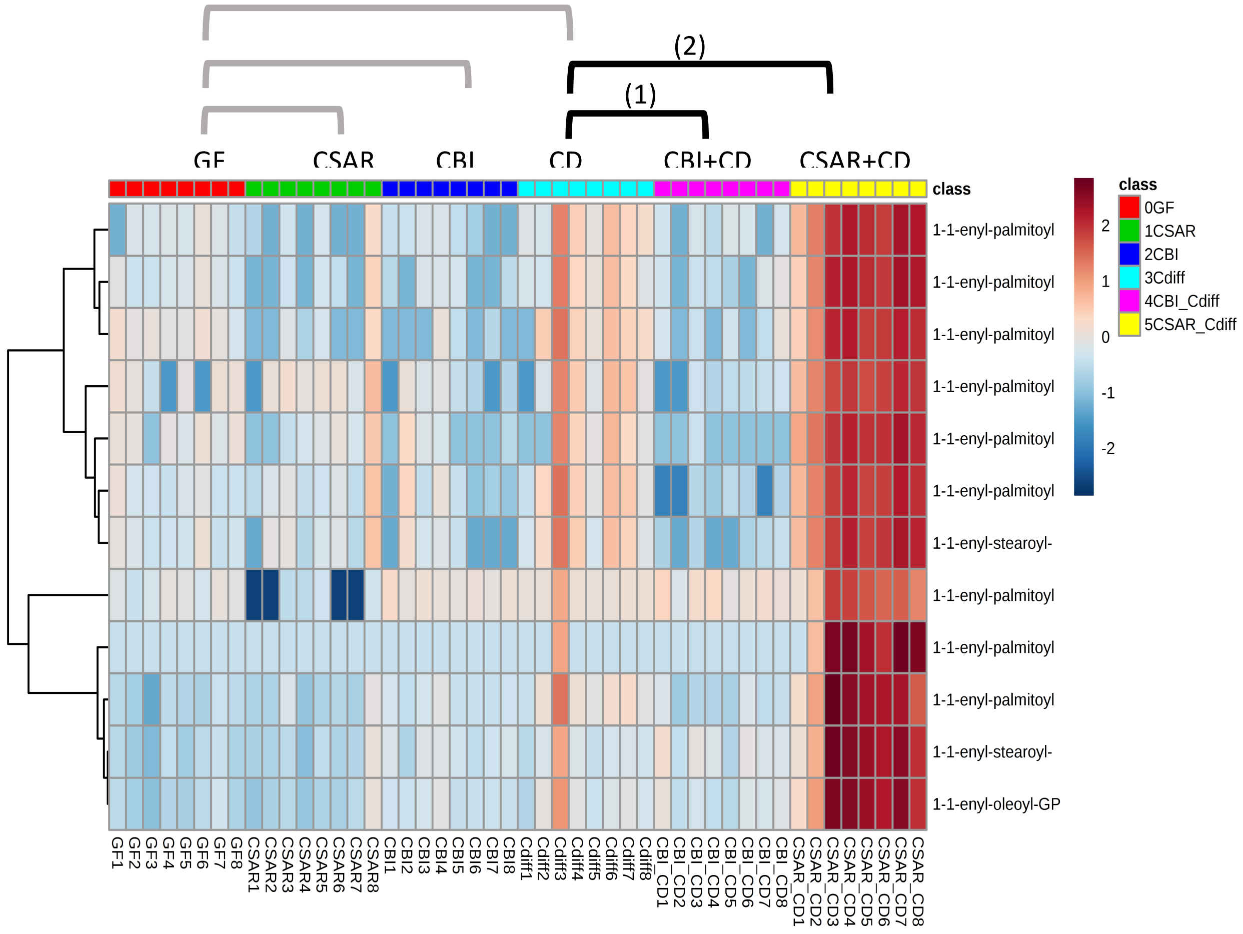


**Significantly Enriched Conditions:**

(1) Enriched in *C. difficile* mono-associated mice vs CBI+ *C. difficile* co-colonized mice at 20h (p=1.72E-03).

(2) Enriched in CSAR+ *C. difficile* co-colonized mice vs *C. difficile* mono-associated mice at 20h (p=1.64E-04).

### 2.16 Primary Bile Acids

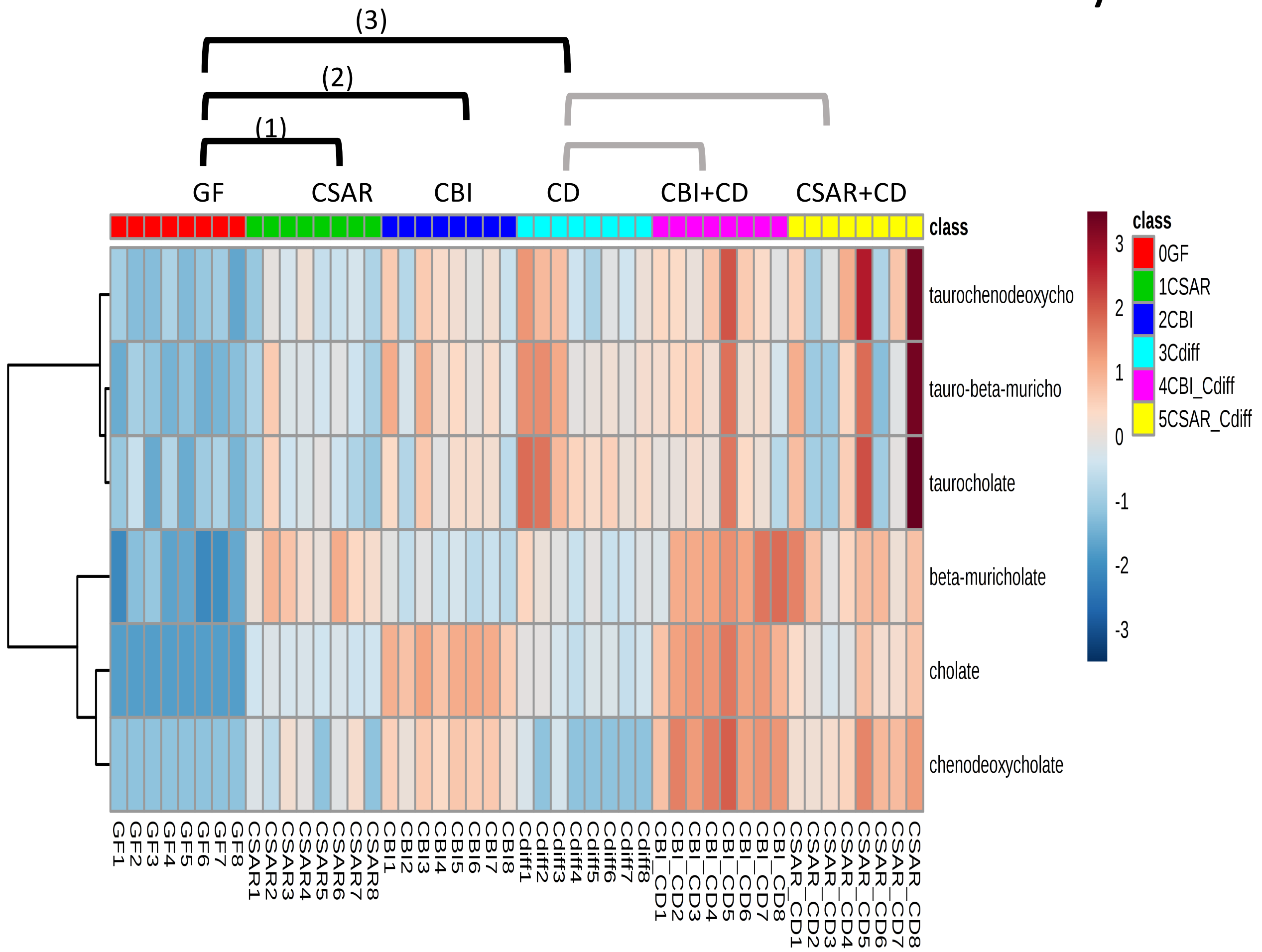


<u>**Significantly Enriched Conditions:**</u>

(1) Enriched in CSAR mono-colonized vs GF mice at 7d (p=1.08E-03).

(2) Enriched in GF vs CBI mono-colonized mice at 7d (p=4.14E-03).

(3) Enriched in *C. difficile* mono-colonized vs GF mice at 20h (p=1.52E-02).

### 2.17 Purines-Xanthines and Metabolites

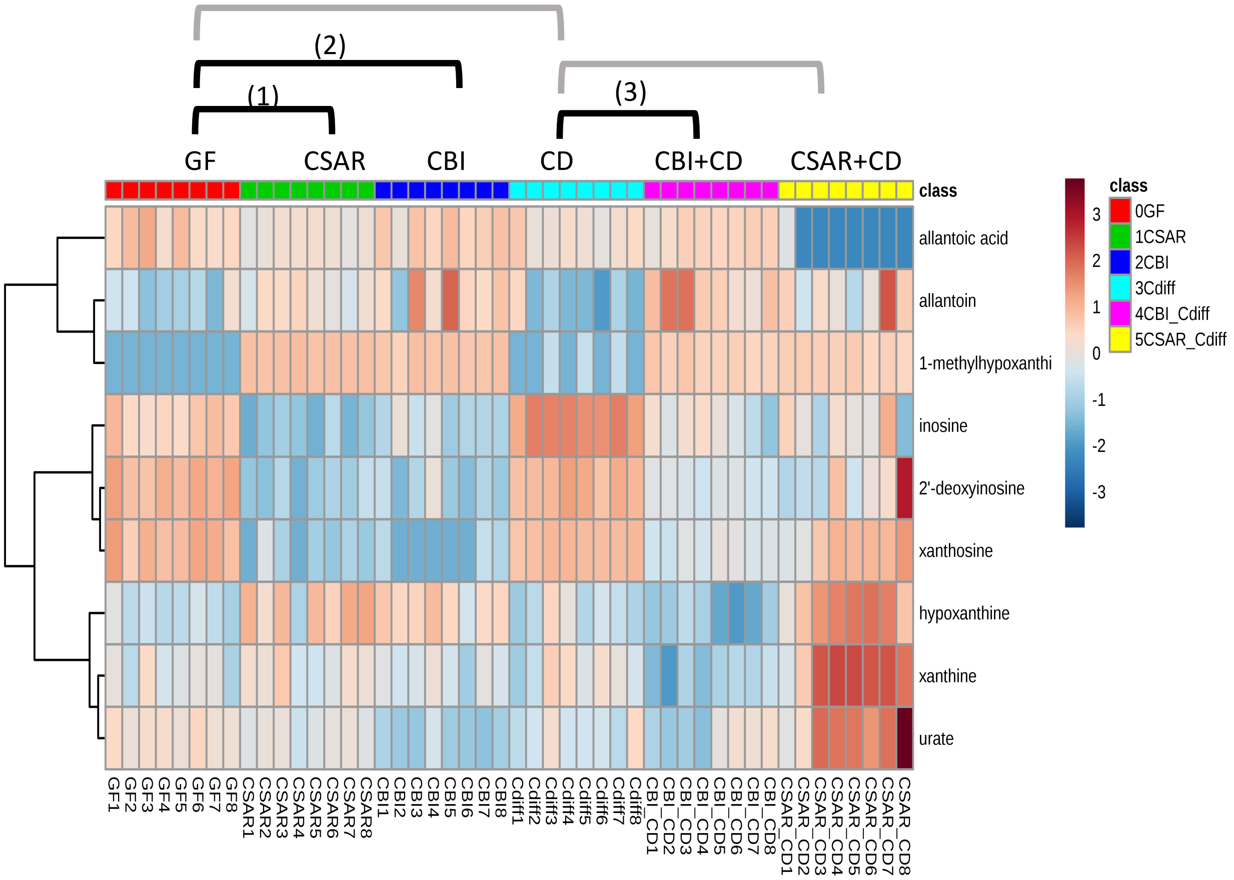


**Significantly Enriched Conditions:**

(1) Enriched in GF vs CSAR mono-colonized mice at 7d (p=2.03E-03).

(2) Enriched in GF vs CBI mono-colonized mice at 7d (p=2.25E-02).

(3) Enriched in *C. difficile* mono-associated mice vs CBI+*C. difficile* co-colonized mice vs at 20h (p=3.35E-02).

### 2.18 Pyrimidines - Cytidines

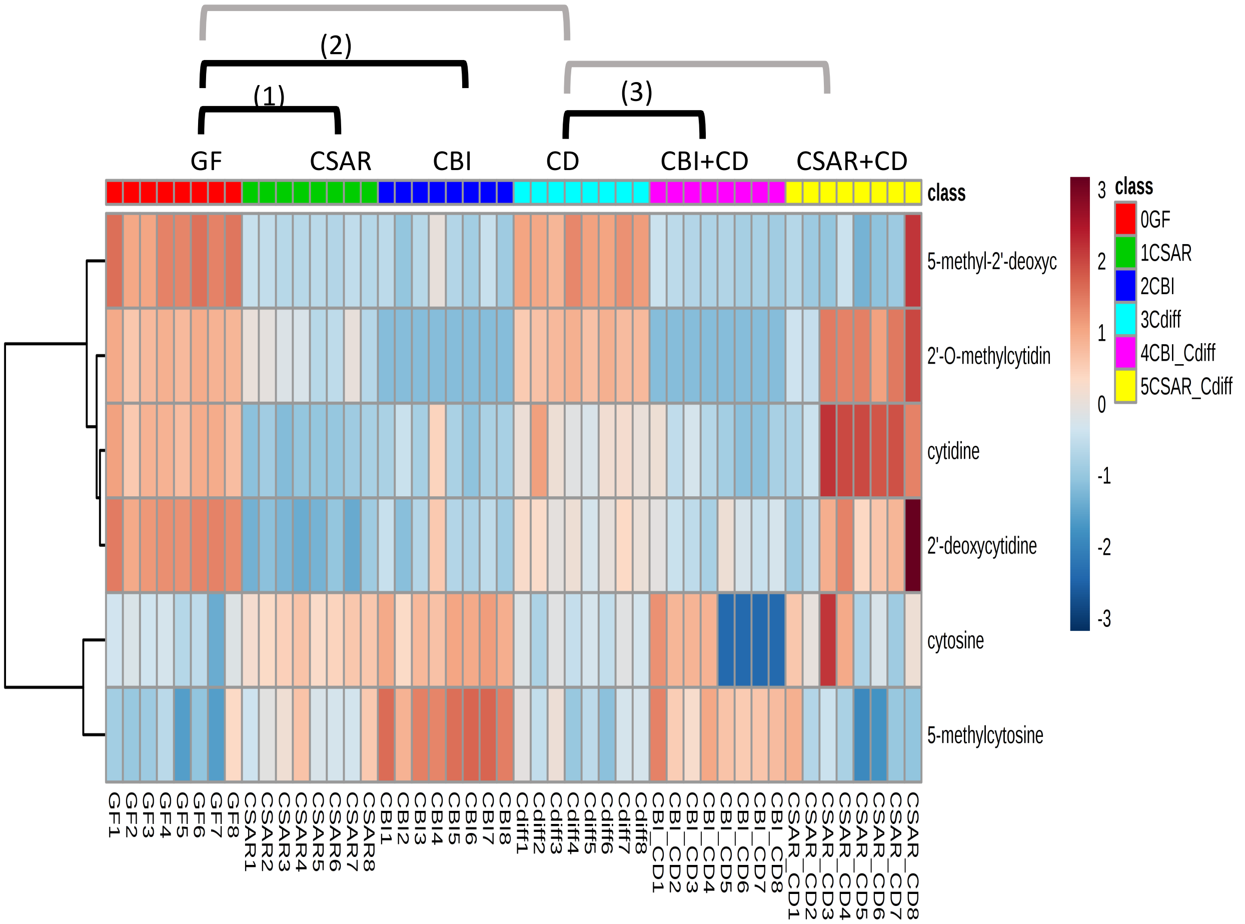


**Significantly Enriched Conditions:**

(1) Enriched in GF vs CSAR mono-colonized mice at 7d (p=1.06E-03).

(2) Enriched in GF vs CBI mono-colonized mice at 7d (p=6.47E-03).

(3) Enriched in *C. difficile* mono-associated mice vs CBI+*C. difficile* co-colonized mice vs at 20h (p=2.58E-02).

### 2.19 Pyrimidines – Thymine and Uracil Containing Compounds

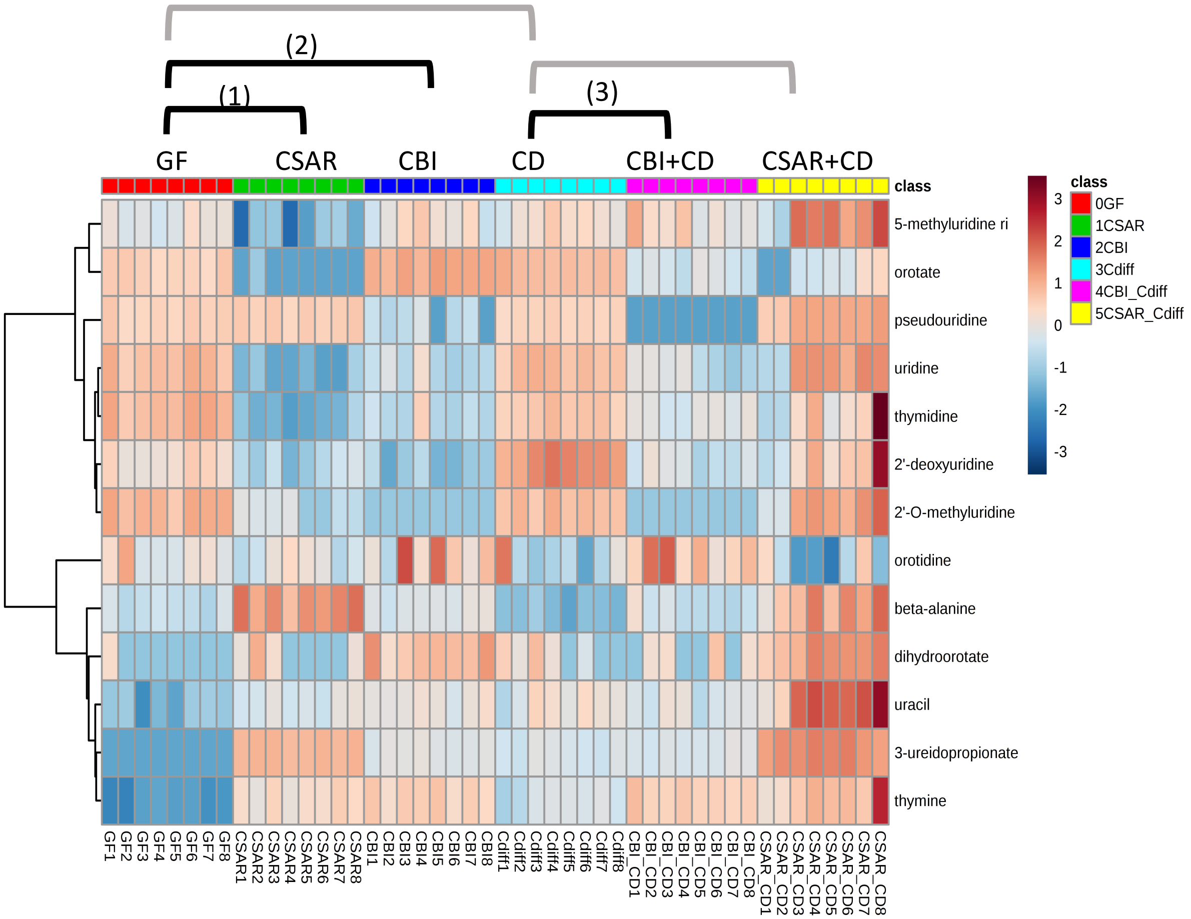


**Significantly Enriched Conditions:**

(1) Enriched in GF vs CSAR mono-colonized mice at 7d (p=1.51E-03).

(2) Enriched in GF vs CBI mono-colonized mice at 7d (p=2.13E-02).

(3) Enriched in *C. difficile* mono-associated mice vs CBI+*C. difficile* co-colonized mice vs at 20h (p=2.76E-02).

### 2.20 SCFA

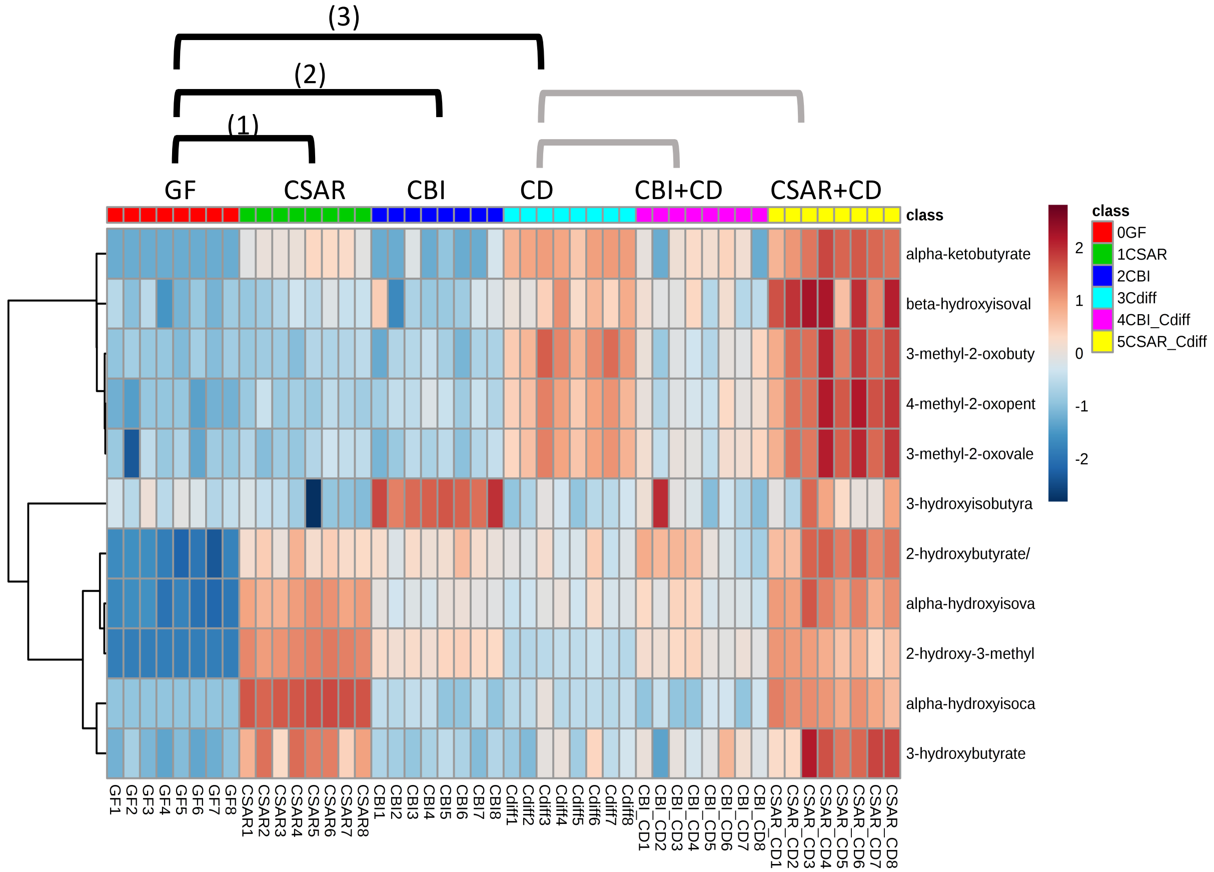


**<u>Significantly Enriched Conditions:</u>**

(1) Enriched in CSAR mono-colonized vs GF mice at 7d (p=2.84E-02).

(2) Enriched in CBI mono-colonized vs GF mice at 7d (p=1.82E-02).

(3) Enriched in *C. difficile* mono-associated vs GF mice at 20h (p=2.76E-04).

### 2.21 SCFA (Non-Volatile)

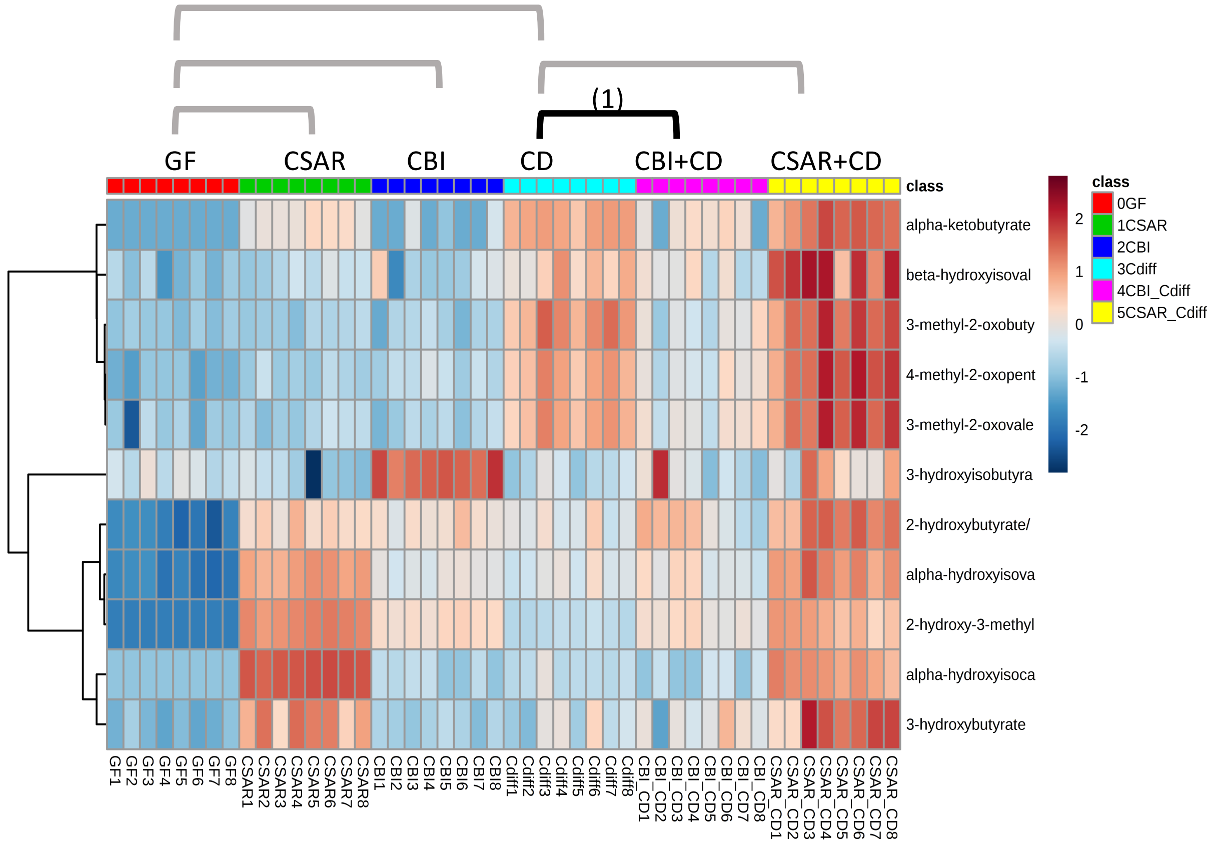


**<u>Significantly Enriched Conditions:</u>**

(1) Enriched in *C. difficile* mono-associated vs CBI+*C. difficile* co-colonized mice at 20h (p=4.60E-02).

### 2.22 Sphingosine Containing Compounds

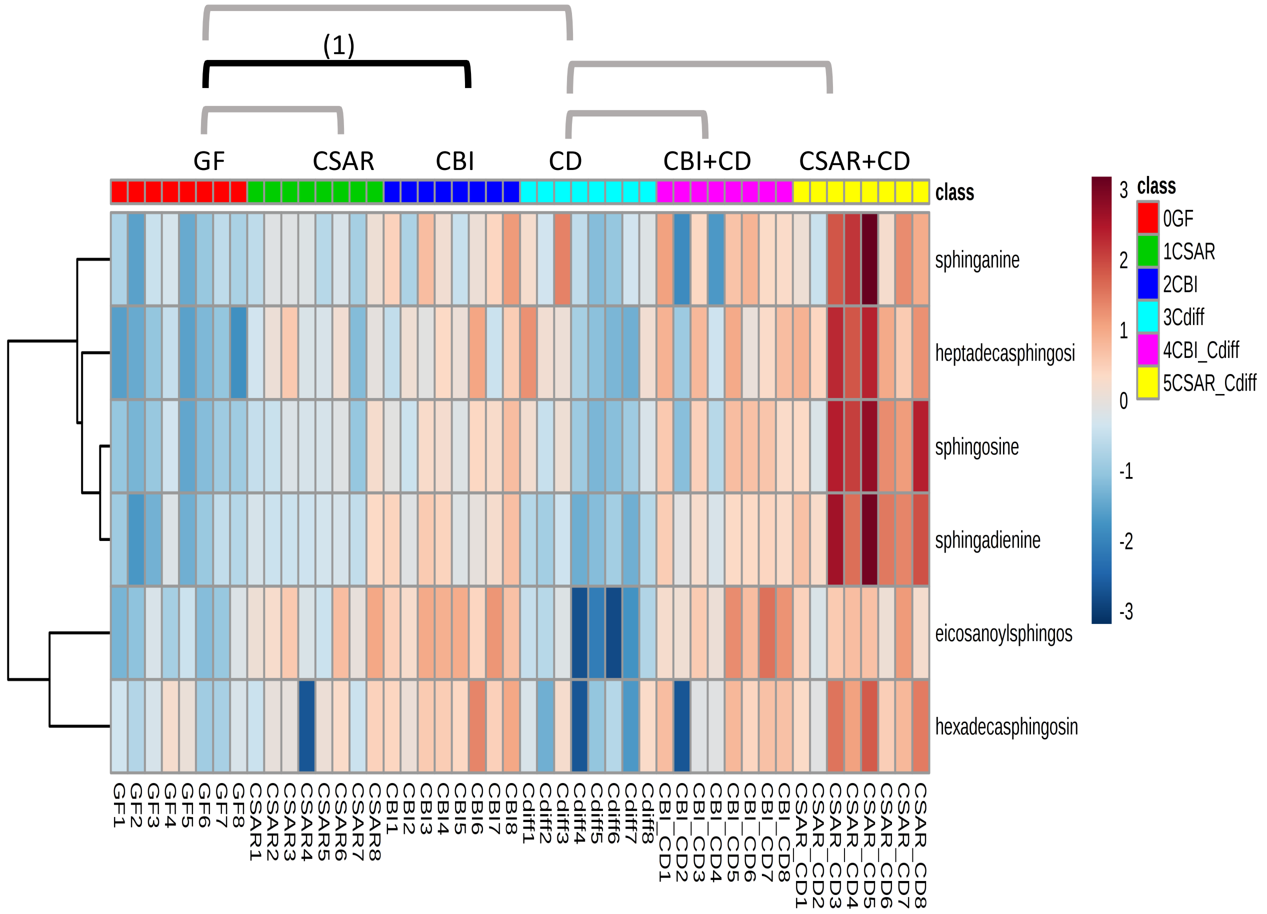


**Significantly Enriched Conditions:**

(1) Enriched in CBI mono-colonized vs GF mice at 7d (p=4.41E-03).

### 2.23 Stickland Acceptor Amino Acids

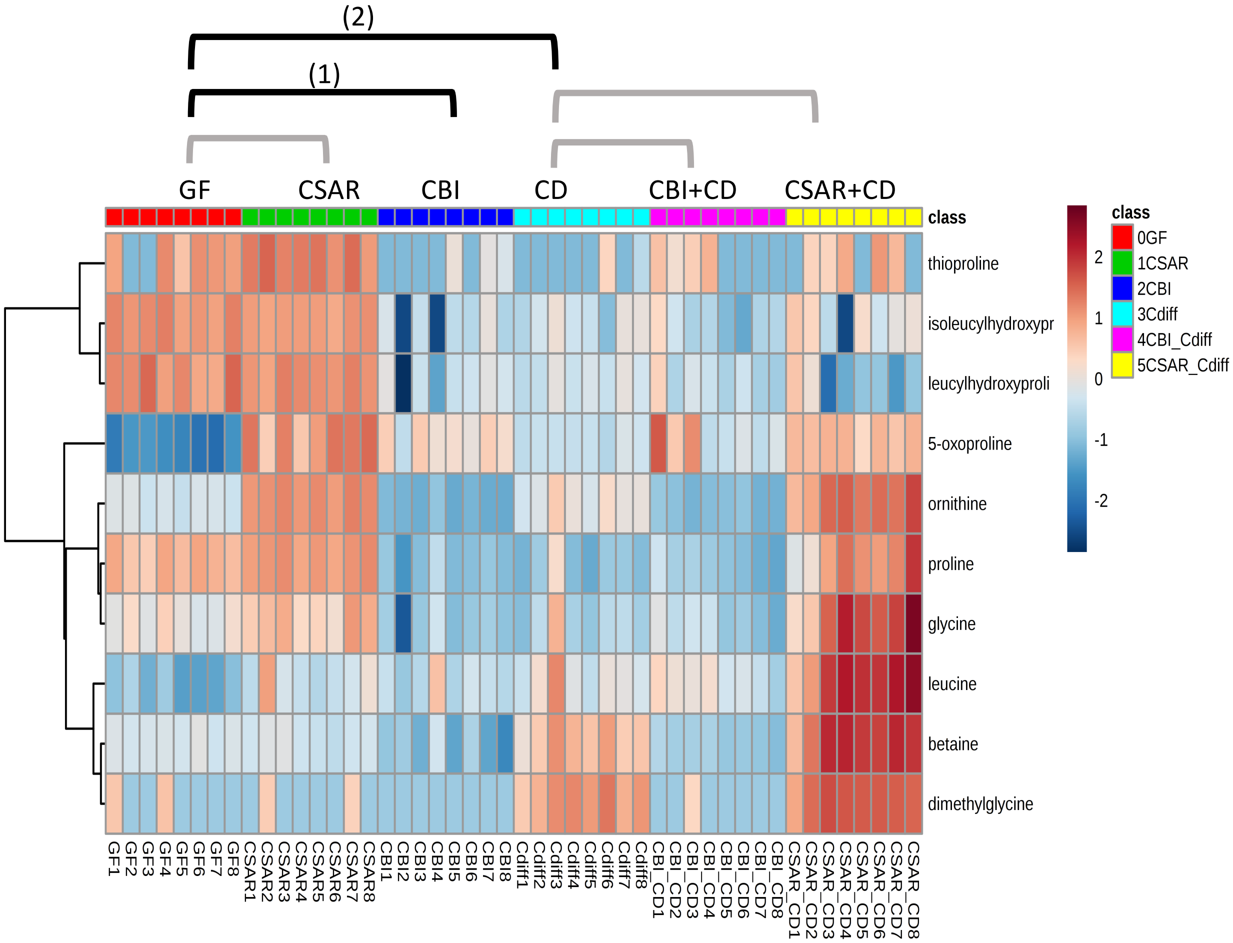


**Significantly Enriched Conditions:**

(1) Enriched in GF vs CBI mono-colonized mice at 7d (p=2.06E-04).

(2) Enriched in GF vs *C. difficile* mono-colonized mice at 20h (p=2.06E-04).

### 2.24 Stickland Donor Amino Acids

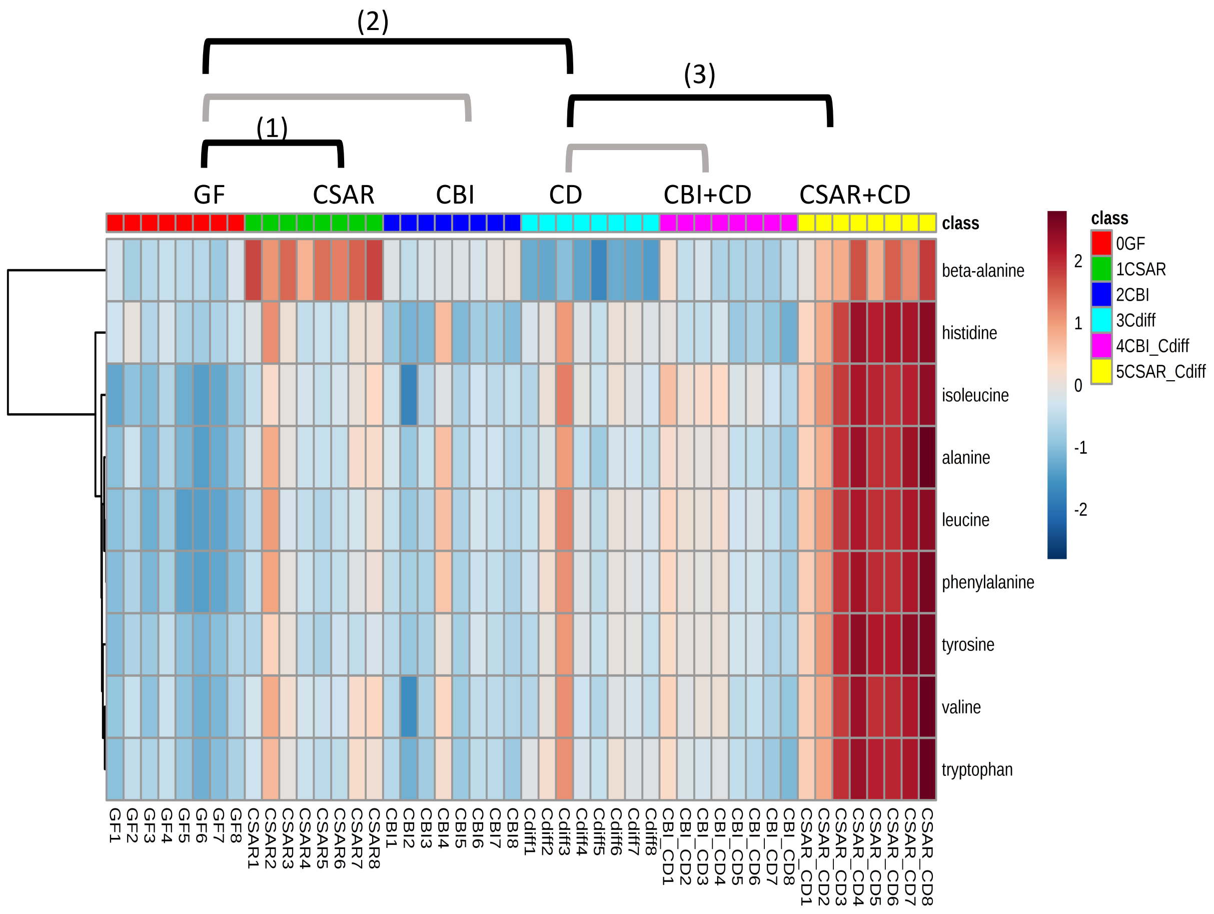


**Significantly Enriched Conditions:**

(1) Enriched in CSAR mono-colonized mice vs GF at 7d (p=5.16E-04).

(2) Enriched in *C. difficile* mono-colonized vs GF mice at 20h (p=3.44E-03).

(3) Enriched in CSAR+*C. difficile* co-colonized vs *C. difficile* mono-colonized mice at 20h (p=4.18E-04).

### 2.25 Sugar Alcohols and Acids

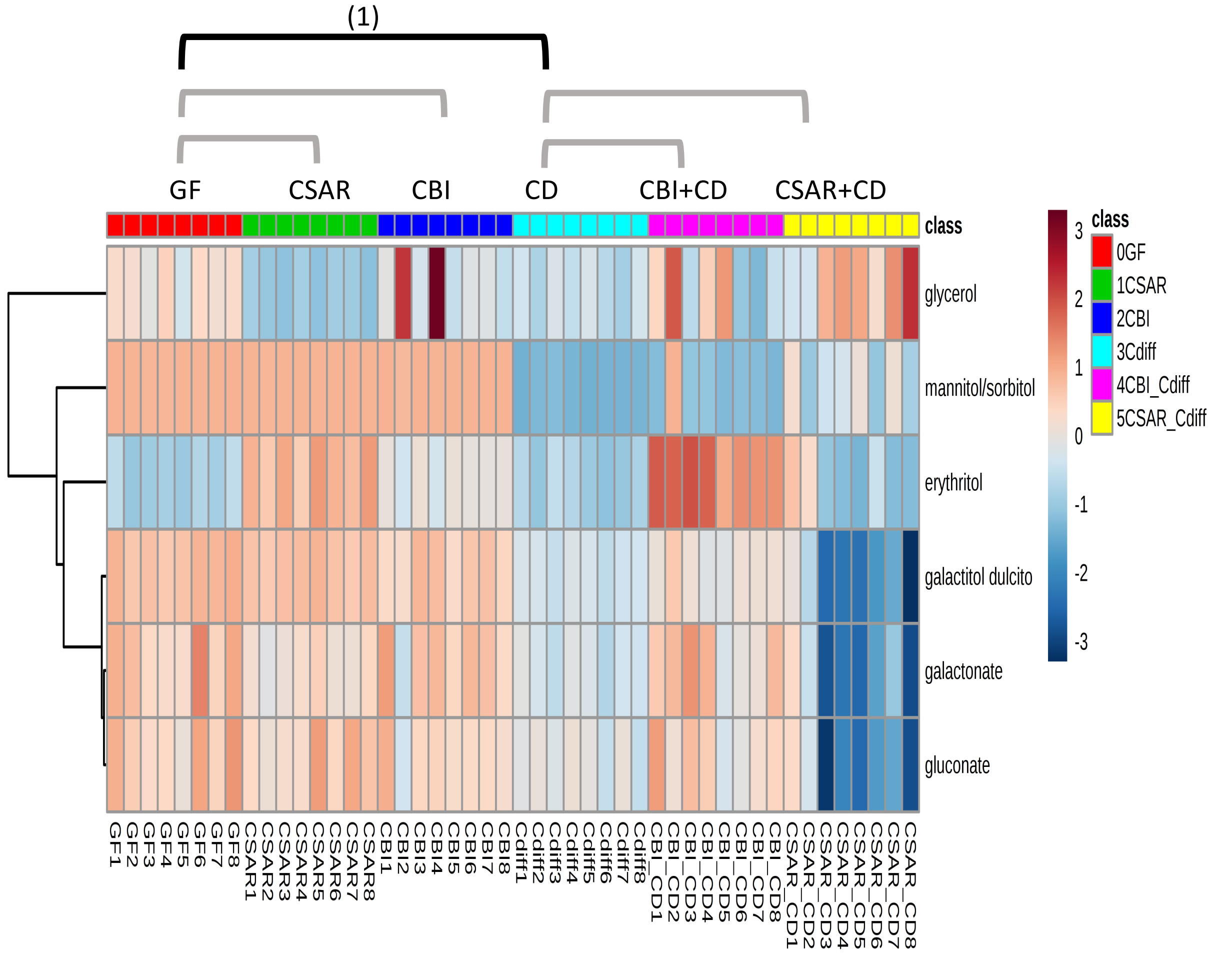


**Significantly Enriched Conditions:**

(1) Enriched in GF vs *C. difficile* mono-colonized mice at 20h (p=3.24E-03).

### 2.26 Vitamins and Cofactors

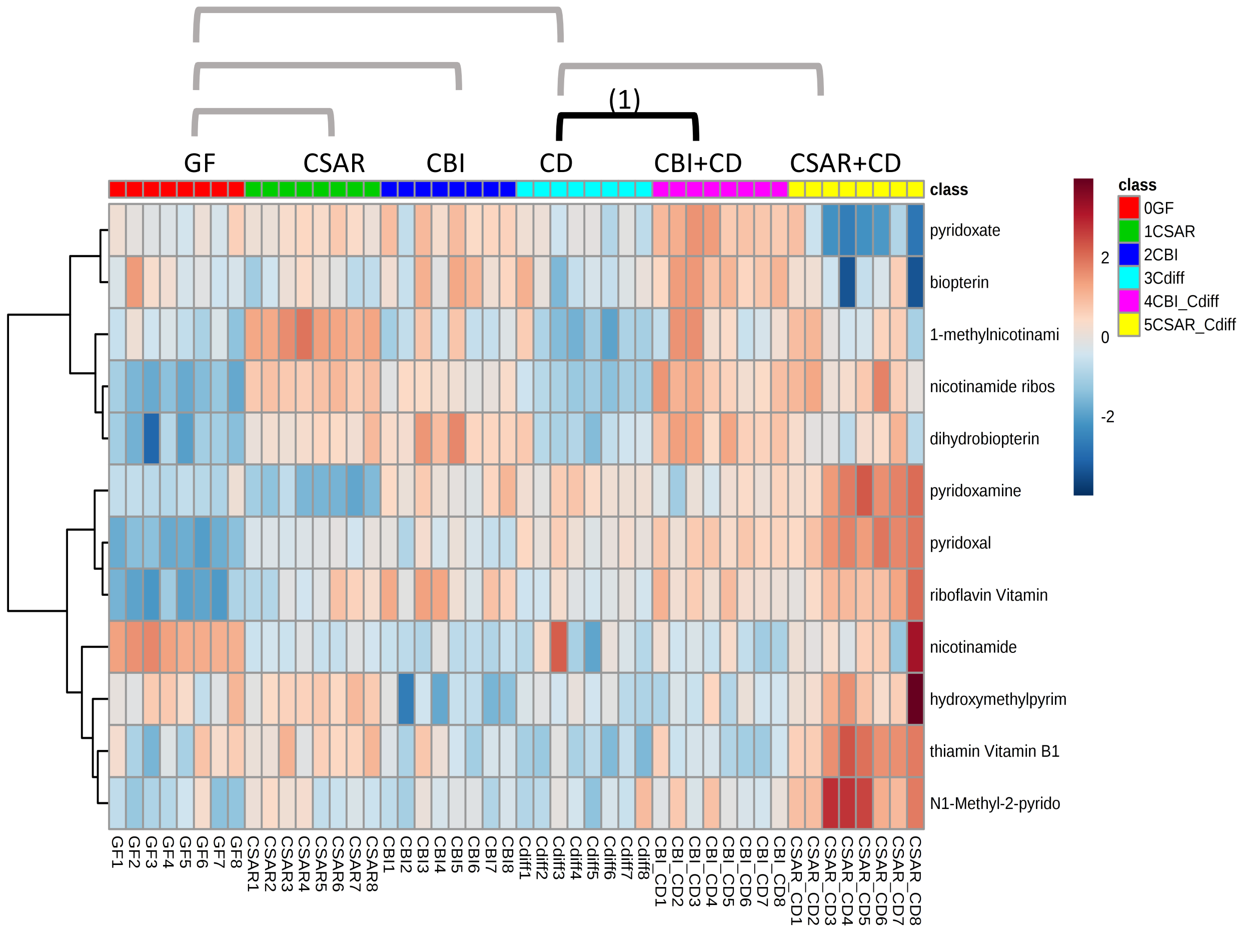


**Significantly Enriched Conditions:**

(1) Enriched in CBI+*C. difficile* co-colonized vs *C. difficile* mono-colonized mice at 20h (p=7.91E-03).
