## Supplemental Data File 5 for "*In vivo* commensal control of *Clostridioides difficile* virulence"

- Version 1.0; 12/23/2019

### Supplemental Data File 5: Transcriptome Analyses: *C. difficile* Enriched Pathways

**Pages:** Introduction, this page
**Page 1**: Sample Key


**Carbon Source Metabolism**
2: 5.1 Aminobutanoate Degradation
3: 5.2 Beta-glucoside Transport and Metabolism
4: 5.3 Butanoate Metabolism
5: 5.4 Dipeptide and Oligopeptide Transport
6: 5.5 Disaccharide Transport and Metabolism
7: 5.6 Ethanolamine Utilization
8: 5.7 Fructose Transport and Metabolism
9: 5.8 Glucose and Maltose Transport and Metabolism
10: 5.9 Mannose, Sorbose and Fructose Transport
11: 5.10 Ornithine Fermentation
12: 5.11 Polysaccharide Metabolism
13: 5.12 Pyruvate Metabolism
14: 5.13 Ribose Transport and Metabolism
15: 5.14 Stickland Fermentations
16: 5.15 Sugar Alcohol and Acids, Transport and Metabolism
17: 5.16 Transport Binding Proteins: Amino Acids and Amines
18: 5.17 Wood-Ljungdahl Pathway
19: 5.18 Xanthines – Transport and Metabolism

**Core Cellular Machinery**
20: 5.19 ATP Synthesis
21: 5.20 DNA Replication
22: 5.21 Protein Translation and Modification
23: 5.22 Ribosomal Structural Components and Modification
24: 5.23: Transcription: RNA synthesis and modification

**Cellular Adhesion and Motility**
25: 5.24 Cell Wall Turnover
26: 5.25 Gram+ Surface Anchored Components

**Vitamin and Micronutrient Metabolism**
27: 5.26 Cation Transport Binding Proteins
28: 5.27 Cobalamin Biosynthesis
29: 5.28 Folate Biosynthesis
30: 5.29 Nitrogenous Compounds: Transport and Metabolism
31: 5.30 Phosphorus and Phosphonate Transport and Metabolism
32: 5.31 Sulfur-containing Compounds: Transport and Metabolism

**Biosynthetic Pathways**
33: 5.32 Cysteine and Methionine Biosynthesis
34: 5.33 Fatty Acid Biosynthesis
35: 5.34 Histidine Biosynthesis
36: 5.35 Terpenoid Backbone Synthesis

**Other Cellular Transport Systems**
37: 5.36 ABC Transport System

**Stress Responses**
38: 5.37 Chemotaxis
39: 5.38 CISPR
40: 5.39 Diffocin Locus Proteins
41: 5.40 Oxidative Stress Responses
42: 5.41 Pathogenicity Islands/Virulence Determinants
43: 5.42 Sporulation

### SDF5: Sample Key

#### 5.13 Ornithine Fermentation

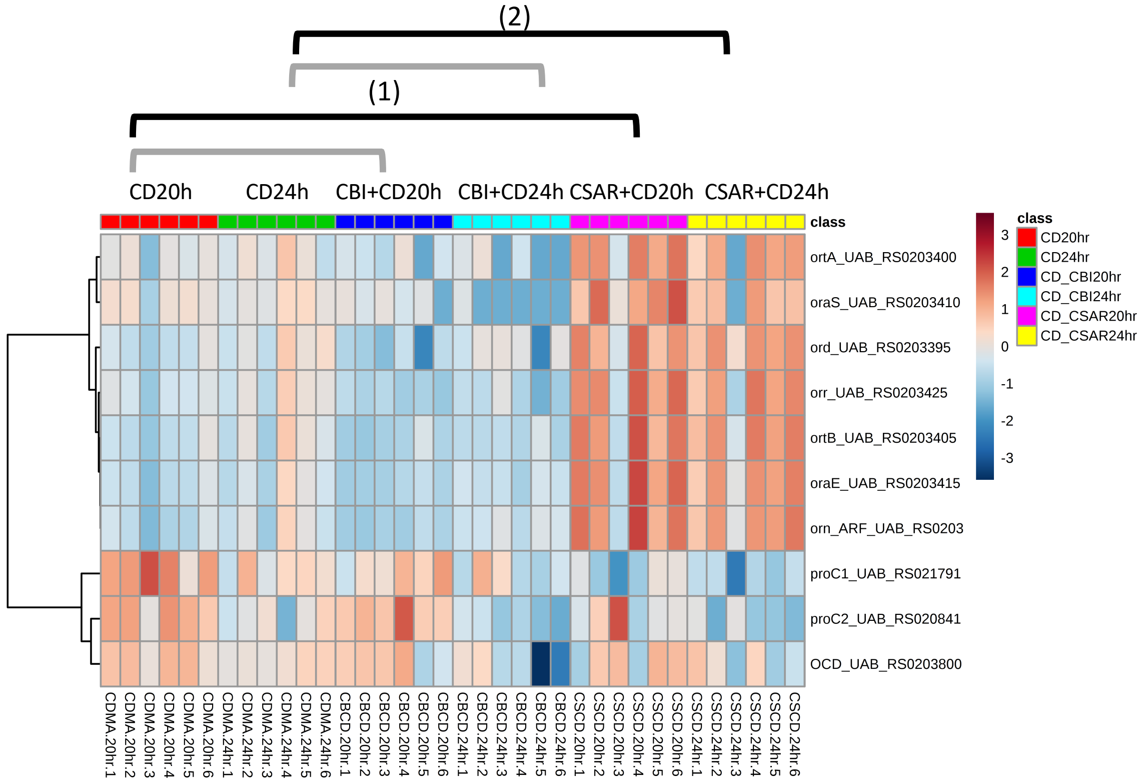


**<u>Guide:</u>**

- Group showing the enriched category is at the top (“5.13 Ornithine Fermentation”)
- Brackets indicate the 4 comparisons evaluated among groups for significant enrichment of genes in the pathway
- Brackets in black show significant comparisons
- Significant comparisons and p values are shown in the text box on the right-hand side

**<u>Significantly Enriched Conditions:</u>**

(1) Enriched in CSAR+*C. difficile* vs *C. difficile* mono-colonized mice at 20h (p=0.0050398).
(2) Enriched CSAR+*C. difficile* vs *C. difficile* mono-colonized mice at 24h (p=0.00193527).

### Carbon Source Transport and Metabolism

#### 5.1 Aminobutanoate degradation

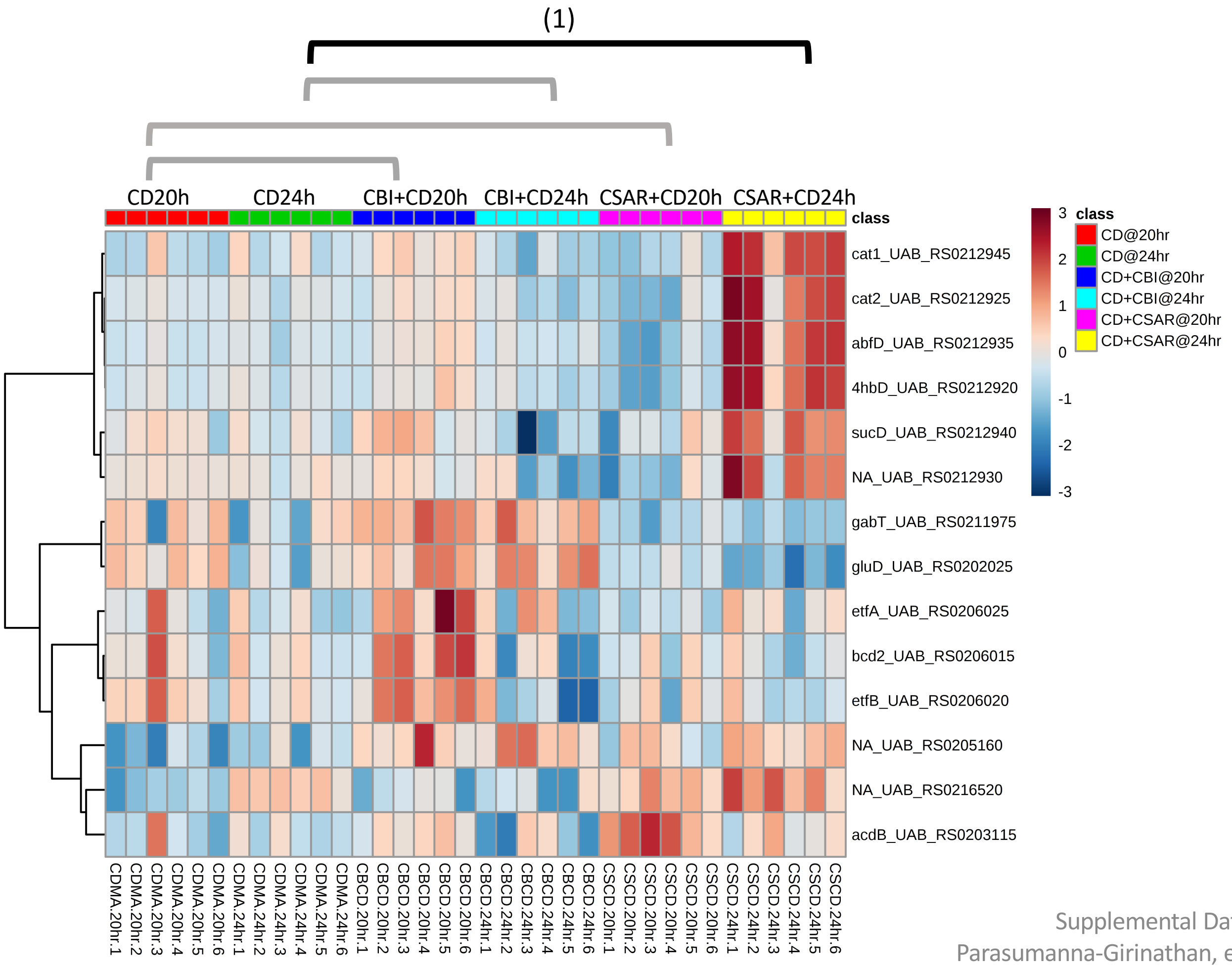


**Significantly Enriched Conditions:**

(1) Enriched in CSAR+*C. difficile* vs *C. difficile* mono-colonized mice at 24h (p=8.73E-03).

### Carbon Source Transport and Metabolism

#### 5.2 Beta-glucoside Transport and Conversion

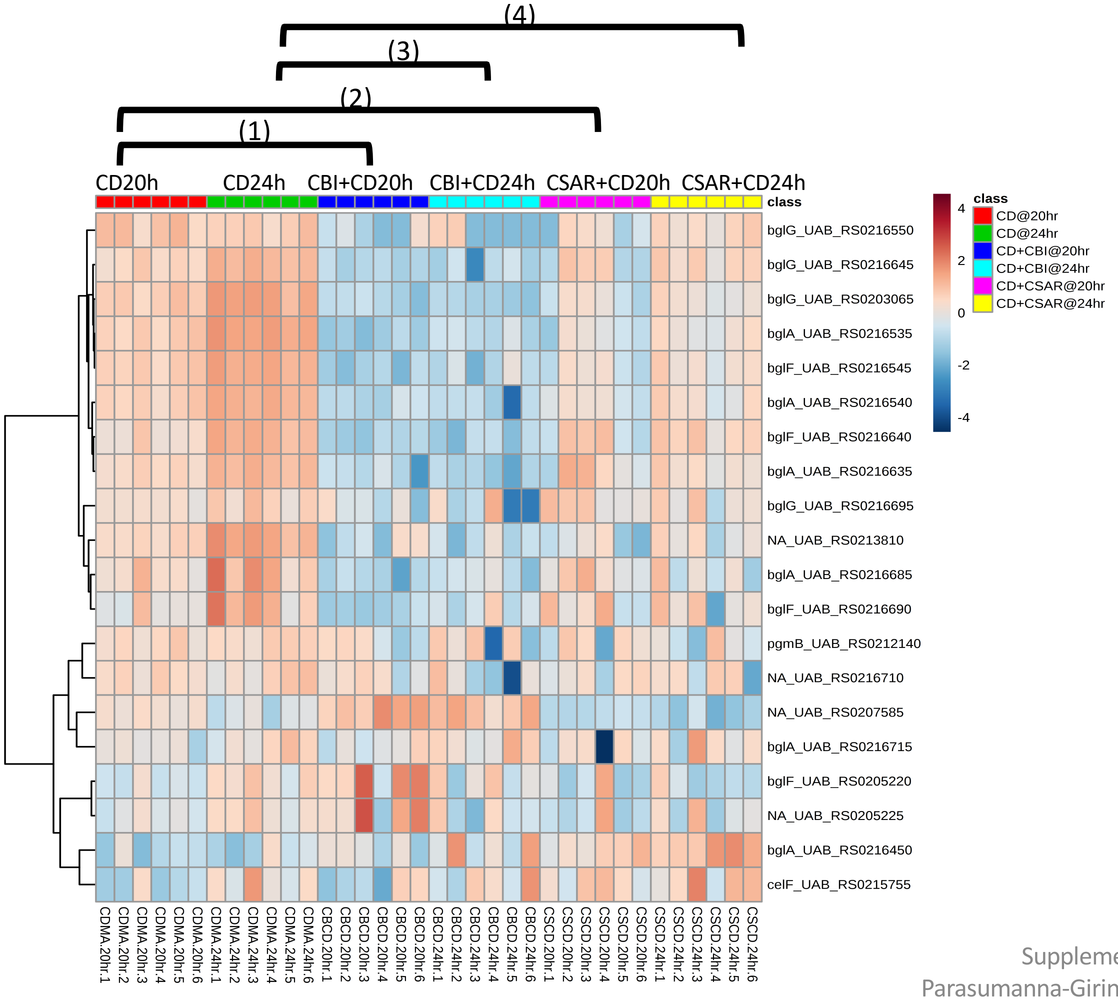


**Significantly Enriched Conditions:**

(1) Enriched in *C. difficile* mono-colonized vs CBI+*C. difficile* mice at 20h (p=8.86E-09).

(2) Enriched in *C. difficile* mono-colonized vs CSAR+*C. difficile* mice at 20h (p=3.01E-02).

(3) Enriched in *C. difficile* mono-colonized vs CBI+*C. difficile* mice at 24h (p=9.43E-04).

(4) Enriched in *C. difficile* mono-colonized vs CSAR+*C. difficile* mice at 24h (p=9.70E-03).

### Carbon Source Transport and Metabolism

#### 5.3 Butanoate Metabolism

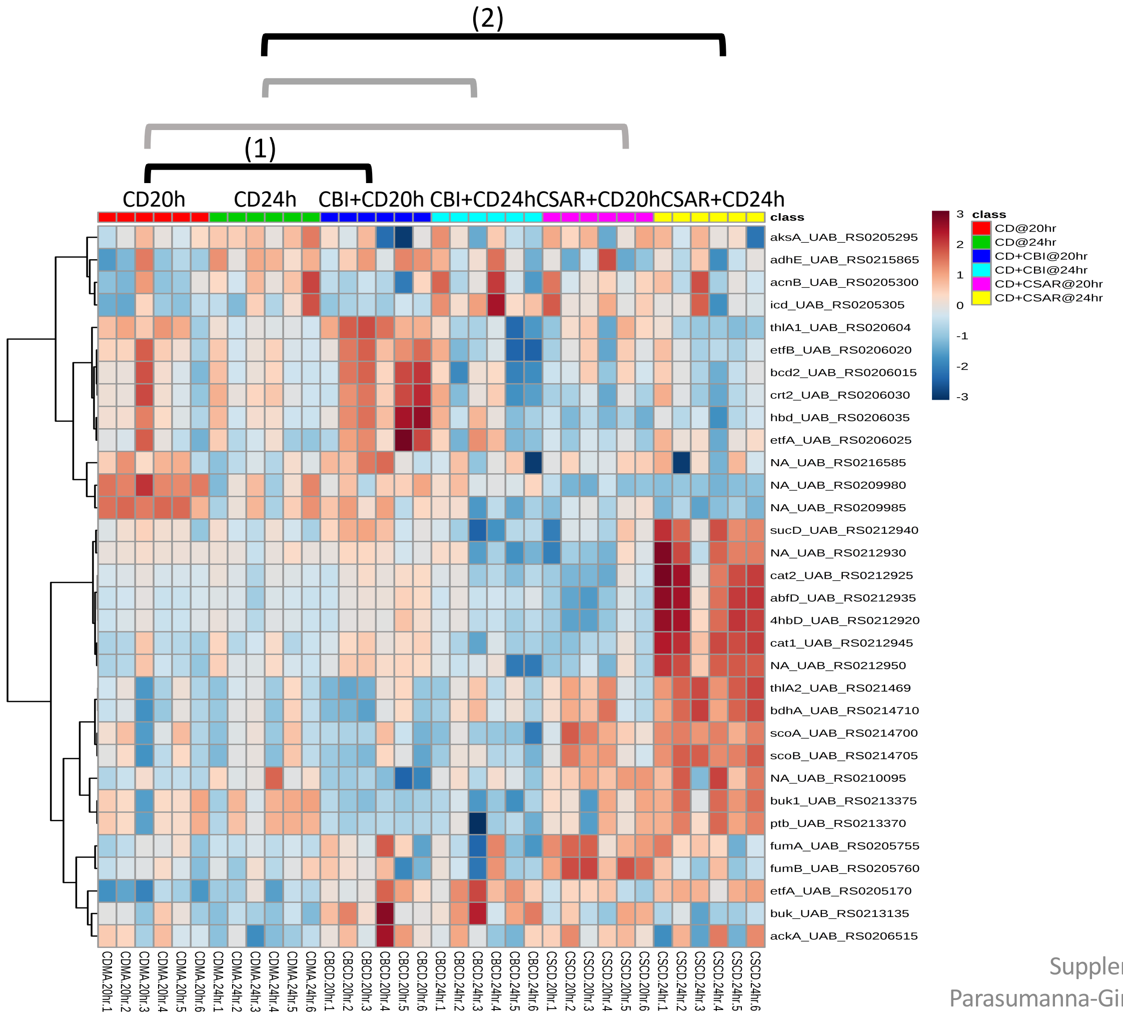


**<u>Significantly Enriched Conditions:</u>**

(1) Enriched in CBI+*C. difficile* vs *C. difficile* mono-colonized mice at 20h (p=6.61E-03).

(2) Enriched in CSAR+*C. difficile* vs *C. difficile* mono-colonized mice at 24h (p=8.84E-03).

### Carbon Source Transport and Metabolism

#### 5.4: Dipeptide and Oligopeptide Transport

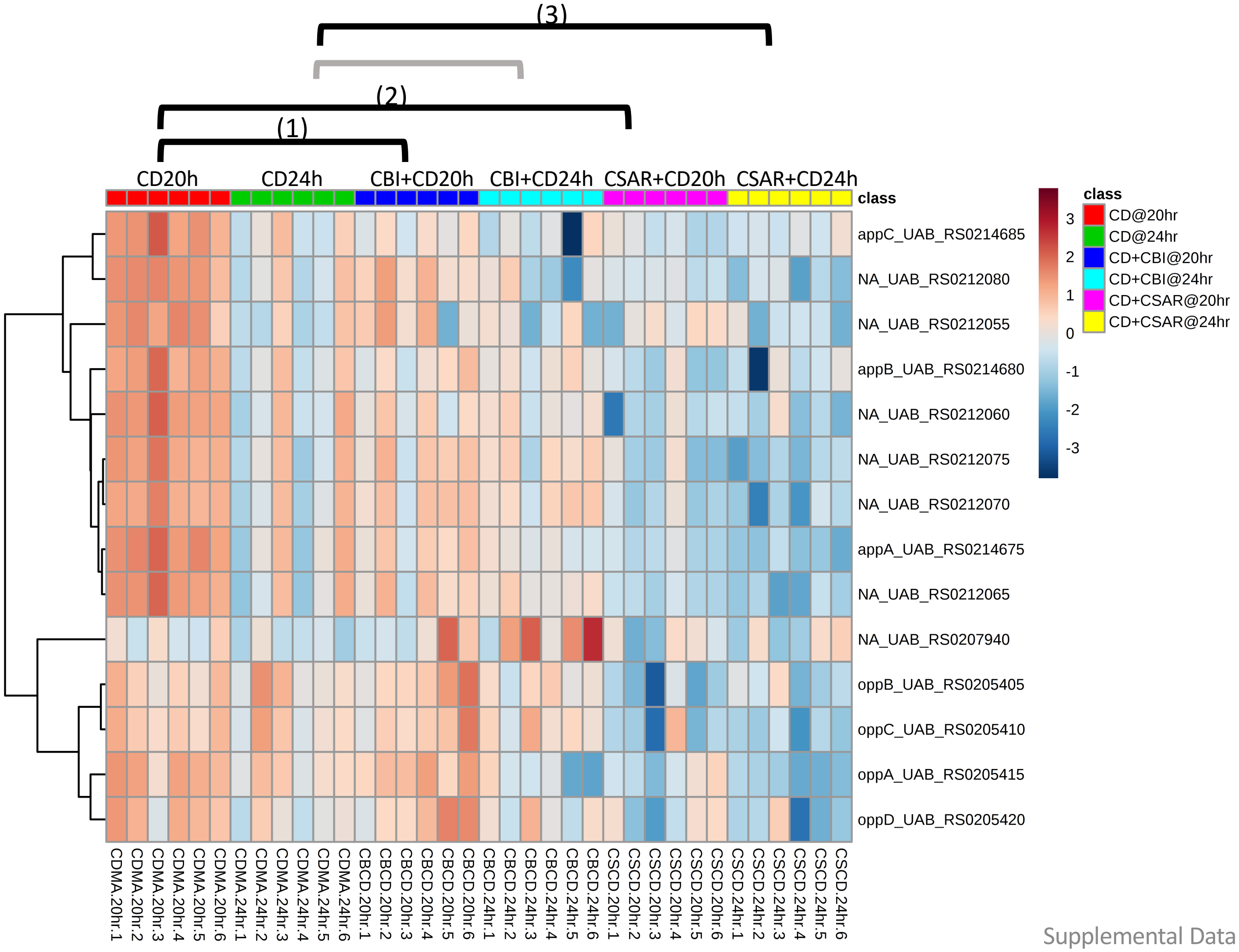


**Significantly Enriched Conditions:**

(1) Enriched in *C. difficile* mono-colonized vs CBI+*C. difficile* mice at 20h (p=3.03E-04).

(2) Enriched in *C. difficile* mono-colonized vs CSAR+*C. difficile* mice at 20h (p=8.34E-10).

(3) Enriched in *C. difficile* mono-colonized vs CSAR+*C. difficile* mice at 24h (p=4.44E-04).

### Carbon Source Transport and Metabolism

#### 5.5 Disaccharide Transport & Metabolism

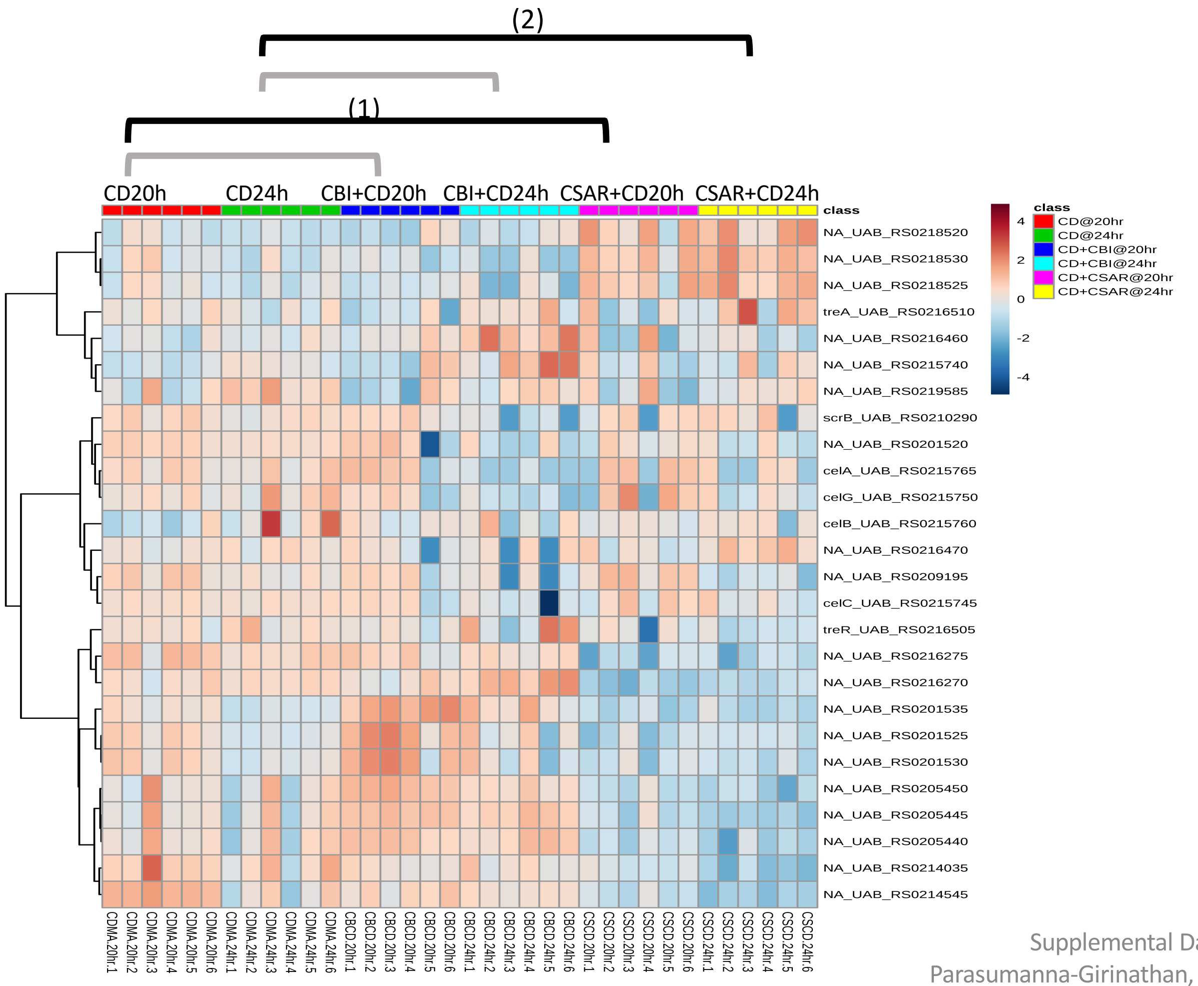


**Significantly Enriched Conditions:**

(1) Enriched in *C. difficile* mono-colonized vs CSAR+*C. difficile* mice at 20h (p=5.62E-03).

(2) Enriched in *C. difficile* mono-colonized vs CSAR+*C. difficile* mice at 24h (p=1.66E-02).

### Carbon Source Transport and Metabolism

#### 5.6 Ethanolamine Utilization

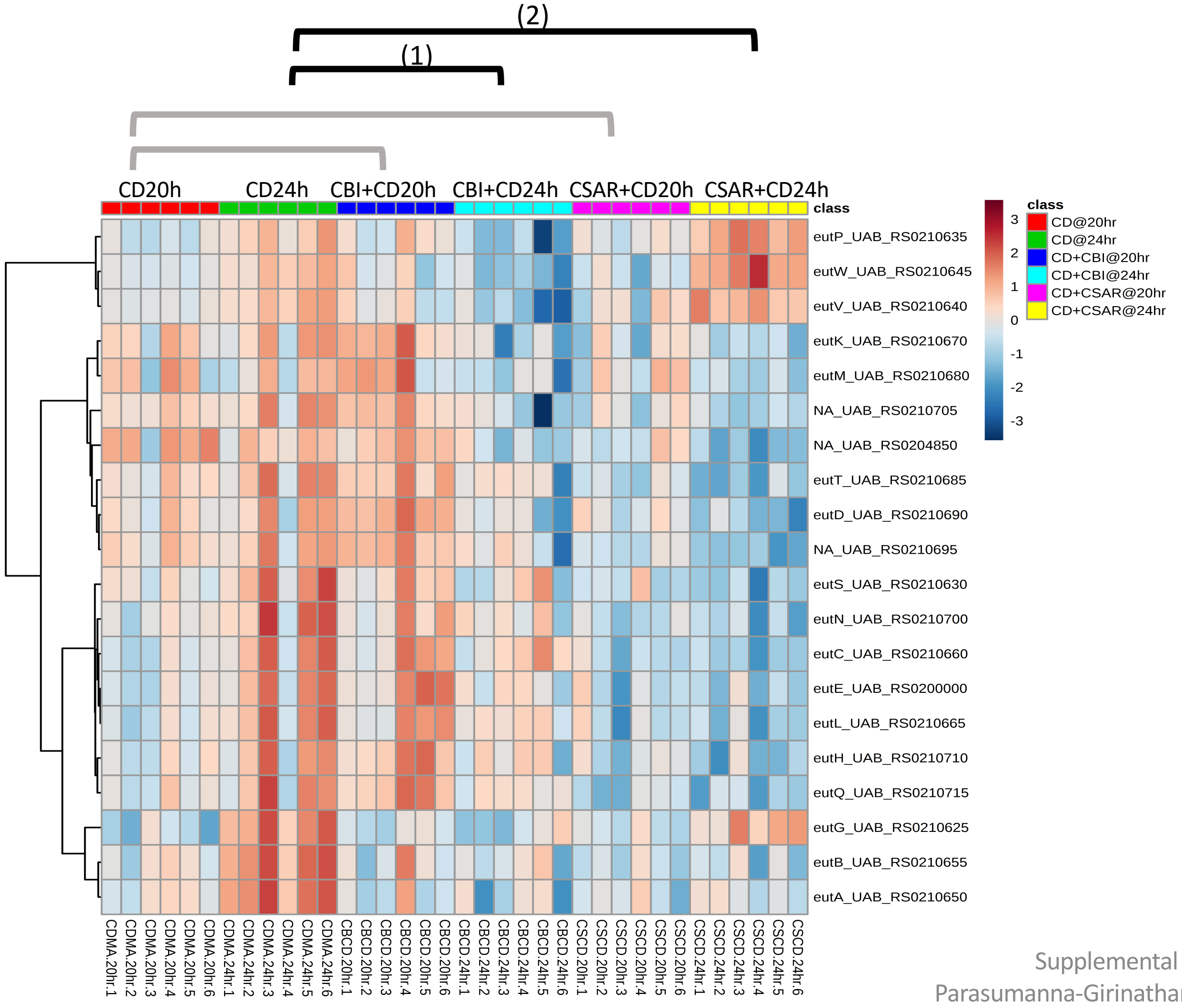


**Significantly Enriched Conditions:**

(1) Enriched in *C. difficil*e mono-colonized vs CBI+*C. difficile* mice at 24h (p=1.21E-08).

(2) Enriched in *C. difficil*e mono-colonized vs CSAR+*C. difficile* mice at 24h (p=1.04E-09).

### Carbon Source Transport and Metabolism

#### 5.7 Fructose Transport & Conversion

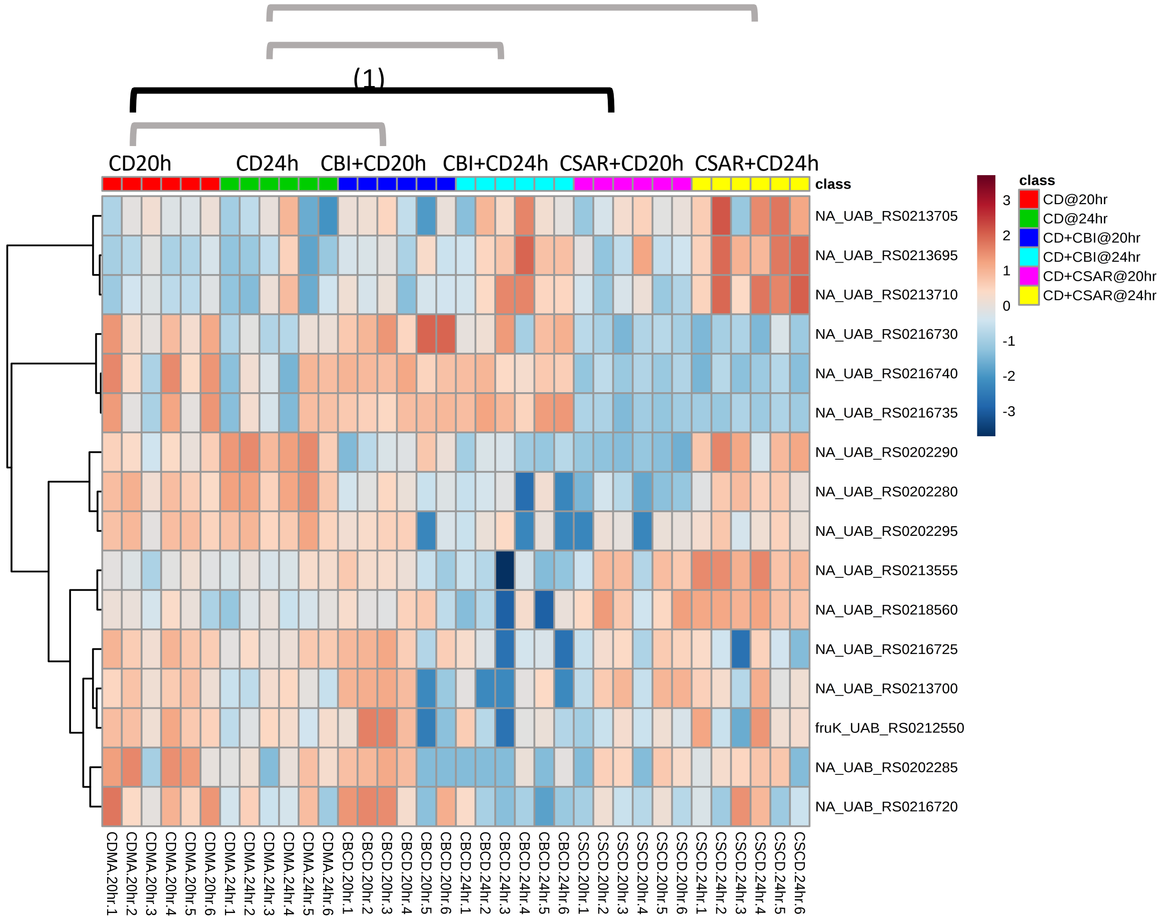


**Significantly Enriched Conditions:**

(1) Enriched in *C. difficile* mono-colonized vs CSAR+*C. difficile* mice at 20h (p=1.90E-03).

### Carbon Source Transport and Metabolism

#### 5.8 Glucose and Maltose Transport & Conversion

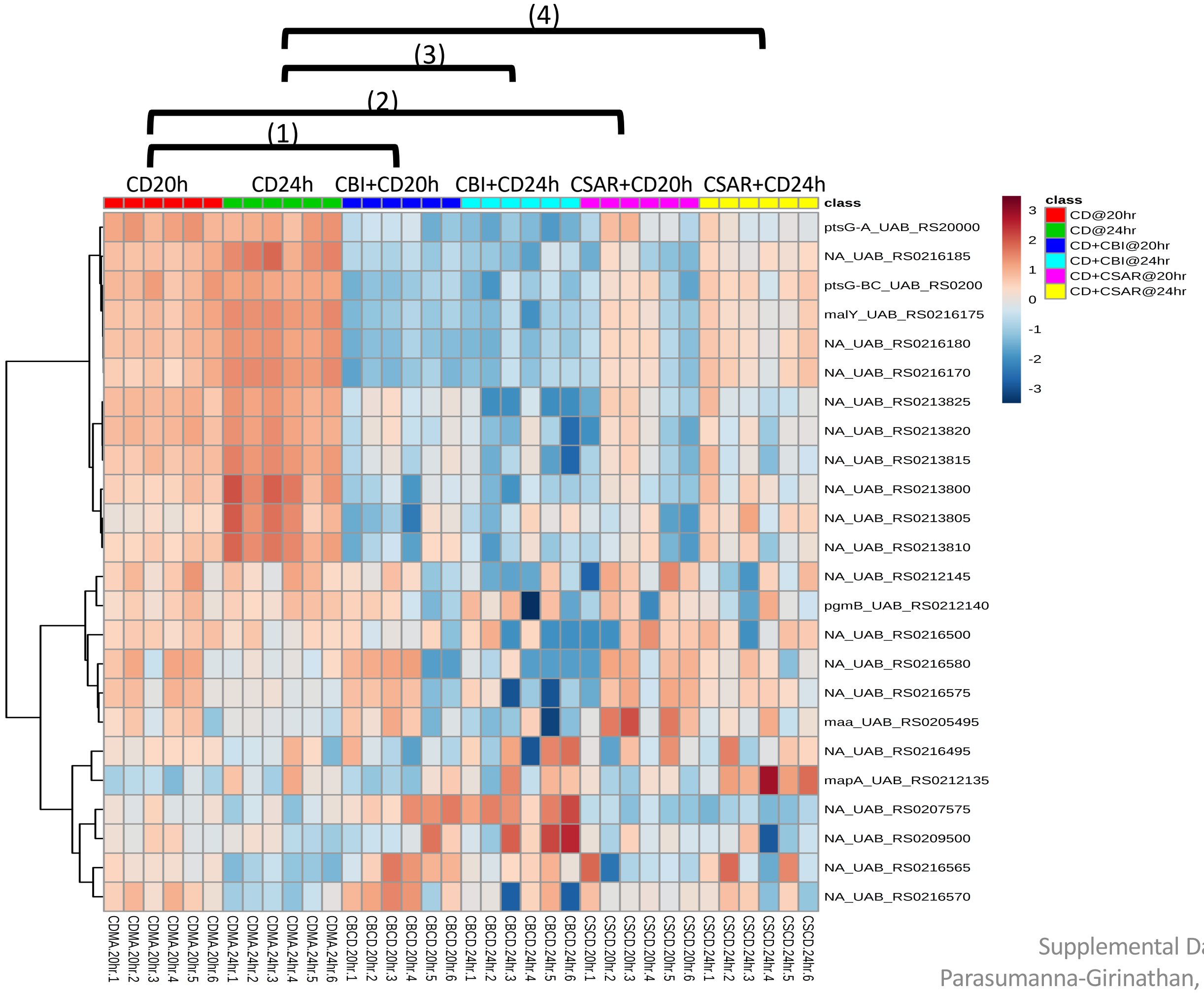


**Significantly Enriched Conditions:**

(1) Enriched in *C. difficil*e mono-colonized vs CBI+*C. difficile* mice at 20h (p=4.31E-08).

(2) Enriched in *C. difficil*e mono-colonized vs CSAR+*C. difficile* mice at 20h (p=1.35E-04).

(3) Enriched in *C. difficil*e mono-colonized vs CBI+*C. difficile* mice at 24h (p=3.06E-05).

(4) Enriched in *C. difficil*e mono-colonized vs CSAR+*C. difficile* mice at 24h (p=1.65E-04).

### Carbon Source Transport and Metabolism

#### 5.9 Mannose, Sorbose, Fructose Transport & Conversion

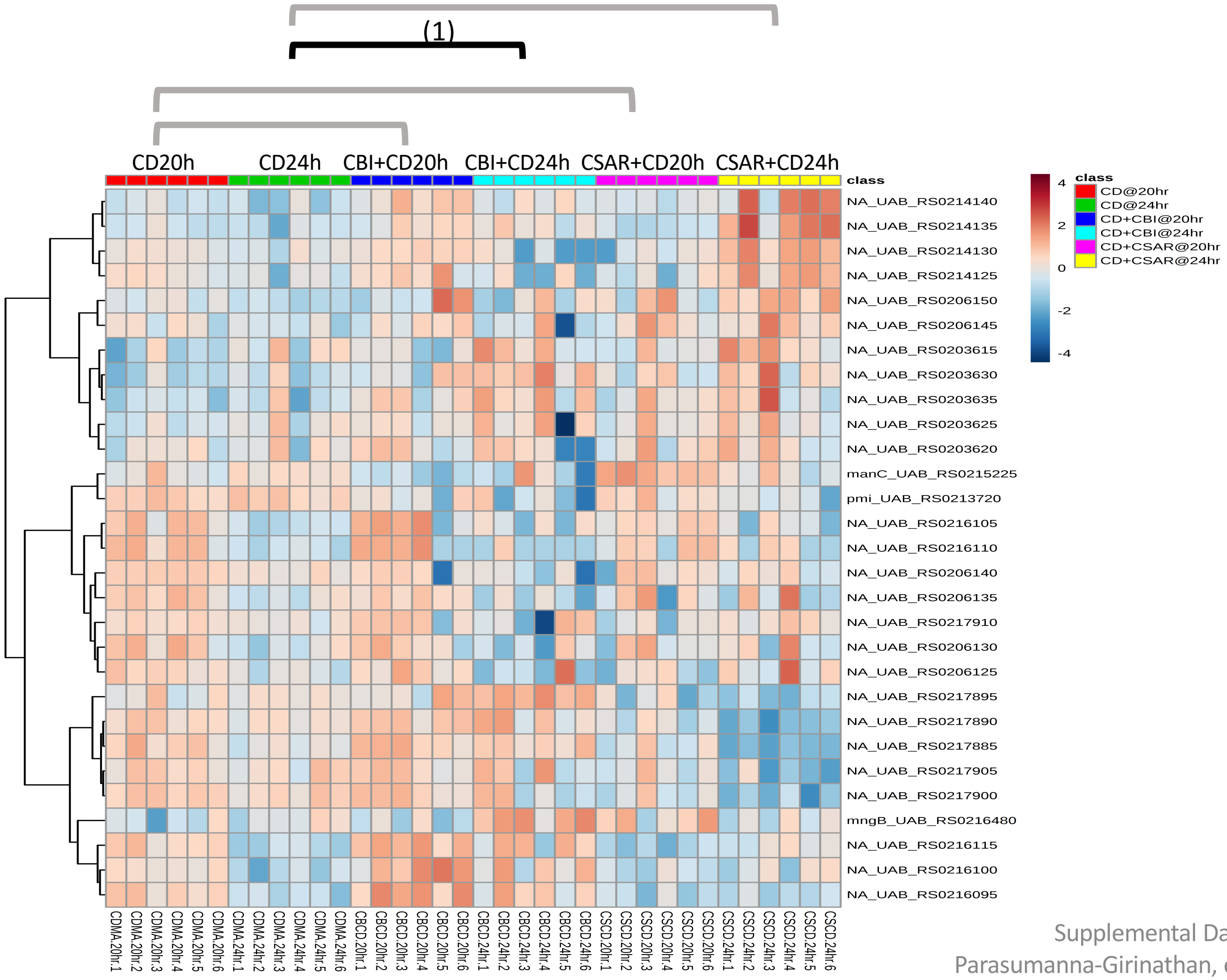


**Significantly Enriched Conditions:**

(1) Enriched in CBI+*C. difficile* vs *C. difficile* mono-colonized mice at 24h (p=4.39E-02).

### Carbon Source Transport and Metabolism

#### 5.10 Ornithine Fermentation

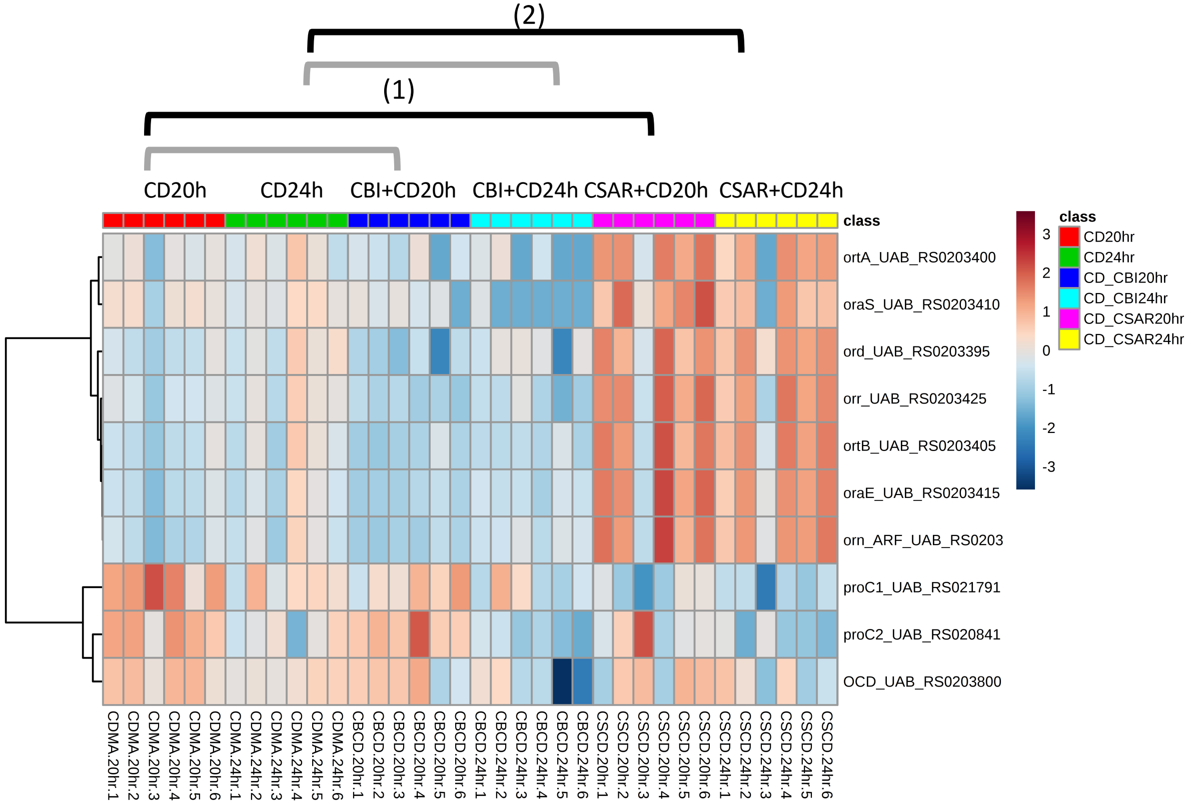


**Significantly Enriched Conditions:**

(1) Enriched in CSAR+*C. difficile* vs *C. difficile* mono-colonized mice at 20h (p=4.24E-03).

(2) Enriched CSAR+*C. difficile* vs *C. difficile* mono-colonized mice at 24h (p=1.94E-03).

### Carbon Source Transport and Metabolism

#### 5.11 Polysaccharide Degradation

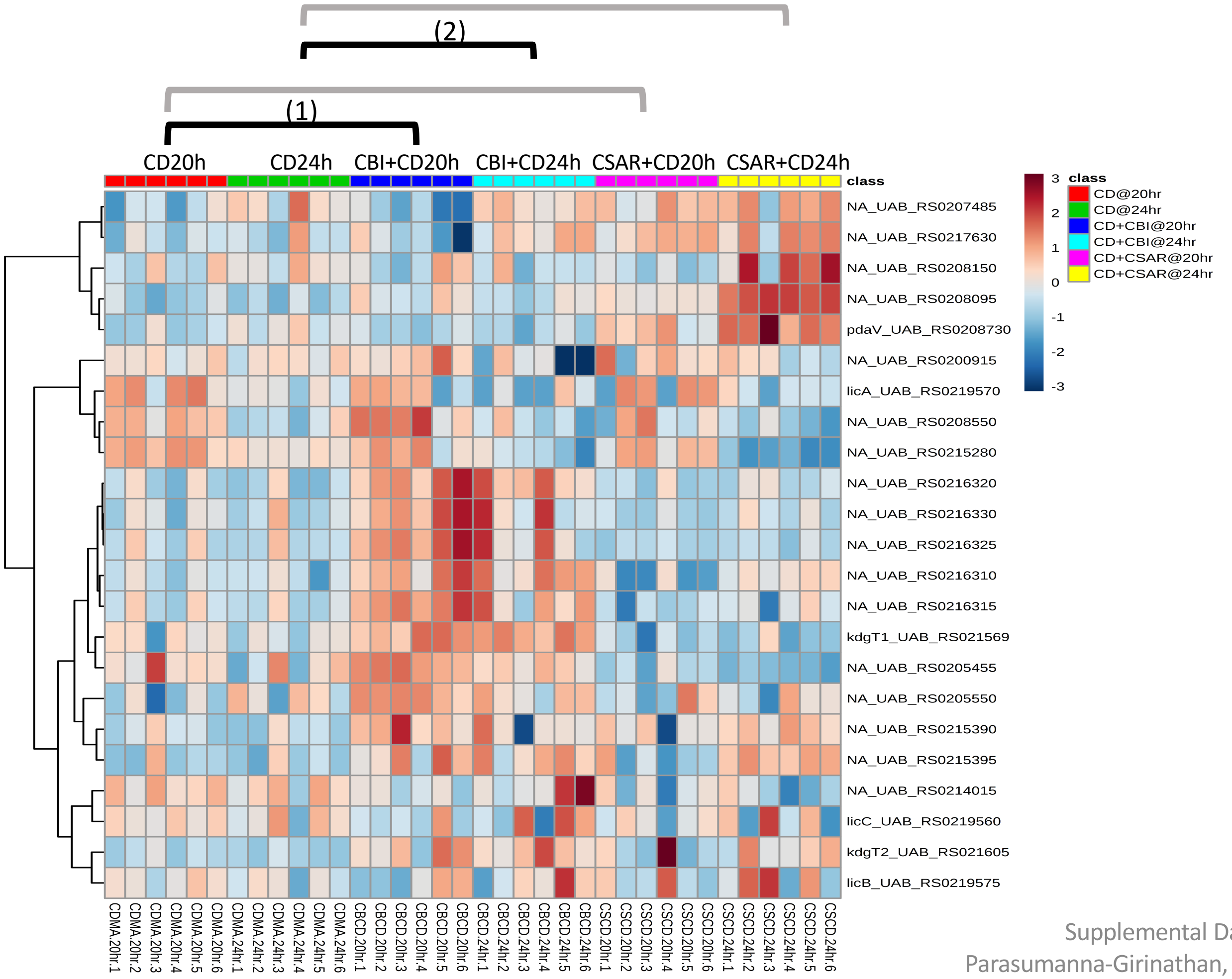


**Significantly Enriched Conditions:**

(1) Enriched in CBI+*C. difficile* vs *C. difficile* mono-colonized mice at 20h (p=5.18E-07).

(2) Enriched in CBI+*C. difficile* vs *C. difficile* mono-colonized mice at 24h (p=2.36E-02).

### Carbon Source Transport and Metabolism

#### 5.12 Pyruvate Metabolism

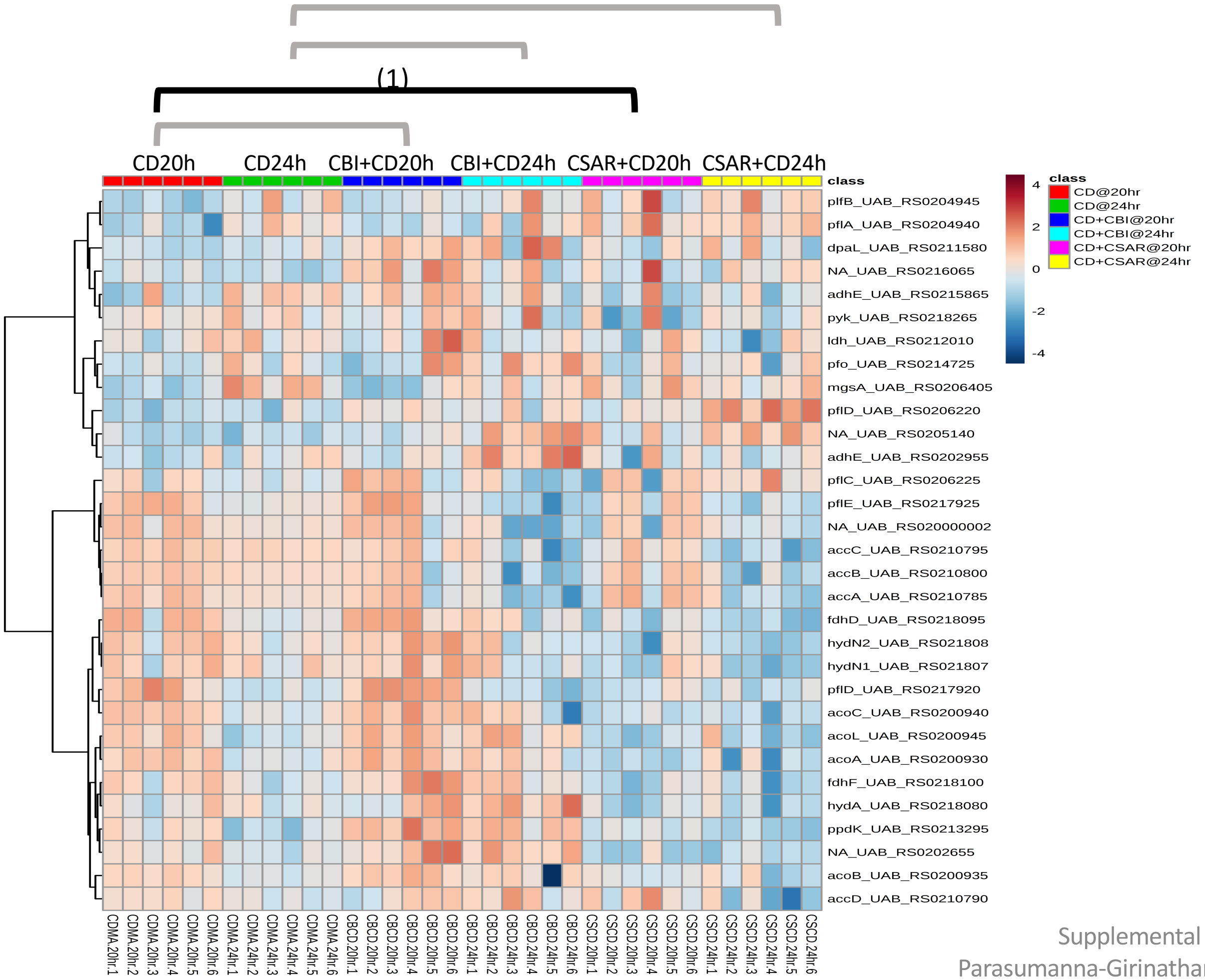


**Significantly Enriched Conditions:**

(1) Enriched in *C. difficile* mono-colonized vs CSAR+*C. difficile* mice at 20h (p=3.43E-02).

### Carbon Source Transport and Metabolism

#### 5.13 Ribose Transport & Metabolism

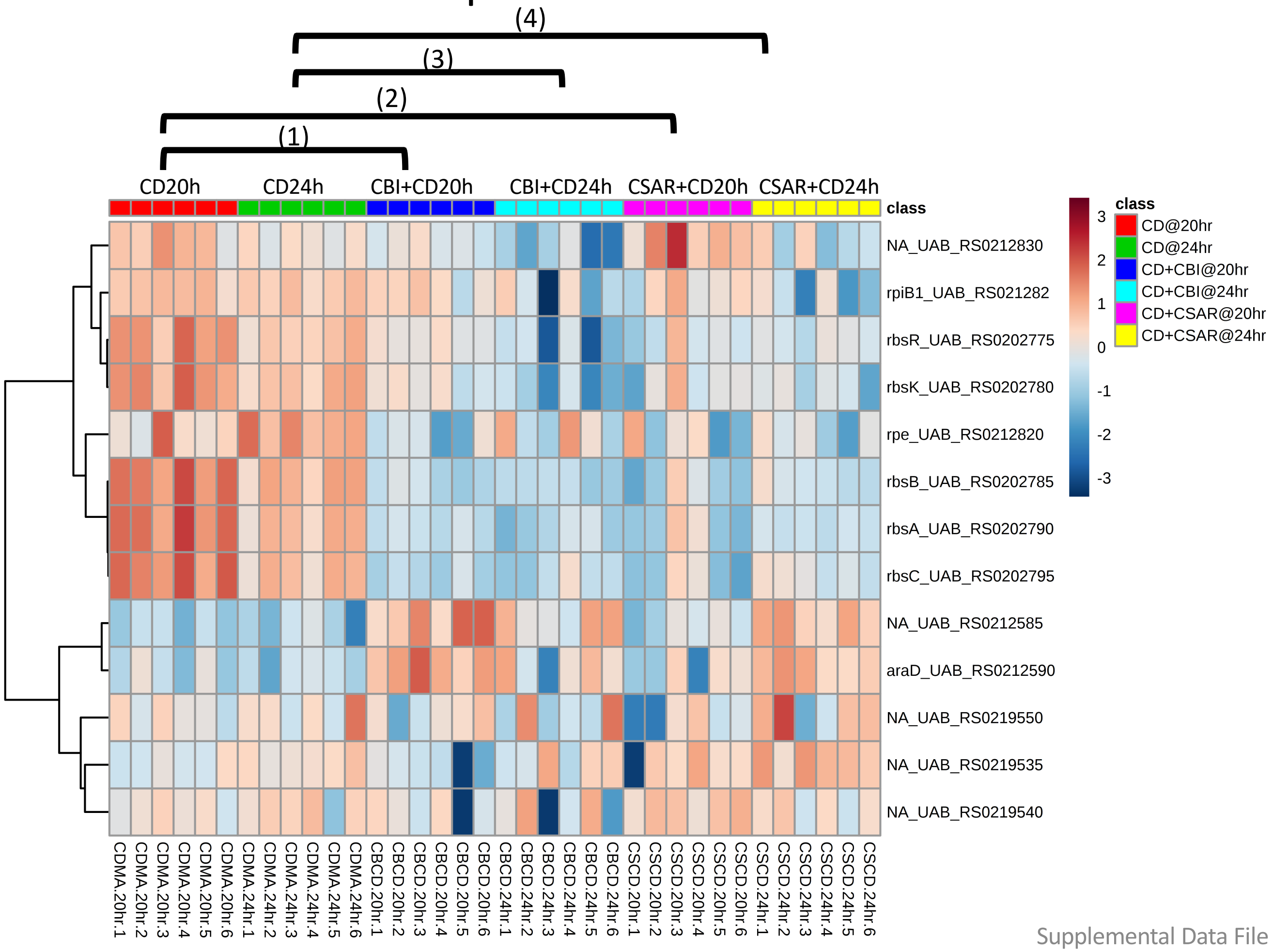


**Significantly Enriched Conditions:**

(1) Enriched in *C. difficile* mono-colonized vs CBI+*C. difficile* mice at 20h (p=2.04E-04).

(2) Enriched in *C. difficile* mono-colonized vs CSAR+*C. difficile* mice at 20h (p=3.58E-02).

(3) Enriched in *C. difficile* mono-colonized vs CBI+*C. difficile* mice at 24h (p=2.77E-03).

(4) Enriched in *C. difficile* mono-colonized vs CSAR+*C. difficile* mice at 24h (p=7.20E-03).

### Carbon Source Transport and Metabolism

#### 5.14 Stickland Fermentation

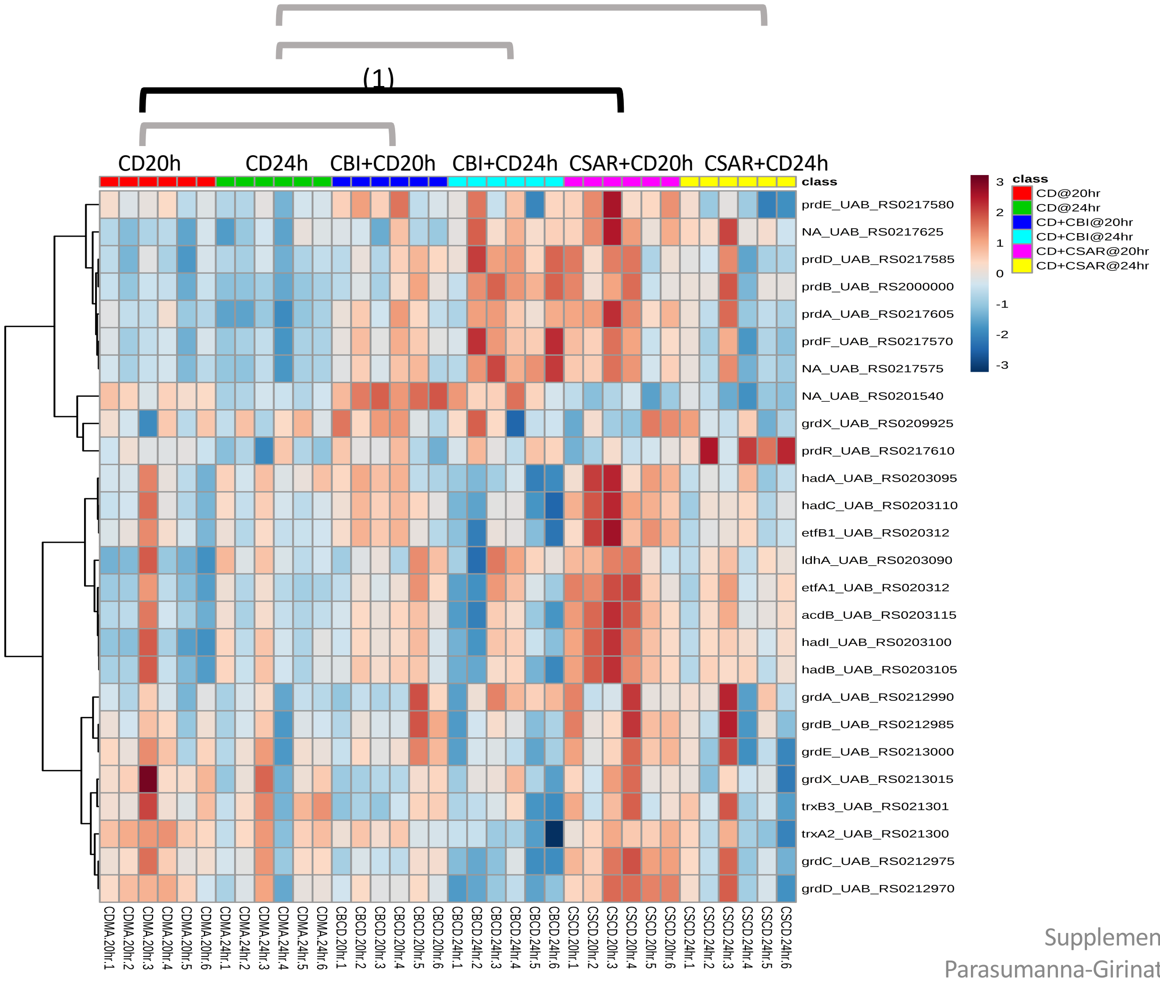


**Significantly Enriched Conditions:**

(1) Enriched in CSAR+*C. difficile vs C. difficile* mono-colonized mice at 20h (p=4.855E-03).

### Carbon Source Transport and Metabolism

#### 5.15 Sugar Alcohol Transport & Metabolism

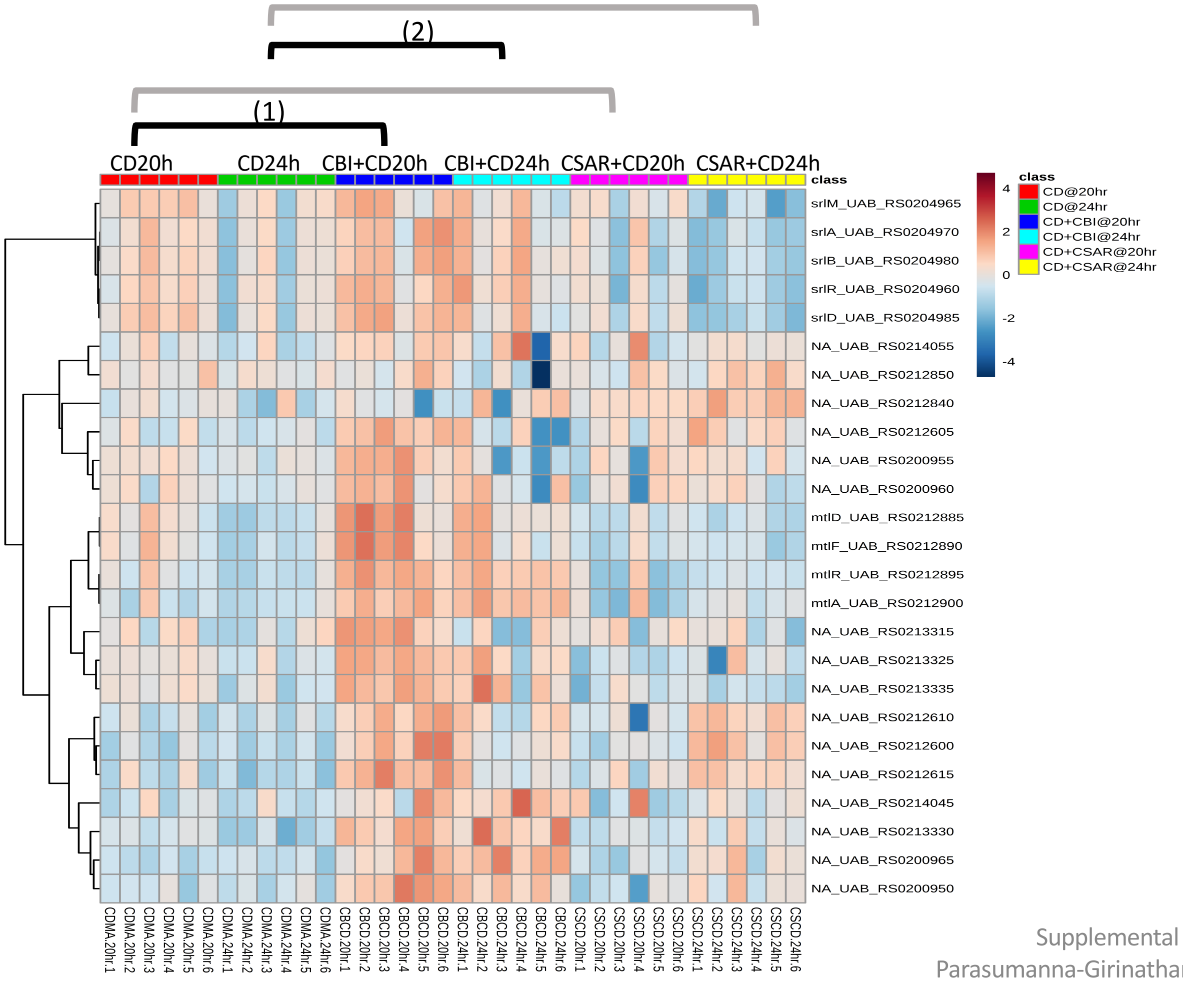


**Significantly Enriched Conditions:**

(1) Enriched in CBI+*C. difficile vs C. difficile* mono-colonized mice at 20h (p=1.09E-13).

(2) Enriched in CBI+*C. difficile vs C. difficile* mono-colonized mice at 24h (p=5.27E-08).

### Carbon Source Transport and Metabolism

#### 5.16 Transport Binding Proteins: Amino acids & amines

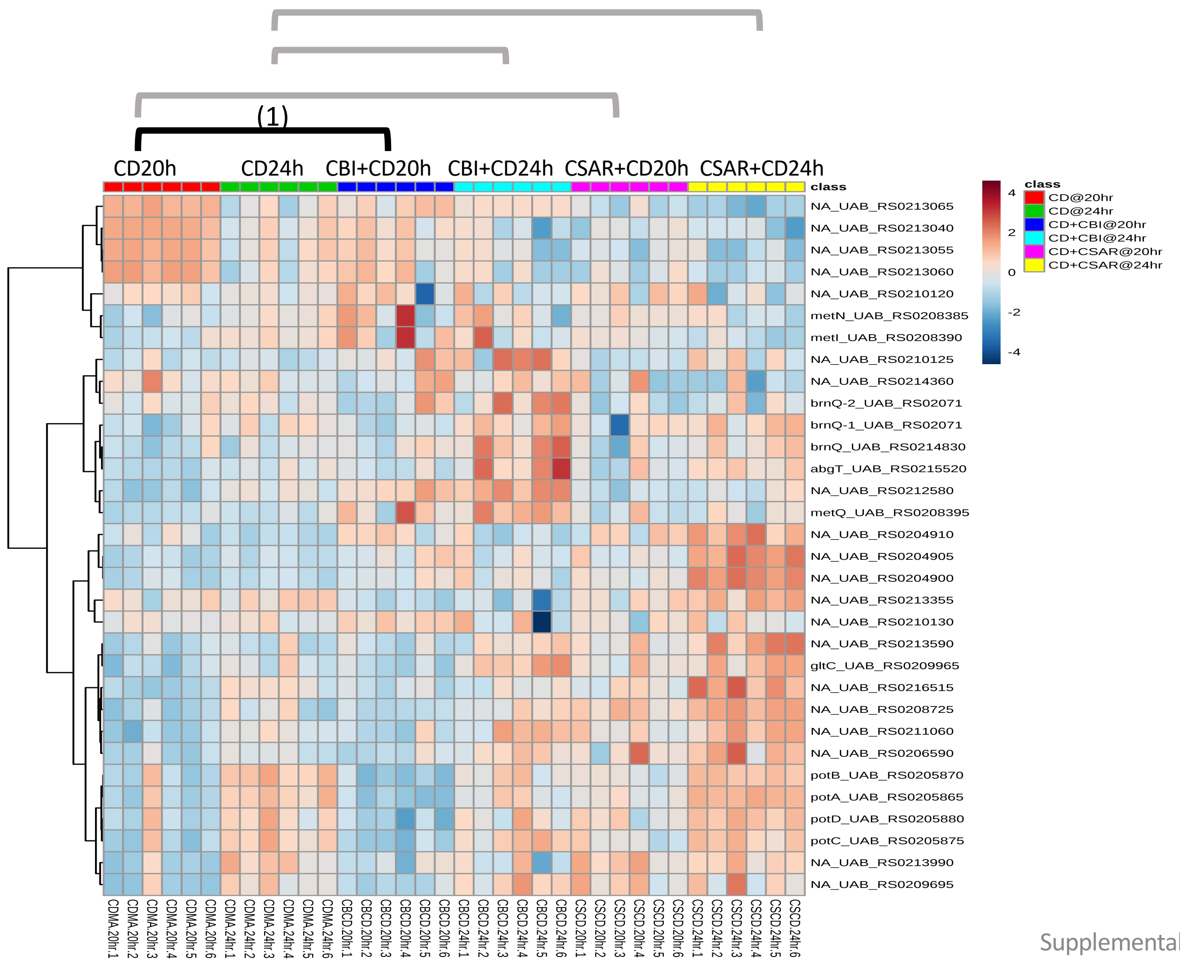


**Significantly Enriched Conditions:**

(1) Enriched in CBI+*C. difficile vs C. difficile* mono-colonized mice at 20h (p=7.27E-03).

### Carbon Source Transport and Metabolism

#### 5.17 Wood-Ljungdahl Pathway

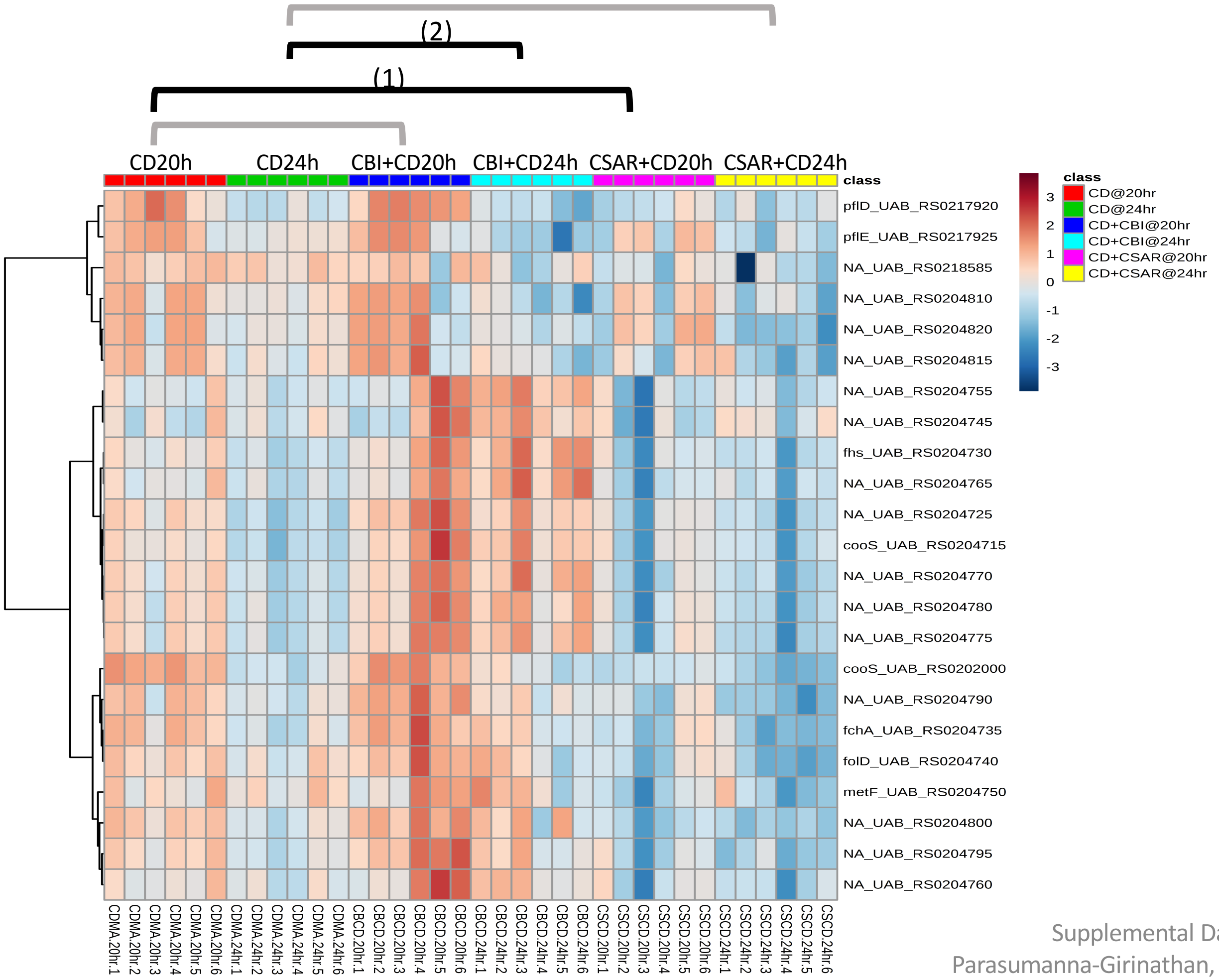


**Significantly Enriched Conditions:**

(1) Enriched in *C. difficile* mono-colonized vs CSAR+*C. difficile* mice at 20h (p=1.79E-03).

(2) Enriched in CBI+*C. difficile vs C. difficile* mono-colonized mice at 24h (p=4.43E-02).

### Carbon Source Transport and Metabolism

#### 5.18 Xanthine, Hypoxanthine: Transport & Metabolism

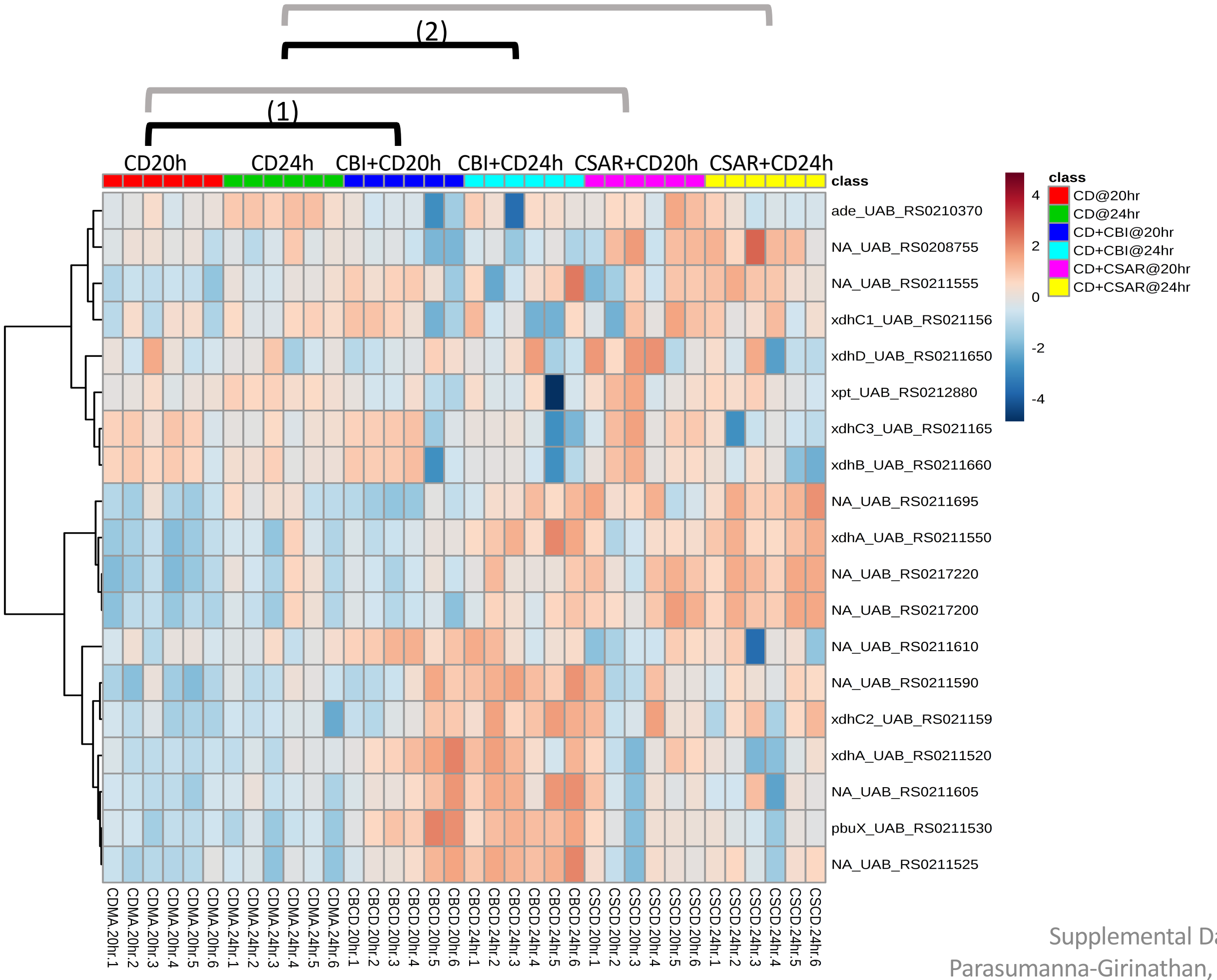


**Significantly Enriched Conditions:**

(1) Enriched in CBI+*C. difficile vs C. difficile* mono-colonized mice at 20h (p=7.41E-05).

(2) Enriched in CBI+*C. difficile vs C. difficile* mono-colonized mice at 24h (p=3.93E-02).

### Core Cellular Machinery

#### 5.19 ATP Synthesis

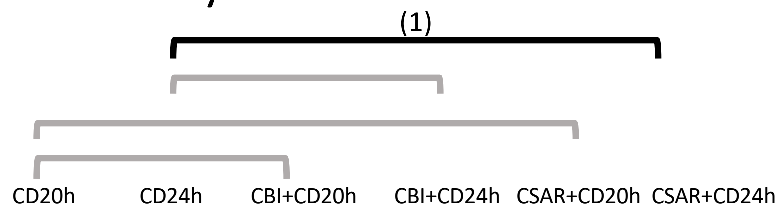


**<u>Significantly Enriched Conditions:</u>**

(1) Enriched in *C. difficil*e mono-colonized vs CBI+*C. difficile* mice at 24h (p=4.78E-02).

### Core Cellular Machinery

#### 5.20 DNA Replication

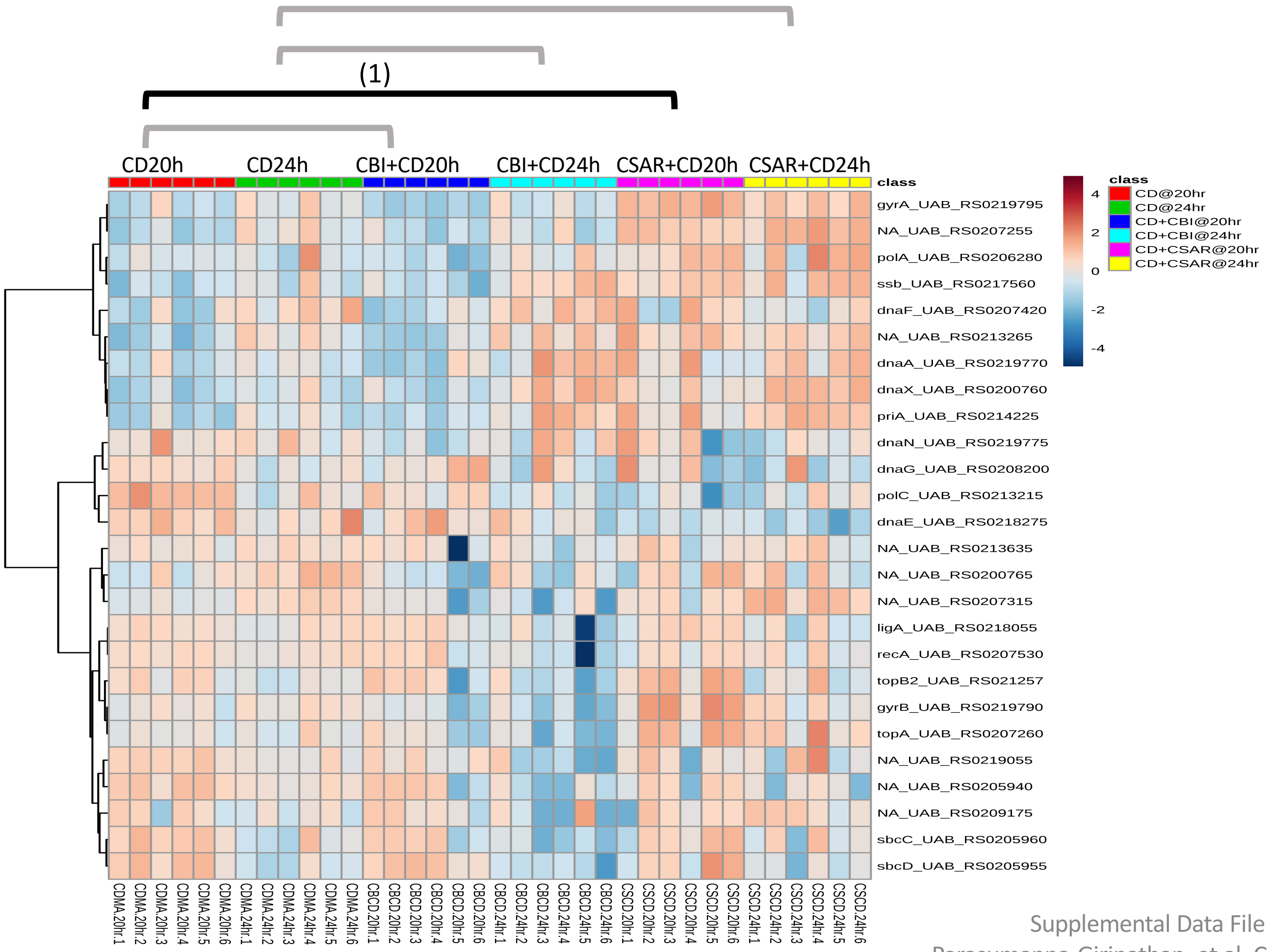


**Significantly Enriched Conditions:**

(1) Enriched in CSAR+*C. difficile vs C. difficile* mono-colonized mice at 20h (p=1.21E-02).

### Core Cellular Machinery

#### 5.21 Protein translation & Modification

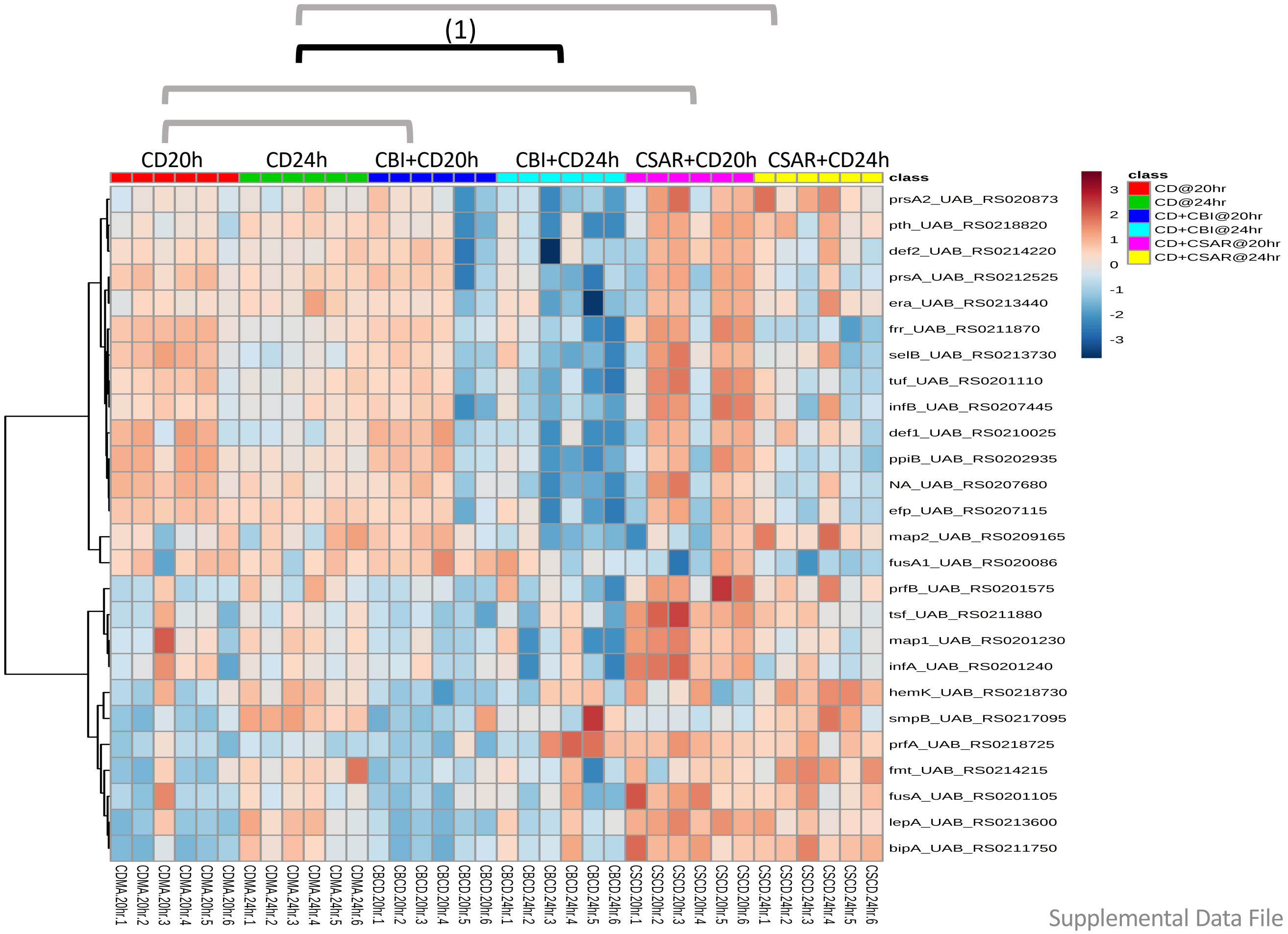


**Significantly Enriched Conditions:**

(1) Enriched in *C. difficile* mono-colonized vs CBI+*C. difficile* mice at 24h (p=1.66E-02).

### Core Cellular Machinery

#### 5.22 Ribosomal Structural Components and Modification

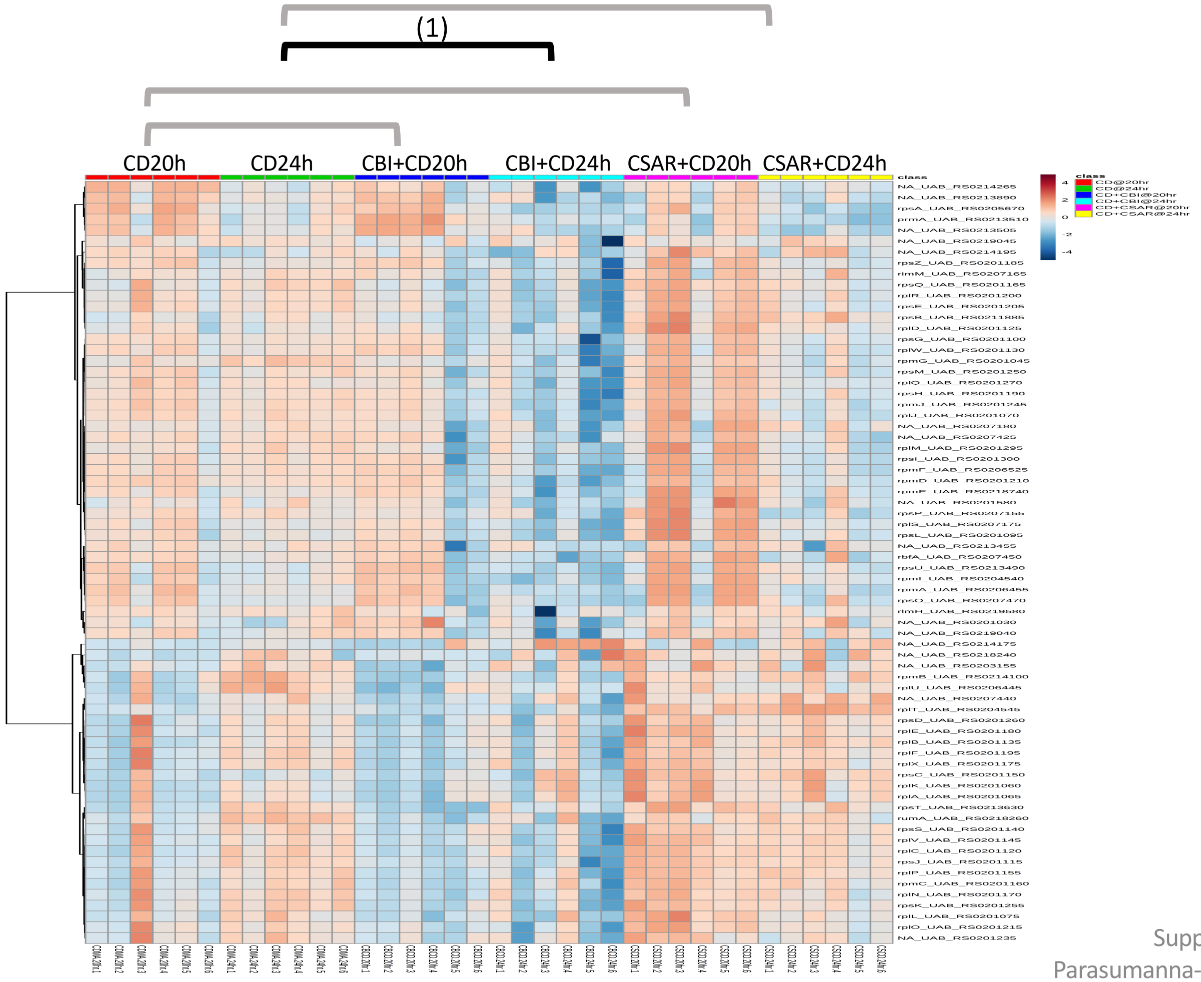


**<u>Significantly Enriched Conditions:</u>**

(1) Enriched in *C. difficil*e mono-colonized vs CBI+*C. difficile* mice at 24h (p=2.59E-11).

### Core Cellular Machinery

#### 5.23 RNA Transcription & Modification

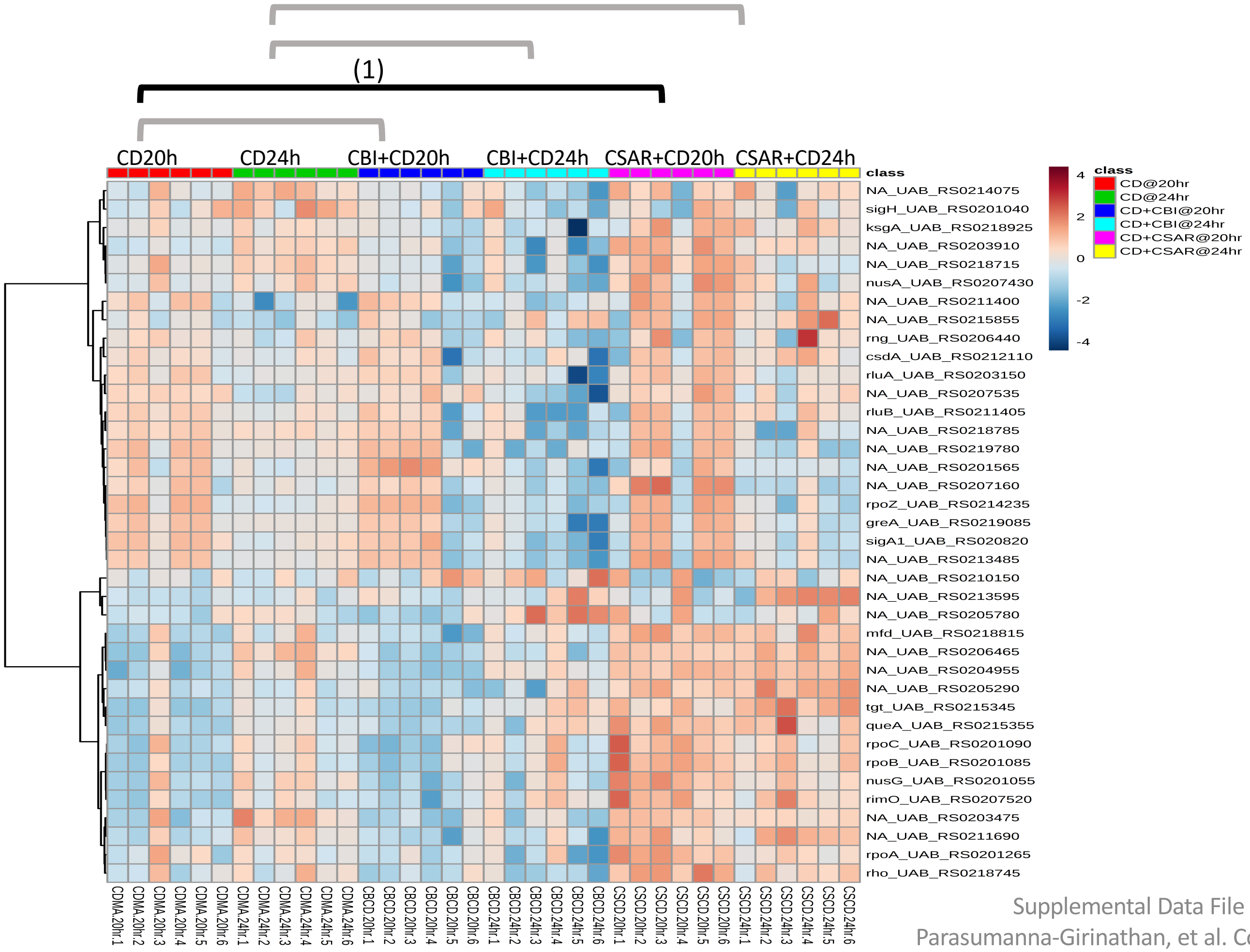


**Significantly Enriched Conditions:**

(1) Enriched in CSAR+*C. difficile vs C. difficile* mono-colonized mice at 20h (p=4.82E-03).

### Cell wall Components

#### 5.24 Cell Wall Turnover

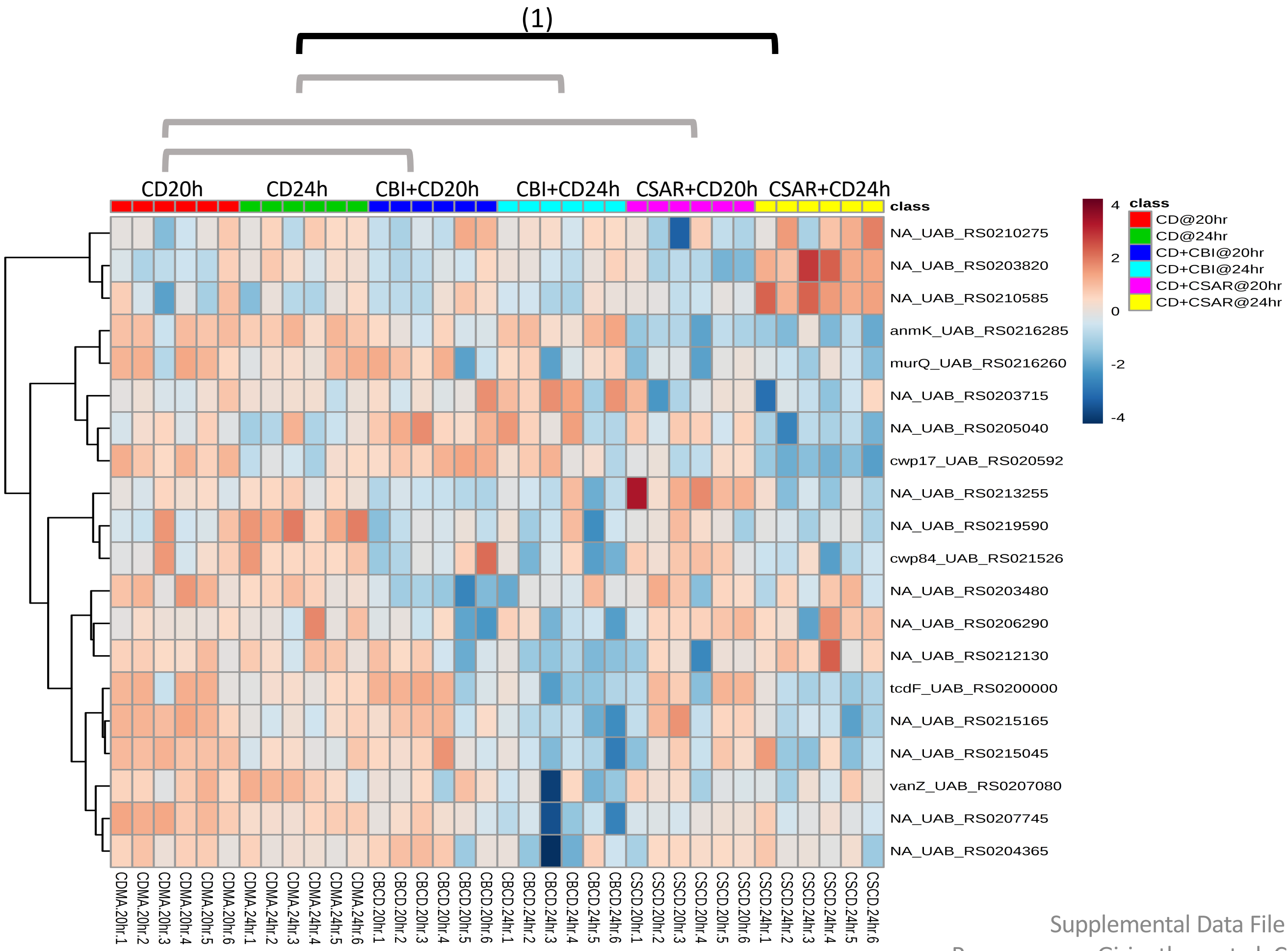


**Significantly Enriched Conditions:**

(1) Enriched in *C. difficile* mono-colonized vs CSAR+*C. difficile* mice at 24h (p=1.89E-02).

### Cell wall Components

#### 5.25 Gram+ Surface Anchored Components

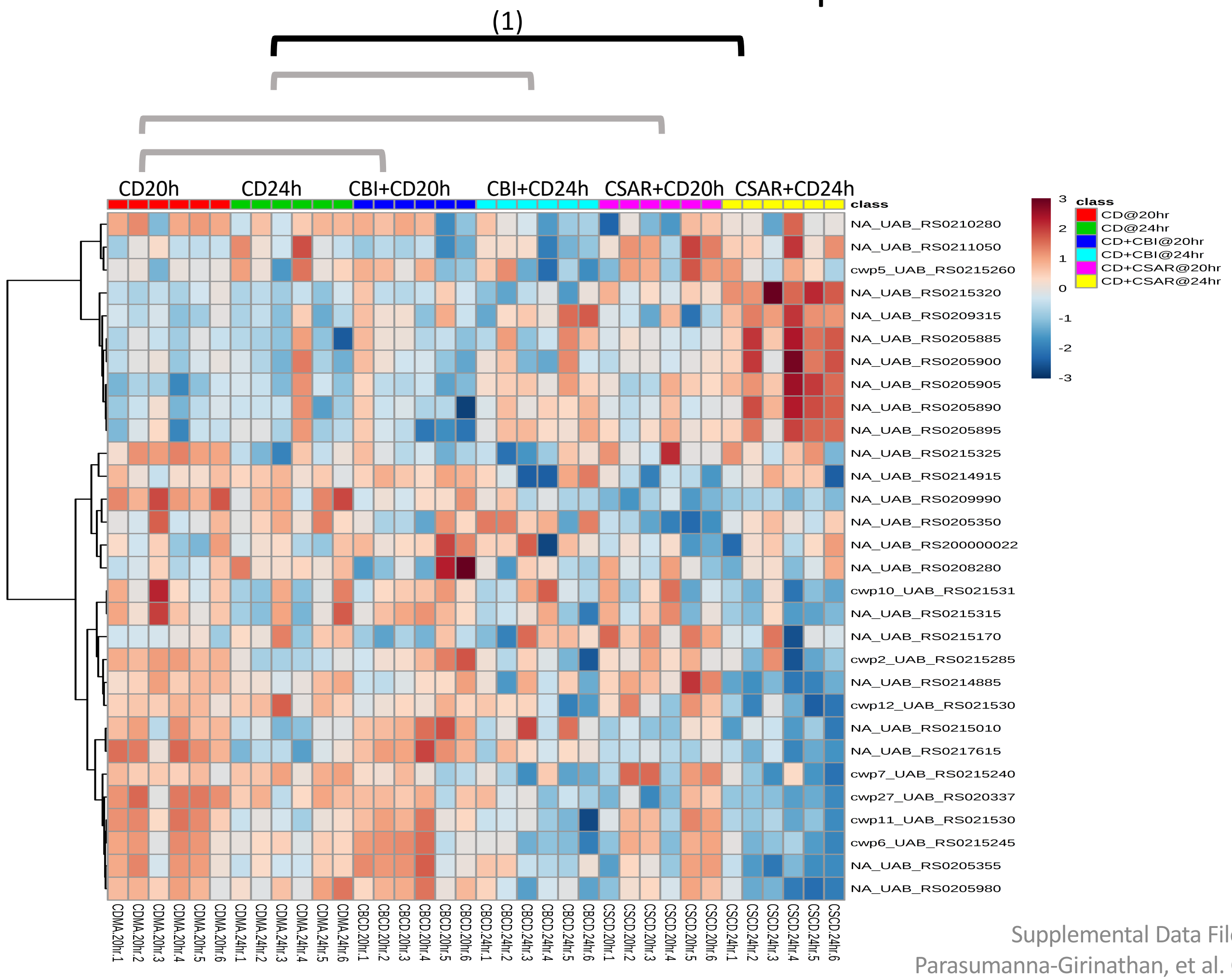


**Significantly Enriched Conditions:**

(1) Enriched in *C. difficile* mono-colonized vs CSAR+*C. difficile* mice at 24h (p=2.63E-02).

### Vitamin and Micronutrient Metabolism

#### 5.26 Cation Transport Binding Proteins

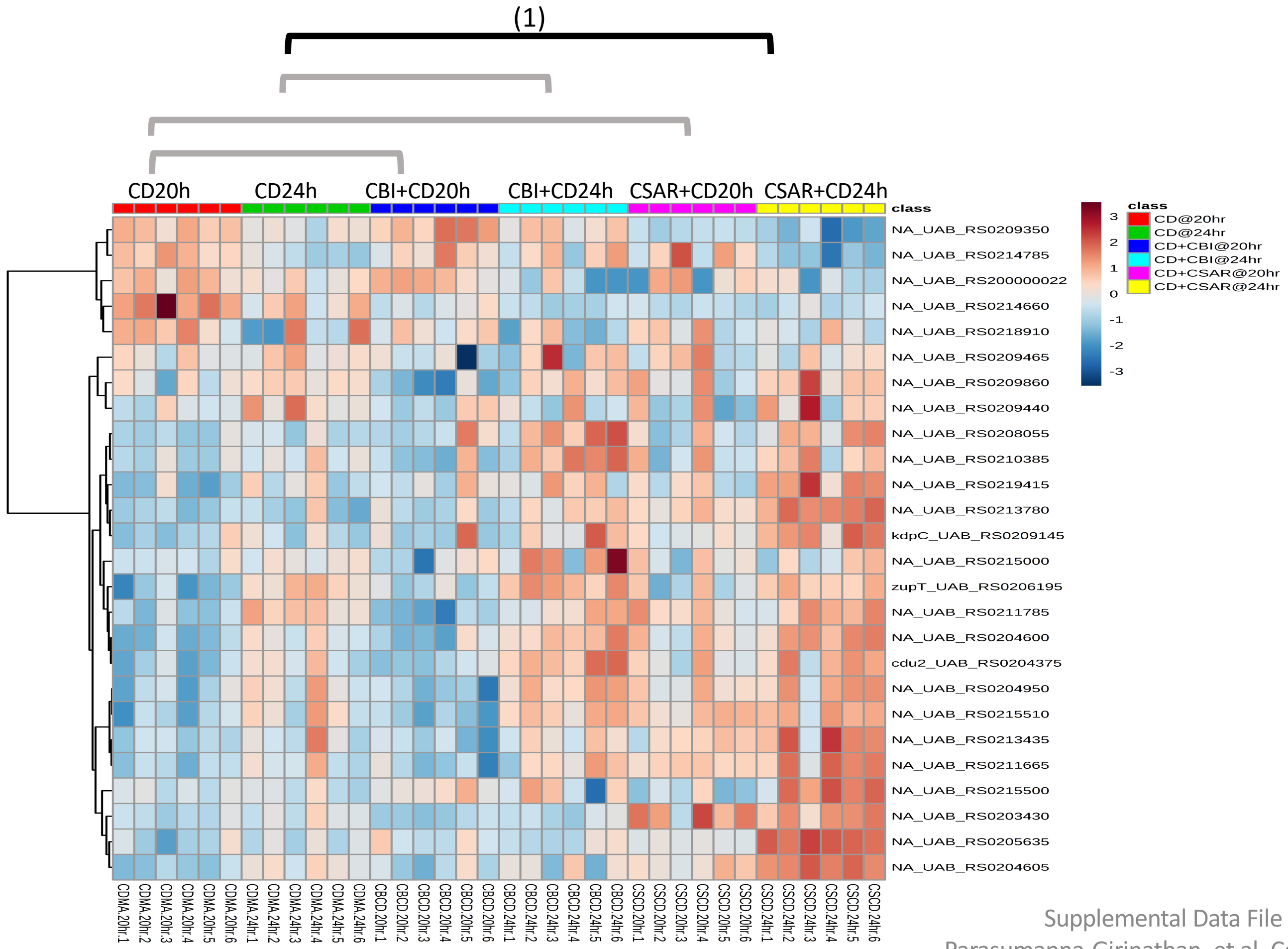


**Significantly Enriched Conditions:**

(1) Enriched in CSAR+*C. difficile vs C. difficile* mono-colonized mice at 24h (p=2.02E-03).

### Vitamin and Micronutrient Metabolism

#### 5.27 Cobalamin Biosynthesis

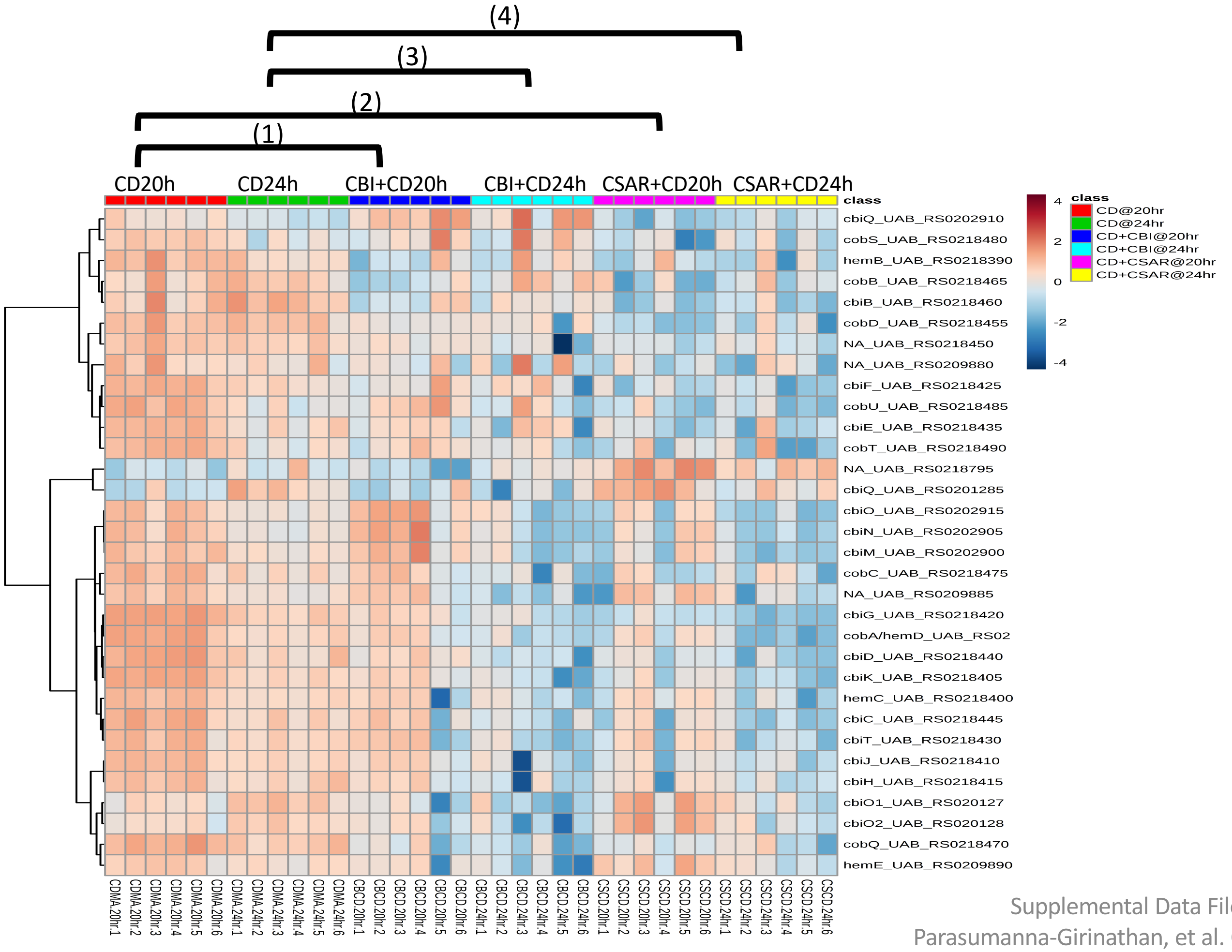


**Significantly Enriched Conditions:**

(1) Enriched in *C. difficile* mono-colonized vs CBI+*C. difficile* mice at 20h (p=7.48E-05).

(2) Enriched in *C. difficile* mono-colonized vs CSAR+*C. difficile* mice at 20h (p=2.54E-11).

(3) Enriched in *C. difficile* mono-colonized vs CBI+*C. difficile* mice at 24h (p=2.12E-06).

(4) Enriched in *C. difficile* mono-colonized vs CSAR+*C. difficile* mice at 24h (p=1.72E-05).

### Vitamin and Micronutrient Metabolism

#### 5.28 Folate Biosynthesis

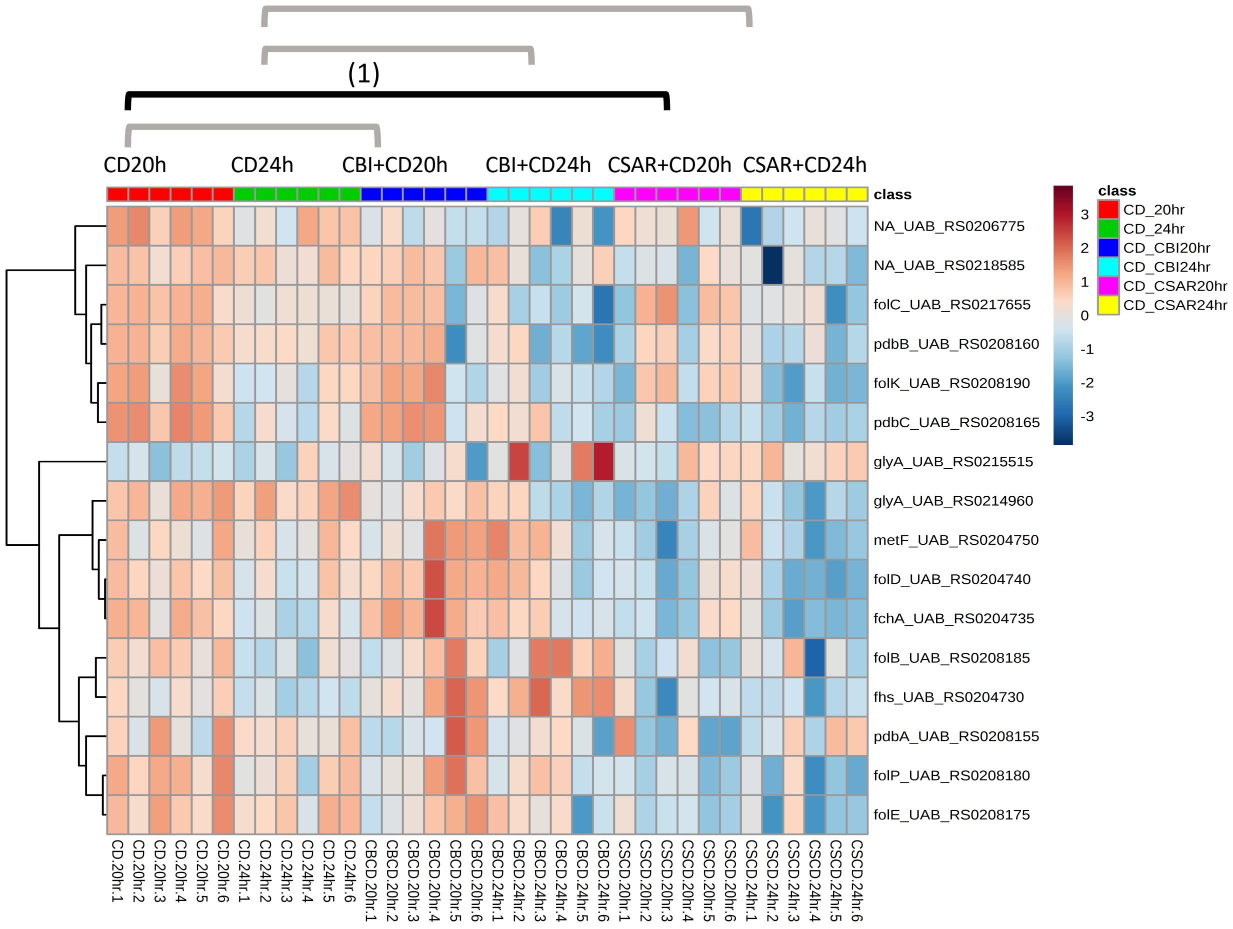


**Significantly Enriched Conditions:**

(1) Enriched in *C. difficile* mono-colonized vs CSAR+*C. difficile* mice at 20h (p=3.69E-04).

### Vitamin and Micronutrient Metabolism

#### 5.29 Nitrogenous compounds Transport & Metabolism

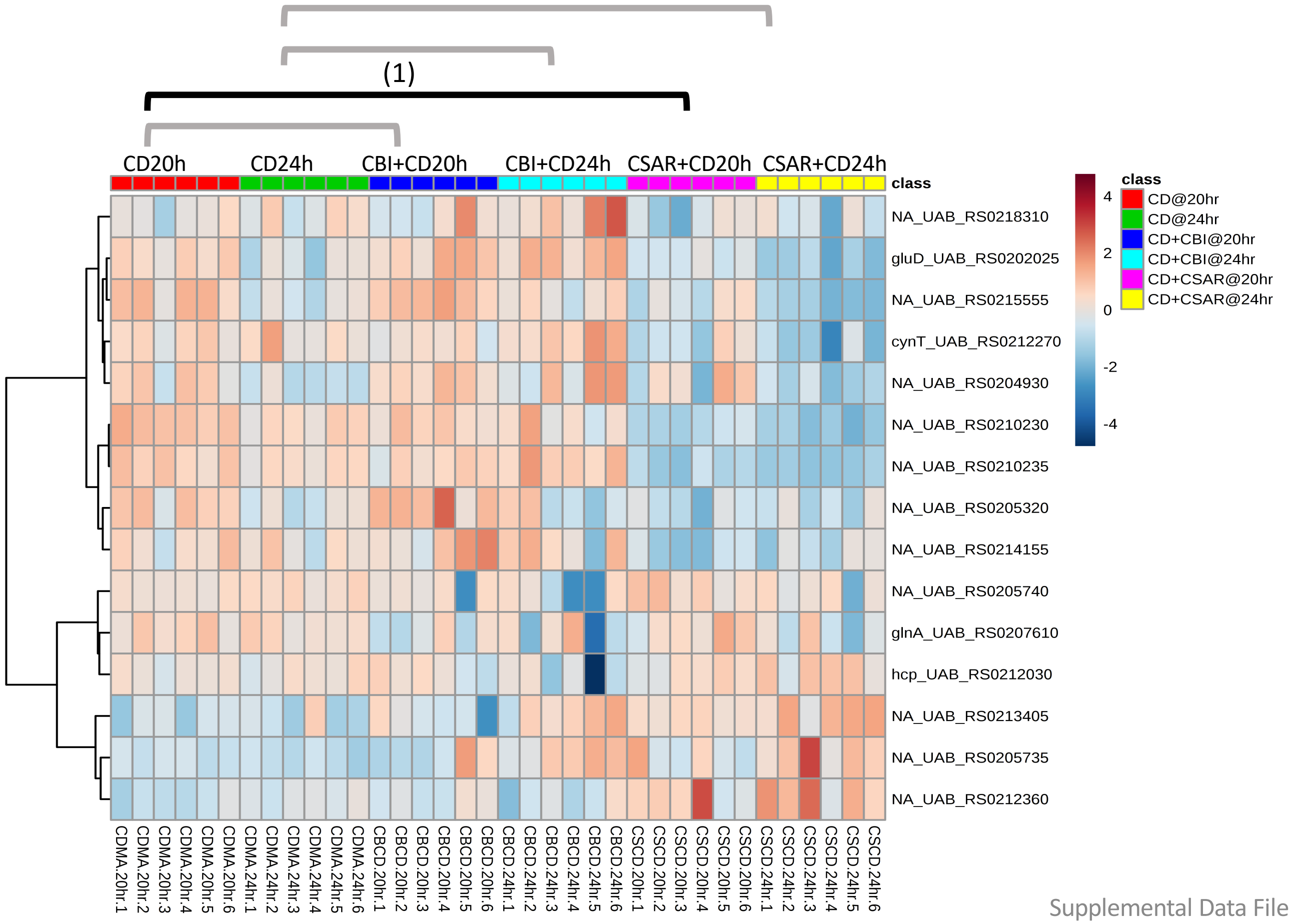


**Significantly Enriched Conditions:**

(1) Enriched in *C. difficile* mono-colonized vs CSAR+*C. difficile* mice at 20h (p=2.32E-02).

### Vitamin and Micronutrient Metabolism

#### 5.30 Phosphorous & Phosphonate Transport & Metabolism

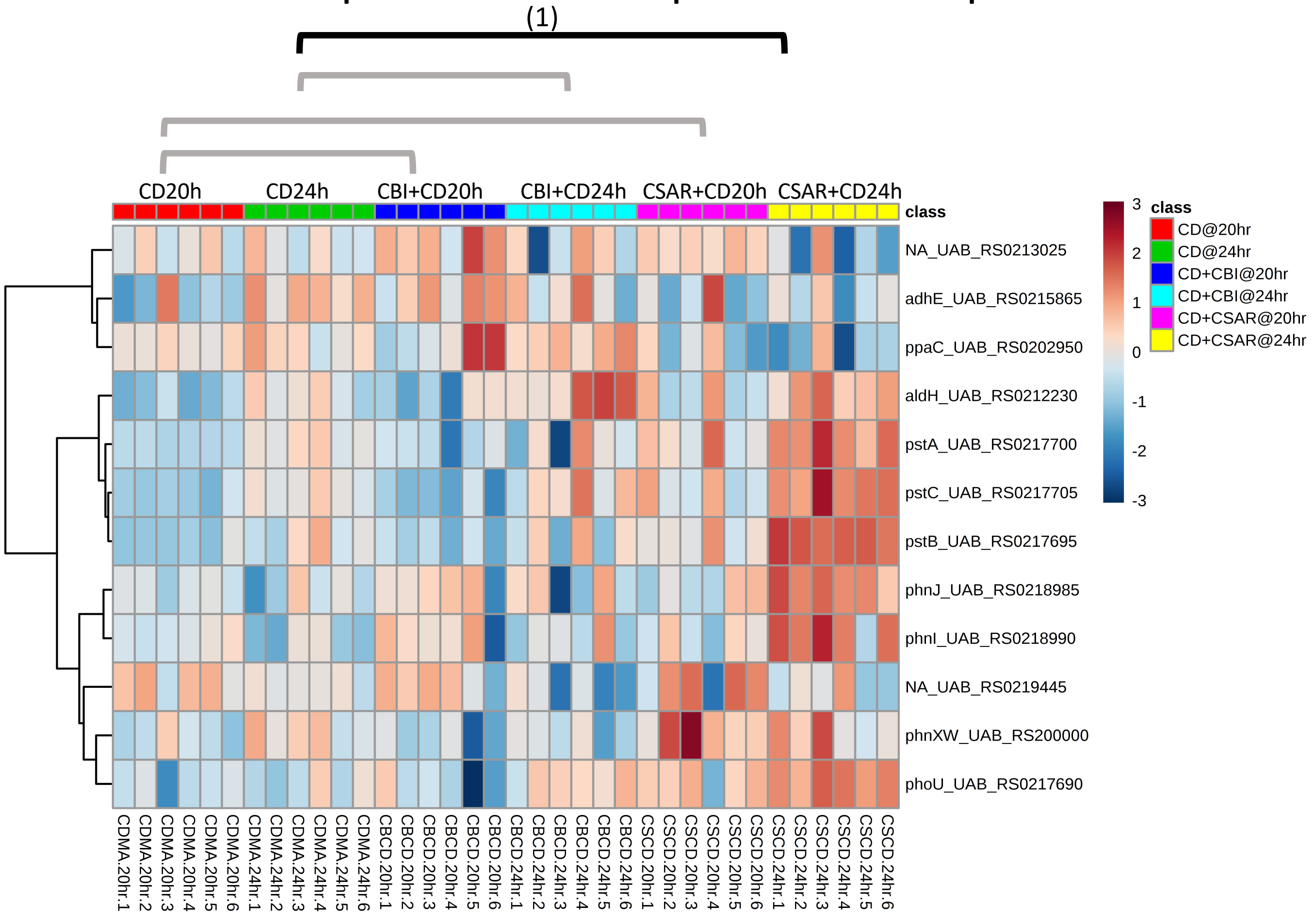


**Significantly Enriched Conditions:**

(1) Enriched in CSAR+*C. difficile vs C. difficile* mono-colonized mice at 24h (p=1.28E-02).

### Vitamin and Micronutrient Metabolism

#### 5.31 Sulphur-containing Compound Transport & Metabolism

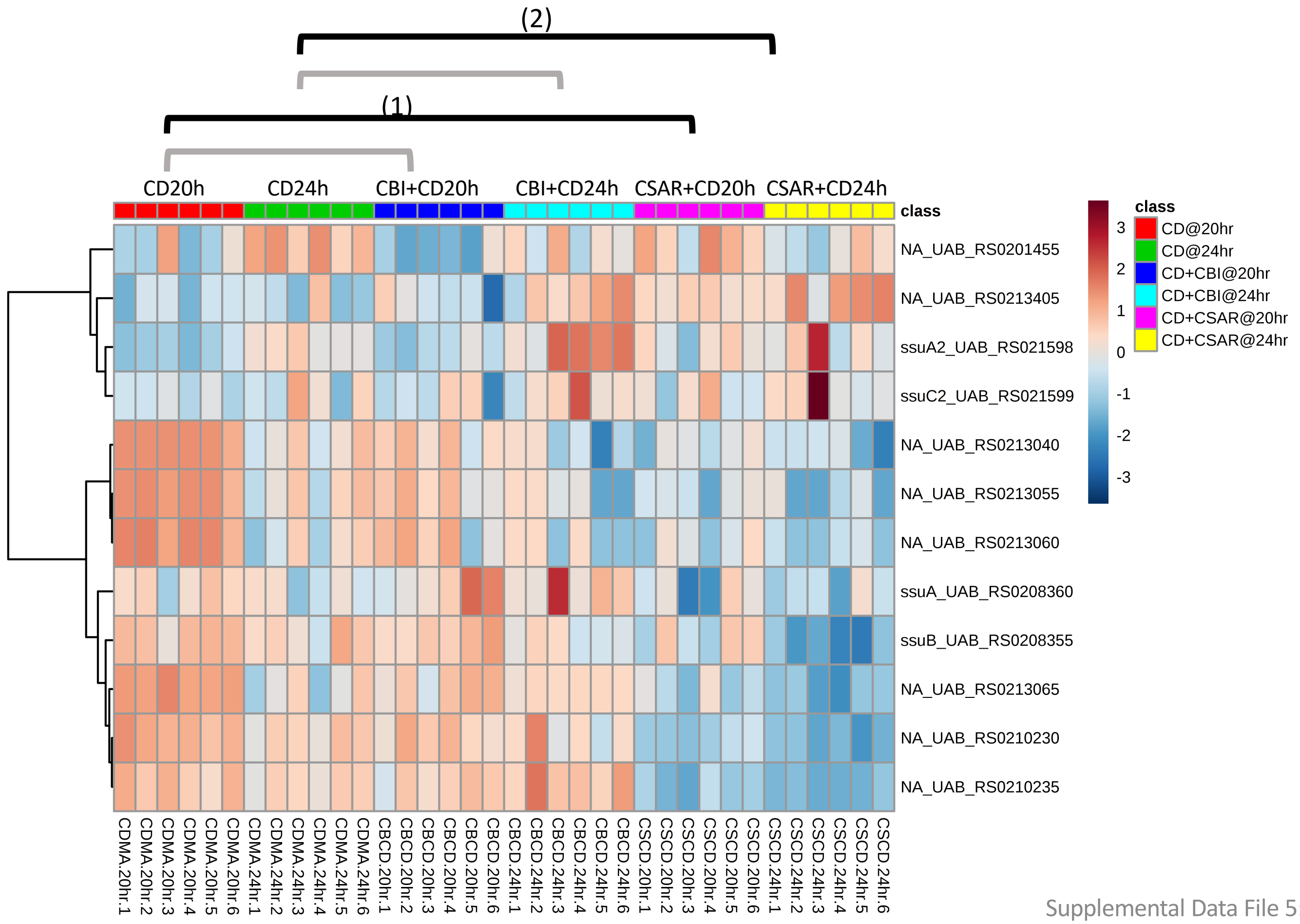


**Significantly Enriched Conditions:**

(1) Enriched in *C. difficil*e mono-colonized vs CSAR+*C. difficile* mice at 20h (p=1.78E-02).

(2) Enriched in *C. difficil*e mono-colonized vs CSAR+*C. difficile* mice at 24h (p=2.81E-03).

### Biosynthetic Pathways

#### 5.32 Cysteine, Homocysteine, Methionine Biosynthesis

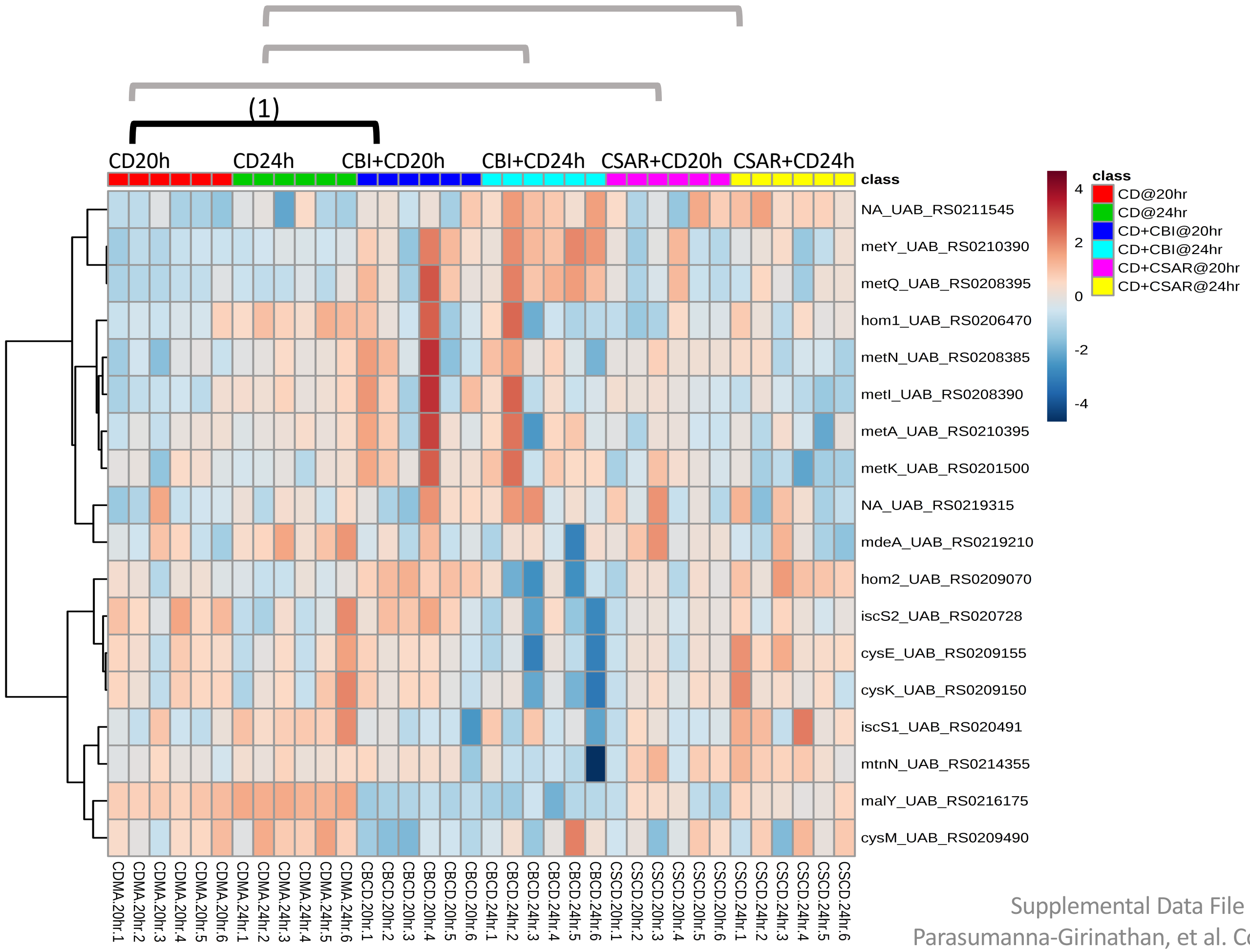


**Significantly Enriched Conditions:**

(1) Enriched in CBI+*C. difficile vs C. difficile* mono-colonized mice at 20h (p=3.94E-04).

### Biosynthetic Pathways

#### 5.33 Fatty Acid Biosynthesis

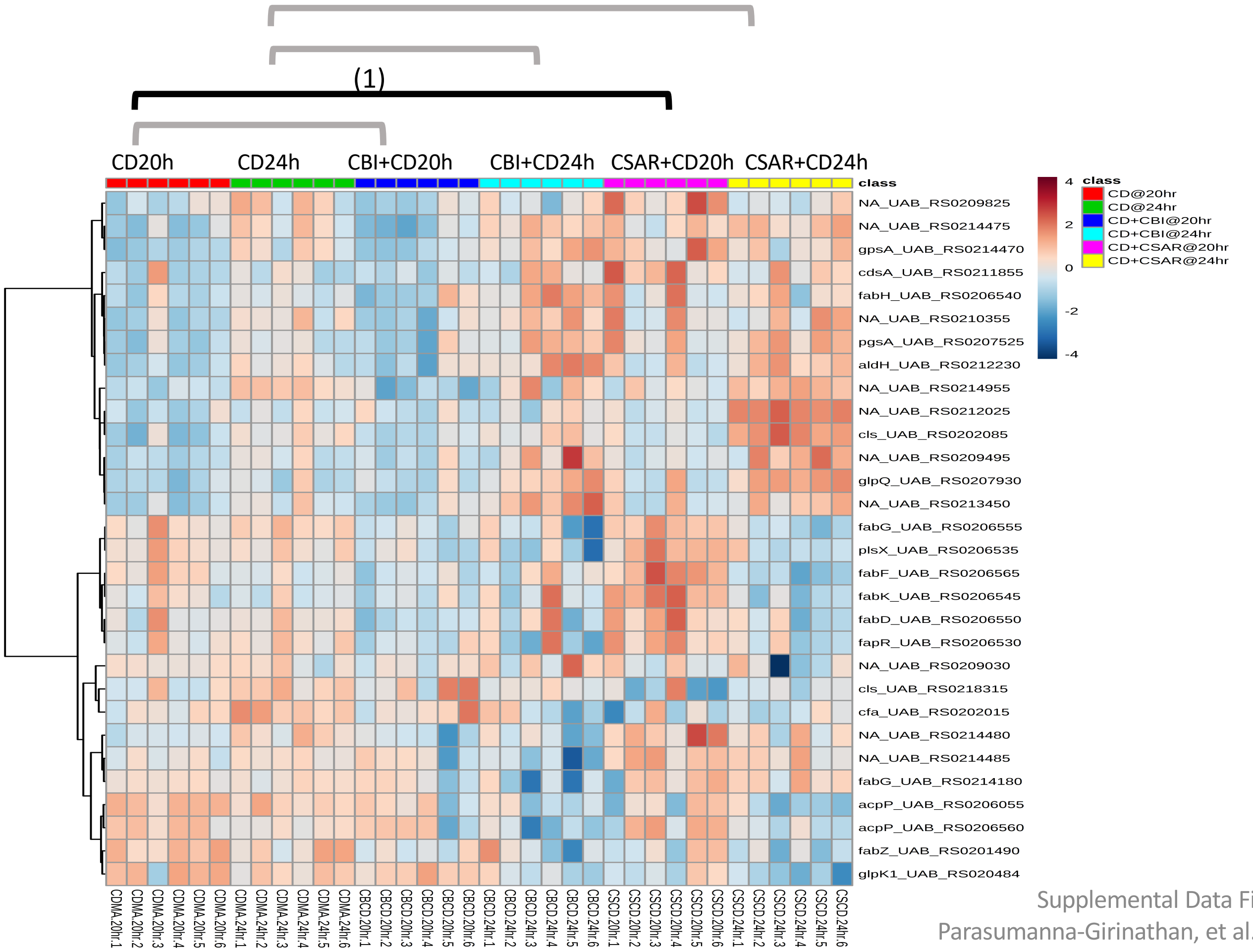


**Significantly Enriched Conditions:**

(1) Enriched in CSAR+*C. difficile vs C. difficile* mono-colonized mice at 20h (p=2.16E-02).

### Biosynthetic Pathways

#### 5.34 Histidine Biosynthesis

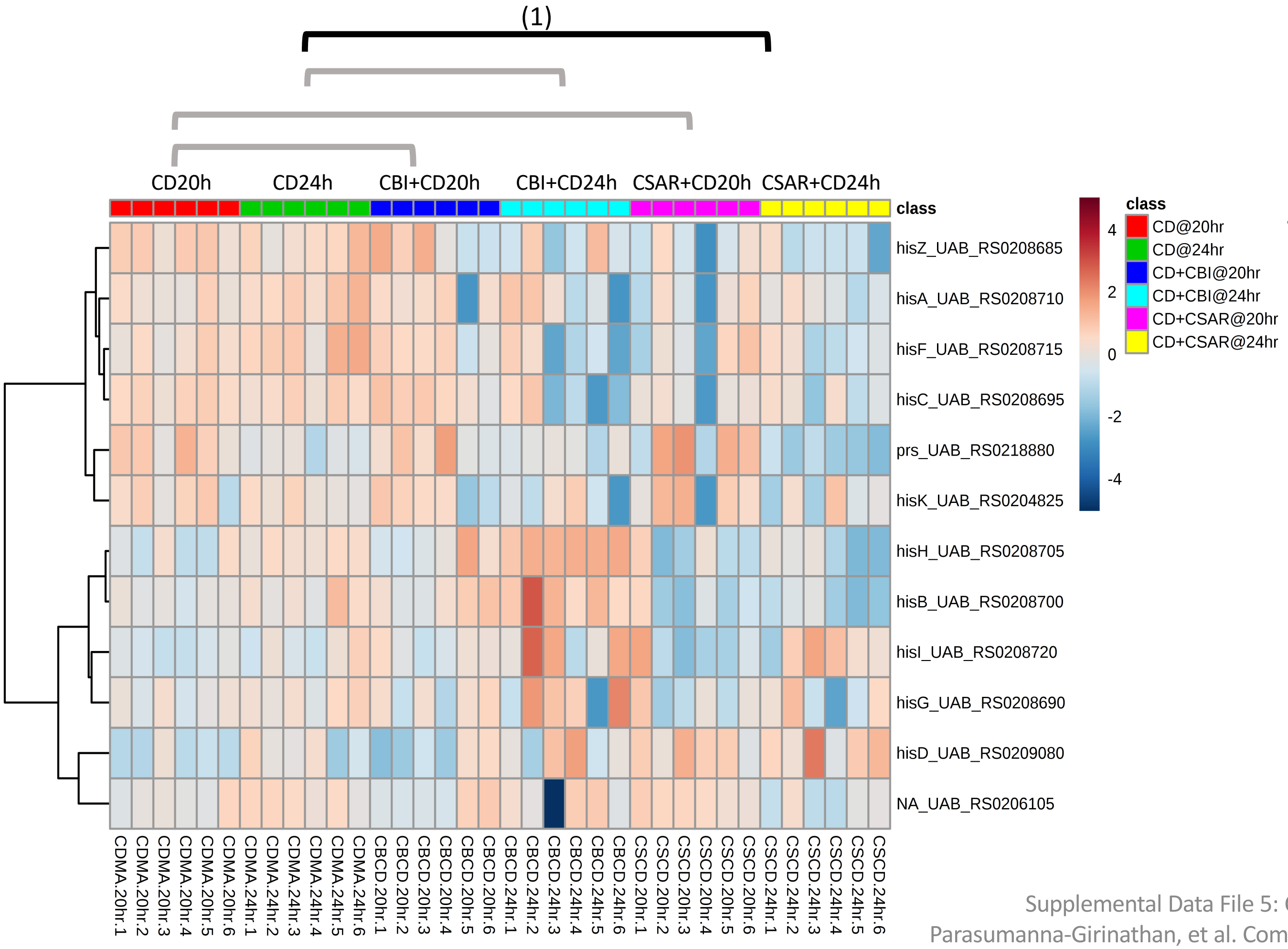


#### Significantly Enriched Conditions:

(1) Enriched in *C. difficile* mono-colonized vs CSAR+*C. difficile* mice at 24h (p=1.70E-02).

### Biosynthetic Pathways

#### 5.35 Terpenoid Backbone Biosynthesis

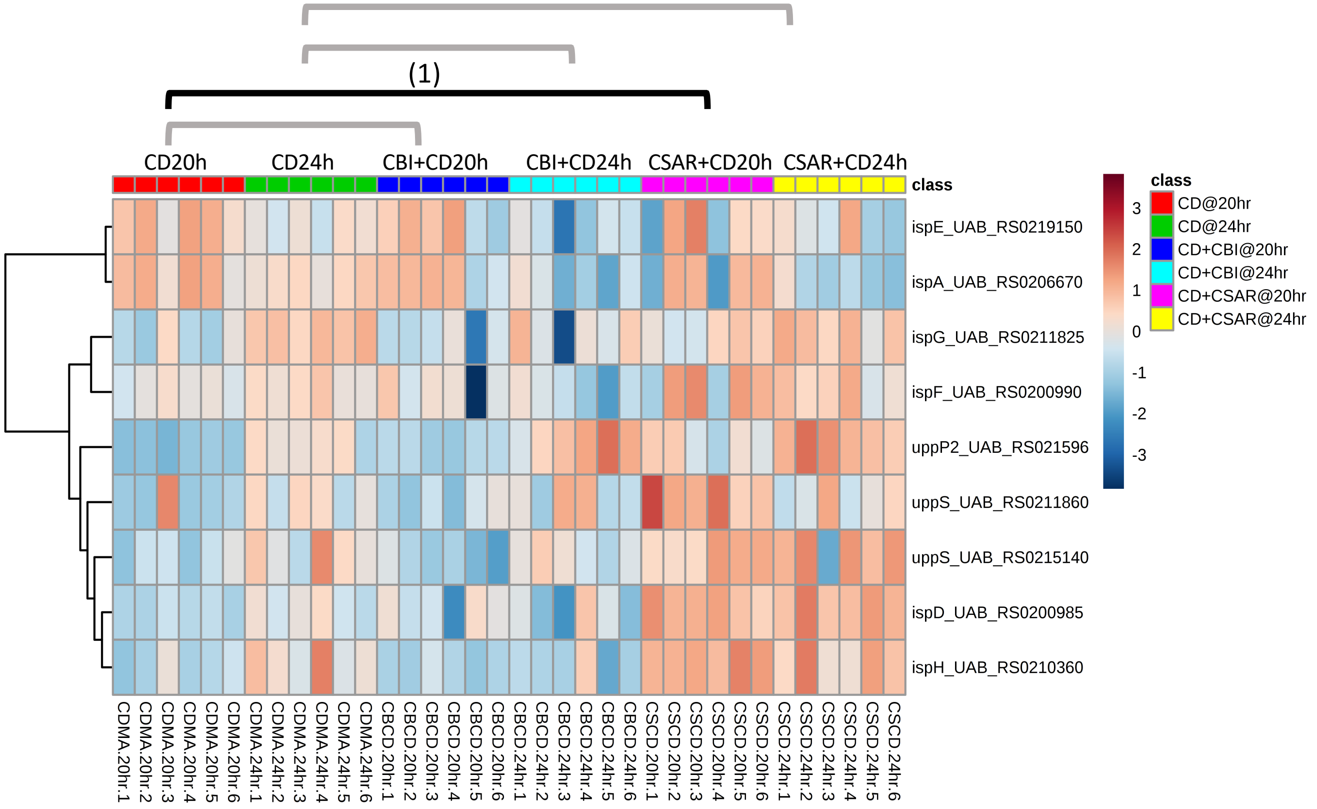


**<u>Significantly Enriched Conditions:</u>**

(1) Enriched in CSAR+*C. difficile vs C. difficile* mono-colonized mice at 20h (p=8.85E-03).

### Other Cellular Transports

#### 5.36 ABC Transport systems

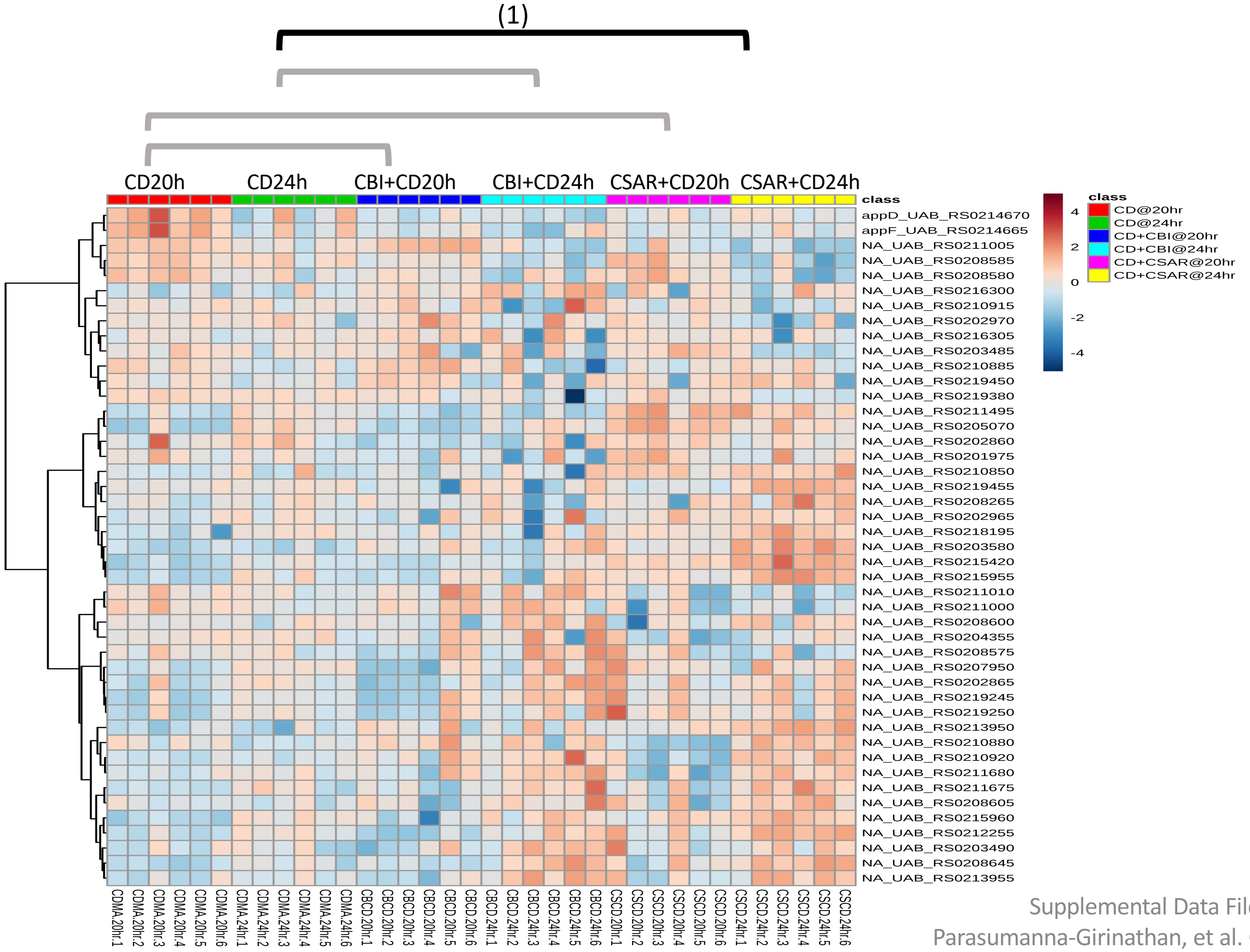


**Significantly Enriched Conditions:**

(1) Enriched in CSAR+*C. difficile vs C. difficile* mono-colonized mice at 24h (p=4.41E-02).

### Stress Responses

#### 5.37 Chemotaxis

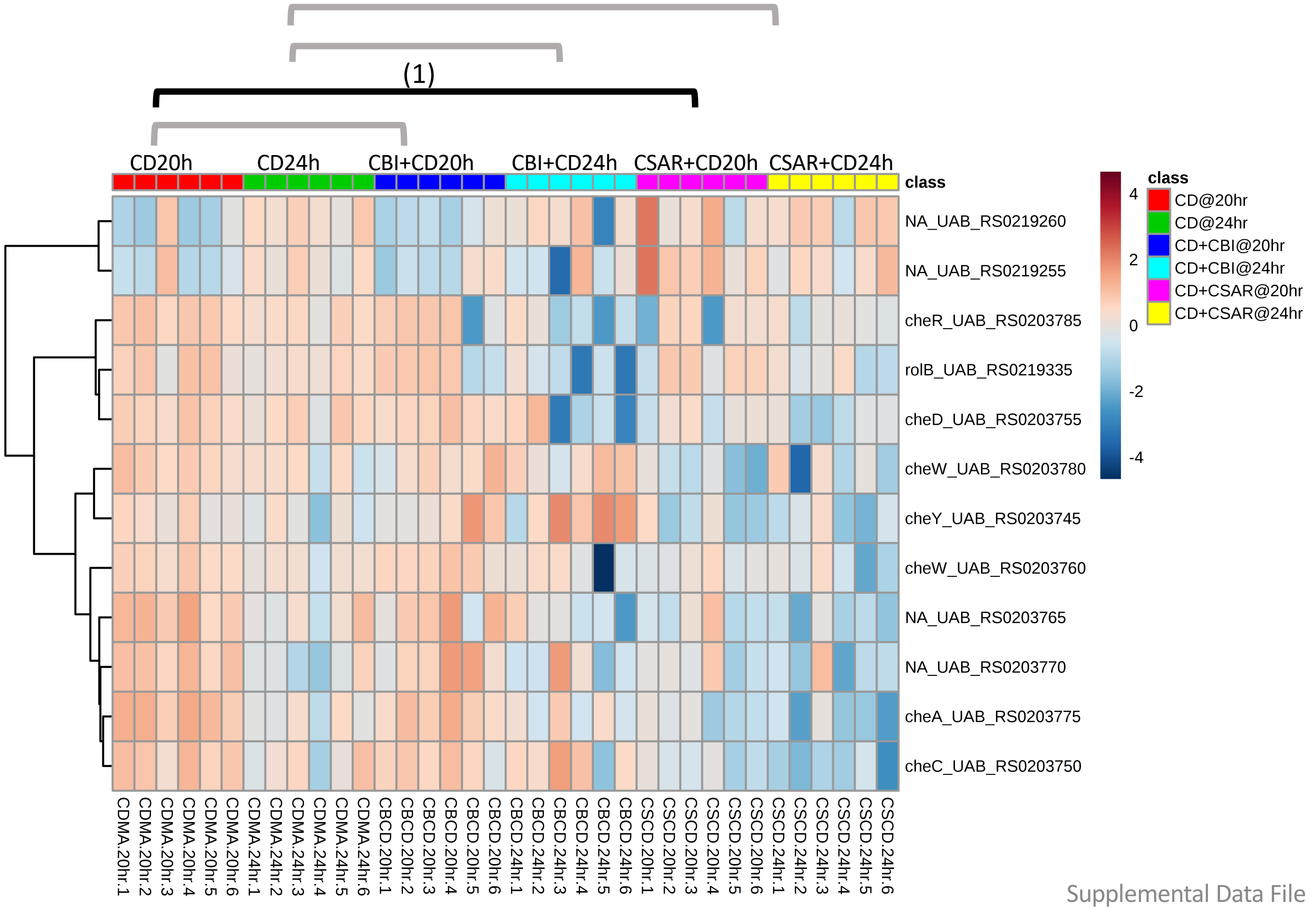


**<u>Significantly Enriched Conditions:</u>**

(1) Enriched in *C. difficile* mono-colonized vs CSAR+*C. difficile* mice at 20h (p=3.21E-02).

### Stress Responses

#### 5.38 CRISPR System

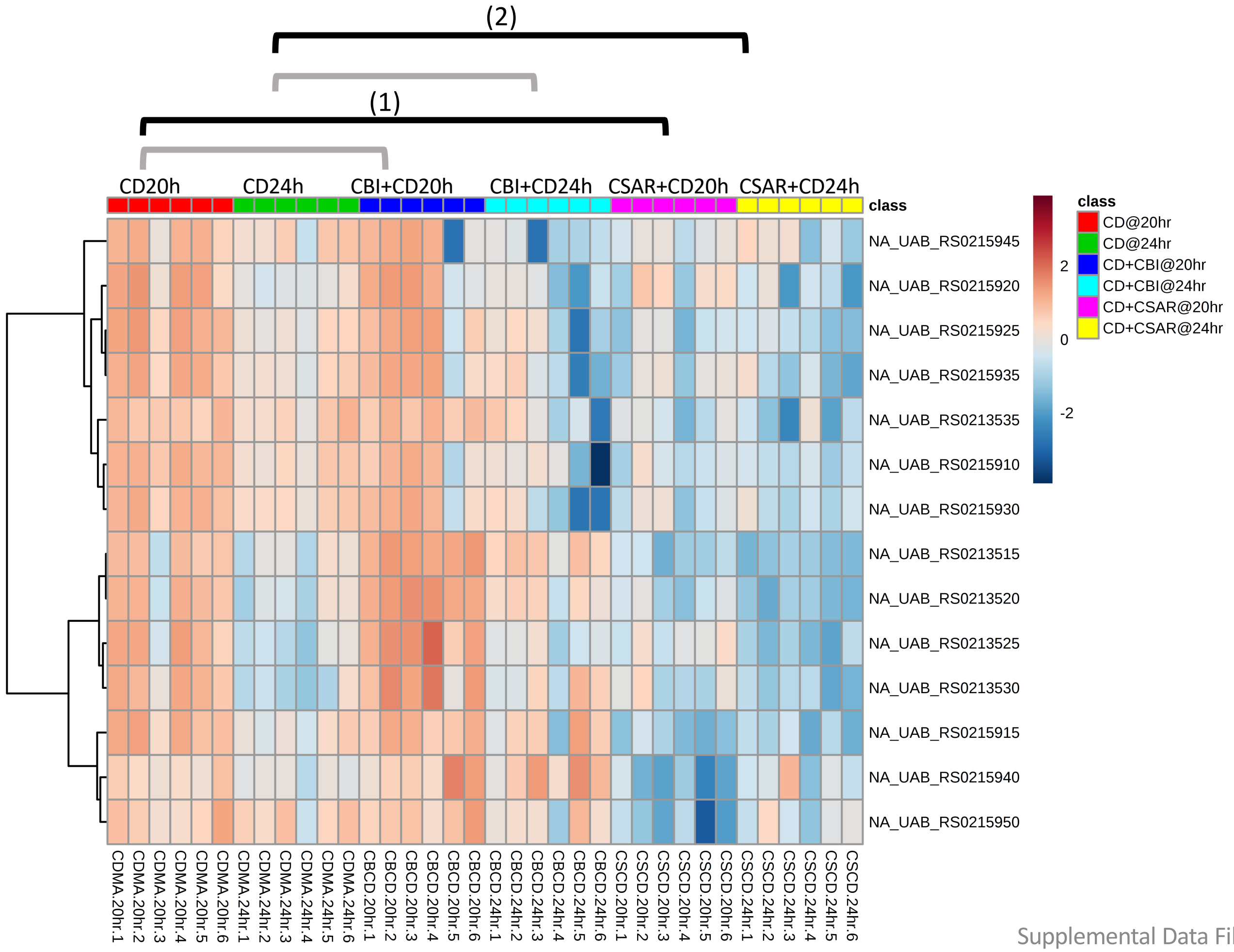


**Significantly Enriched Conditions:**

(1) Enriched in *C. difficile* mono-colonized vs CSAR+*C. difficile* mice at 20h (p=2.86E-11).

(2) Enriched in *C. difficile* mono-colonized vs CSAR+*C. difficile* mice at 24h (p=2.81E-03).

### Stress Responses

#### 5.39 Diffocin Locus Proteins

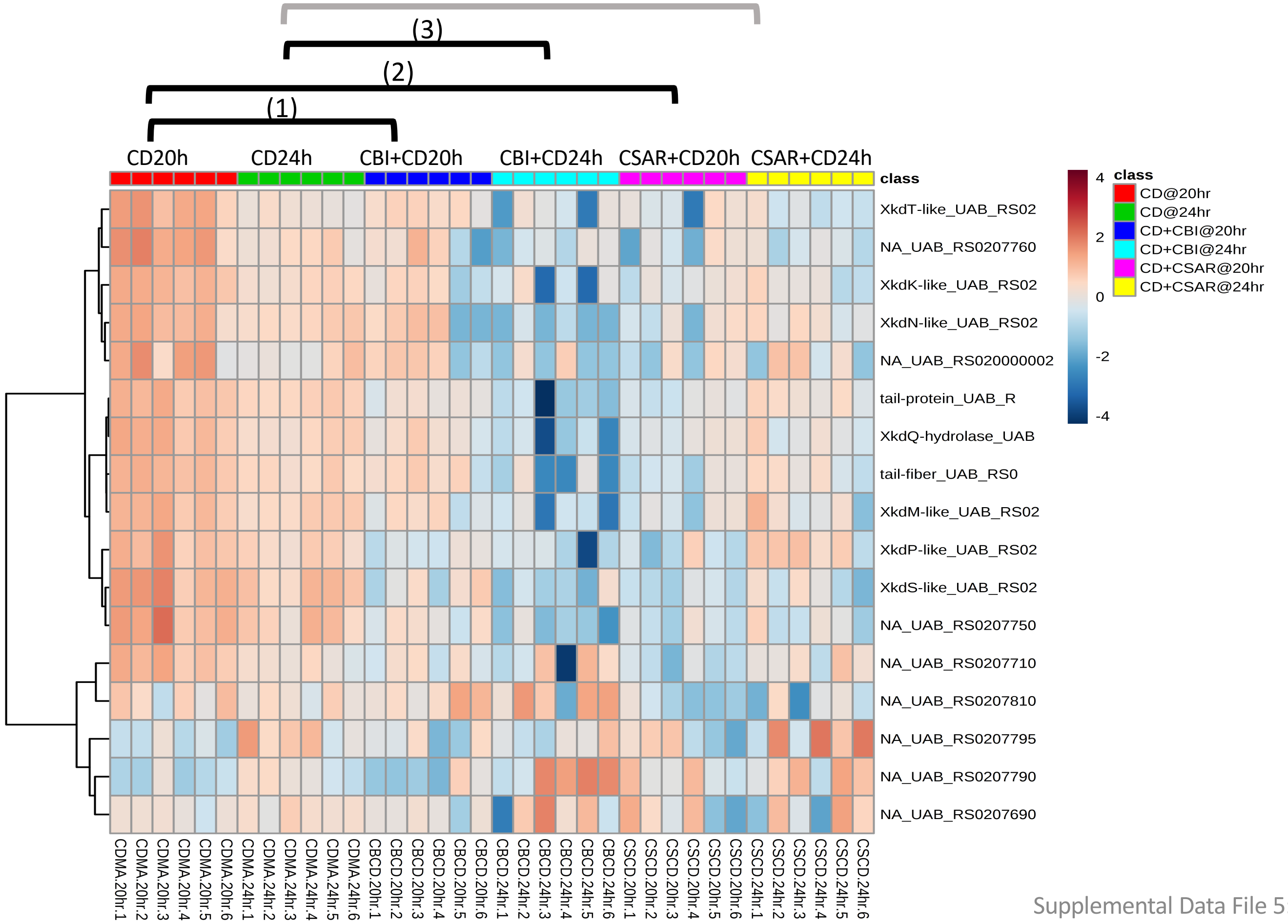


**Significantly Enriched Conditions:**

(1) Enriched in *C. difficile* mono-colonized vs CBI+*C. difficile* mice at 20h (p=6.20E-12).

(2) Enriched in *C. difficile* mono-colonized vs CSAR+*C. difficile* mice at 20h (p=6.38E-10).

(3) Enriched in *C. difficile* mono-colonized vs CBI+*C. difficile* mice at 24h (p=4.72E-04).

### Stress Responses

#### 5.40 Oxidative Stress Responses

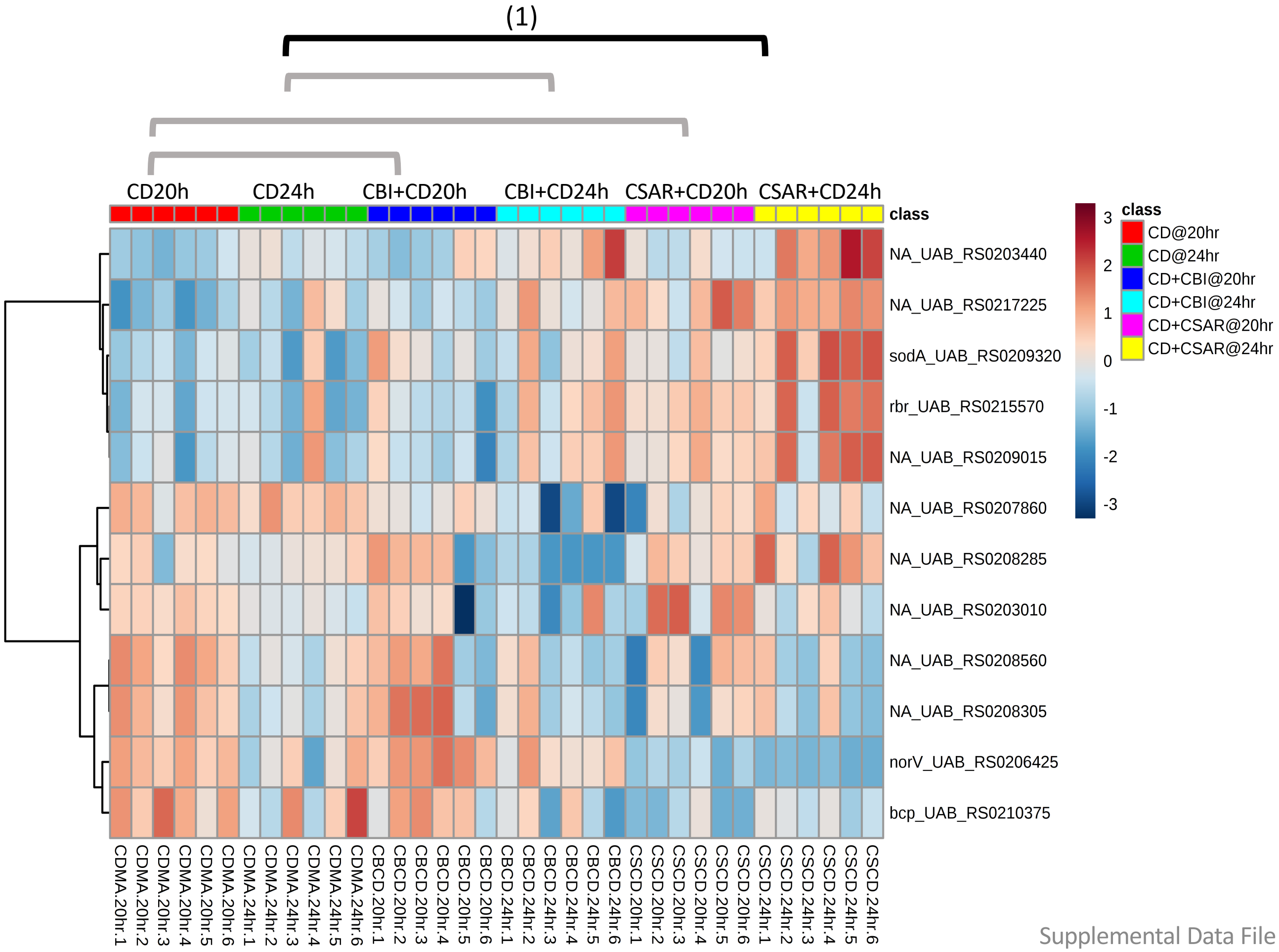


**Significantly Enriched Conditions:**

(1) Enriched in CSAR+*C. difficile vs C. difficile* mono-colonized mice at 24h (p=3.79E-02).

### Stress Responses

#### 5.41 Pathogenicity Island/Virulence Determinants

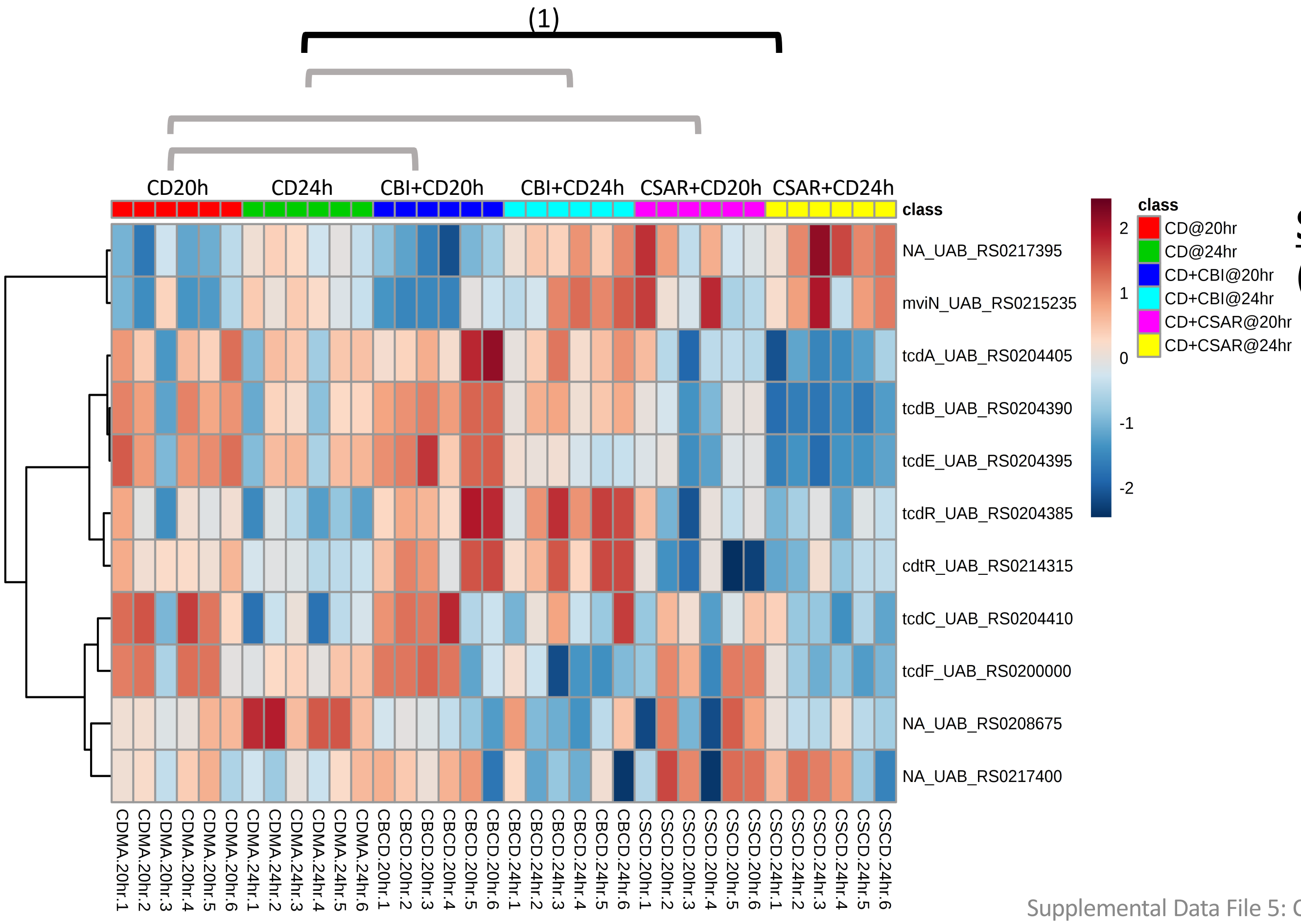


**<u>Significantly Enriched Conditions:</u>**

(1) Enriched in *C. difficile* mono-colonized vs CSAR+*C. difficile* mice at 24h (p=4.07E-02).

### Stress Responses

#### 5.42 Sporulation

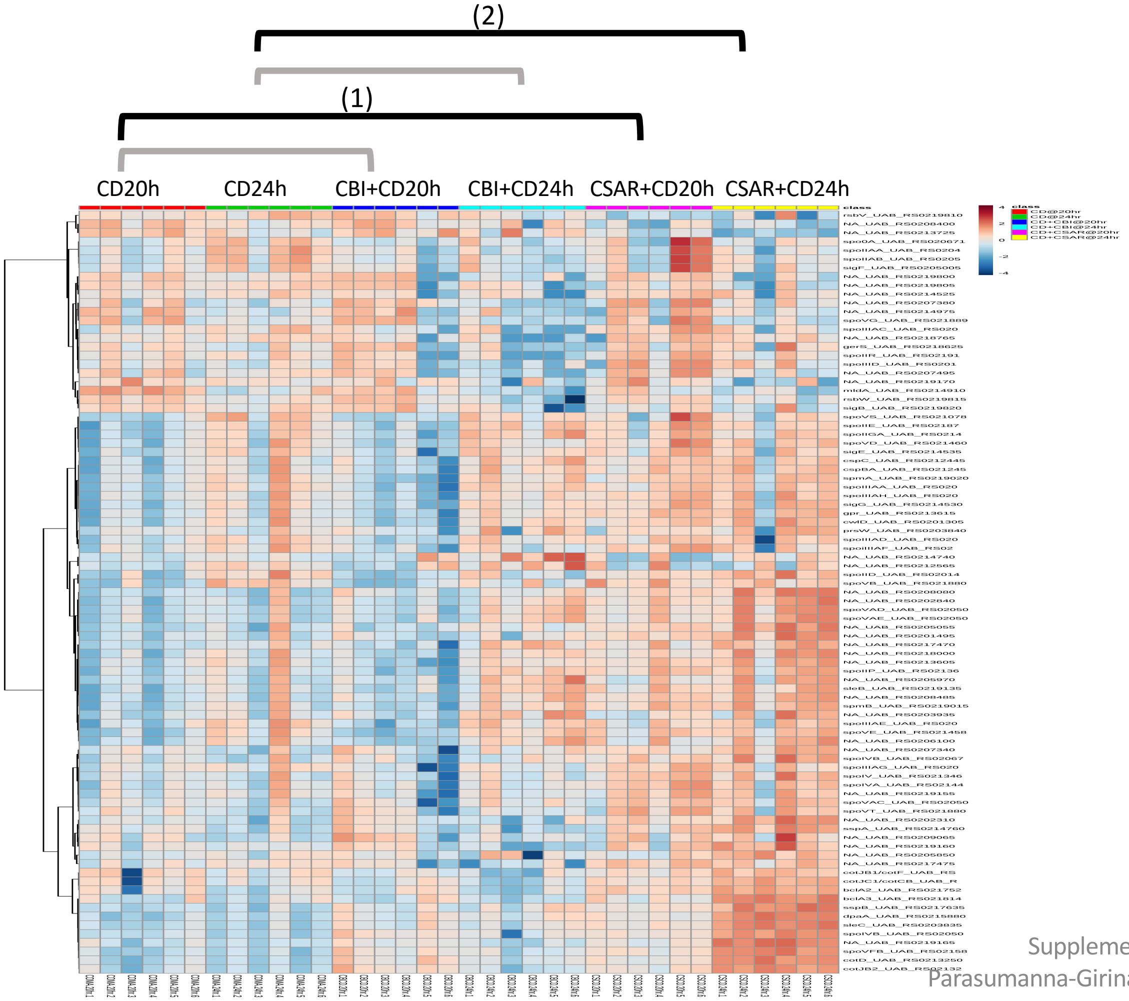


**Significantly Enriched Conditions:**

(1) Enriched in CSAR+*C. difficile vs C. difficile* mono-colonized mice at 20h (p=6.92E-16).

(2) Enriched in CSAR+*C. difficile vs C. difficile* mono-colonized mice at 24h (p=1.63E-06).
