## Supplemental Data File 6 for "*In vivo* commensal control of *Clostridioides difficile* virulence"

• Version 1.0; 12/23/2019

### Supplemental Data File 6: Transcriptome Analyses: *C. sardiniense* Enriched Pathways

**Pages:** Introduction, this page
**Page 1**: Sample Key


**Carbon Source Transport and Metabolism**
2: 6.1 Ascorbate Transport and Metabolism
3: 6.2 Beta-glucoside Transport and Conversion
4: 6.3 Dipeptide and Oligopeptide Transport
5: 6.4 Ethanolamine Utilization
6: 6.5 Mucin Degradation and Metabolism
7: 6.6 Polysaccharide metabolism
8: 6.7 Propanediol Metabolism
9: 6.8 Tagatose and Galactose Transport and Conversion
10: 6.9 Transport Binding Proteins: Amino Acids and Amines
11: 6.10 Xanthine, Hypoxanthine Transport and Metabolism

**Core Cellular Machinery**
12: 6.11 ATP Synthesis
13: 6.12 Electron Transport
14: 6.13 Hydrogenase Systems
15: 6.14 Ribosome Synthesis and Modification

**Cellular Adhesion and Motility**
16: 6.15 Cell Surface Adhesive Organelles

**Vitamin and Micronutrient Metabolism**
17: 6.16 Cobalamin Biosynthesis
18: 6.17 Folic Acid Biosynthesis
19: 6.18 Nickel Transport and Metabolism
20: 6.19 Phosphorus Transport and Metabolism
21: 6.20 Transport Binding Proteins: Cations

**Biosynthetic Pathways**
22: 6.21 Cysteine and Methionine Biosynthesis
23: 6.22 Fatty Acid Biosynthesis

**Bacteriophage and Mobile Elements**
24: 6.23 Putative Phage Locus 1
25: 6.24 Putative Phage Locus 3
26: 6.25 Putative Phage Locus 4

**Stress Responses**
27: 6.26 Sporulation

### SDF6: Sample Key

6.2 Beta-glucoside Transport and Metabolism

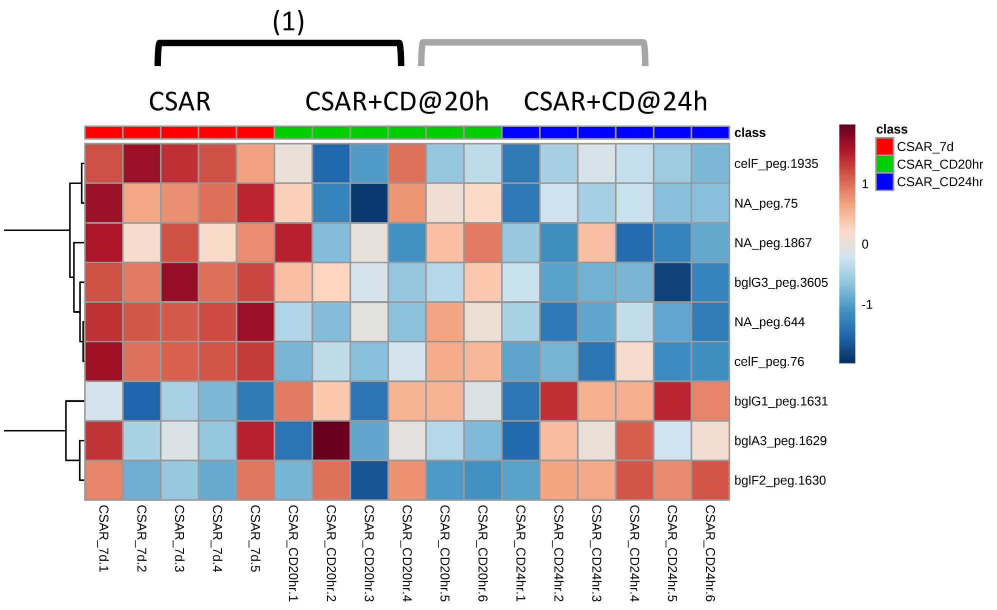


**Guide:**

- Group showing the enriched category is at the top ("6.2 Beta-glucoside Transport and Metabolism")
- Brackets indicate the 2 comparisons evaluated among groups for significant enrichment of genes in the pathway
- Brackets in black show significant comparisons
- Significant comparisons and p values are shown in the text box on the right-hand side

**Significantly Enriched Conditions:**

(1) Enriched in CSAR mono-colonized mice vs CSAR+*C. difficile* co-colonized mice at 20h (p=0.0122061).

### Carbon Source Transport and Metabolism

#### 6.1 Ascorbate Transport and Metabolism

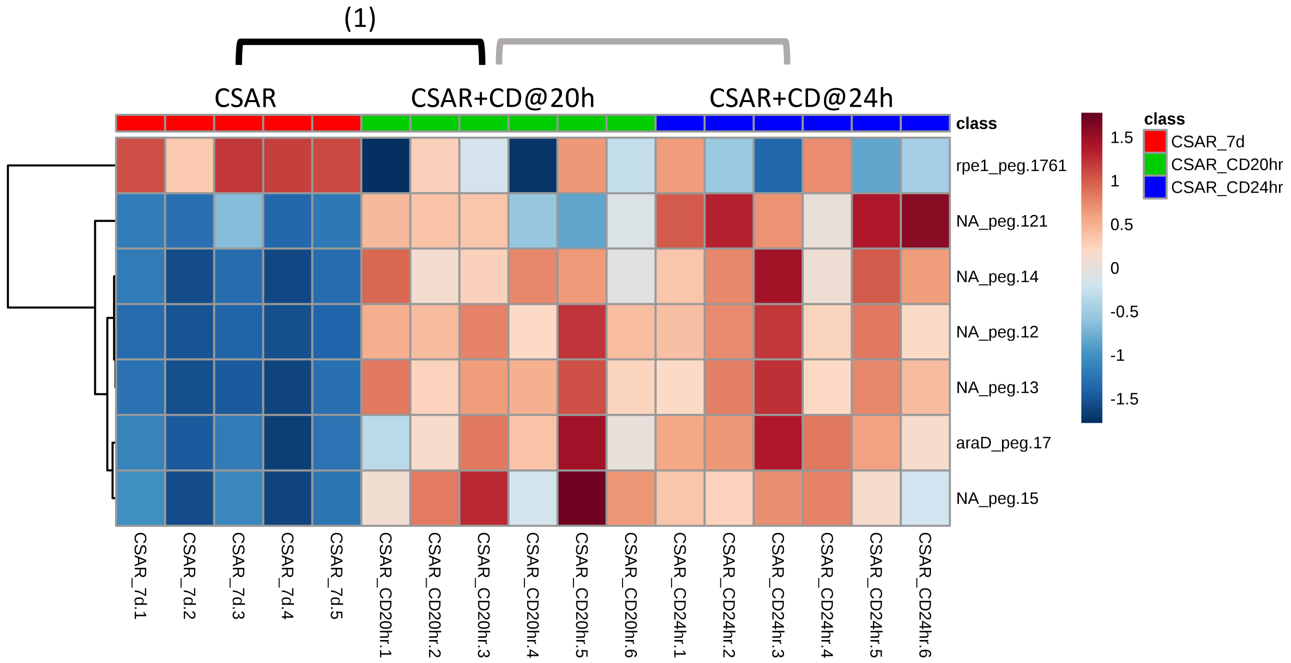


**Significantly Enriched Conditions:**

(1) Enriched in CSAR+*C. difficile* co-colonized *vs* CSAR mono-colonized mice at 20h (p=9.52E-04).

### Carbon Source Transport and Metabolism

#### 6.2 Beta-glucoside Transport and Metabolism

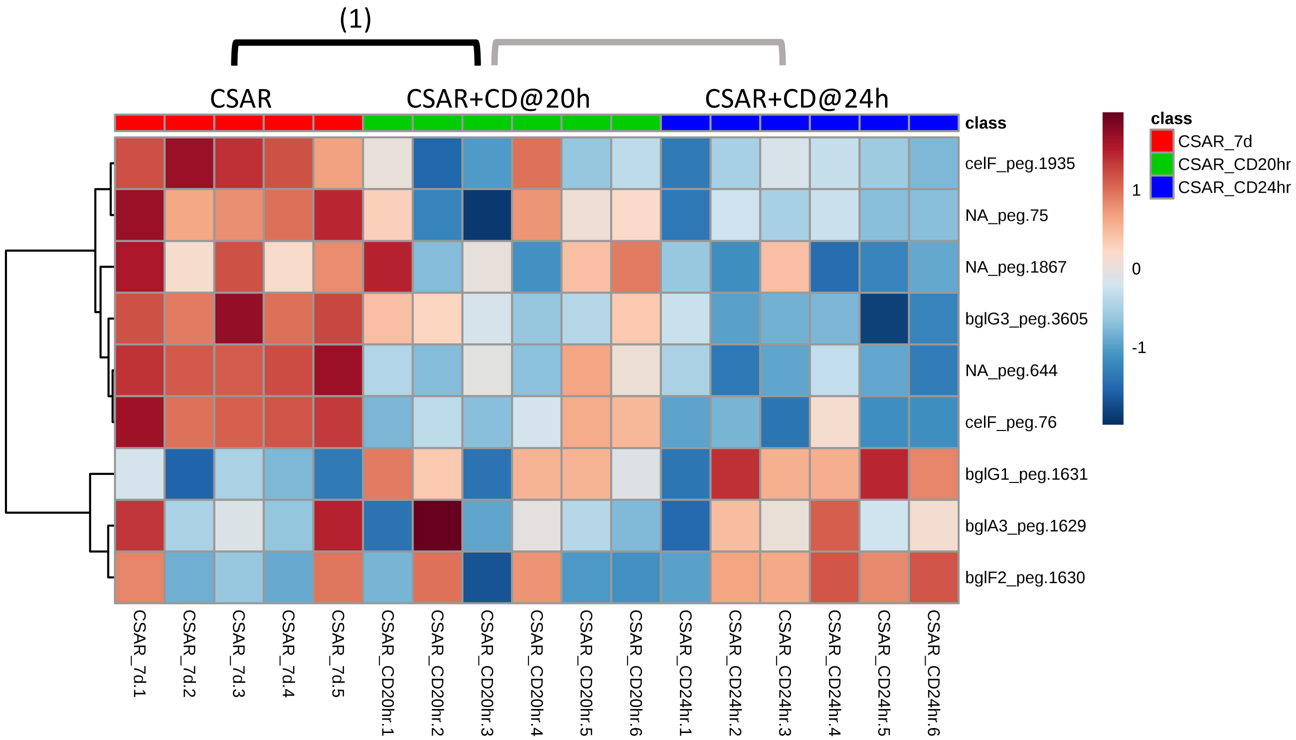


**<u>Significantly Enriched Conditions:</u>**

(1) Enriched in CSAR mono-colonized mice vs CSAR+*C. difficile* co-colonized mice at 20h (p=1.22E-02).

### Carbon Source Transport and Metabolism

#### 6.3 Dipeptide and Oligopeptide Transport

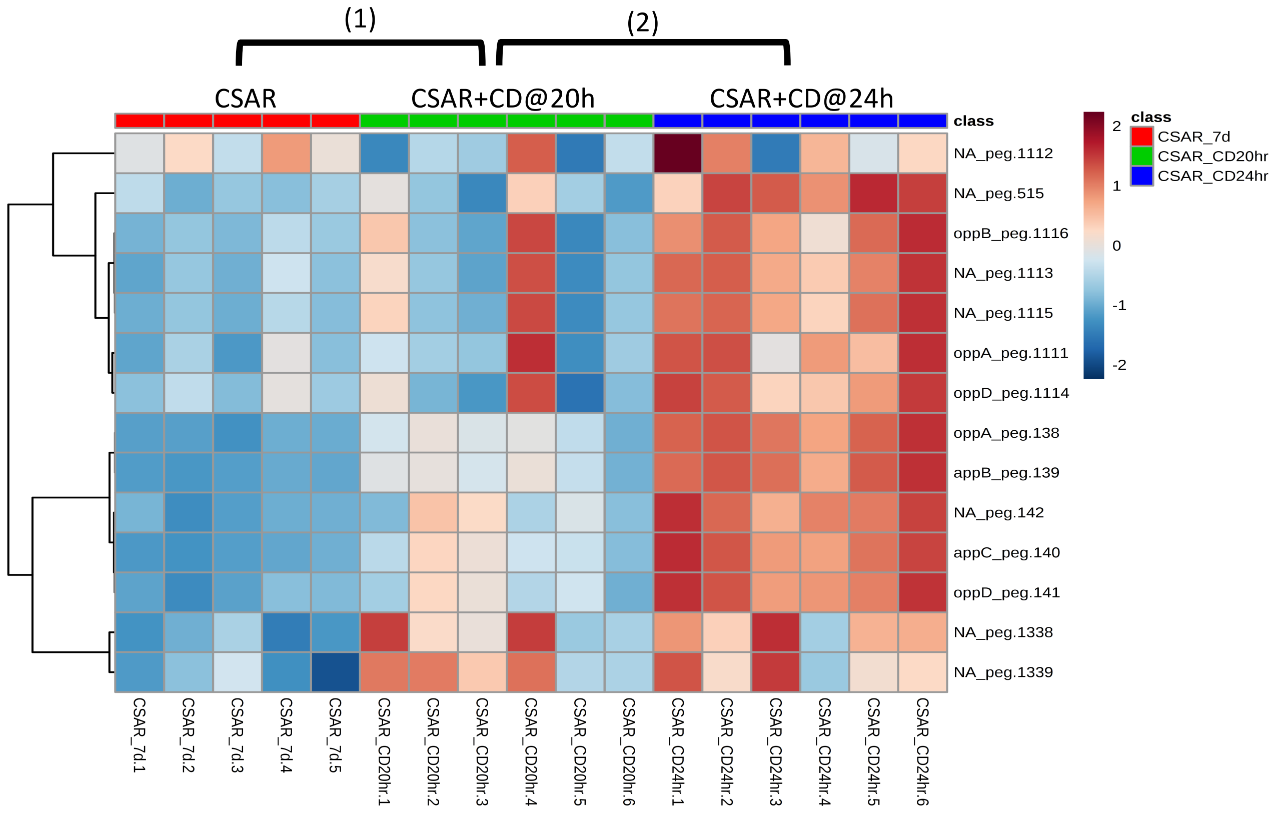


**Significantly Enriched Conditions:**

(1) Enriched in CSAR+*C. difficile* co-colonized *vs* CSAR mono-colonized mice at 20h (p=1.79E-03).

(2) Enriched in CSAR+*C. difficile* co-colonized at 24h *vs* CSAR +*C. difficile* co-colonized at 20h (p=3.42E-04).

### Carbon Source Transport and Metabolism

#### 6.4 Ethanolamine Utilization

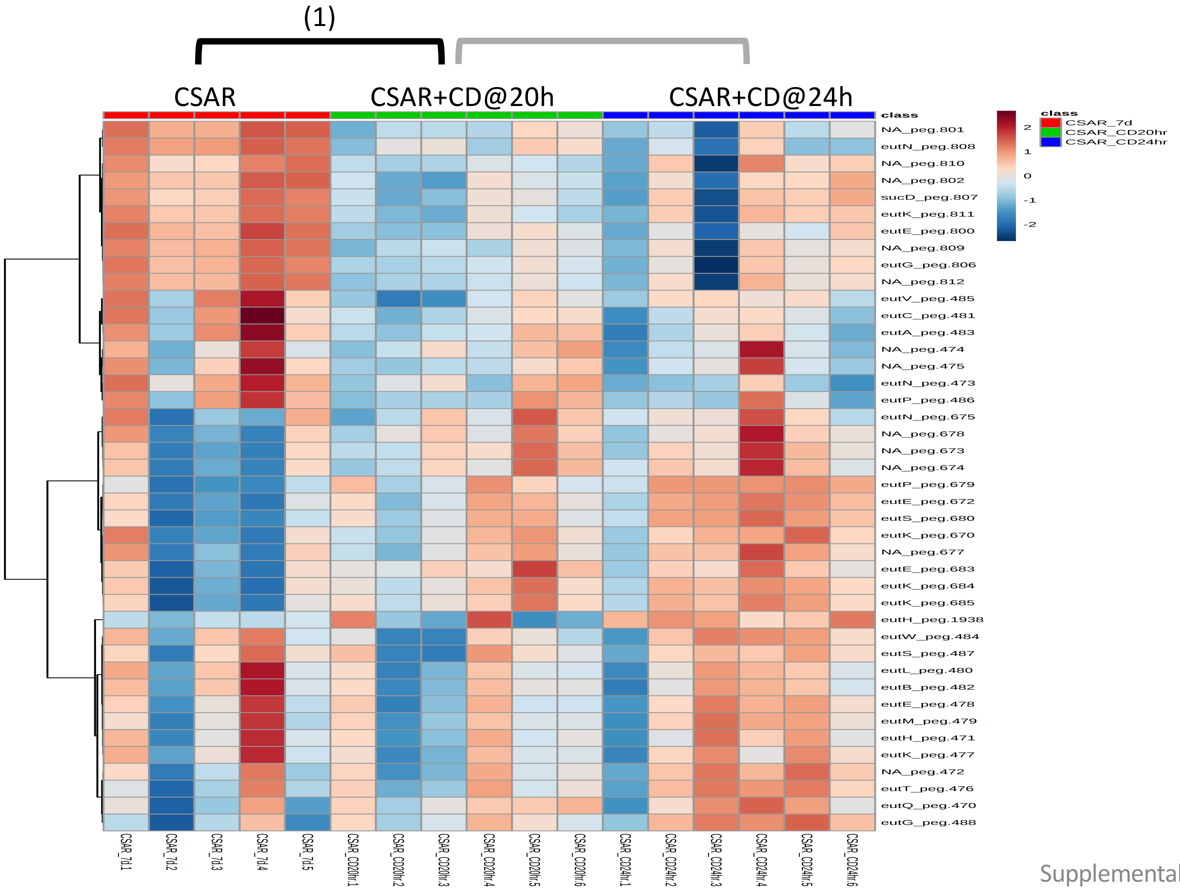


<u>**Significantly Enriched Conditions:**</u>

(1) Enriched in CSAR mono-colonized mice vs CSAR+*C. difficile* co-colonized mice at 20h (p=4.73E-03).

### Carbon Source Transport and Metabolism

#### 6.5 Mucin Degradation and Metabolism

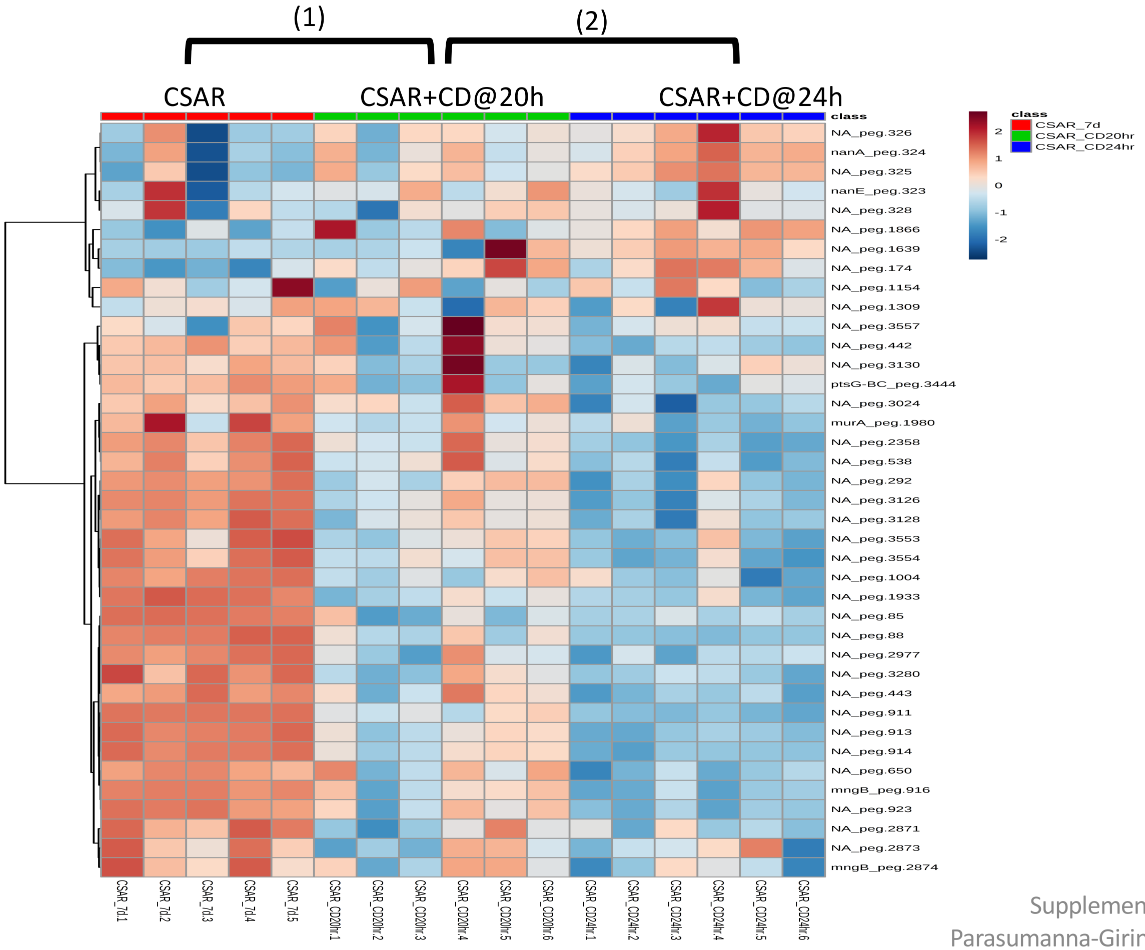


**Significantly Enriched Conditions:**

(1) Enriched in CSAR mono-colonized mice vs CSAR+*C. difficile* co-colonized mice at 20h (p=2.58E-06).

(2) Enriched in CSAR+*C. difficile* co-colonized at 20h *vs* CSAR +*C. difficile* co-colonized at 24h (p=6.27E-04).

### Carbon Source Transport and Metabolism

#### 6.6 Polysaccharide Metabolism

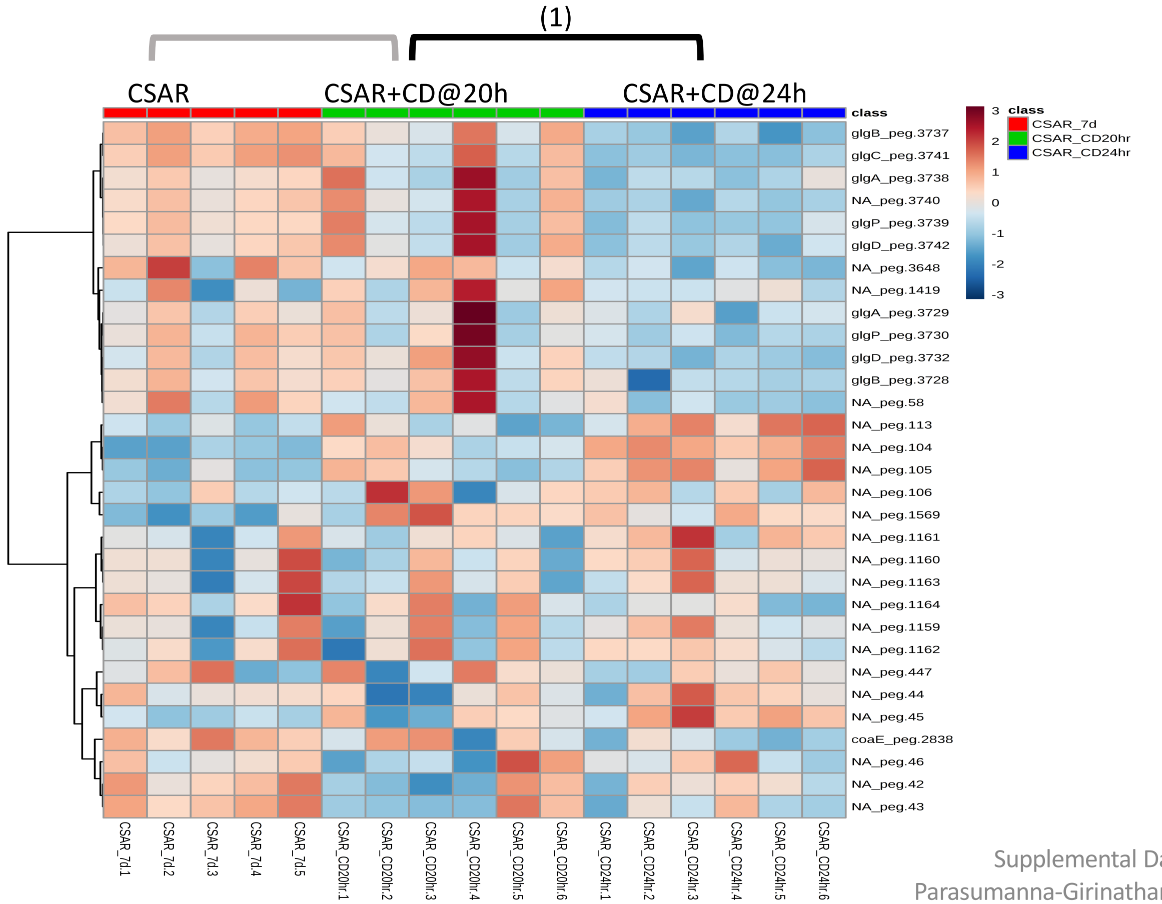


**Significantly Enriched Conditions:**

(1) Enriched in CSAR+*C. difficile* co-colonized at 20h *vs* CSAR +*C. difficile* co-colonized at 24h (p=2.15E-02).

### Carbon Source Transport and Metabolism

#### 6.7 Propanediol Metabolism

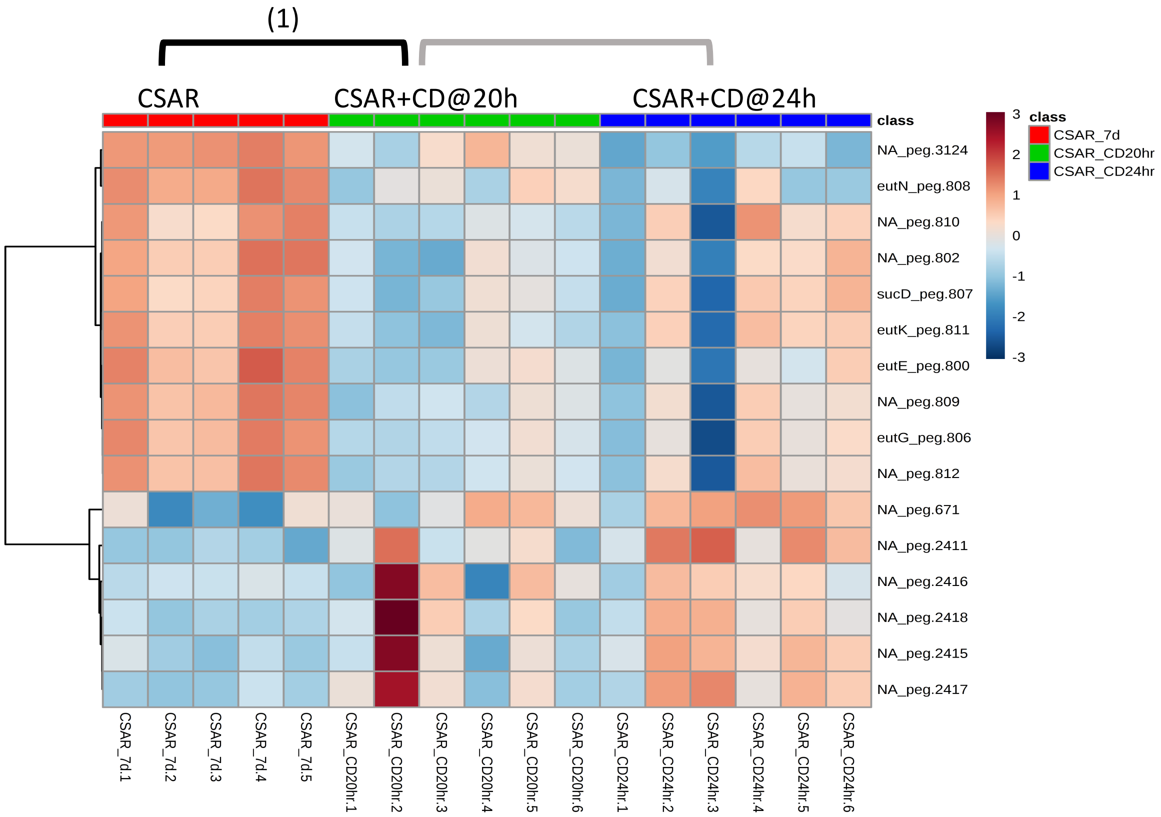


**Significantly Enriched Conditions:**

(1) Enriched in CSAR mono-colonized mice vs CSAR+*C. difficile* co-colonized mice at 20h (p=8.53E-05).

### Carbon Source Transport and Metabolism

#### 6.8 Tagatose, Galactose Transport and Metabolism

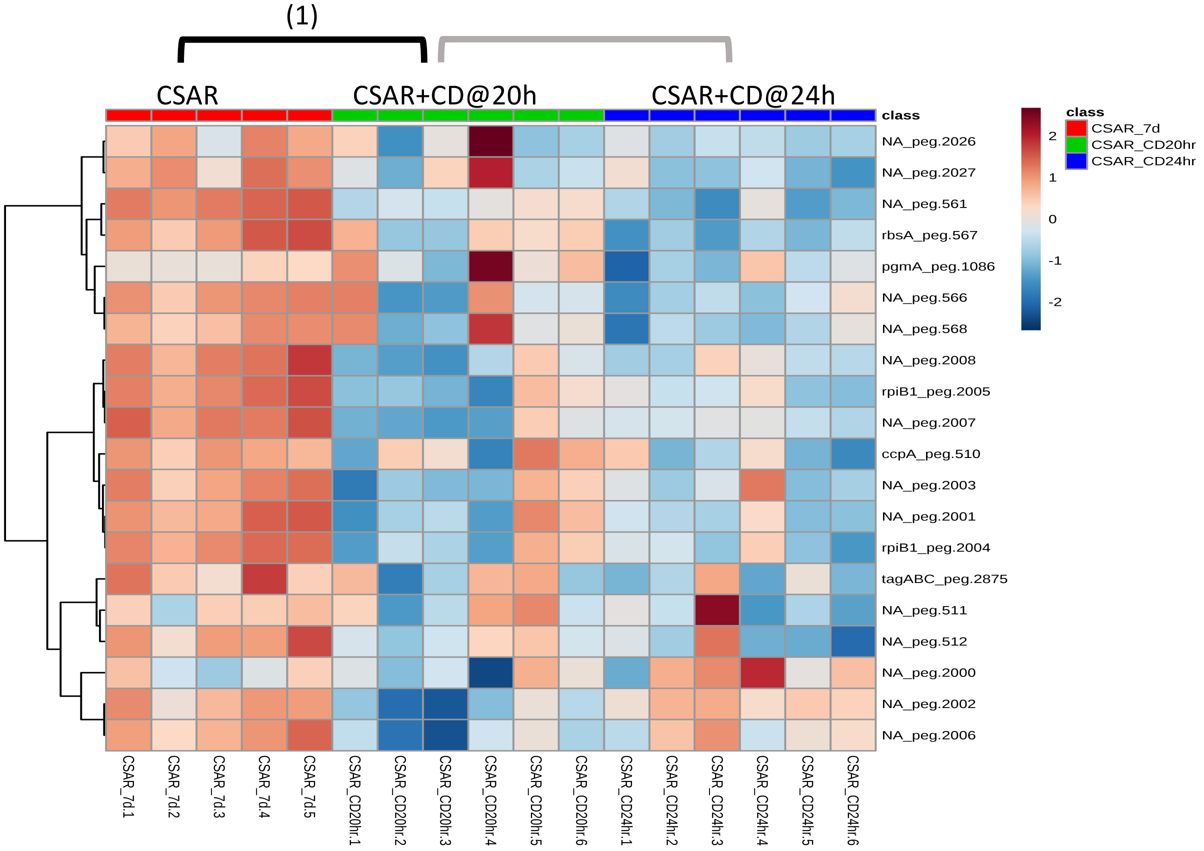


**Significantly Enriched Conditions:**

(1) Enriched in CSAR mono-colonized mice vs CSAR+*C. difficile* co-colonized mice at 20h (p=8.08E-04).

### Carbon Source Transport and Metabolism

#### 6.9 Transport Binding Proteins: Amino Acids and Amines

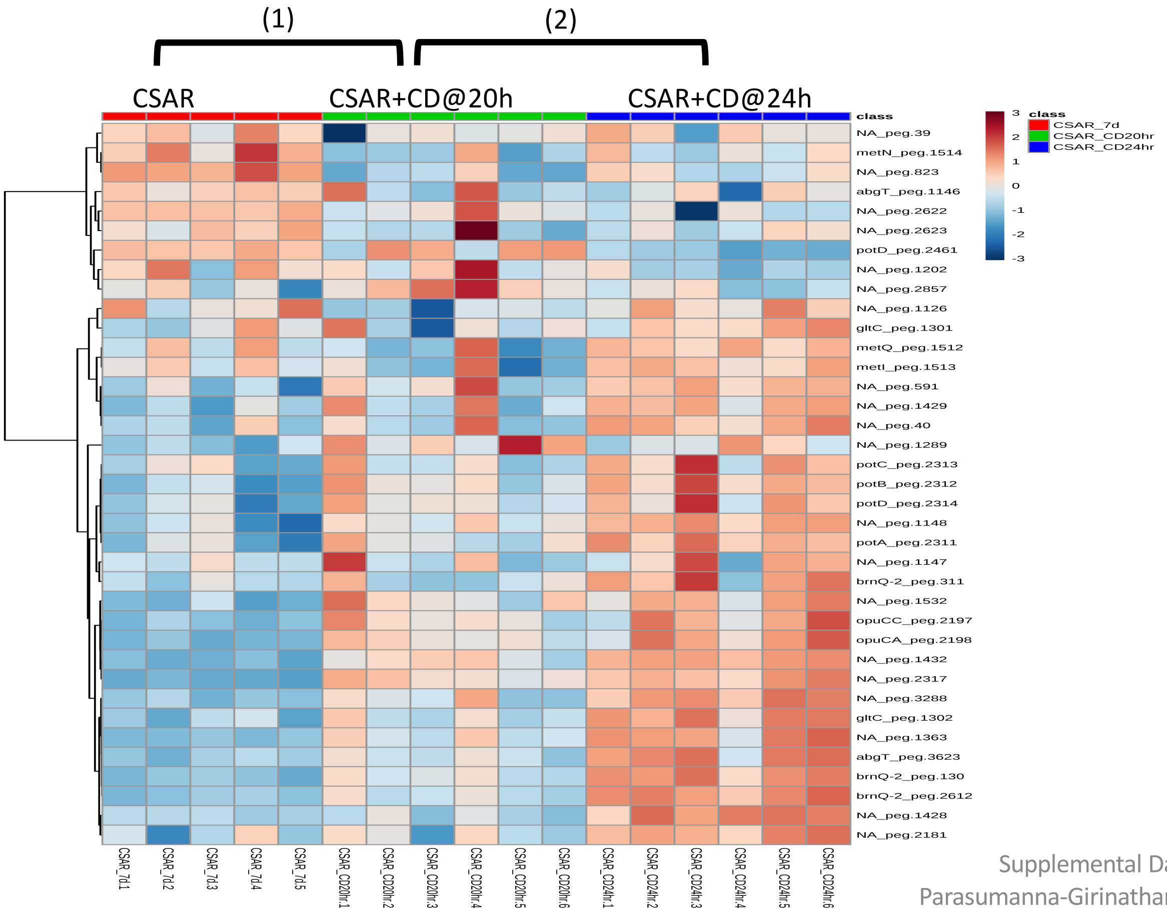


**<u>Significantly Enriched Conditions:</u>**

(1) Enriched in CSAR+*C. difficile* co-colonized vs CSAR mono-colonized mice at 20h (p=9.25E-03).

(2) Enriched in CSAR+*C. difficile* co-colonized at 24hr vs CSAR+*C. difficile* co-colonized at 20hr (p=3.22E-05).

### Carbon Source Transport and Metabolism

#### 6.10 Xanthine, Hypoxanthine Transport and Metabolism

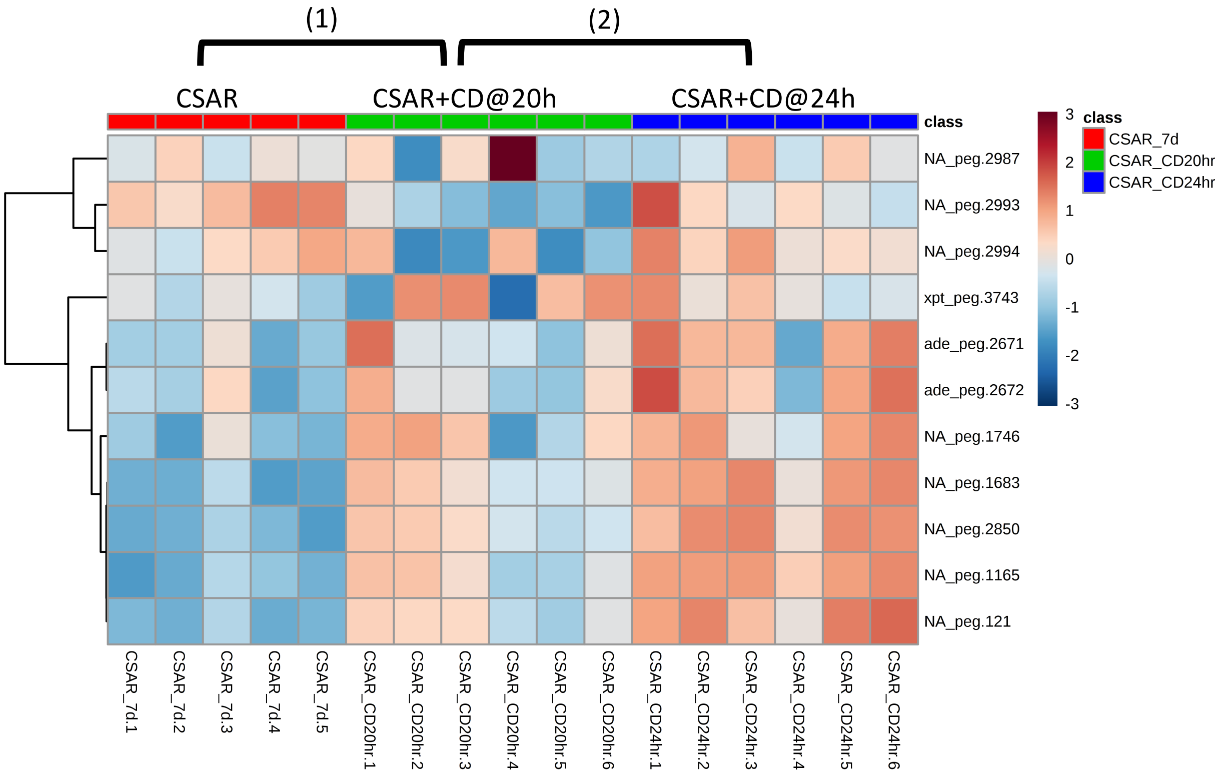


**Significantly Enriched Conditions:**

(1) Enriched in CSAR+*C. difficile* co-colonized vs CSAR mono-colonized mice at 20h (p=3.28E-02).

(2) Enriched in CSAR+*C. difficile* co-colonized at 24hr vs CSAR+*C. difficile* co-colonized at 20hr (p=2.00E-02).

### Core Cellular Machinery

#### 6.11 ATP Synthesis

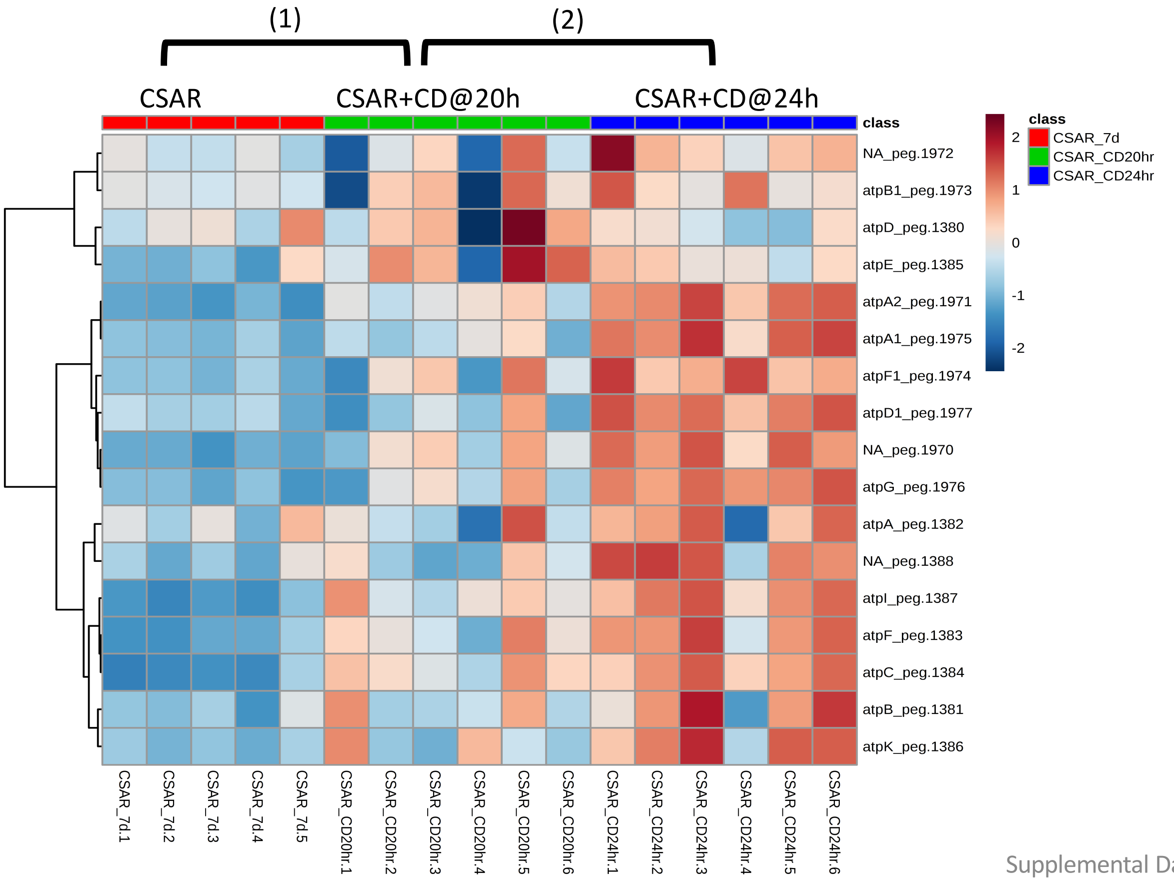


**Significantly Enriched Conditions:**

(1) Enriched in CSAR+*C. difficile* co-colonized vs CSAR mono-colonized mice at 20h (p=3.41E-02).

(2) Enriched in CSAR+*C. difficile* co-colonized at 24hr vs CSAR+*C. difficile* co-colonized at 20hr (p=7.79E-05)

### Core Cellular Machinery

#### 6.12 Electron Transport

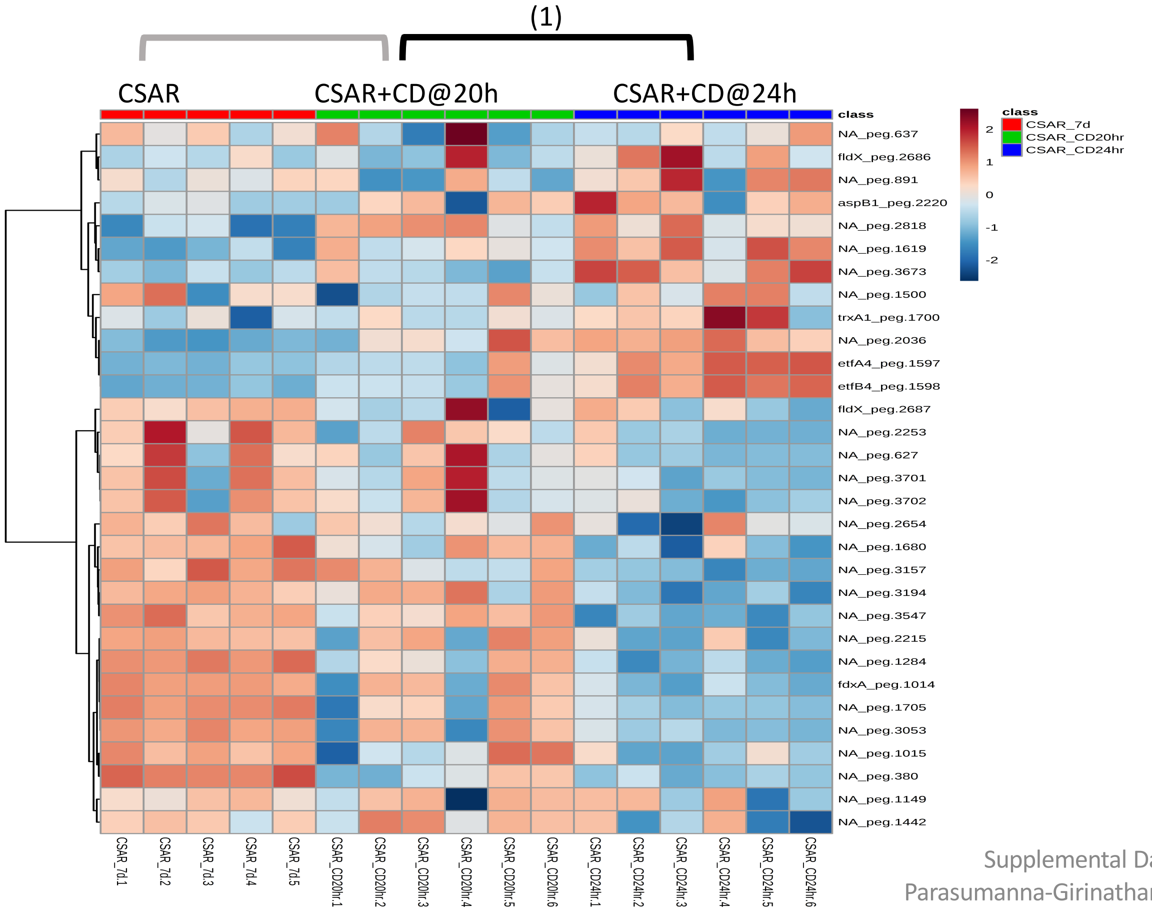


**Significantly Enriched Conditions:**

(1) Enriched in CSAR+*C. difficile* co-colonized at 20hr vs CSAR+*C. difficile* co-colonized at 24hr (p=4.92E-02)

### Core Cellular Machinery

#### 6.13 Hydrogenase Systems

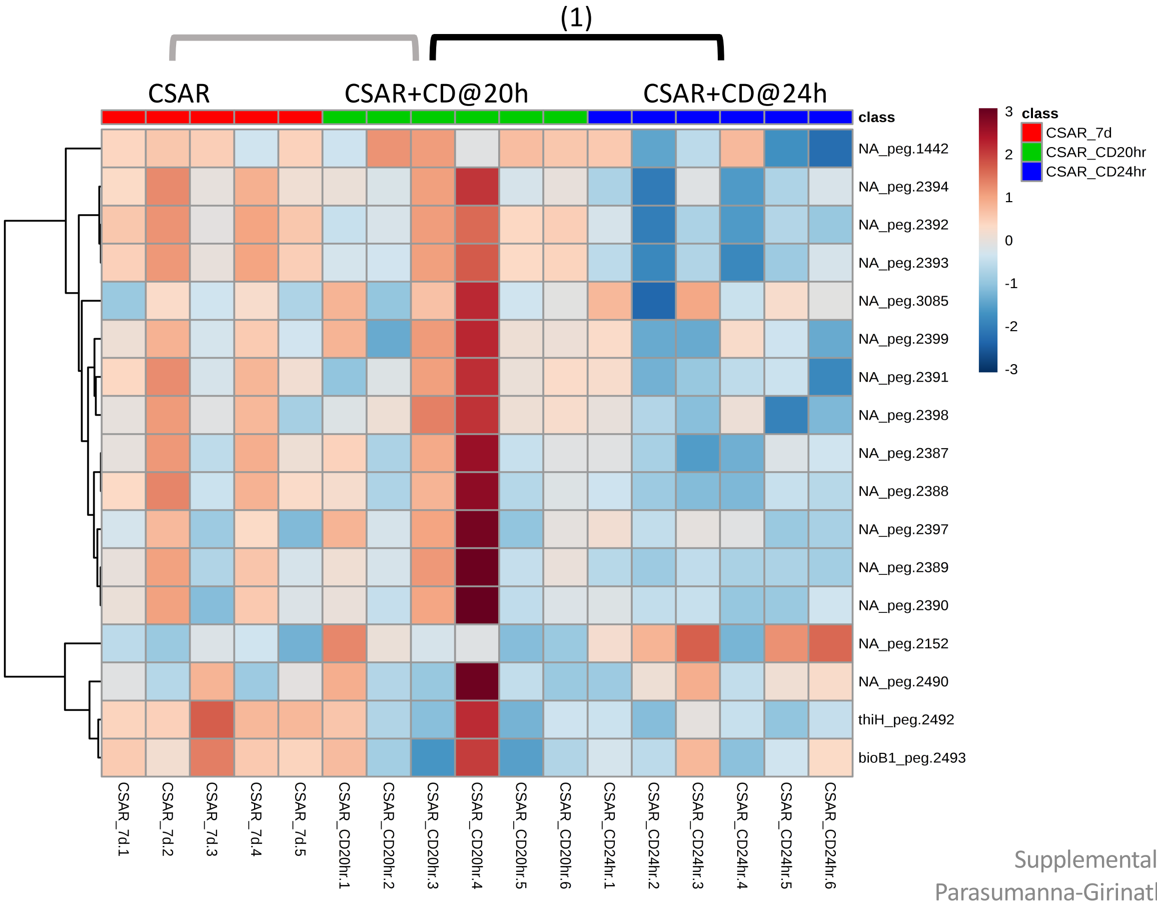


**Significantly Enriched Conditions:**

(1) Enriched in CSAR+*C. difficile* co-colonized at 20hr vs CSAR+*C. difficile* co-colonized at 24hr (p=7.12E-04).

### Core Cellular Machinery

#### 6.14 Ribosomal Structural Components and Modification

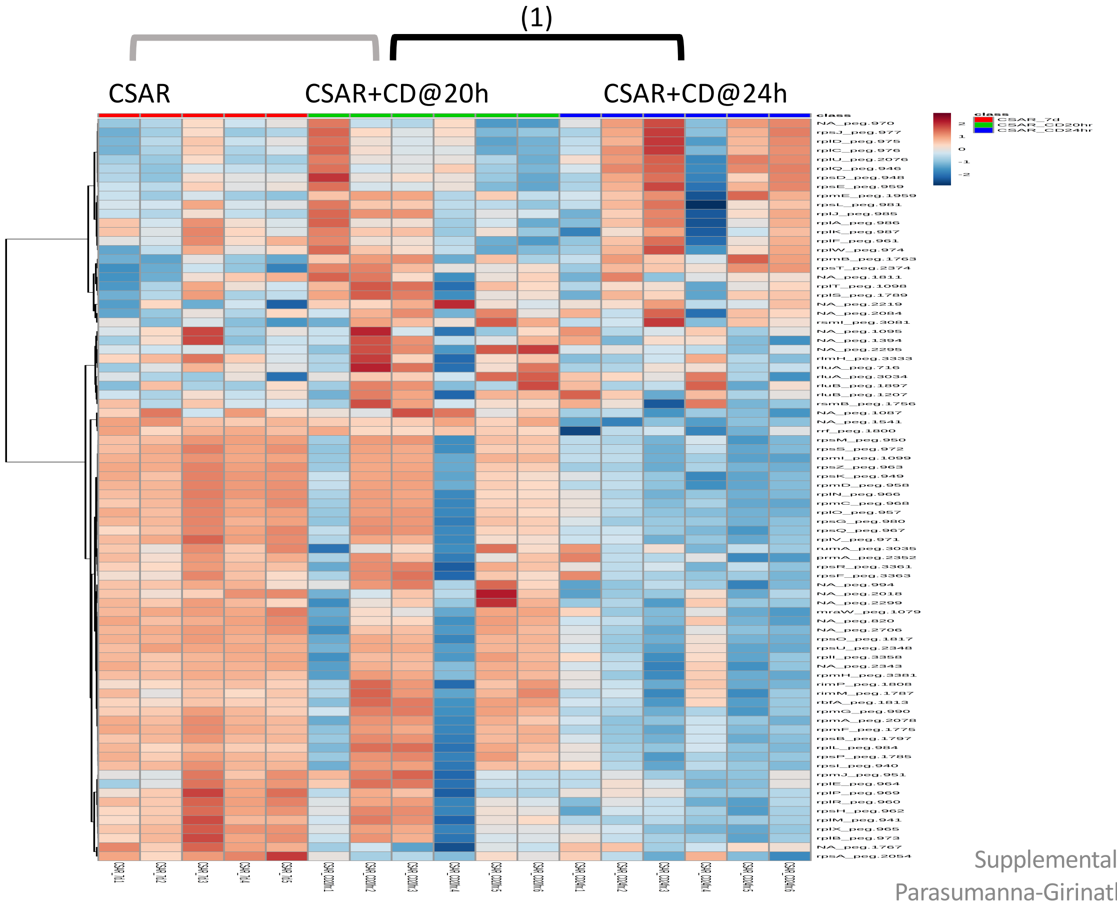


**<u>Significantly Enriched Conditions:</u>**

(1) Enriched in CSAR+*C. difficile* co-colonized at 20hr vs CSAR+*C. difficile* co-colonized at 24hr (p=5.49E-04).

### Cellular Adhesion and Motility

#### 6.15 Cell Surface Adhesive Organelles

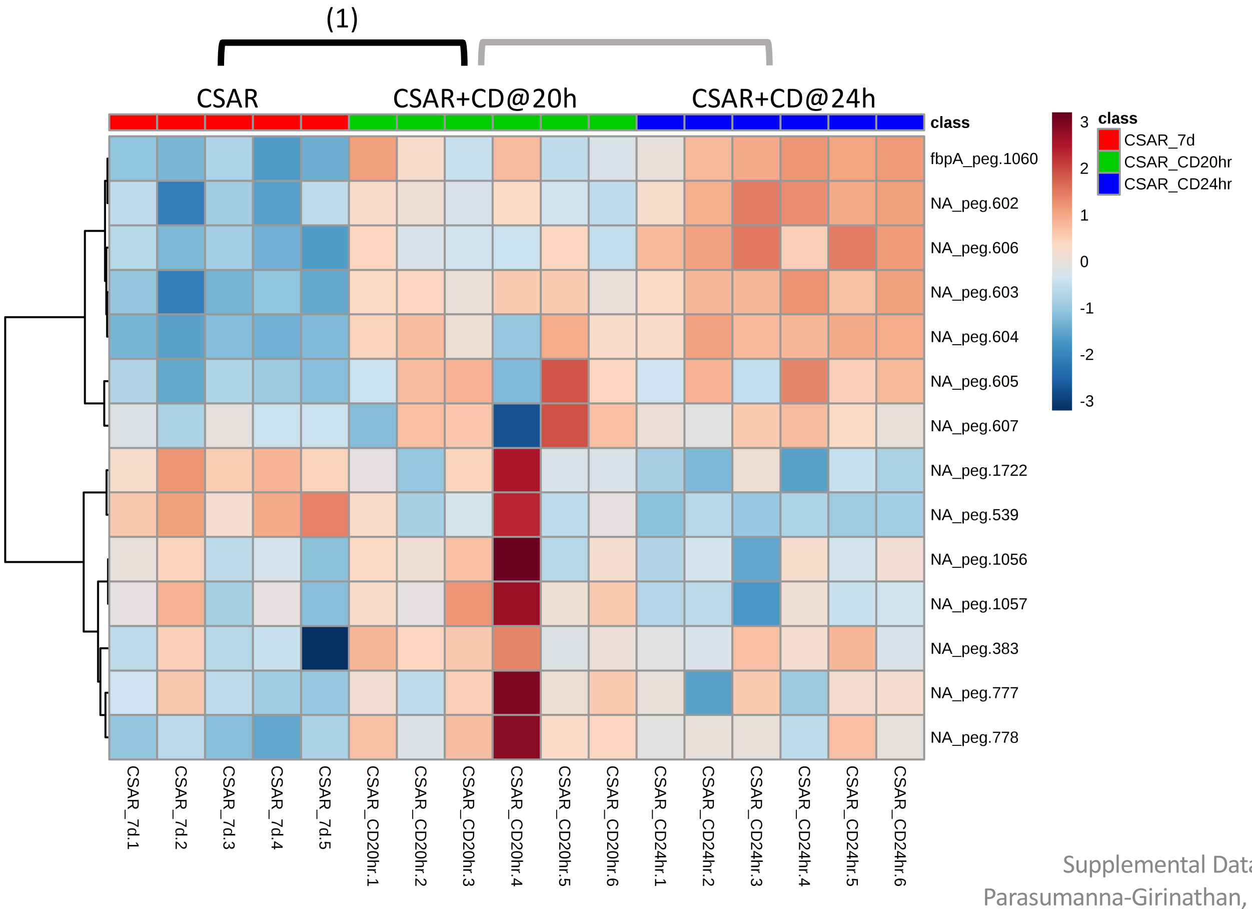


**Significantly Enriched Conditions:**

(1) Enriched in CSAR+*C. difficile* co-colonized mice vs CSAR mono-colonized at 20h (p=6.59E-05).

### Vitamins and Micronutrient Metabolism

#### 6.16 Cobalamin Biosynthesis

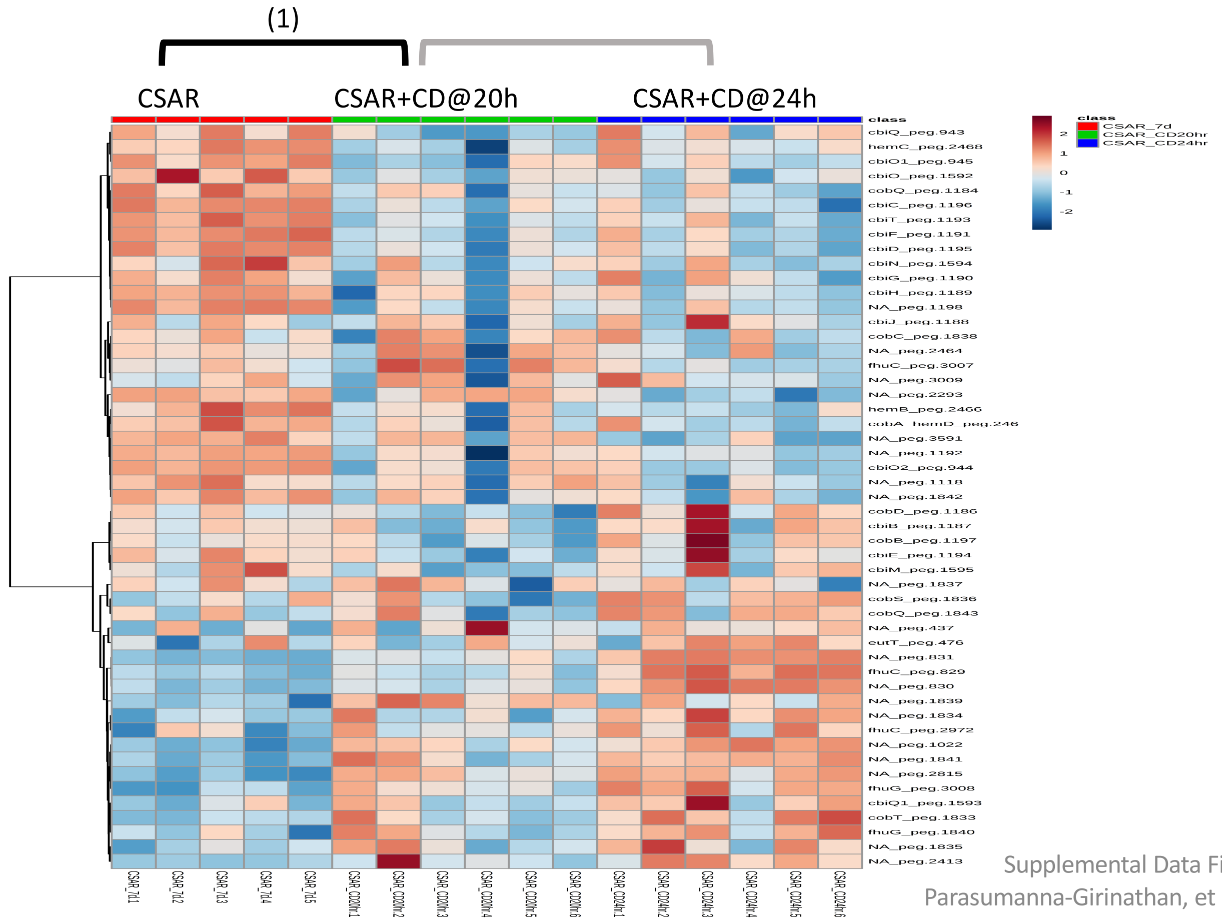


**Significantly Enriched Conditions:**

(1) Enriched in CSAR mono-colonized mice vs CSAR+*C. difficile* co-colonized mice at 20h (p=8.56E-03).

### Vitamins and Micronutrient Metabolism

#### 6.17 Folate Biosynthesis

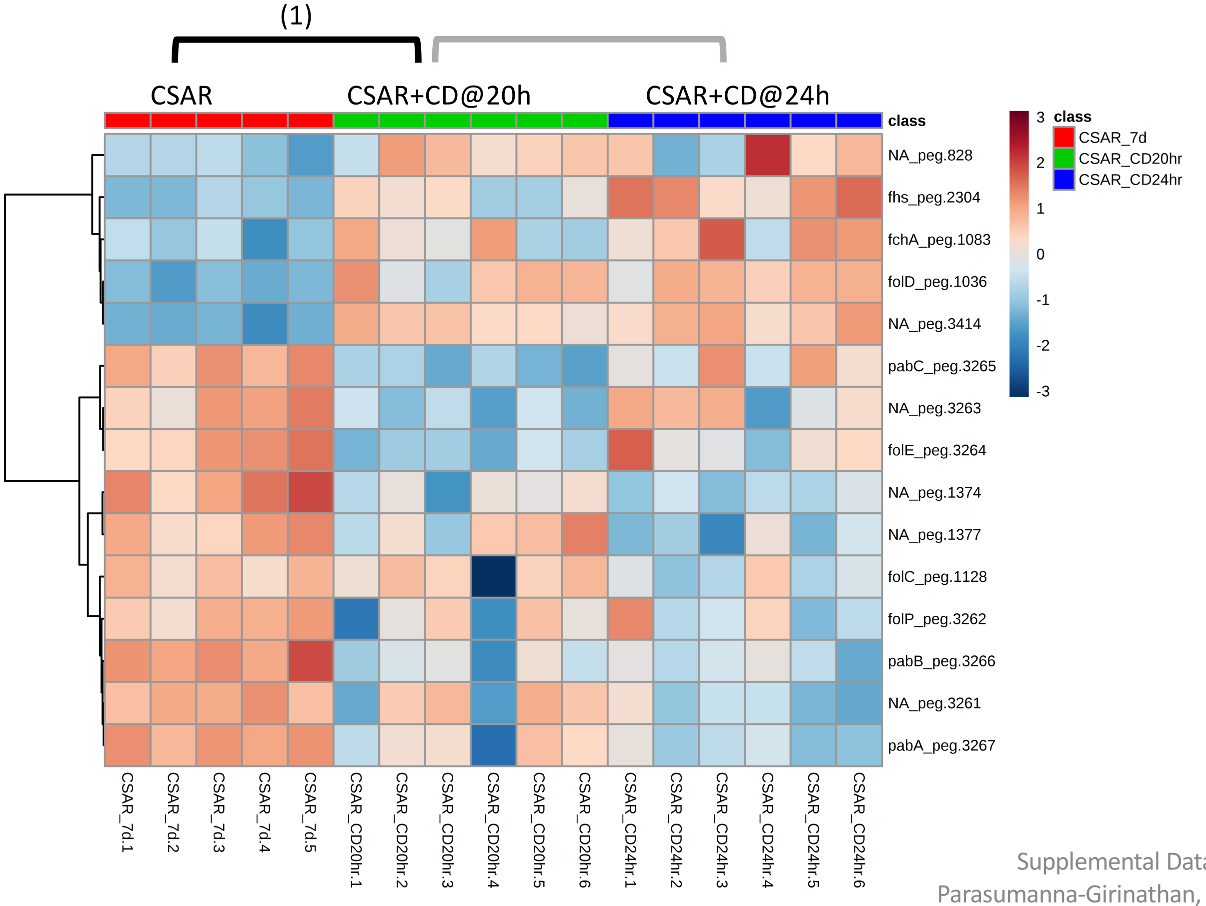


**Significantly Enriched Conditions:**

(1) Enriched in CSAR mono-colonized mice vs CSAR+*C. difficile* co-colonized mice at 20h (p=2.85E-02).

### Vitamins and Micronutrient Metabolism

#### 6.18 Nickel Transport and Metabolism

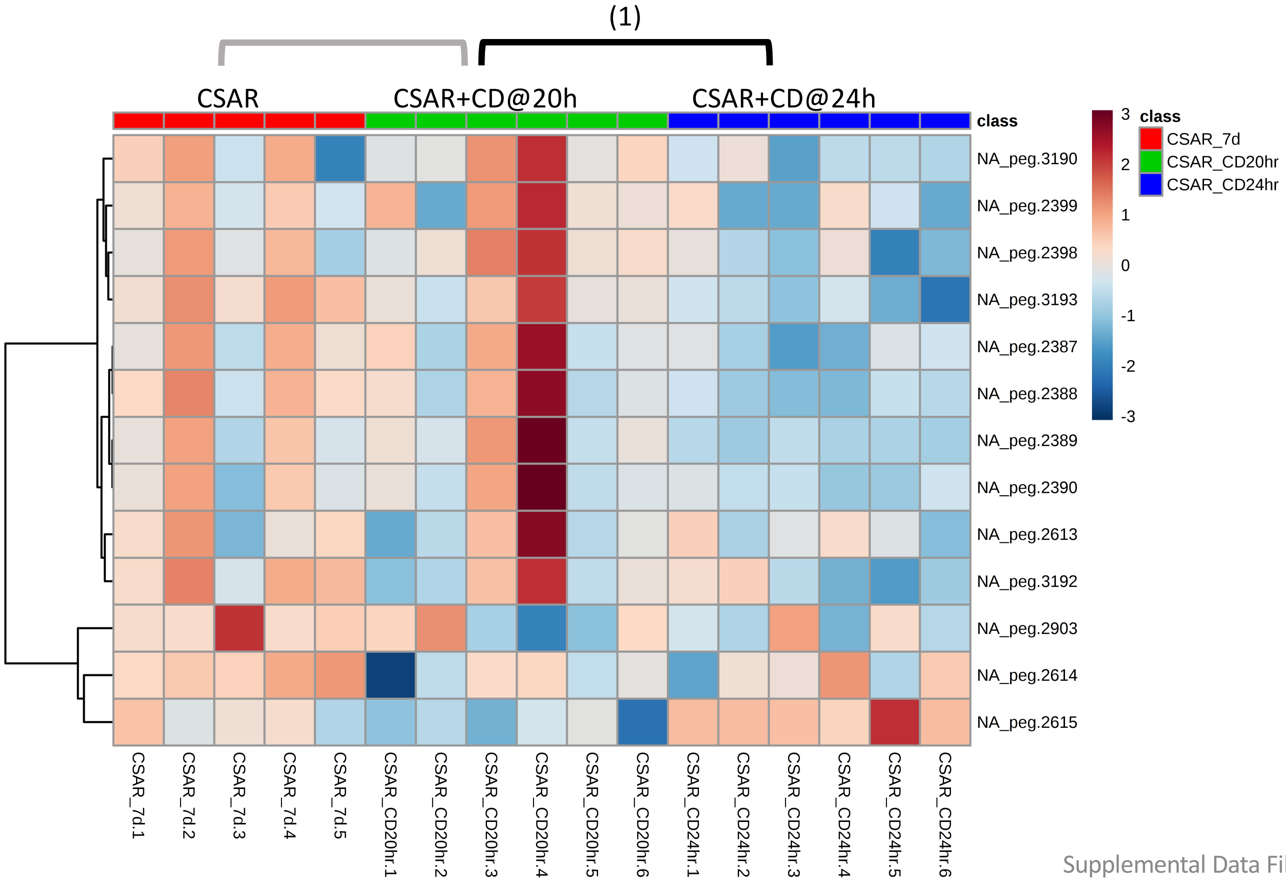


**Significantly Enriched Conditions:**

(1) Enriched in CSAR+*C. difficile* co-colonized mice at 20h vs CSAR+*C. difficile* co-colonized mice at 24h (p=01.25E-03).

### Vitamins and Micronutrient Metabolism

#### 6.19 Phosphorus Transport and Metabolism

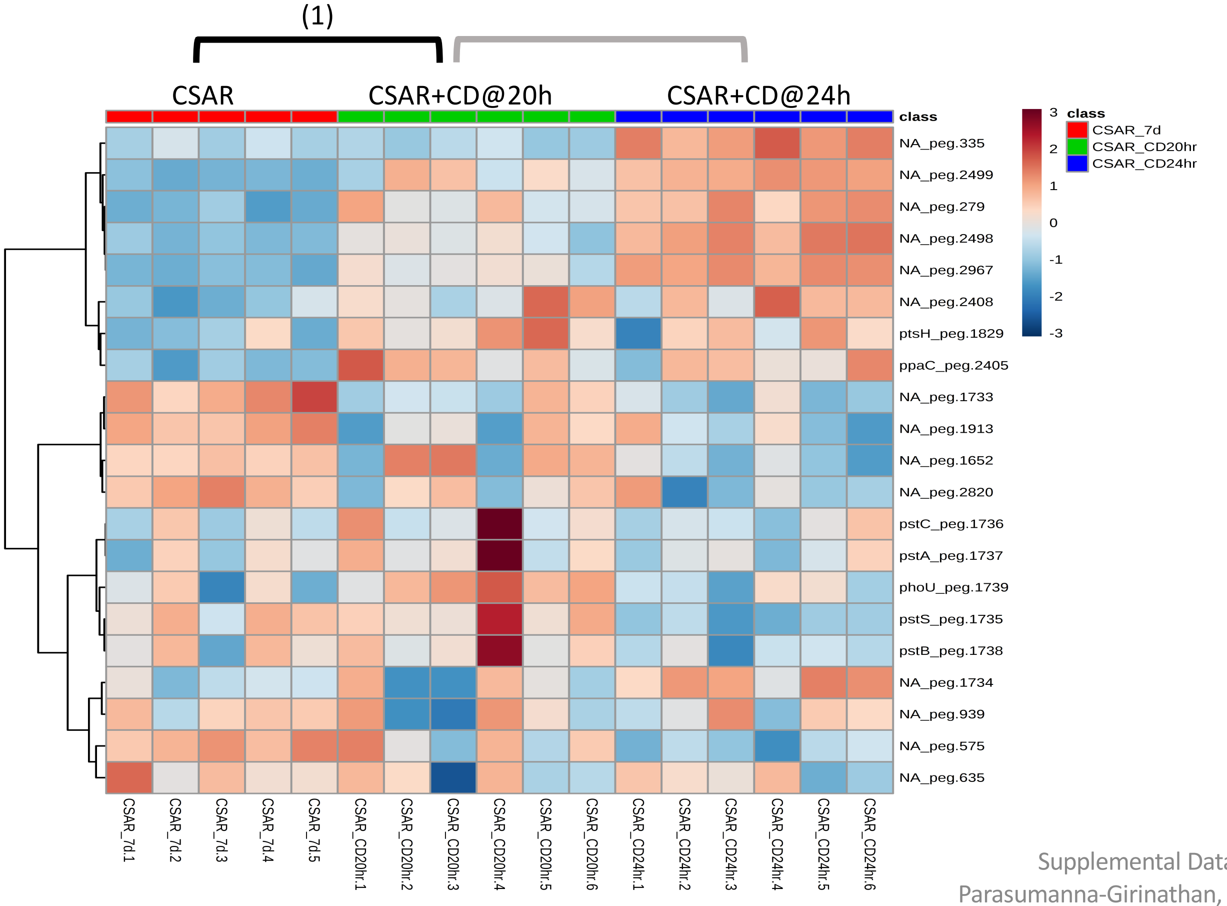


**Significantly Enriched Conditions:**

(1) Enriched in CSAR+*C. difficile* co-colonized vs CSAR mono-colonized mice at 20h (p=9.82E-03).

### Vitamins and Micronutrient Metabolism

#### 6.20 Transport Binding Proteins: Cations

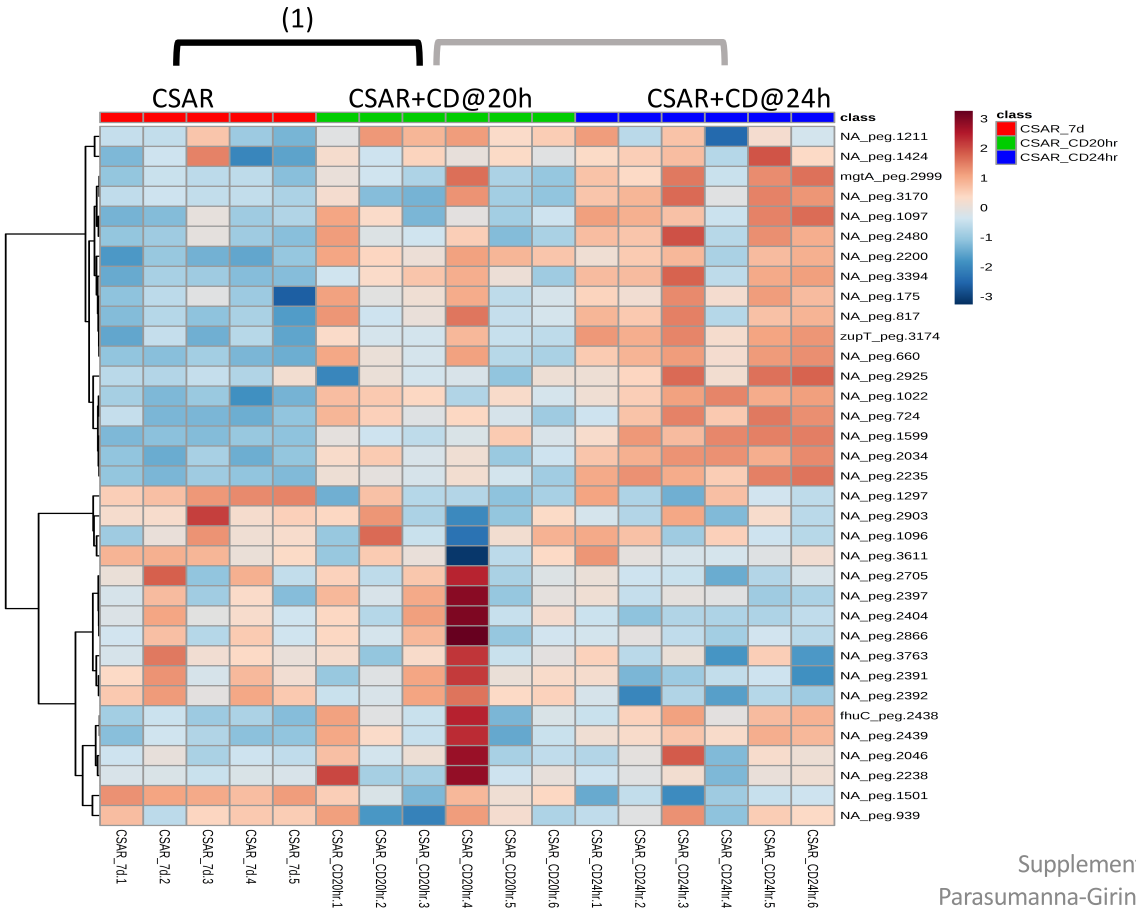


**Significantly Enriched Conditions:**

(1) Enriched in CSAR+*C. difficile* co-colonized vs CSAR mono-colonized mice at 20h (p=1.99E-03).

### Biosynthetic Pathways

#### 6.21 Cysteine and Methionine Biosynthesis

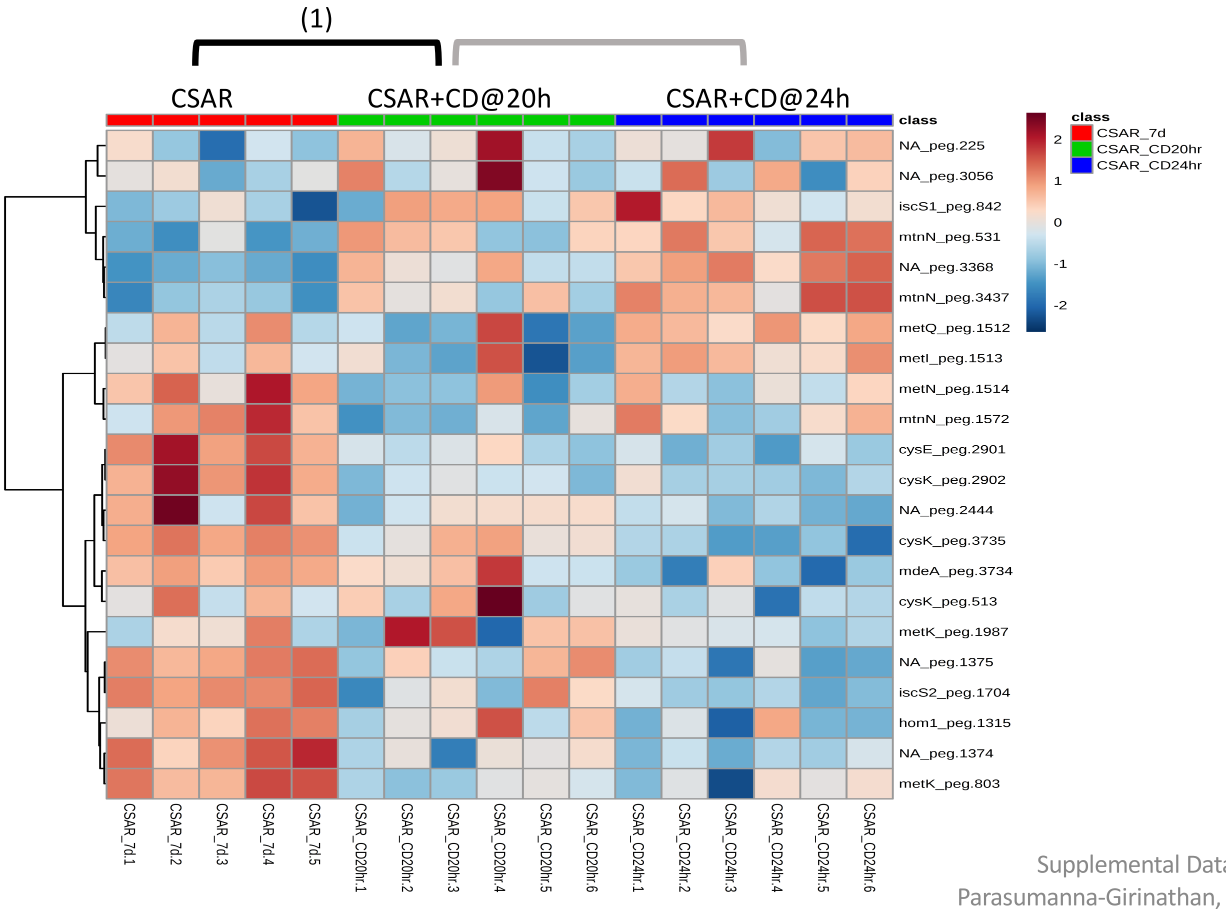


**Significantly Enriched Conditions:**

(1) Enriched in CSAR mono-colonized mice vs CSAR+*C. difficile* co-colonized mice at 20h (p=6.55E-03).

### Biosynthetic Pathways

#### 6.22 Fatty Acid Biosynthesis and Metabolism

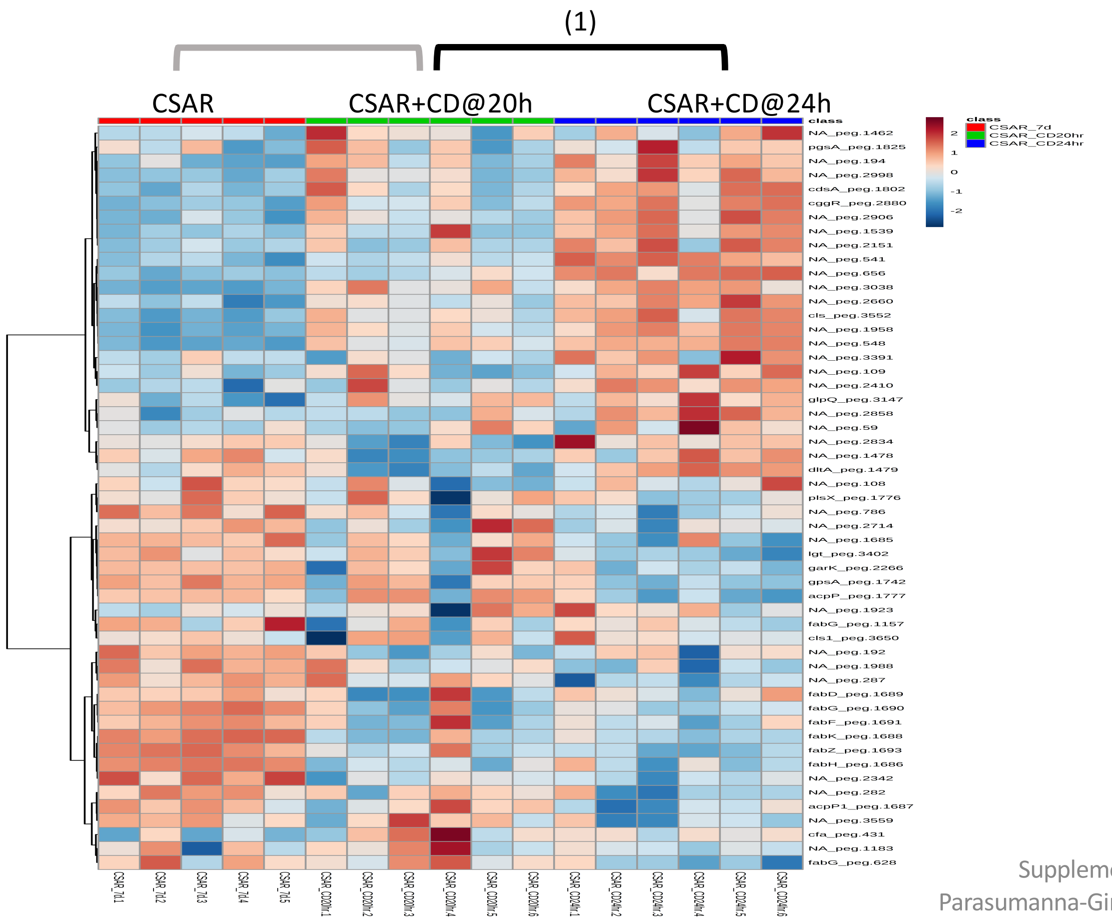


**Significantly Enriched Conditions:**

(1) Enriched in CSAR+*C. difficile* co-colonized mice at 24h vs CSAR+*C. difficile* co-colonized mice at 20h (p=1.02E-02).

### Bacteriophage and Mobile Elements

#### 6.23 Putative Phage Locus 1

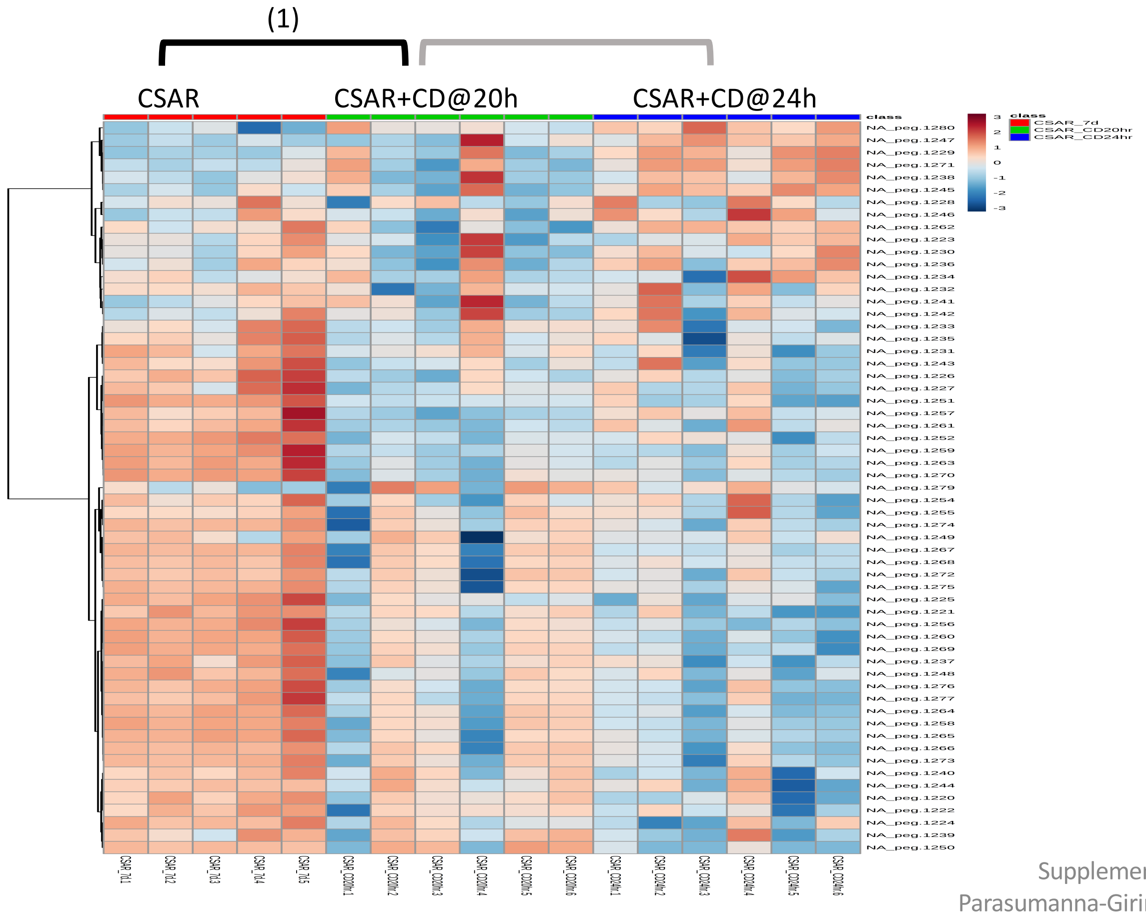


**Significantly Enriched Conditions:**

(1) Enriched in CSAR mono-colonized mice vs CSAR+*C. difficile* co-colonized mice at 20h (p=3.84E-07).

### Bacteriophage and Mobile Elements

#### 6.24 Putative Phage Locus 3

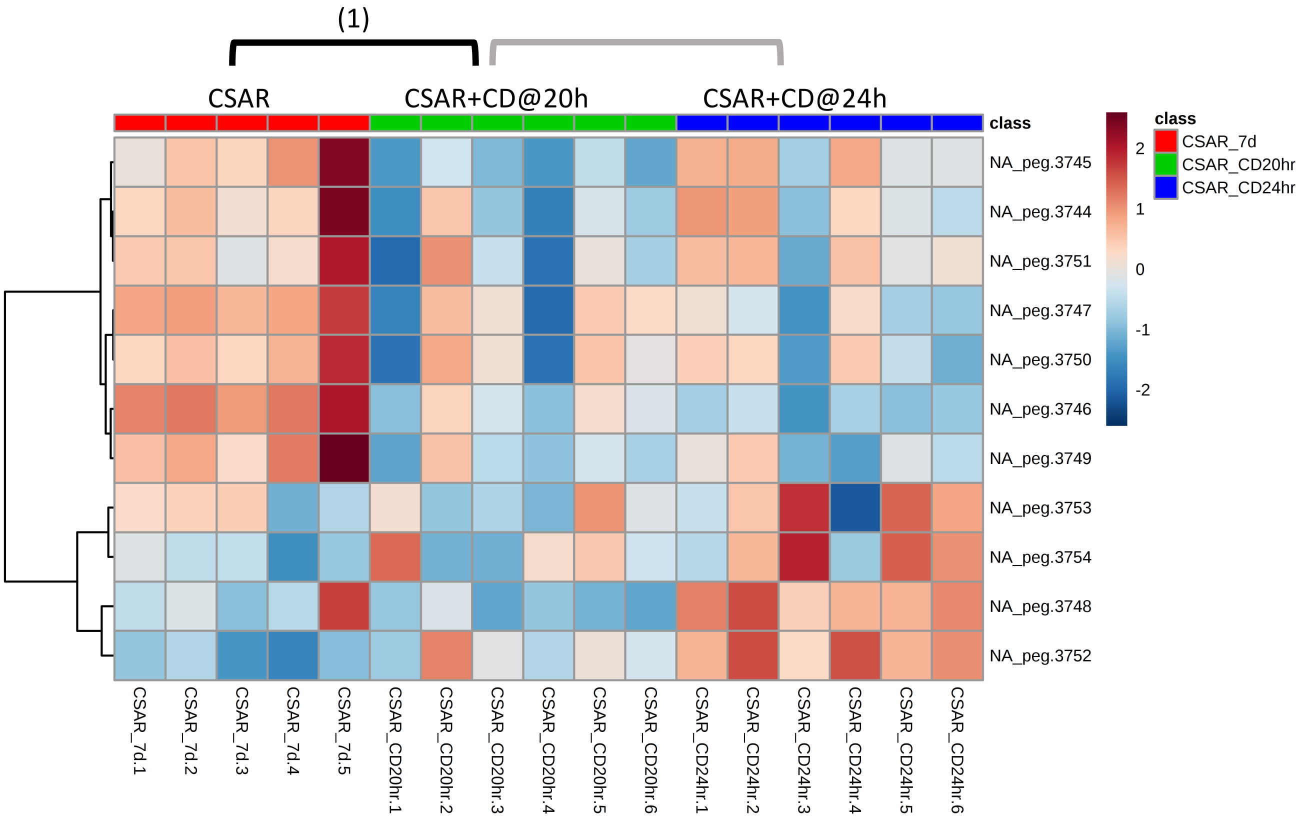


**Significantly Enriched Conditions:**

(1) Enriched in CSAR mono-colonized mice vs CSAR+*C. difficile* co-colonized mice at 20h (p=2.64E-02).

### Bacteriophage and Mobile Elements

#### 6.25 Putative Phage Locus 4

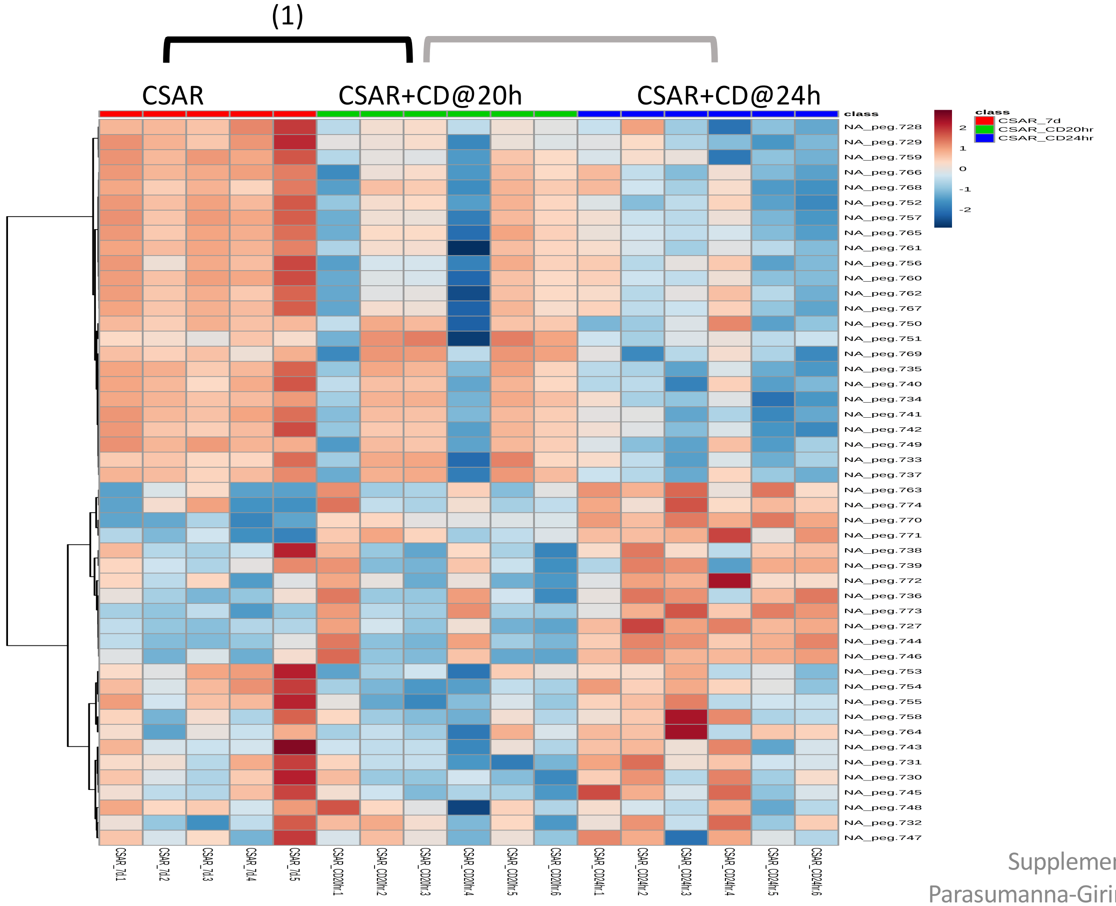


**<u>Significantly Enriched Conditions:</u>**

(1) Enriched in CSAR mono-colonized mice vs CSAR+*C. difficile* co-colonized mice at 20h (p=4.28E-02).

### Stress Responses

#### 6.26 Sporulation

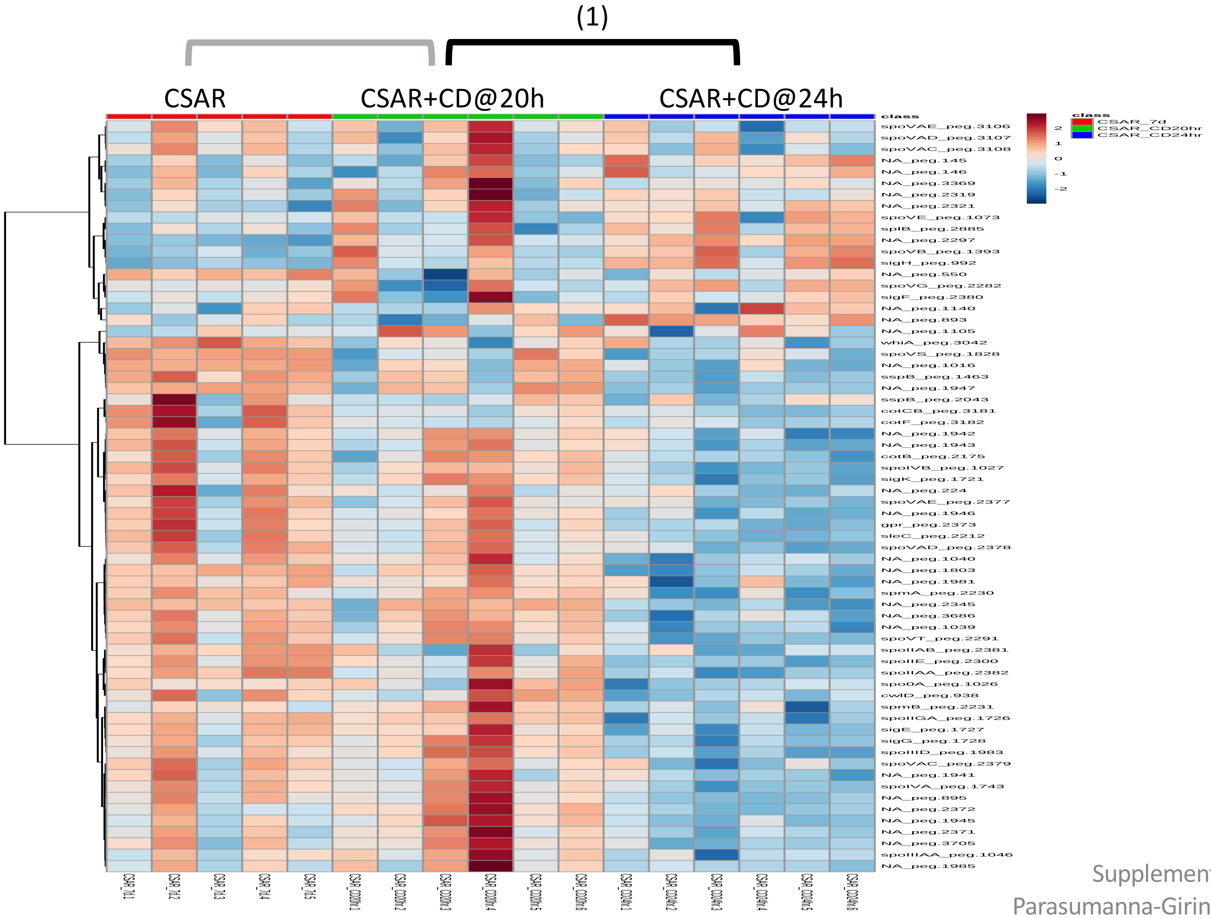


**Significantly Enriched Conditions:**

(1) Enriched in CSAR+*C. difficile* co-colonized mice at 20h vs CSAR+*C. difficile* co-colonized mice at 24h (p=1.02E-02).
