## Supplemental Data File 7 for "*In vivo* commensal control of *Clostridioides difficile* virulence"

- Version 1.0; 12/23/2019

### Supplemental Data File 7: Transcriptome Analyses: *C. bifermentans* Enriched Pathways

**Page 0:** Introduction, this page
**Page 1**: Sample Key


**Carbon Source Transport and Metabolism**
Page 2: 7.1 Butanoate Metabolism
3: 7.2 Ethanolamine Utilization
4: 7.3 Polyamine metabolism
5: 7.4 Ribose Transport and Metabolism
6: 7.5 Stickland Fermentations
7: 7.6 Transport Binding Proteins: Amino Acids and Amines

**Core Cellular Machinery**
8: 7.7 Amino-acyl tRNA synthesis and modification
9: 7.8 ATP Synthesis
10: 7.9 Cell division
11: 7.10 Chaperone Systems
12: 7.11 DNA modification
13: 7.12 Electron Transport
14: 7.13 Protein translation and modification
15: 7.14 Ribosome Synthesis and Modification

**Cellular Adhesion and Motility**
Page 16: 7.15 Flagellar Motility

**Cell Wall Components**
17: 7.16 Gram+ Peptidoglycan and Teichoic Acid

**Vitamin and Micronutrient Metabolism**
18: 7.17 Cobalamin Biosynthesis
19: 7.18 Folic Acid Biosynthesis
20: 7.19 Flavodoxin and Riboflavin Biosynthesis
21: 7.20 Iron Transport and Storage
22: 7.21 Sulfur-Compound Transport and Metabolism
23: 7.22 Transport Binding Proteins: Cations

**Other Transport Systems**
Page 24: 7.23 ABC Transport Systems
25: 7.24 Multi-drug Transport Systems

**Bacteriophage and Mobile Elements**
26: 7.25 Putative Phage

**Stress Responses**
27: 7.26 Cellular Stress Responses

### SDF7: Sample Key

7.9: ATP Synthesis

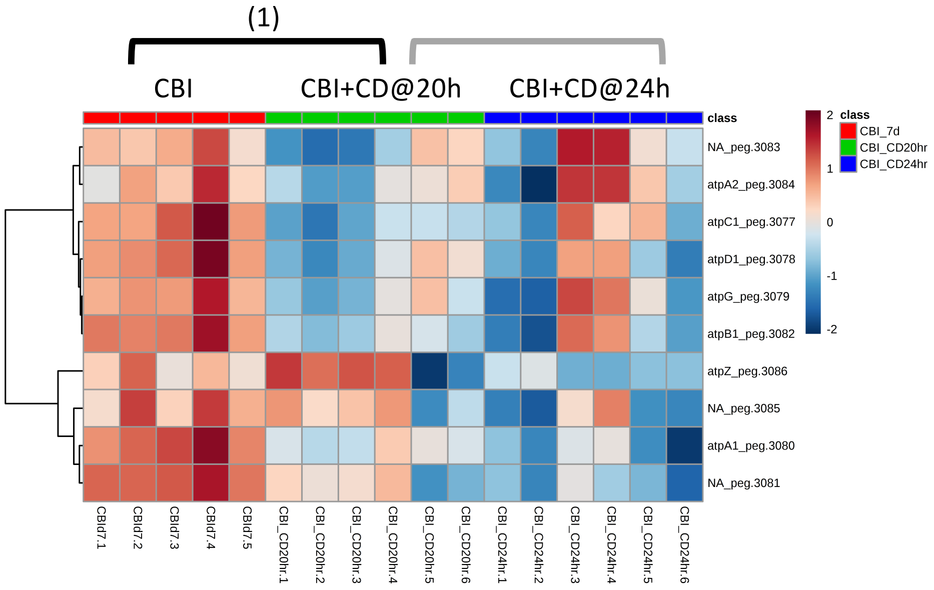


**Guide:**

- Group showing the enriched category is at the top ("7.9 ATP Synthesis")
- Brackets indicate the 2 comparisons evaluated among groups for significant enrichment of genes in the pathway
- Brackets in black show significant comparisons
- Significant comparisons and p values are shown in the text box on the right-hand side

**Significantly Enriched Conditions:**

(1) Enriched in CBI-mono-colonized mice vs CBI+*C. difficile* co-colonized mice at 20h (p=0.0421123).

### Carbon Source Transport and Metabolism

#### 7.1 Butanoate Metabolism

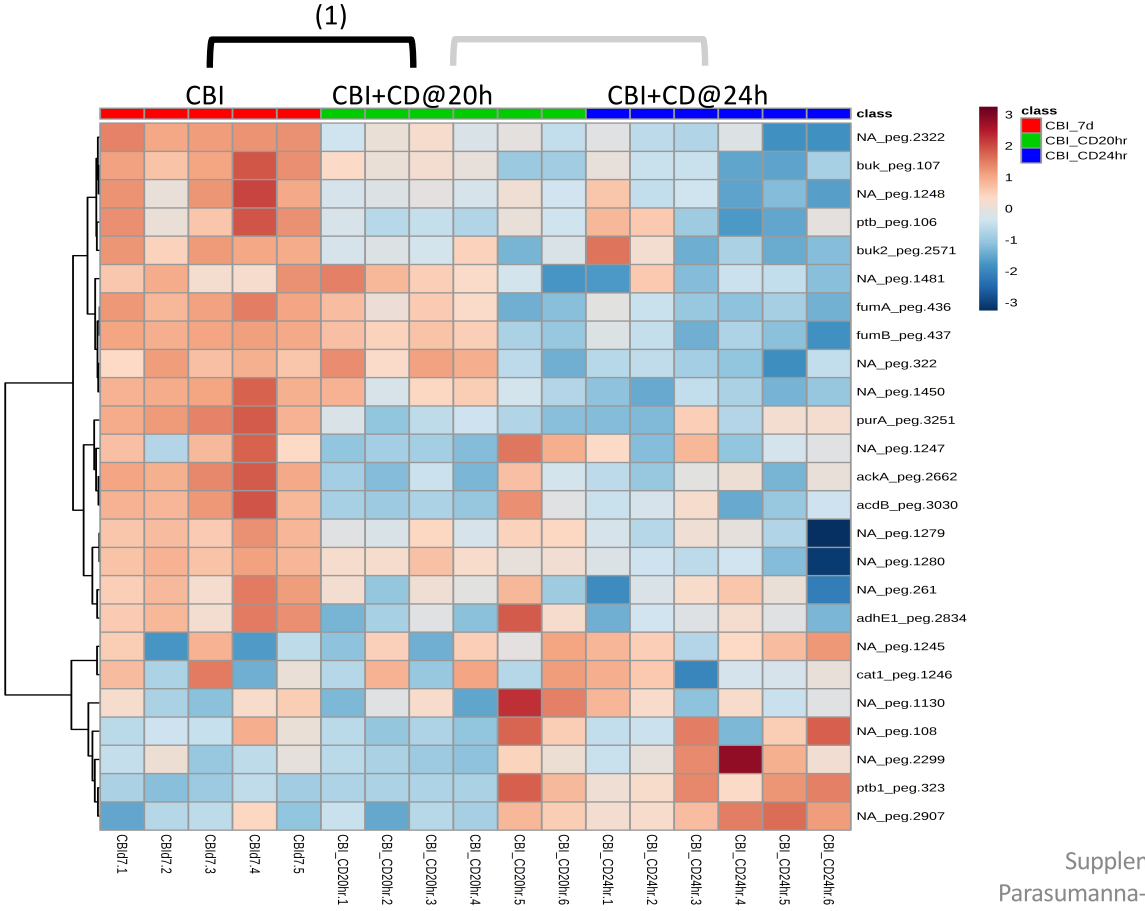


**Significantly Enriched Conditions:**

(1) Enriched in CBI-mono-colonized mice vs CBI+*C. difficile* co-colonized mice at 20h (p=2.04E-02).

### Carbon Source Transport and Metabolism

#### 7.3 Ethanolamine Utilization

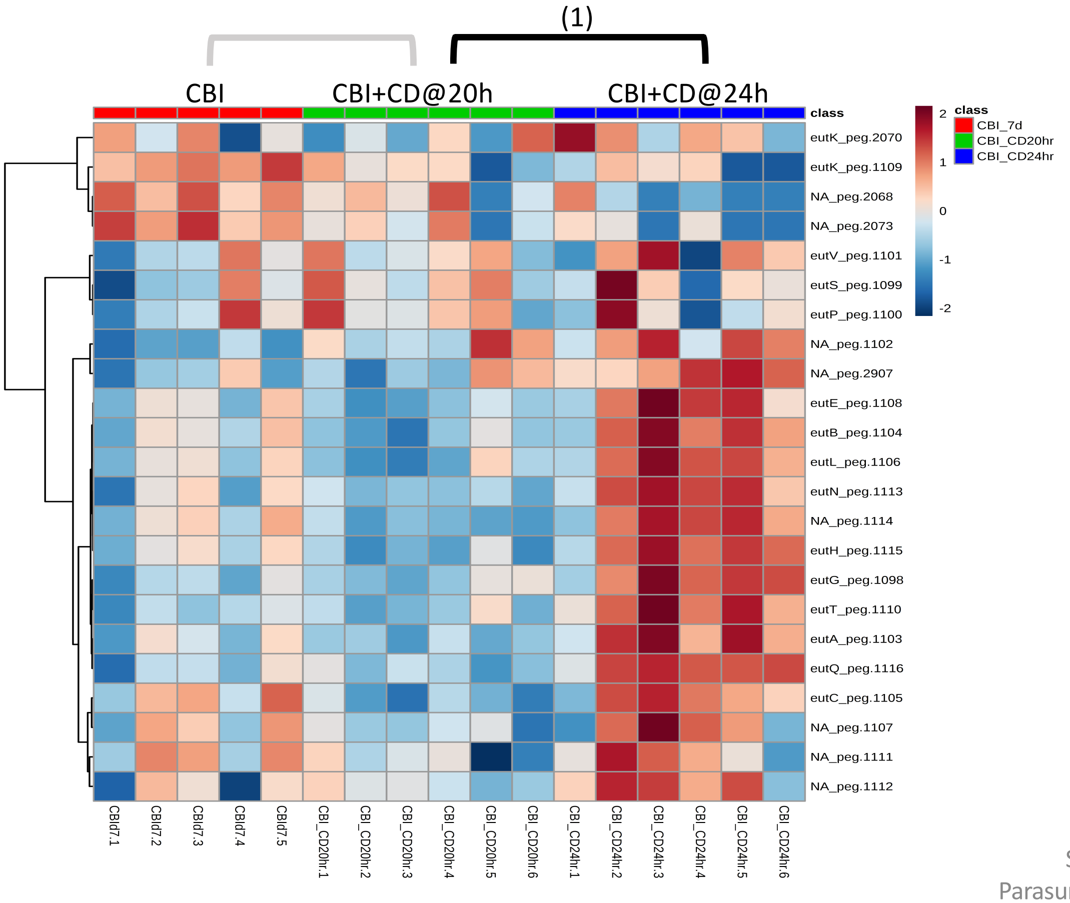


**Significantly Enriched Conditions:**

(1) Enriched in CBI+*C. difficile* co-colonized at 24h *vs* CBI +*C. difficile* co-colonized at 20h (p=1.16E-06).

### Carbon Source Transport and Metabolism

#### 7.3 Polyamine Metabolism

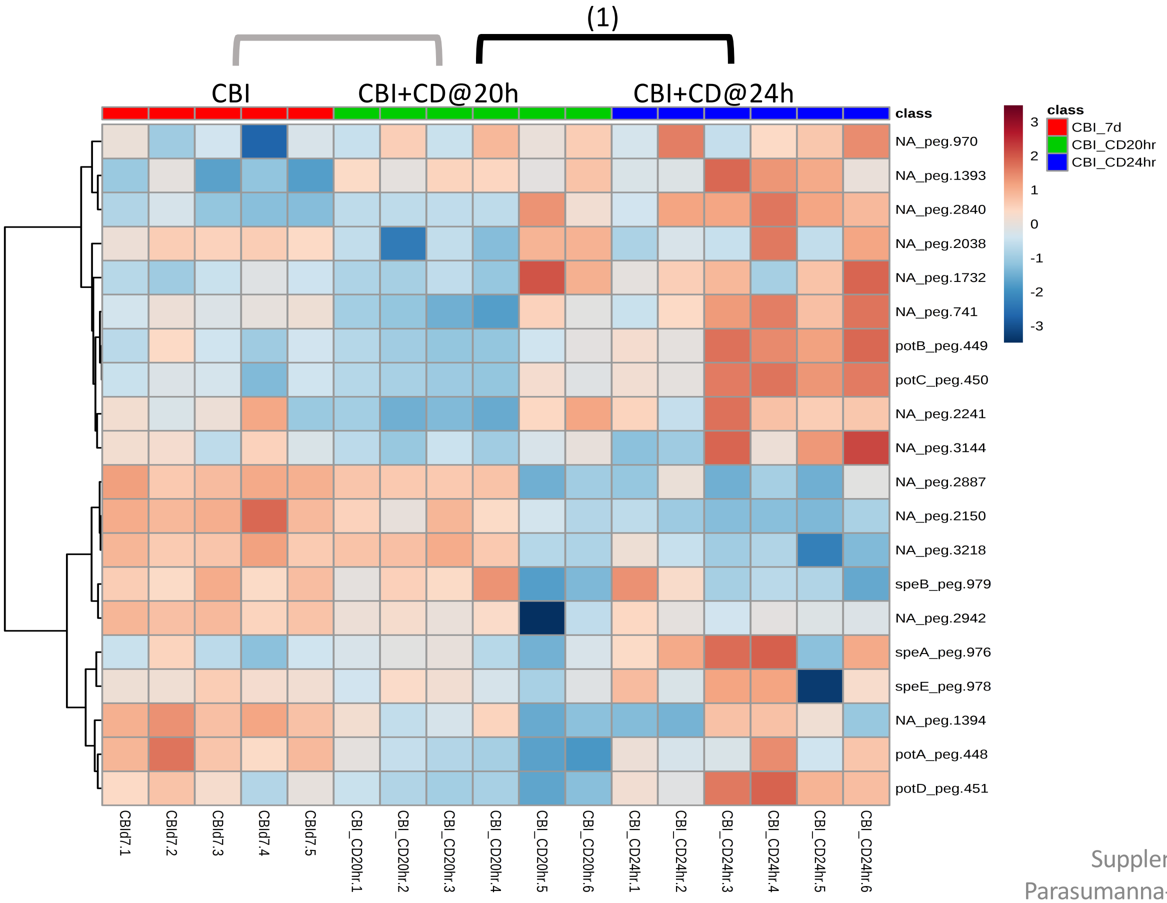


**<u>Significantly Enriched Conditions</u>:**

(1) Enriched in CBI+*C. difficile* co-colonized at 24h *vs* CBI +*C. difficile* co-colonized at 20h (p=7.89E-03).

### Carbon Source Transport and Metabolism

#### 7.4 Ribose Metabolism

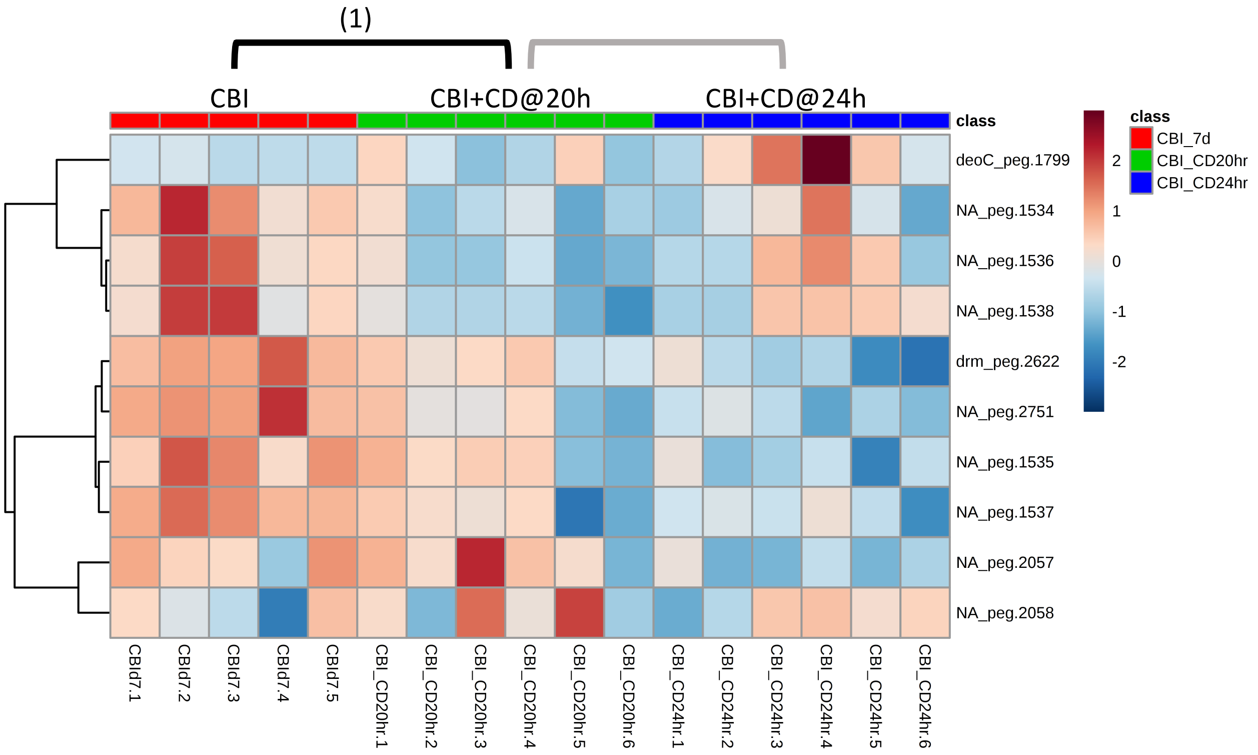


**Significantly Enriched Conditions:**

(1) Enriched in CBI-mono-colonized mice vs CBI+*C. difficile* co-colonized mice at 20h (p=4.63E-02).

### Carbon Source Transport and Metabolism

#### 7.5 Stickland Fermentations

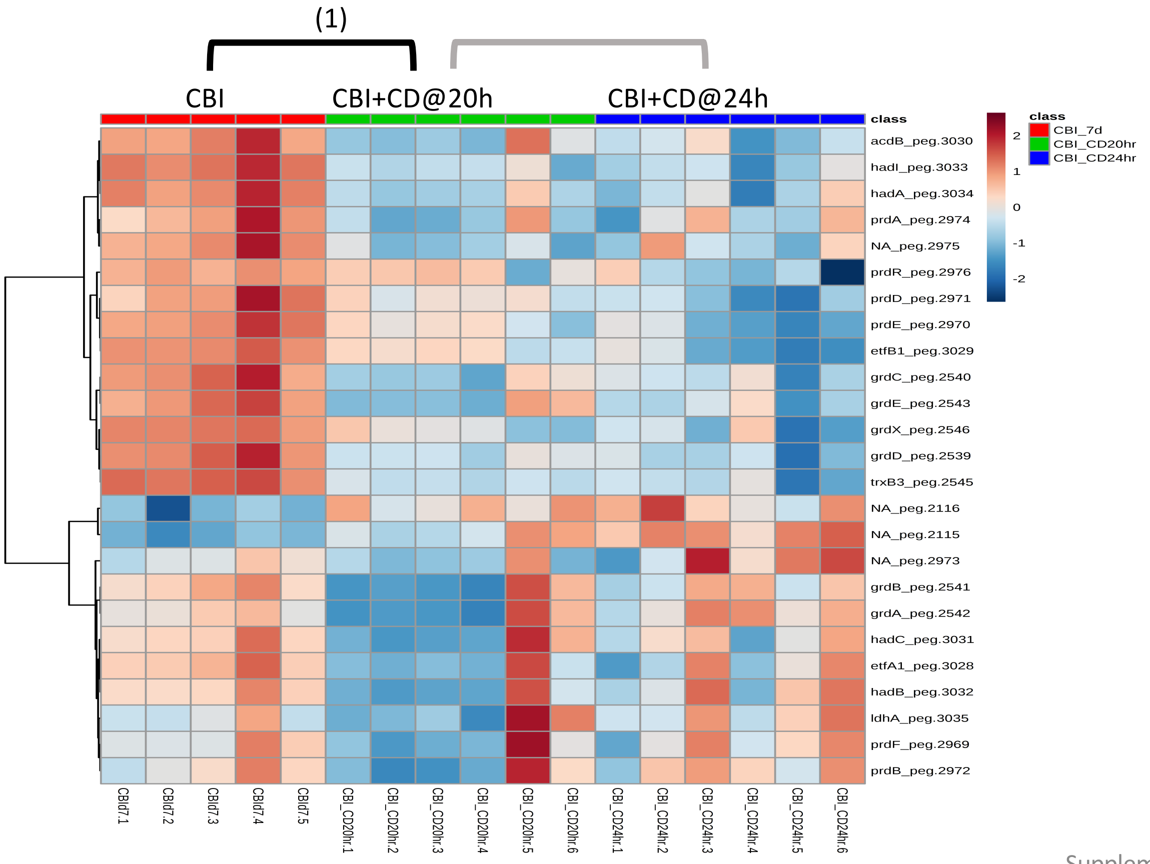


**Significantly Enriched Conditions:**

(1) Enriched in CBI-mono-colonized mice vs CBI+*C. difficile* co-colonized mice at 20h (p=1.40E-03).

### Carbon Source Transport and Metabolism

#### 7.6 Transport Binding Proteins: Amino Acids and Amines

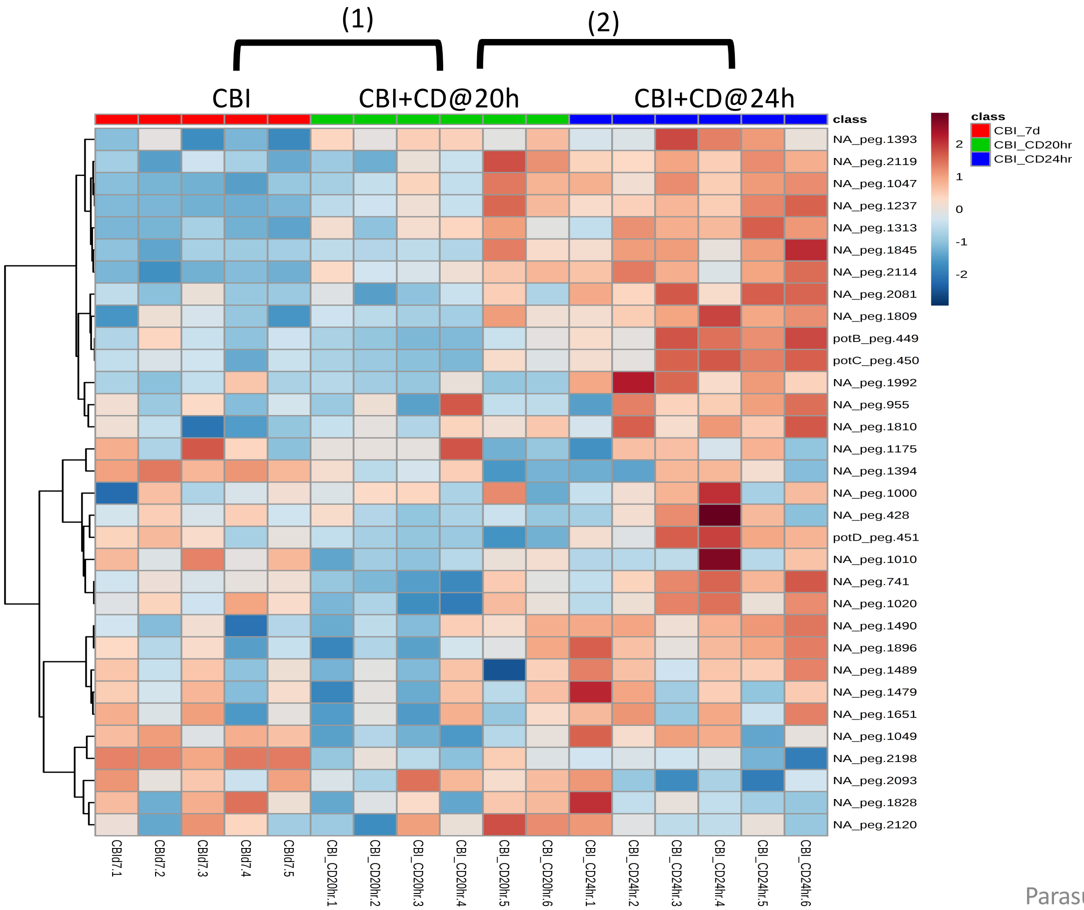


**Significantly Enriched Conditions:**

(1) Enriched in CBI+*C. difficile* co-colonized *vs* CBI mono-colonized mice at 20h (p=1.17E-02).

(2) Enriched in CBI+*C. difficile* co-colonized at 24h *vs* CBI +*C. difficile* co-colonized at 20h (p=3.39E-05).

### Core Cellular Machinery

#### 7.7 Amino-acyl tRNA Synthesis and Modification

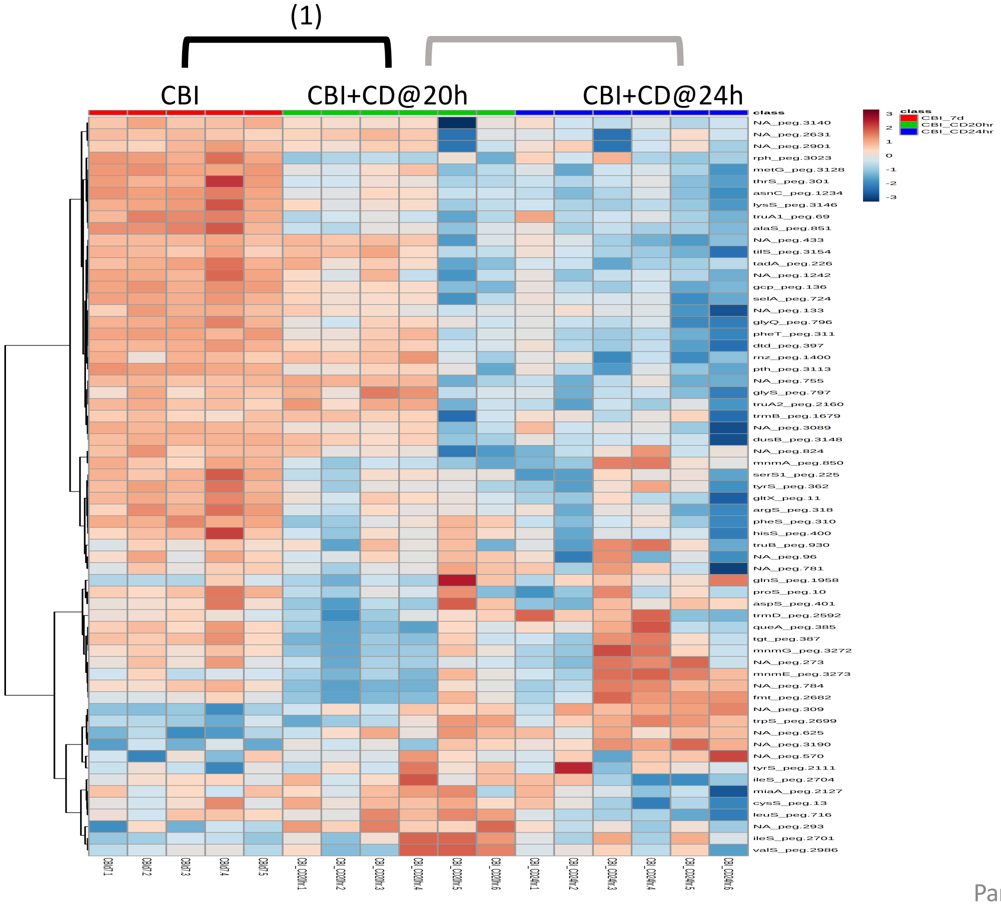


**Significantly Enriched Conditions:**

(1) Enriched in CBI-mono-colonized mice vs CBI+*C. difficile* co-colonized mice at 20h (p=3.60E-03).

### Core Cellular Machinery

#### 7.8 ATP Synthesis

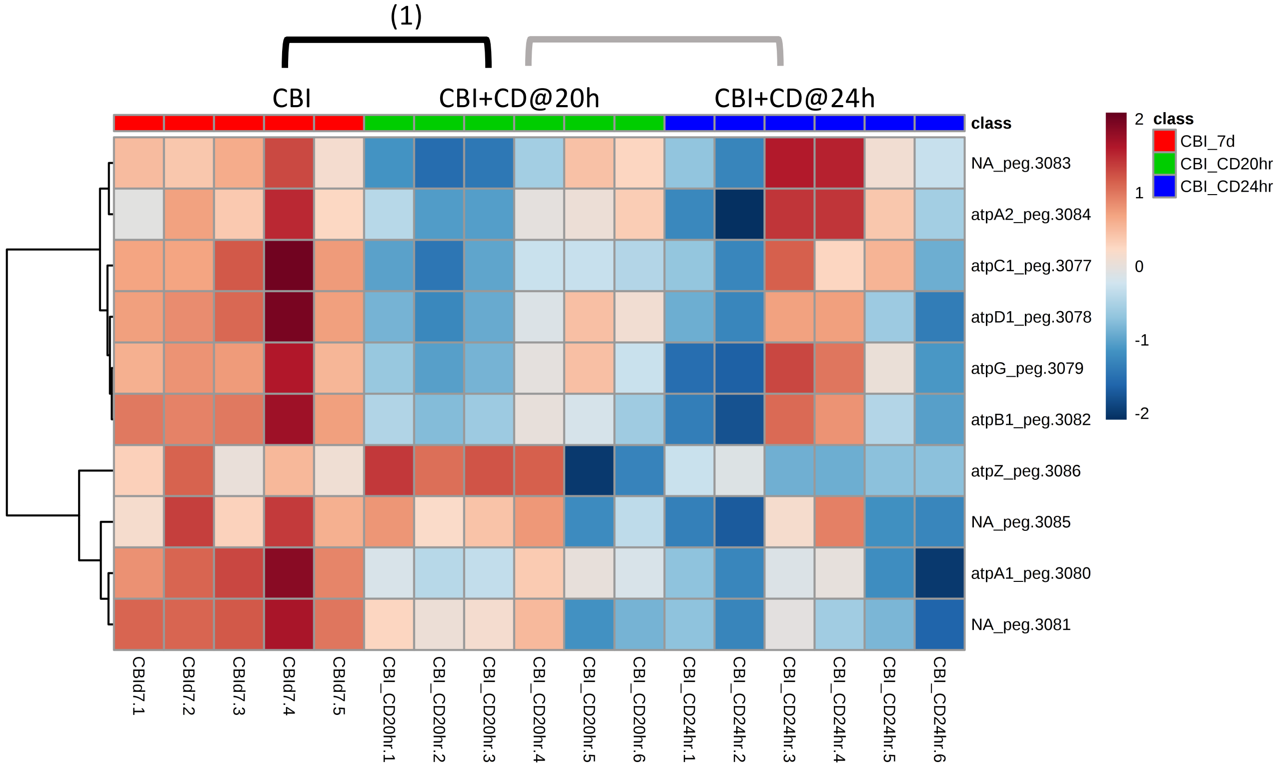


**Significantly Enriched Conditions:**

(1) Enriched in CBI-mono-colonized mice vs CBI+*C. difficile* co-colonized mice at 20h (p=4.21E-02).

### Core Cellular Machinery

#### 7.9 Cell division

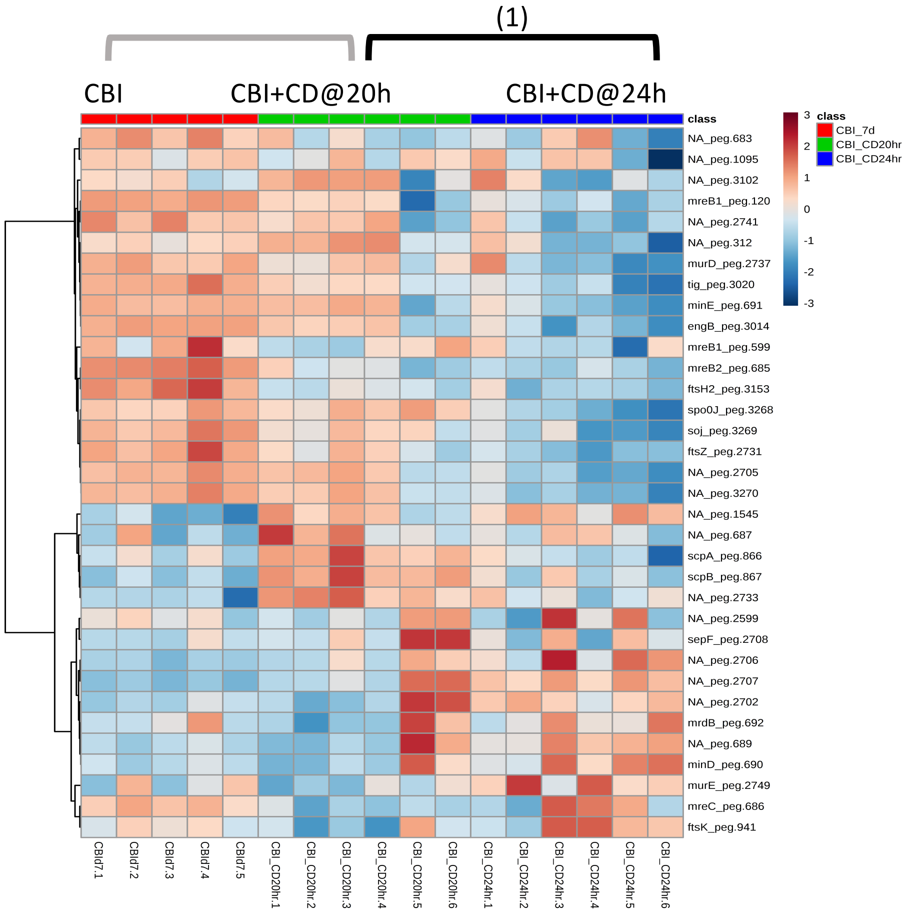


**Significantly Enriched Conditions:**

(1) Enriched in CBI+*C. difficile* co-colonized at 20h *vs* CBI +*C. difficile* co-colonized at 24h (p=1.77E-02).

#### 7.10 Chaperone Systems

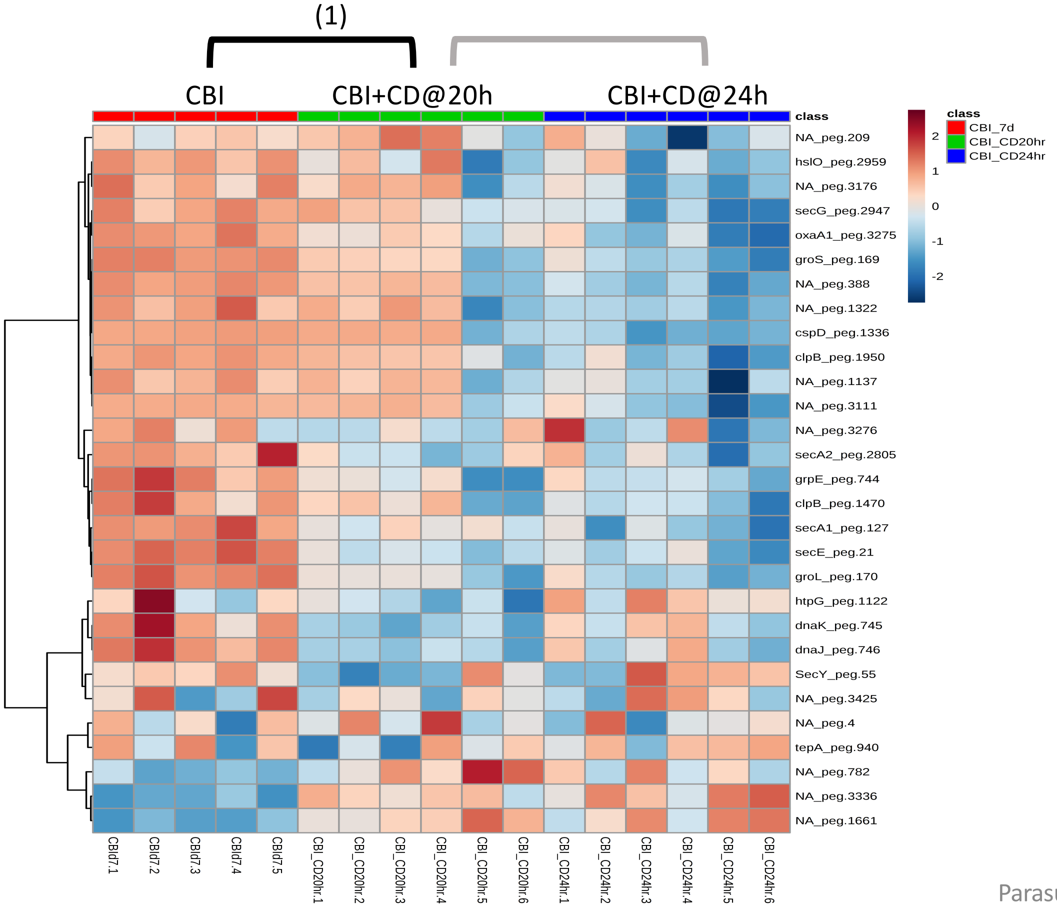


**<u>Significantly Enriched Conditions:</u>**

(1) Enriched in CBI-mono-colonized mice vs CBI+*C. difficile* co-colonized mice at 20h (p=9.54E-03).

### Core Cellular Machinery

#### 7.11 DNA Modification

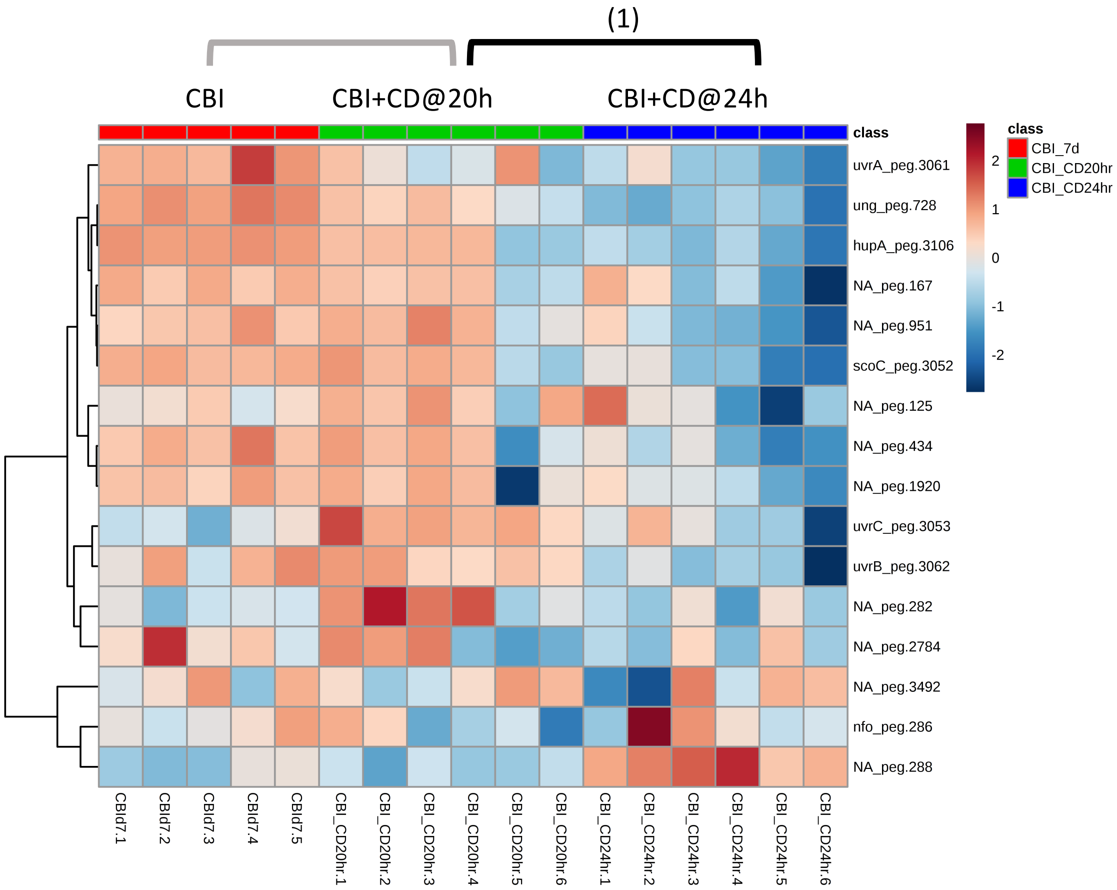


**Significantly Enriched Conditions:**

(1) Enriched in CBI+*C. difficile* co-colonized at 20h *vs* CBI +*C. difficile* co-colonized at 24h (p=4.00E-02).

### Core Cellular Machinery

#### 7.12 Electron Transport

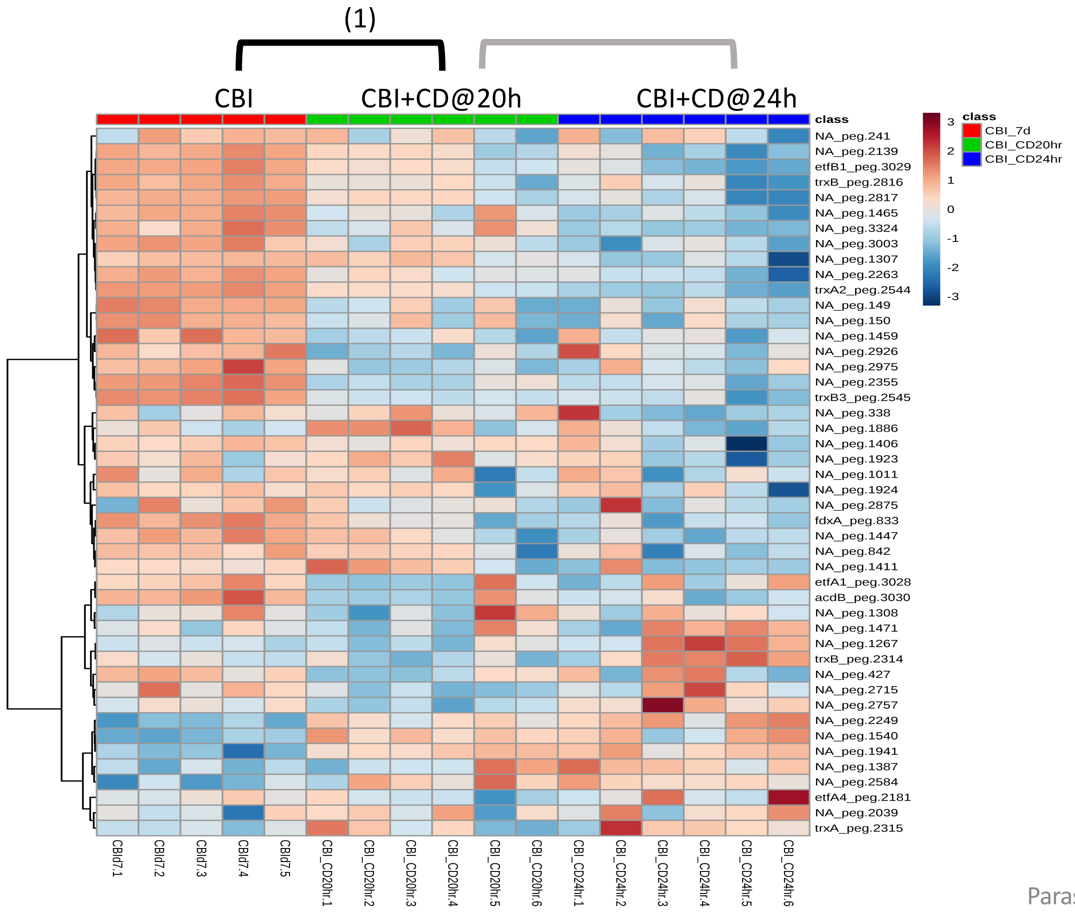


**<u>Significantly Enriched Conditions:</u>**

(1) Enriched in CBI-mono-colonized mice vs CBI+*C. difficile* co-colonized mice at 20h (p=4.01E-03).

#### Core Cellular Machinery

##### 7.13 Protein Translation and Modification

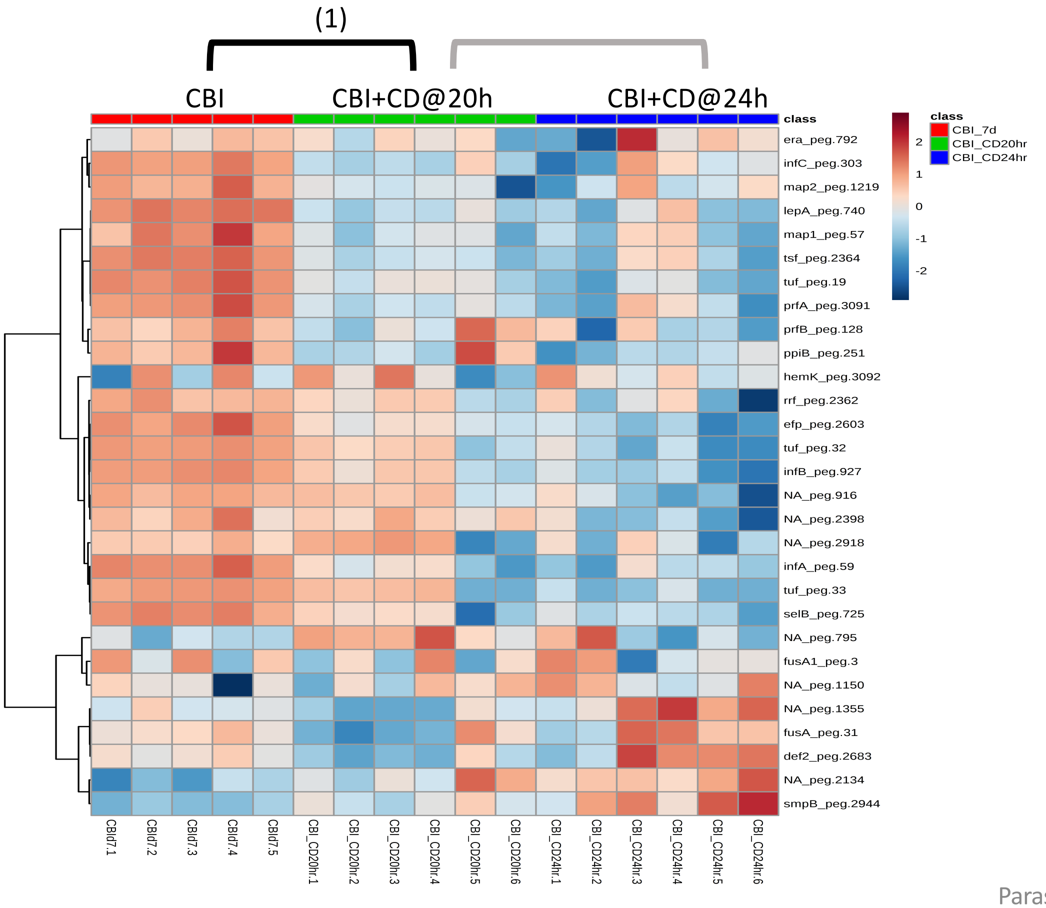


**Significantly Enriched Conditions:**

(1) Enriched in CBI-mono-colonized mice vs CBI+*C. difficile* co-colonized mice at 20h (p=8.17E-03).

### Core Cellular Machinery

#### 7.14 Ribosomal Structural Components and Modification

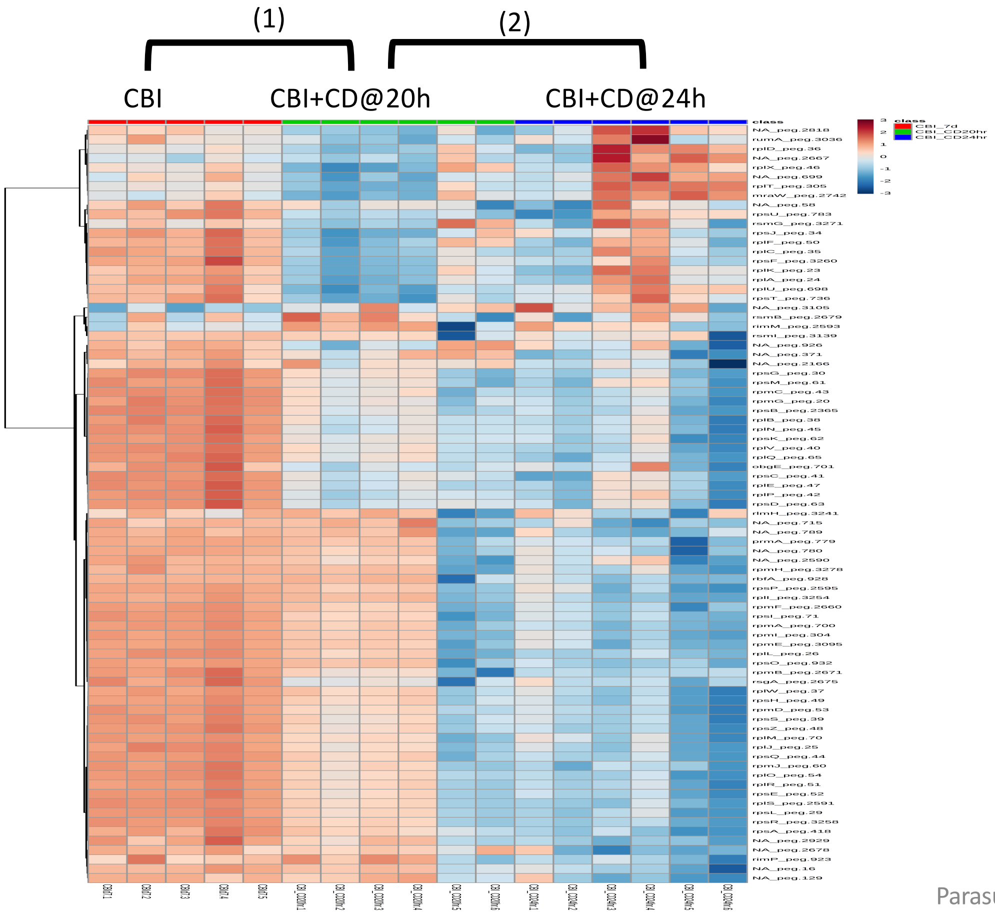


**<u>Significantly Enriched Conditions:</u>**

(1) Enriched in CBI-mono-colonized mice vs CBI+*C. difficile* co-colonized mice at 20h (p=3.37E-16).

(2) Enriched in CBI+*C. difficile* co-colonized at 20h *vs* CBI +*C. difficile* co-colonized at 24h (p=3.34E-08).

### Cell Adhesion and Motility

#### 7.15 Flagellar Motility

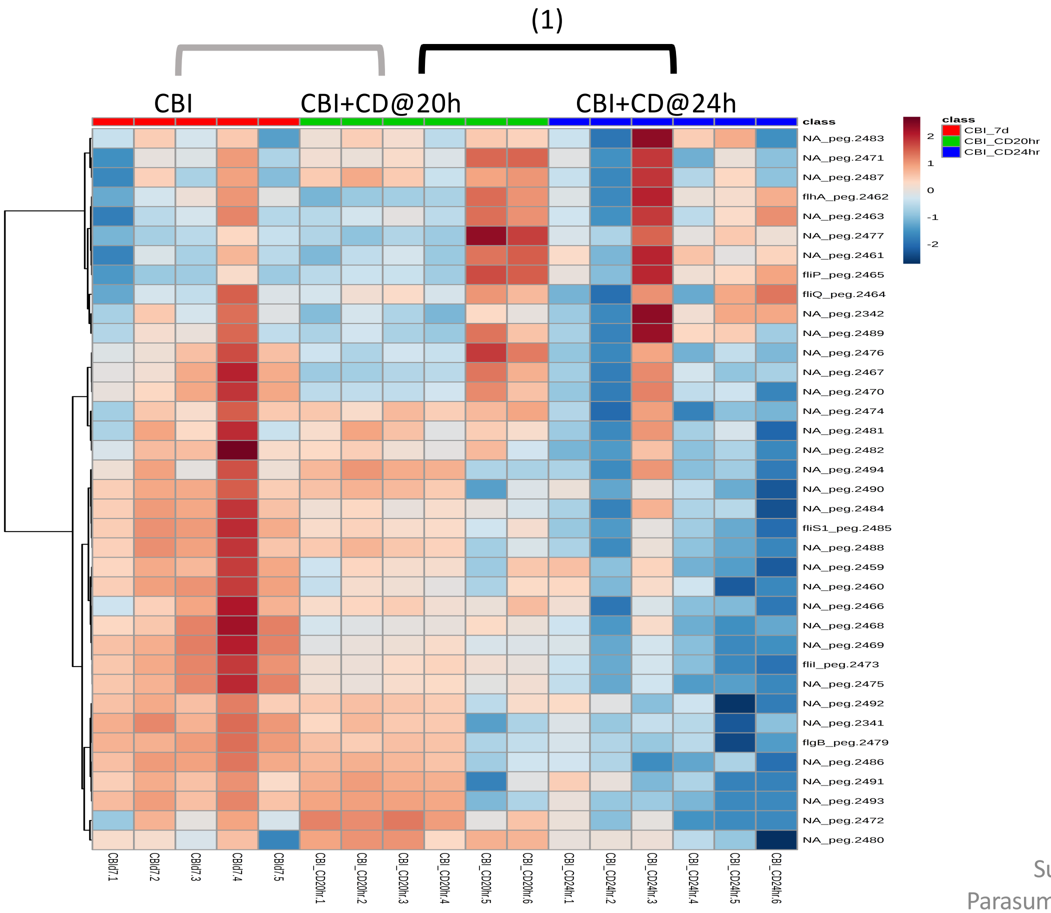


**<u>Significantly Enriched Conditions:</u>**

(1) Enriched in CBI+*C. difficile* co-colonized at 20h *vs* CBI +*C. difficile* co-colonized at 24h (p=1.24E-03).

### Cell Wall Components

#### 7.16 Gram Positive Peptidoglycan and Teichoic Acids

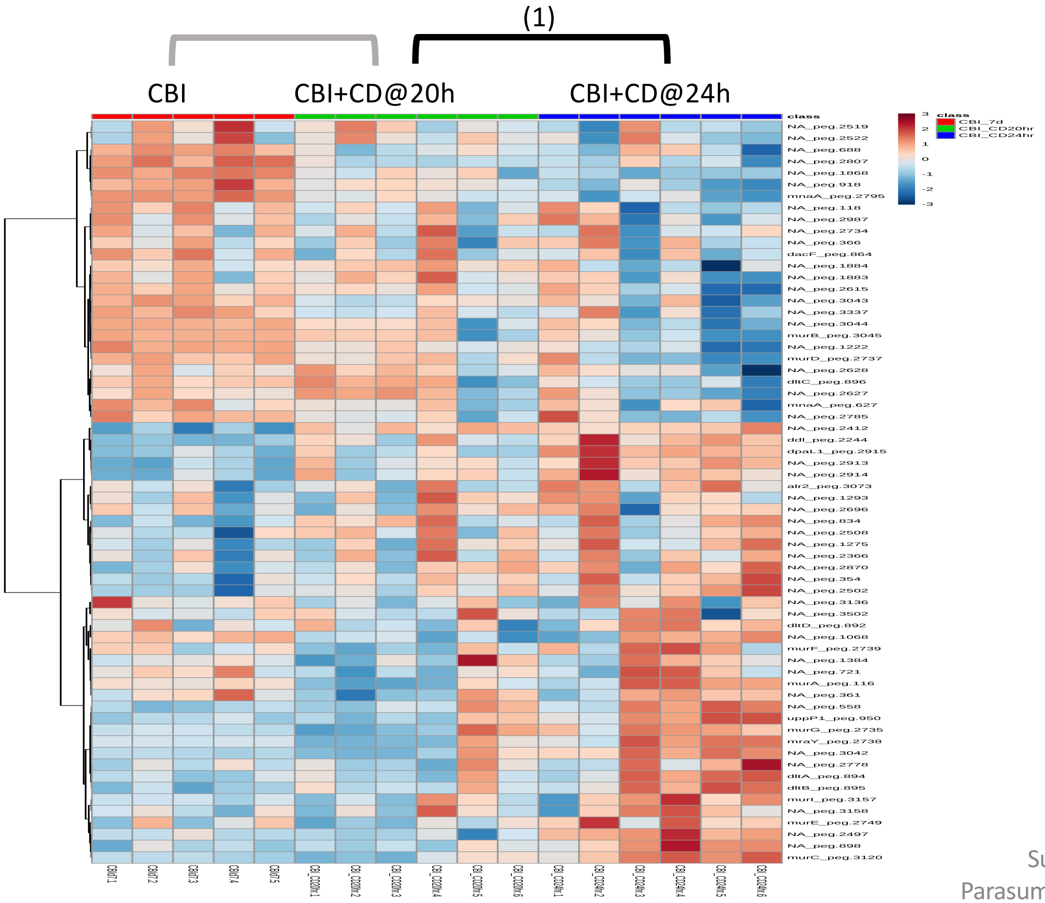


**Significantly Enriched Conditions:**

(1) Enriched in CBI+*C. difficile* co-colonized at 24h *vs* CBI +*C. difficile* co-colonized at 20h (p=2.65E-02).

### Vitamins and Micronutrient Metabolism

#### 7.17 Cobalamin Biosynthesis

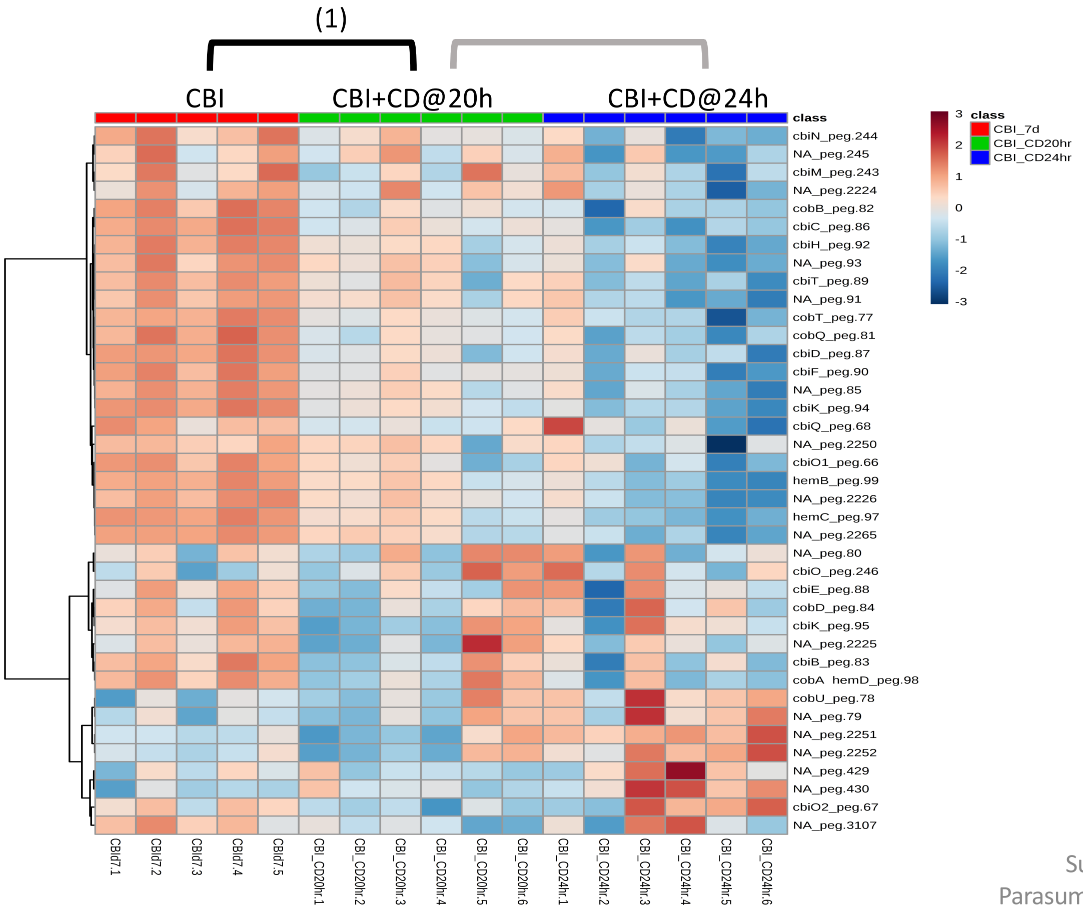


**Significantly Enriched Conditions:**

(1) Enriched in CBI-mono-colonized mice vs CBI+*C. difficile* co-colonized mice at 20h (p=2.99E-02).

### Vitamins and Micronutrient Metabolism

#### 7.18 Folate Biosynthesis

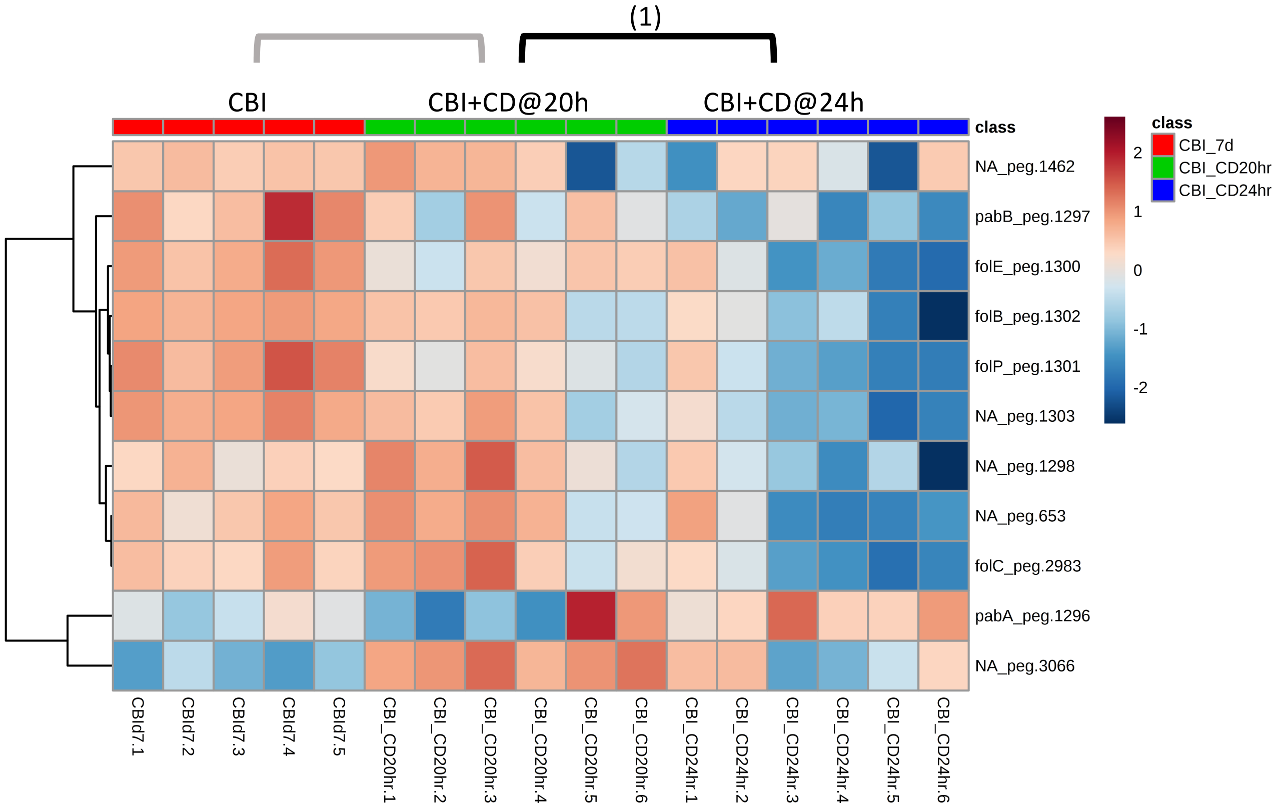


**Significantly Enriched Conditions:**

(1) Enriched in CBI+*C. difficile* co-colonized at 20h *vs* CBI +*C. difficile* co-colonized at 24h (p=3.94E-03).

### Vitamins and Micronutrient Metabolism

#### 7.19 Flavodoxins and Riboflavin Biosynthesis

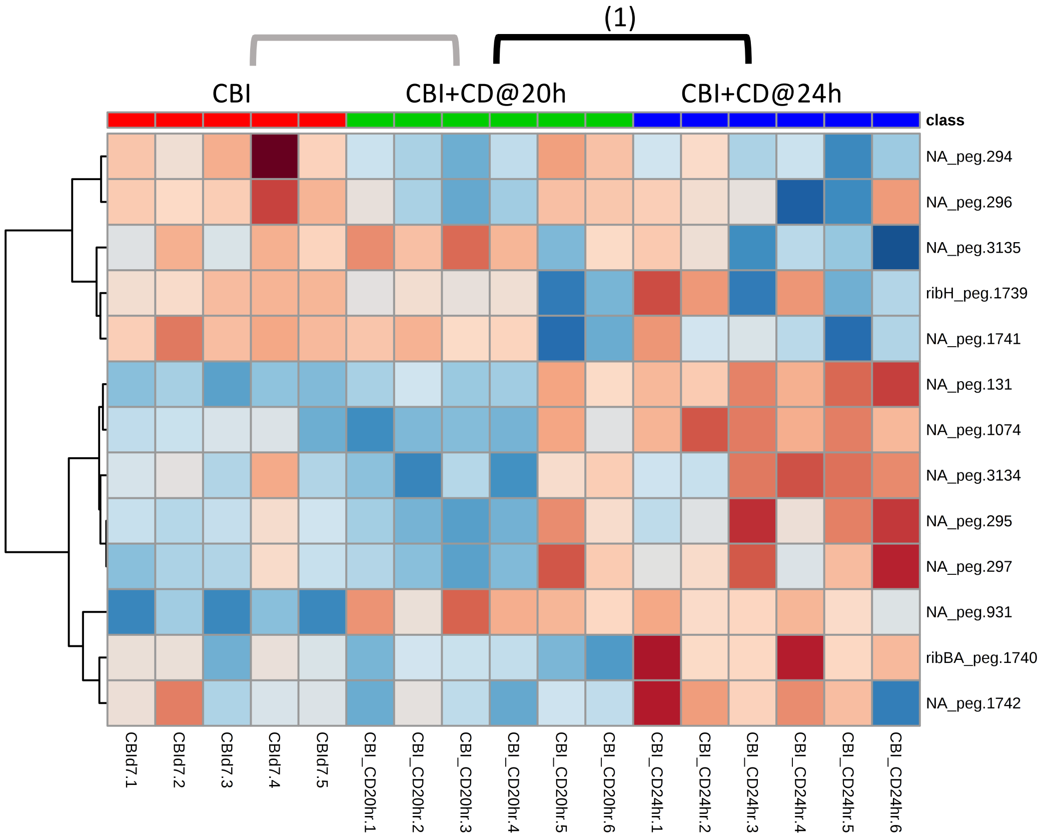


**Significantly Enriched Conditions:**

(1) Enriched in CBI+*C. difficile* co-colonized at 24h *vs* CBI +*C. difficile* co-colonized at 20h (p=1.05E-02).

### Vitamins and Micronutrient Metabolism

#### 7.20 Iron Transport and Storage

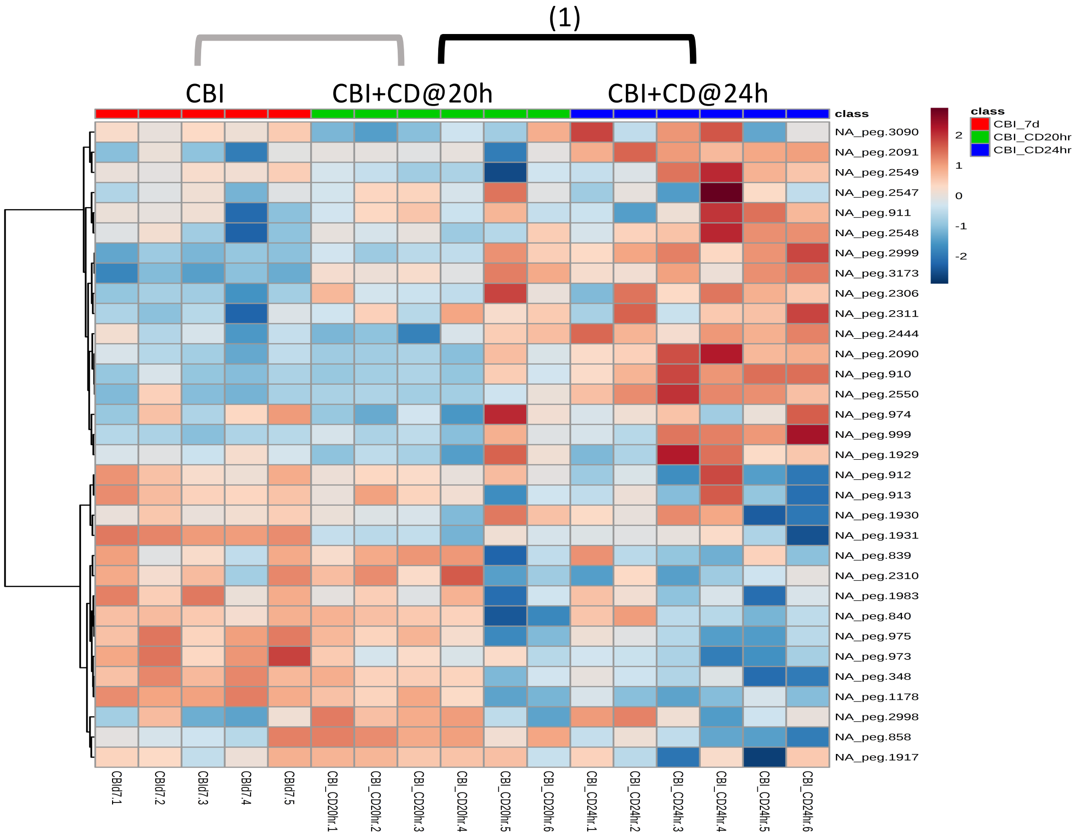


**Significantly Enriched Conditions:**

(1) Enriched in CBI+*C. difficile* co-colonized at 24h *vs* CBI +*C. difficile* co-colonized at 20h (p=2.88E-02).

### Vitamins and Micronutrient Metabolism

#### 7.21 Sulfur-Compound Transport and Metabolism

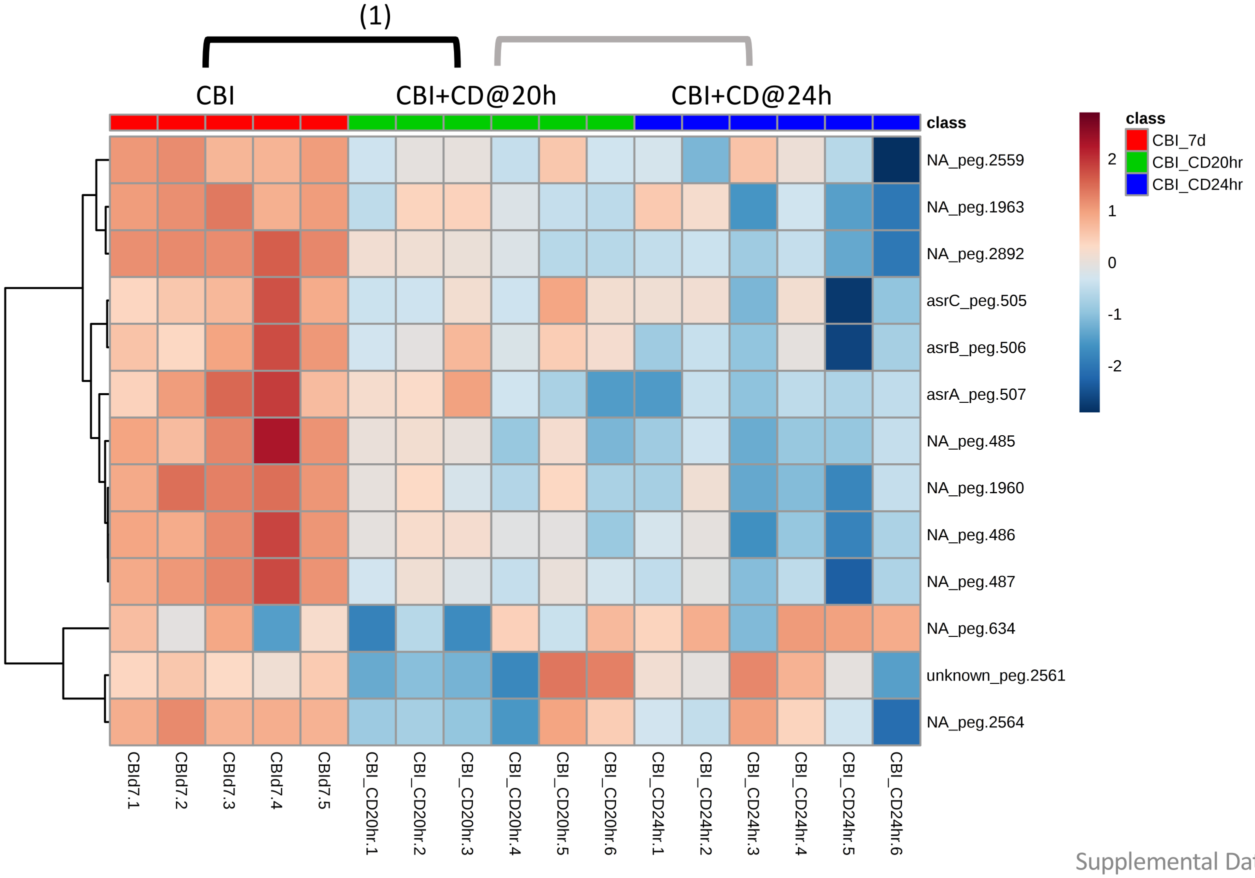


**<u>Significantly Enriched Conditions:</u>**

(1) Enriched in CBI-mono-colonized mice vs CBI+*C. difficile* co-colonized mice at 20h (p=4.80E-03).

### Vitamins and Micronutrient Metabolism

#### 7.22 Transport Binding Proteins: Cations

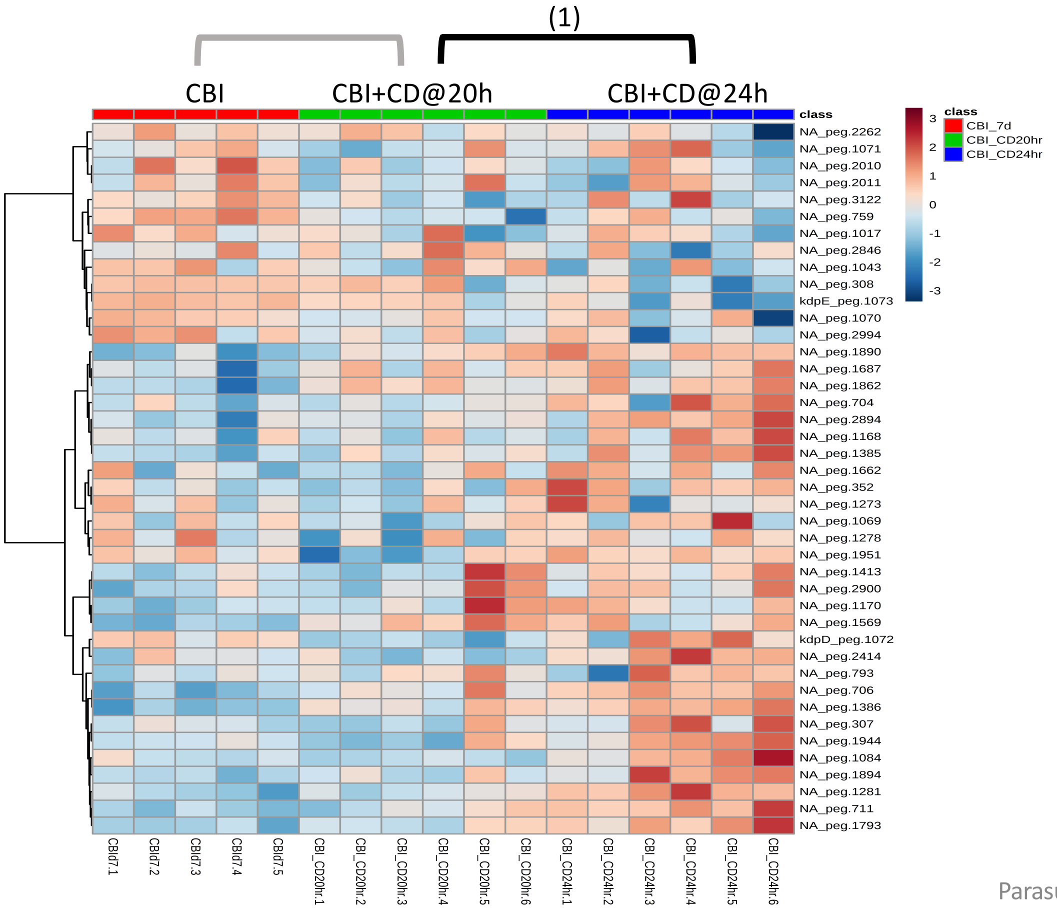


**Significantly Enriched Conditions:**

(1) Enriched in CBI+*C. difficile* co-colonized at 24h *vs* CBI +*C. difficile* co-colonized at 20h (p=5.88E-03).

### Other Transport Systems

#### 7.23 ABC Transport Systems

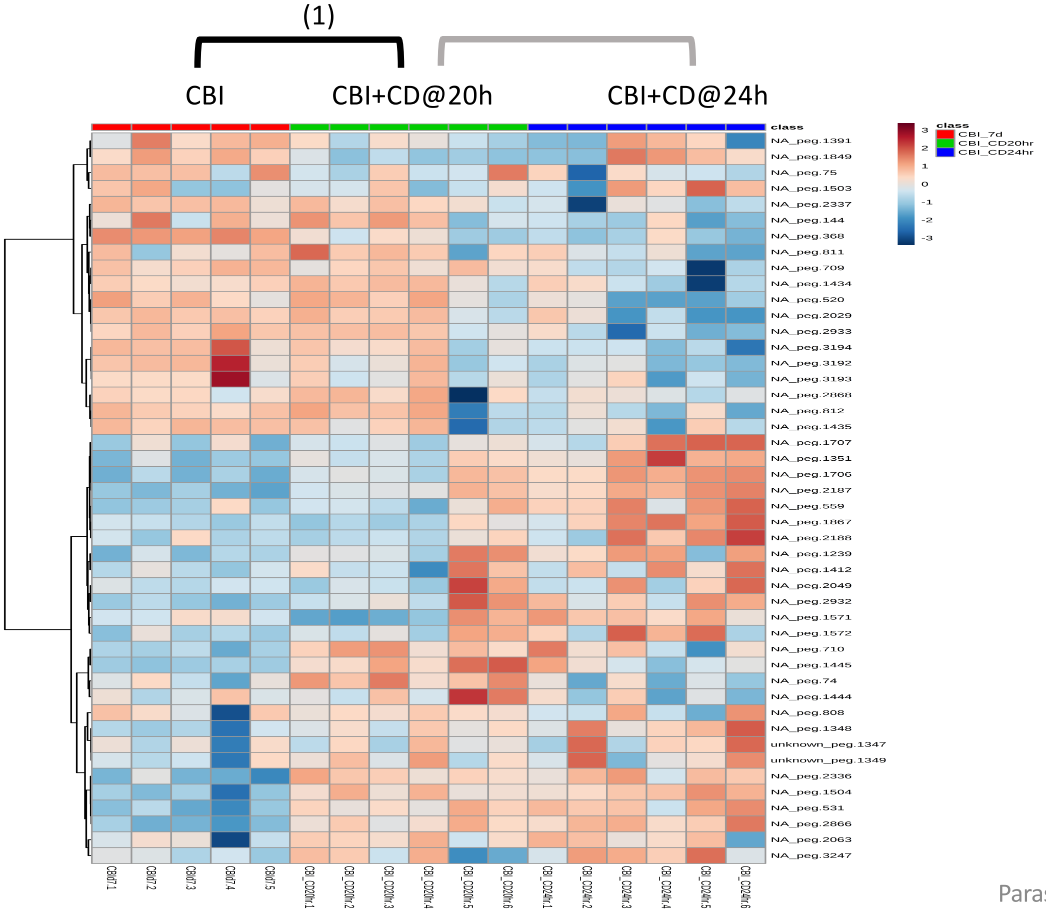


**<u>Significantly Enriched Conditions:</u>**

(1) Enriched in CBI+*C. difficile* co-colonized *vs* CBI mono-colonized mice at 20h (p=8.58E-03).

### Other Transport Systems

#### 7.24 Multi-Drug Family Transport Systems

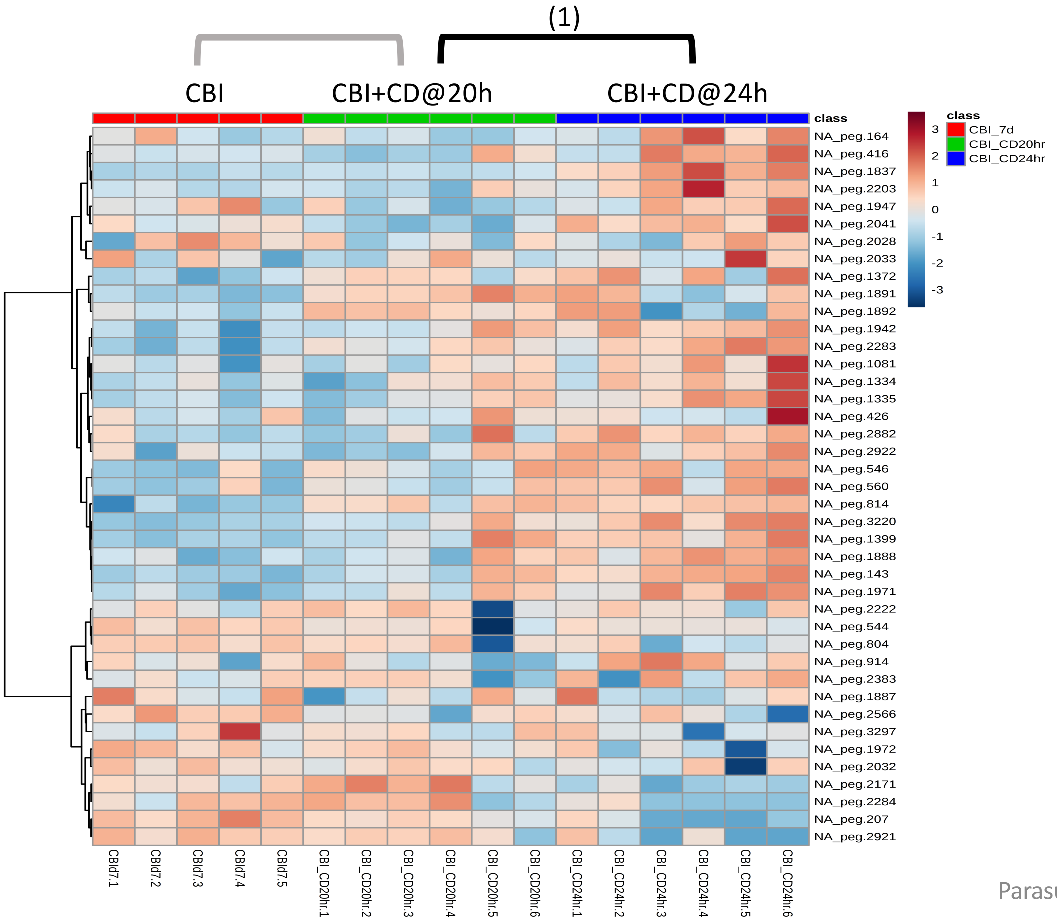


**Significantly Enriched Conditions:**

(1) Enriched in CBI+*C. difficile* co-colonized at 24h *vs* CBI +*C. difficile* co-colonized at 20h (p=4.32E-02).

### Bacteriophage and Mobile Elements

#### 7.25 Putative Phage Locus Genes

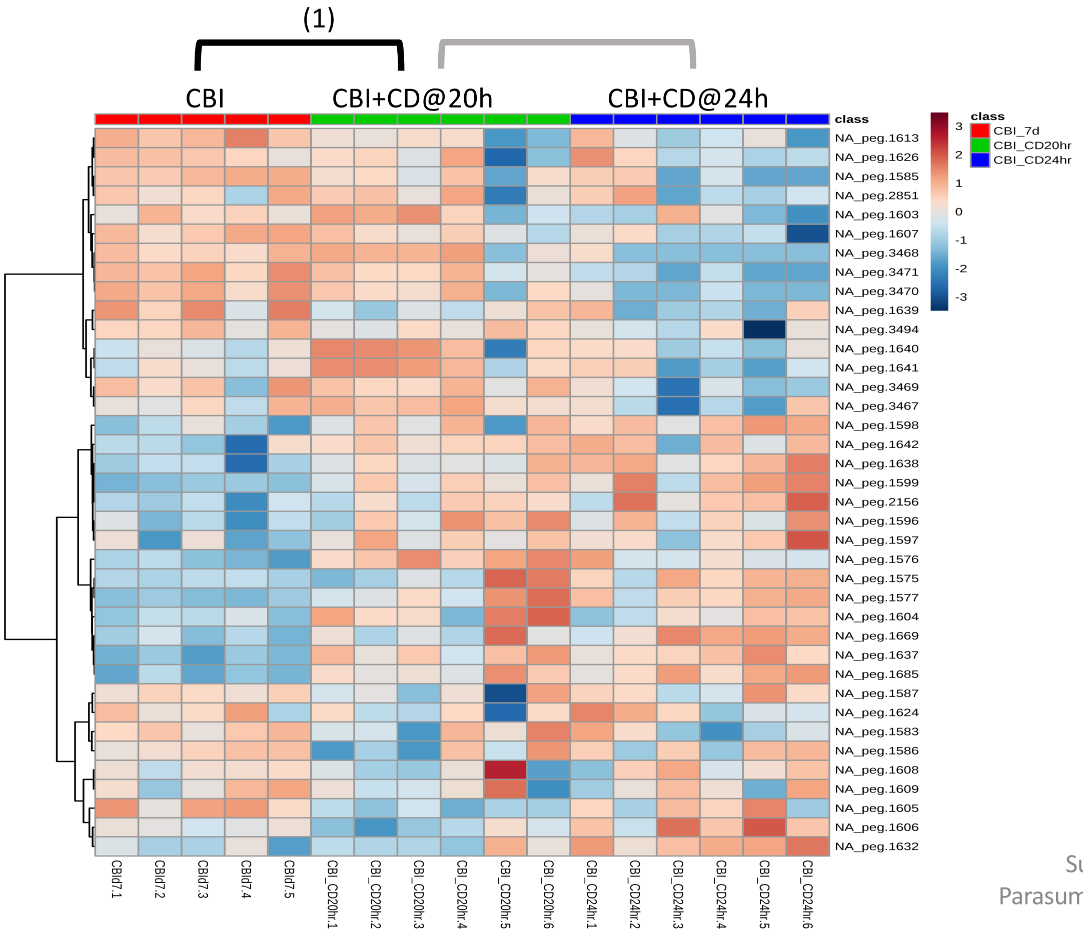


**Significantly Enriched Conditions:**

(1) Enriched in CBI+*C. difficile* co-colonized *vs* CBI mono-colonized mice at 20h (p=6.12E-04).

### Stress Responses

#### 7.26 Cellular Stress Responses

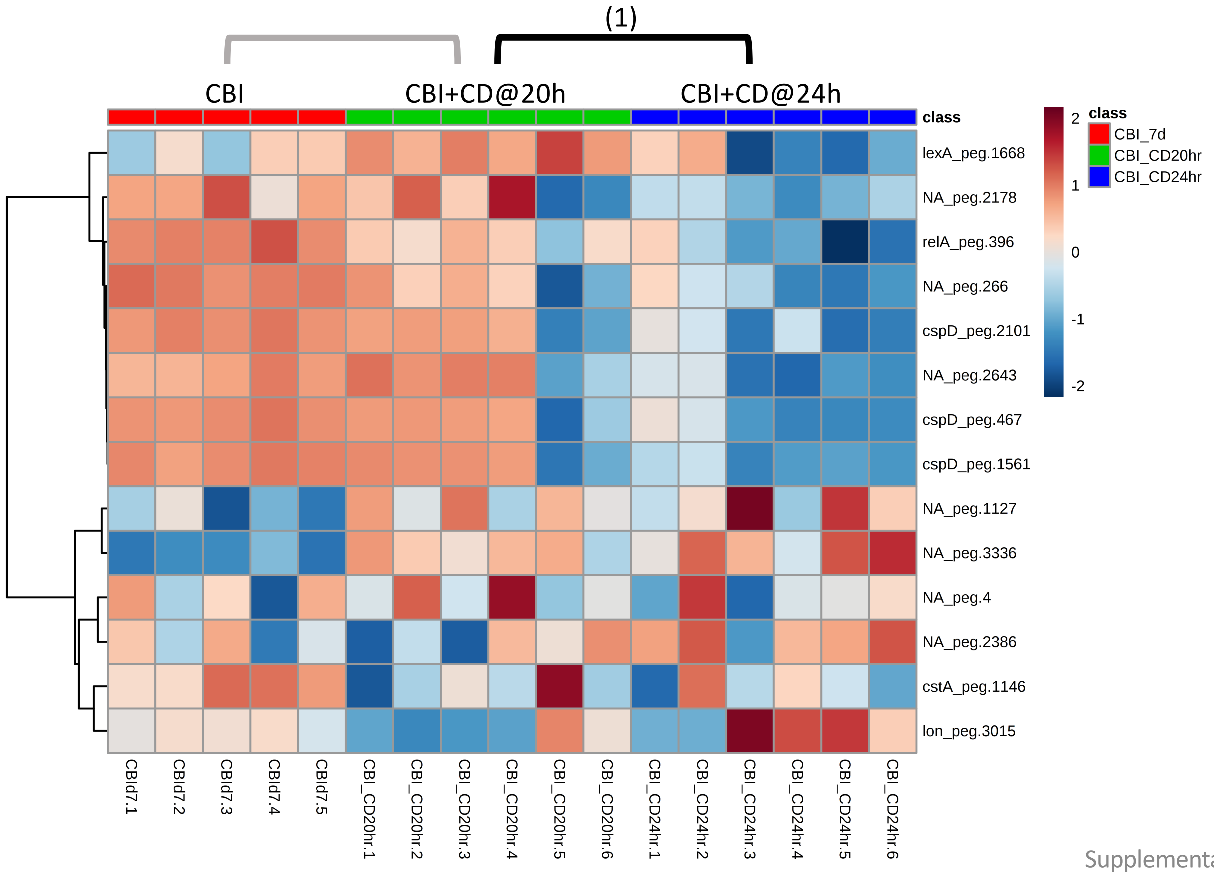


**<u>Significantly Enriched Conditions:</u>**

(1) Enriched in CBI+*C. difficile* co-colonized at 20h *vs* CBI +*C. difficile* co-colonized at 24h (p=2.27E-02).
