## Supplemental Data File 10 for "*In vivo* commensal control of *Clostridioides difficile* virulence"

### Slide 1
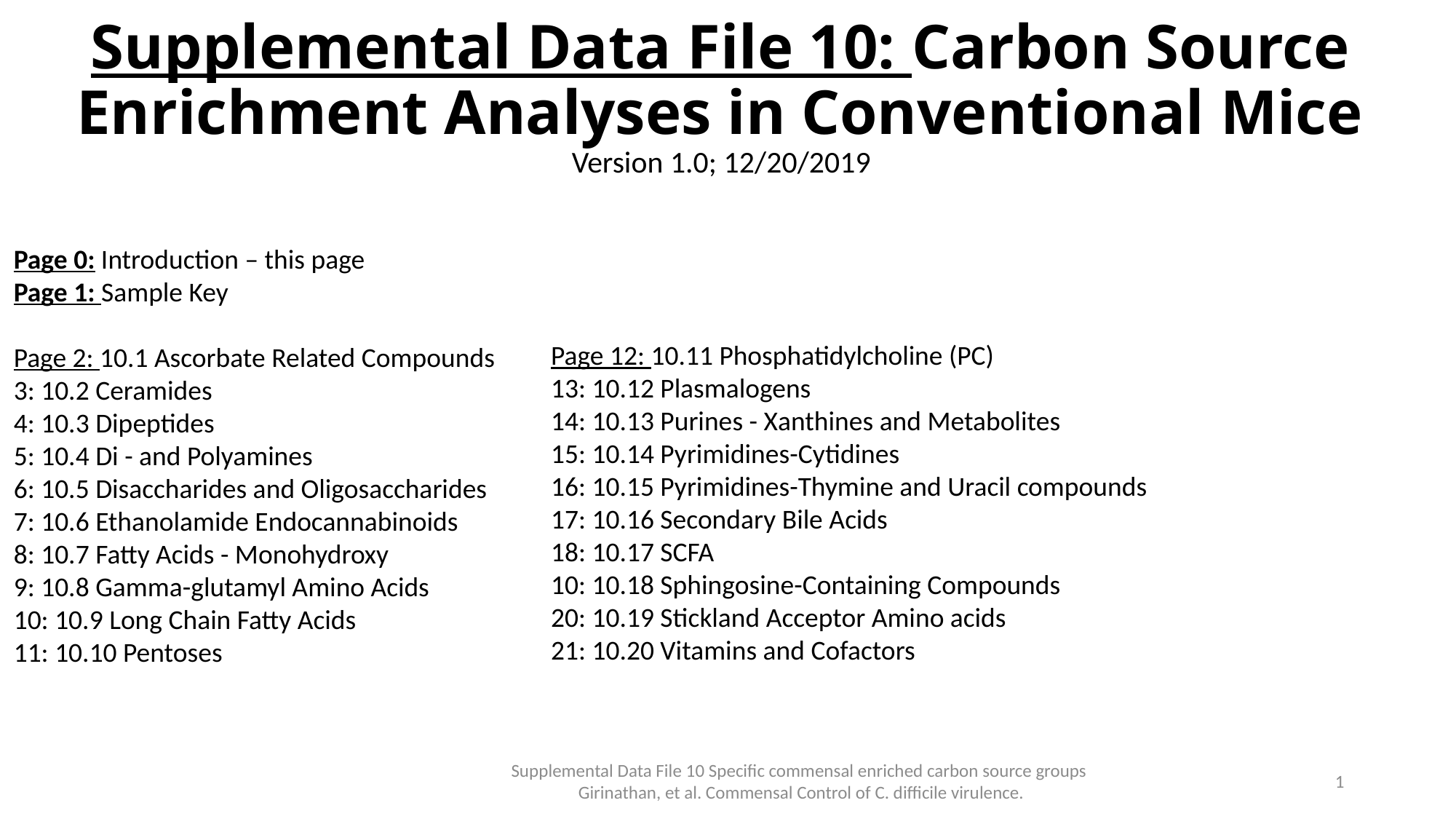

Supplemental Data File 10: Carbon Source Enrichment Analyses in Conventional Mice
Version 1.0; 12/20/2019
Page 0: Introduction – this page
Page 1: Sample Key
Page 2: 10.1 Ascorbate Related Compounds
3: 10.2 Ceramides
4: 10.3 Dipeptides
5: 10.4 Di - and Polyamines
6: 10.5 Disaccharides and Oligosaccharides
7: 10.6 Ethanolamide Endocannabinoids
8: 10.7 Fatty Acids - Monohydroxy
9: 10.8 Gamma-glutamyl Amino Acids
10: 10.9 Long Chain Fatty Acids
11: 10.10 Pentoses
Page 12: 10.11 Phosphatidylcholine (PC)
13: 10.12 Plasmalogens
14: 10.13 Purines - Xanthines and Metabolites
15: 10.14 Pyrimidines-Cytidines
16: 10.15 Pyrimidines-Thymine and Uracil compounds
17: 10.16 Secondary Bile Acids
18: 10.17 SCFA
10: 10.18 Sphingosine-Containing Compounds
20: 10.19 Stickland Acceptor Amino acids
21: 10.20 Vitamins and Cofactors
Supplemental Data File 10 Specific commensal enriched carbon source groups Girinathan, et al. Commensal Control of C. difficile virulence.
0

### Slide 2
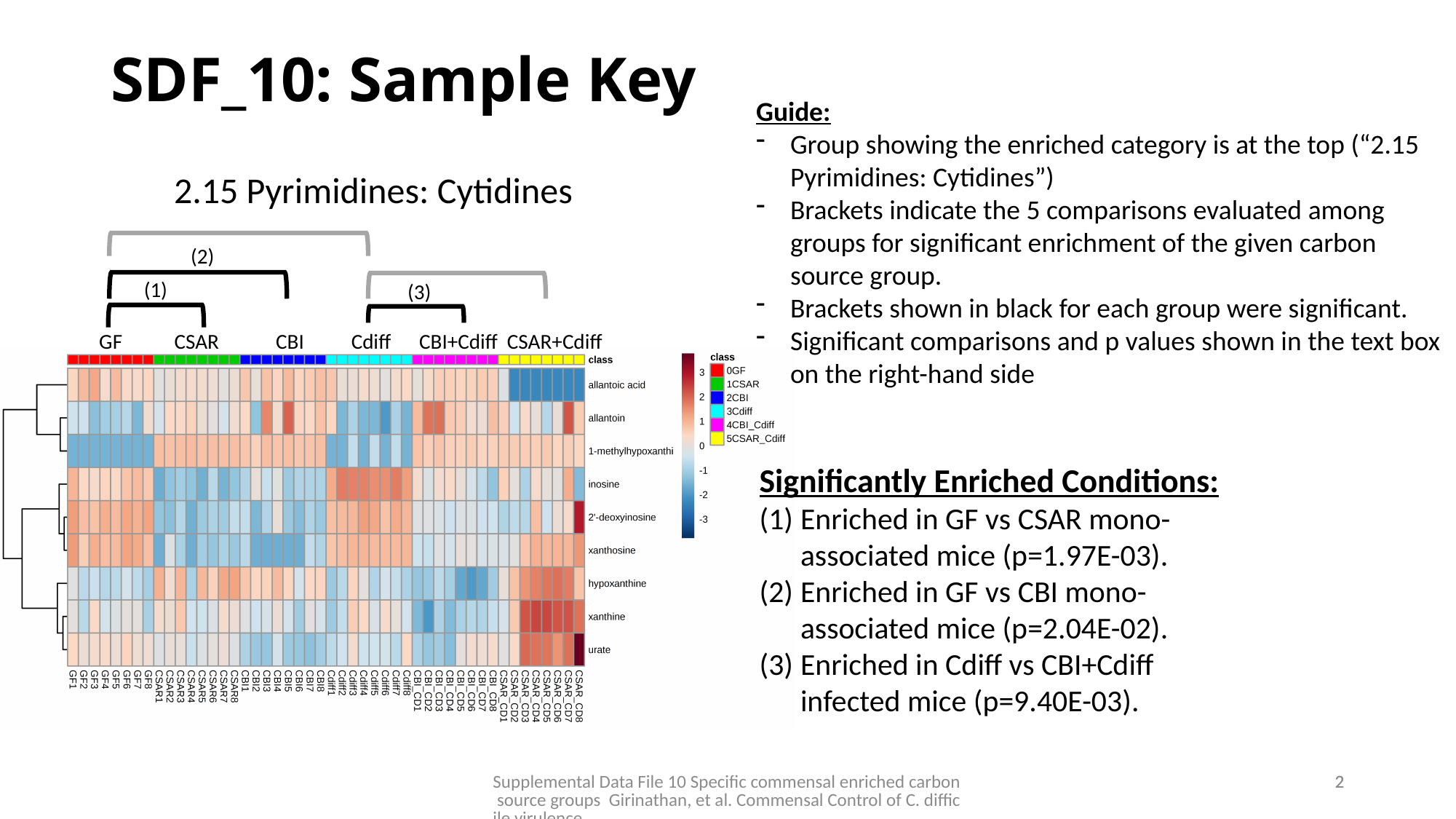

SDF_10: Sample Key
Guide:
Group showing the enriched category is at the top (“2.15 Pyrimidines: Cytidines”)
Brackets indicate the 5 comparisons evaluated among groups for significant enrichment of the given carbon source group.
Brackets shown in black for each group were significant.
Significant comparisons and p values shown in the text box on the right-hand side
2.15 Pyrimidines: Cytidines
(2)
(1)
(3)
GF CSAR CBI Cdiff CBI+Cdiff CSAR+Cdiff
Significantly Enriched Conditions:
Enriched in GF vs CSAR mono-associated mice (p=1.97E-03).
Enriched in GF vs CBI mono-associated mice (p=2.04E-02).
Enriched in Cdiff vs CBI+Cdiff infected mice (p=9.40E-03).
Supplemental Data File 10 Specific commensal enriched carbon source groups Girinathan, et al. Commensal Control of C. difficile virulence.
1
1

### Slide 3
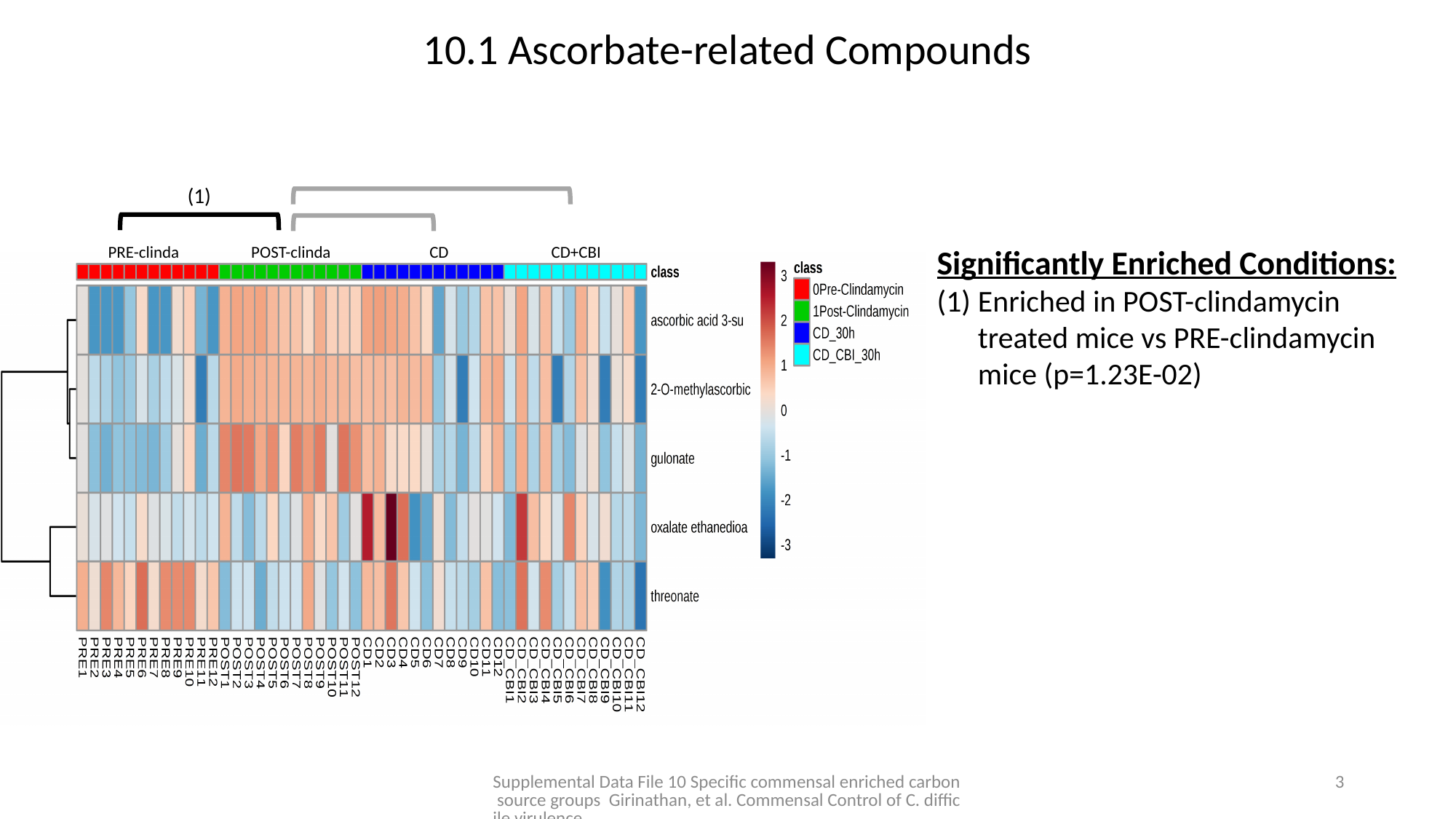

10.1 Ascorbate-related Compounds
(1)
PRE-clinda POST-clinda CD CD+CBI
Significantly Enriched Conditions:
Enriched in POST-clindamycin treated mice vs PRE-clindamycin mice (p=1.23E-02)
Supplemental Data File 10 Specific commensal enriched carbon source groups Girinathan, et al. Commensal Control of C. difficile virulence.
2

### Slide 4
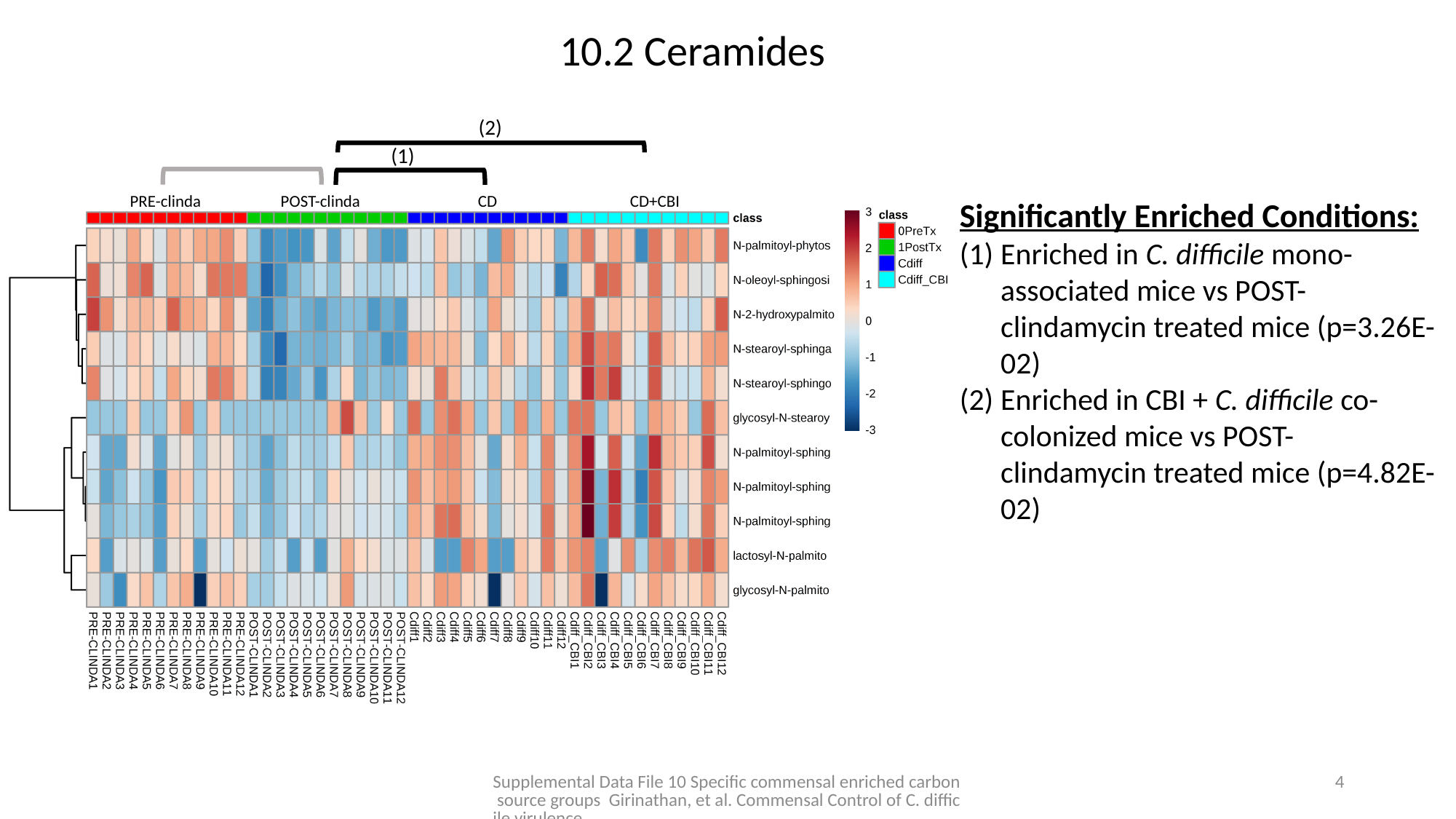

10.2 Ceramides
(2)
(1)
PRE-clinda POST-clinda CD CD+CBI
Significantly Enriched Conditions:
Enriched in C. difficile mono-associated mice vs POST-clindamycin treated mice (p=3.26E-02)
Enriched in CBI + C. difficile co-colonized mice vs POST-clindamycin treated mice (p=4.82E-02)
Supplemental Data File 10 Specific commensal enriched carbon source groups Girinathan, et al. Commensal Control of C. difficile virulence.
3

### Slide 5
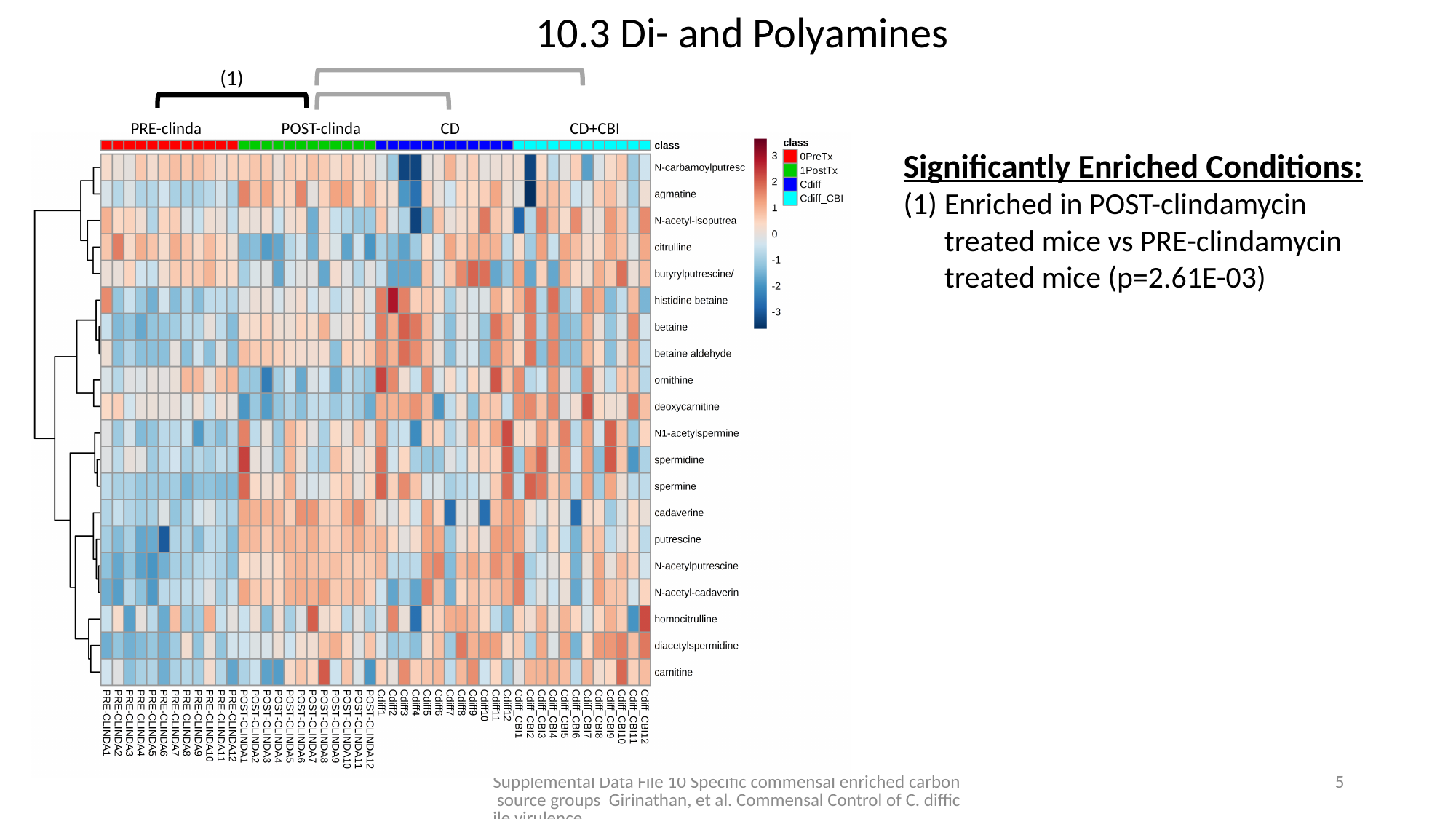

10.3 Di- and Polyamines
(1)
PRE-clinda POST-clinda CD CD+CBI
Significantly Enriched Conditions:
Enriched in POST-clindamycin treated mice vs PRE-clindamycin treated mice (p=2.61E-03)
Supplemental Data File 10 Specific commensal enriched carbon source groups Girinathan, et al. Commensal Control of C. difficile virulence.
4

### Slide 6
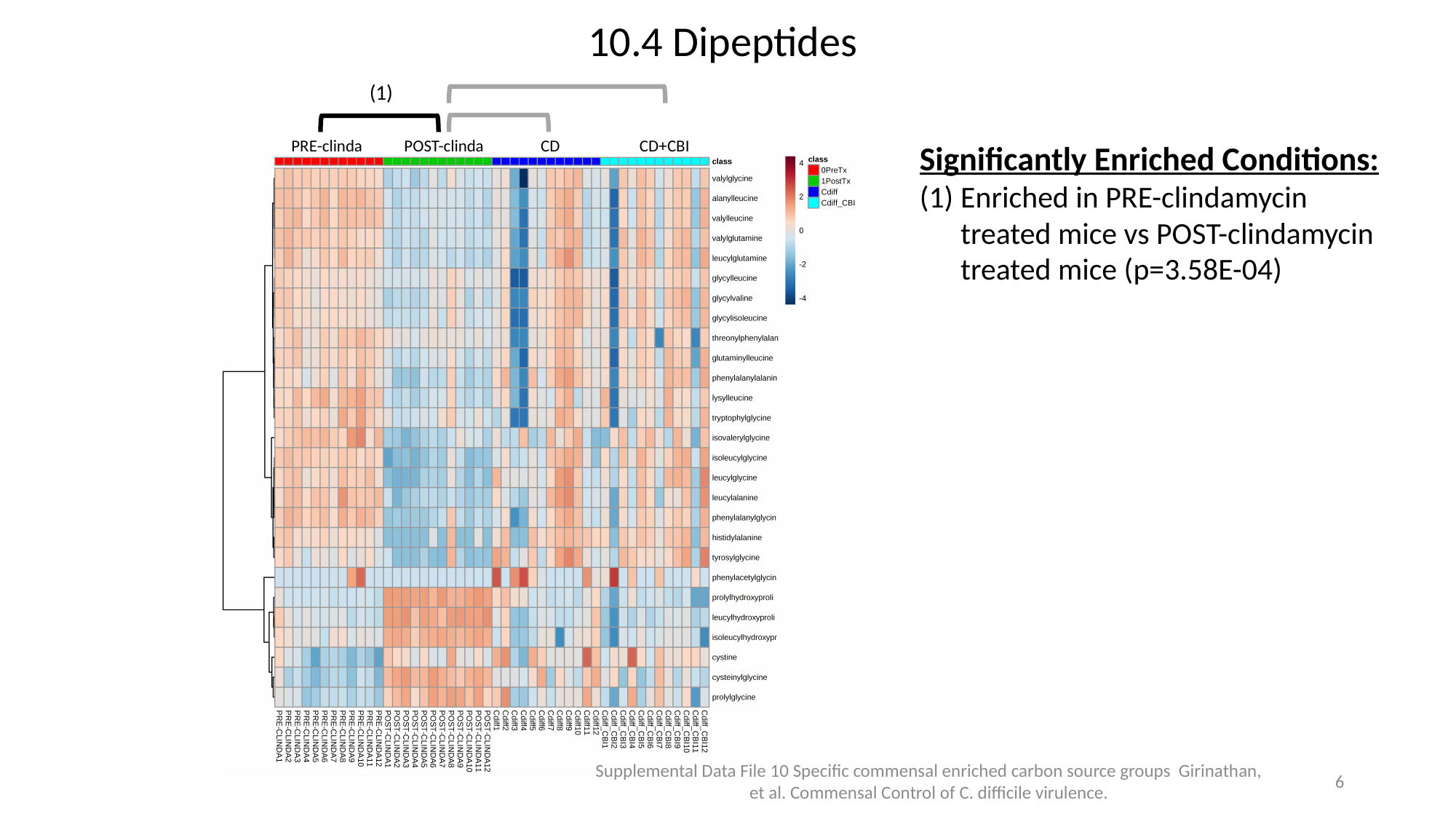

10.4 Dipeptides
(1)
PRE-clinda POST-clinda CD CD+CBI
Significantly Enriched Conditions:
Enriched in PRE-clindamycin treated mice vs POST-clindamycin treated mice (p=3.58E-04)
Supplemental Data File 10 Specific commensal enriched carbon source groups Girinathan, et al. Commensal Control of C. difficile virulence.
5

### Slide 7
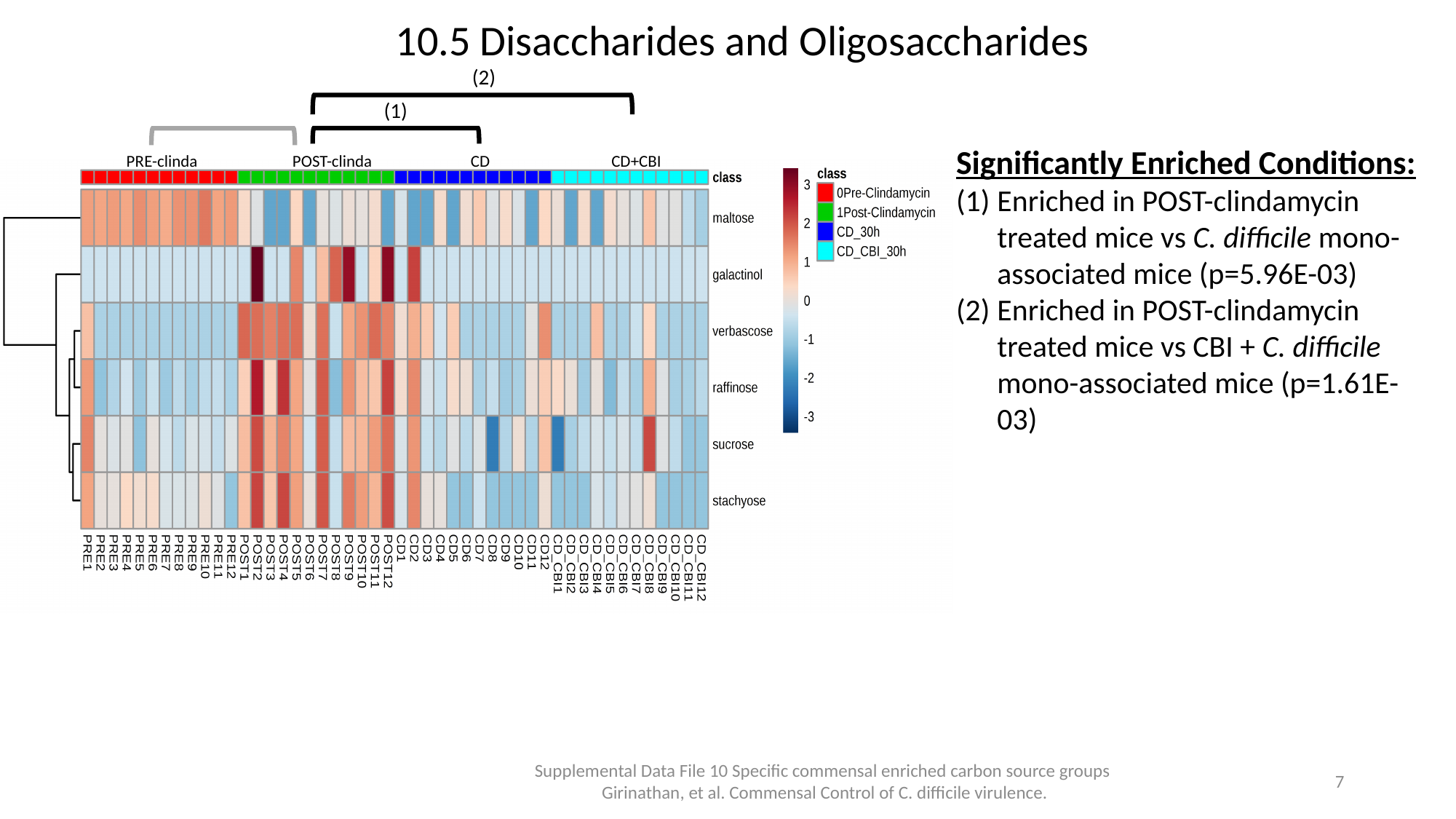

10.5 Disaccharides and Oligosaccharides
(2)
(1)
Significantly Enriched Conditions:
Enriched in POST-clindamycin treated mice vs C. difficile mono-associated mice (p=5.96E-03)
Enriched in POST-clindamycin treated mice vs CBI + C. difficile mono-associated mice (p=1.61E-03)
PRE-clinda POST-clinda CD CD+CBI
Supplemental Data File 10 Specific commensal enriched carbon source groups Girinathan, et al. Commensal Control of C. difficile virulence.
6

### Slide 8
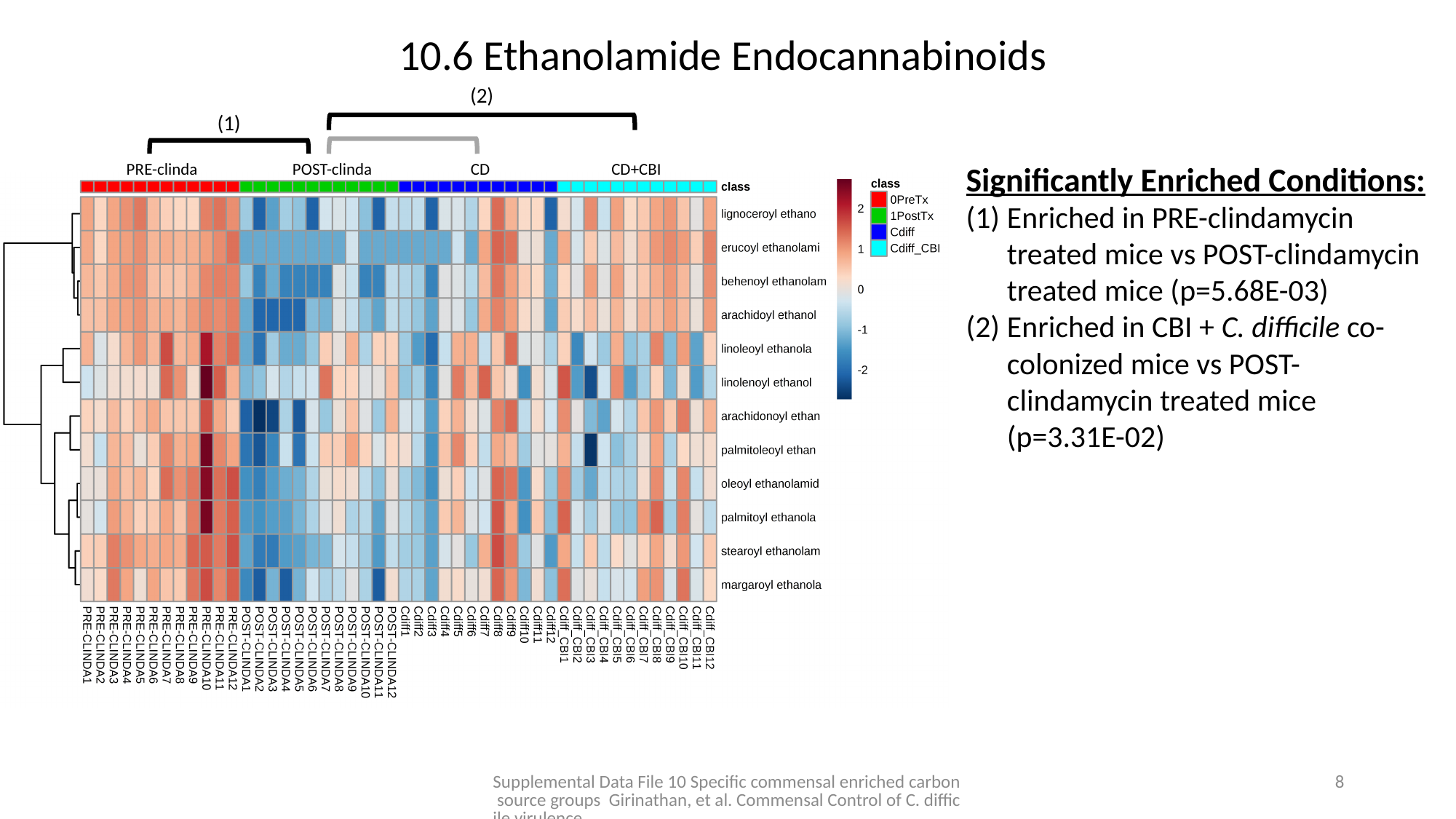

10.6 Ethanolamide Endocannabinoids
(2)
(1)
PRE-clinda POST-clinda CD CD+CBI
Significantly Enriched Conditions:
Enriched in PRE-clindamycin treated mice vs POST-clindamycin treated mice (p=5.68E-03)
Enriched in CBI + C. difficile co-colonized mice vs POST-clindamycin treated mice (p=3.31E-02)
Supplemental Data File 10 Specific commensal enriched carbon source groups Girinathan, et al. Commensal Control of C. difficile virulence.
7

### Slide 9
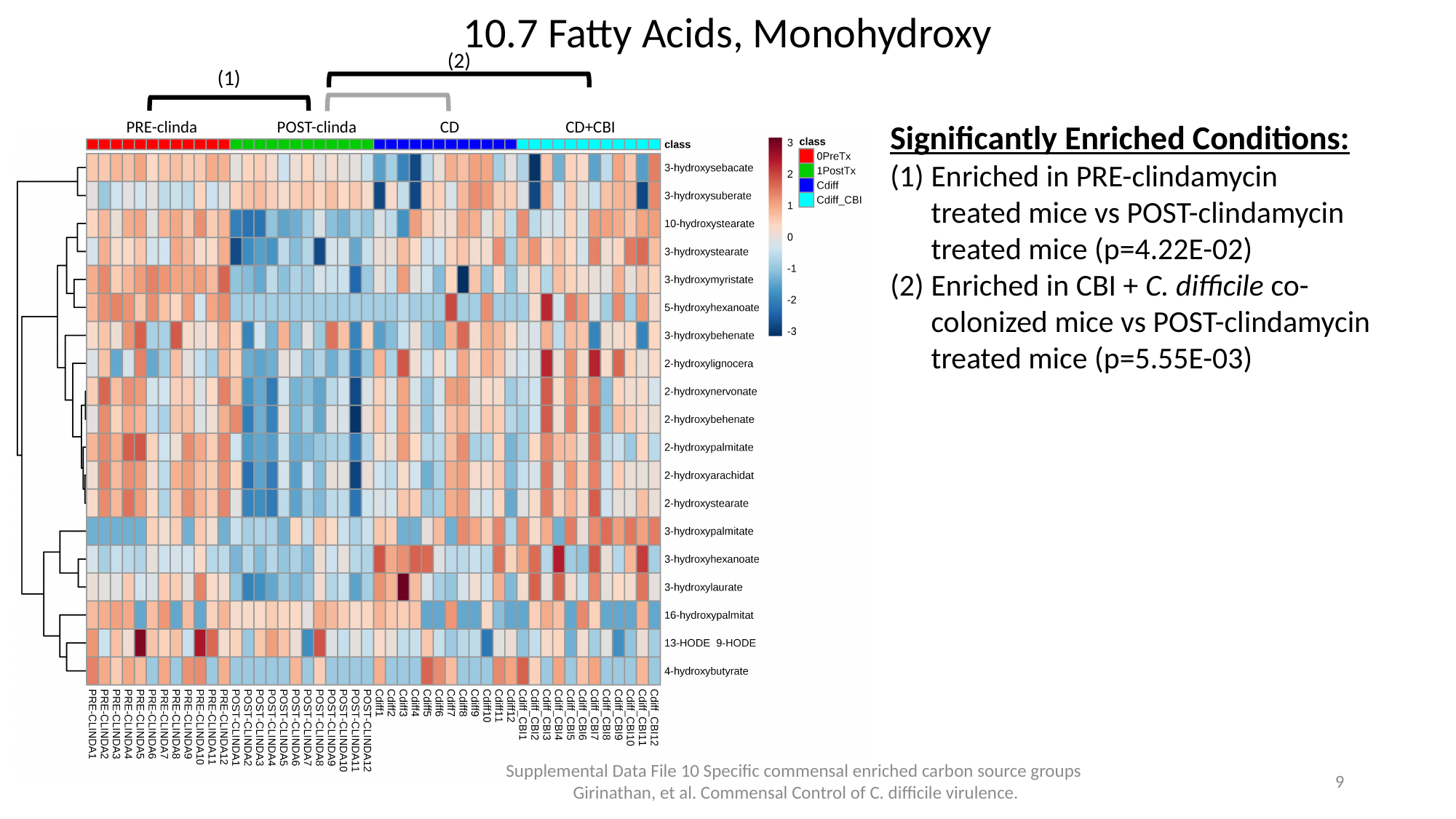

10.7 Fatty Acids, Monohydroxy
(2)
(1)
PRE-clinda POST-clinda CD CD+CBI
Significantly Enriched Conditions:
Enriched in PRE-clindamycin treated mice vs POST-clindamycin treated mice (p=4.22E-02)
Enriched in CBI + C. difficile co-colonized mice vs POST-clindamycin treated mice (p=5.55E-03)
Supplemental Data File 10 Specific commensal enriched carbon source groups Girinathan, et al. Commensal Control of C. difficile virulence.
8

### Slide 10
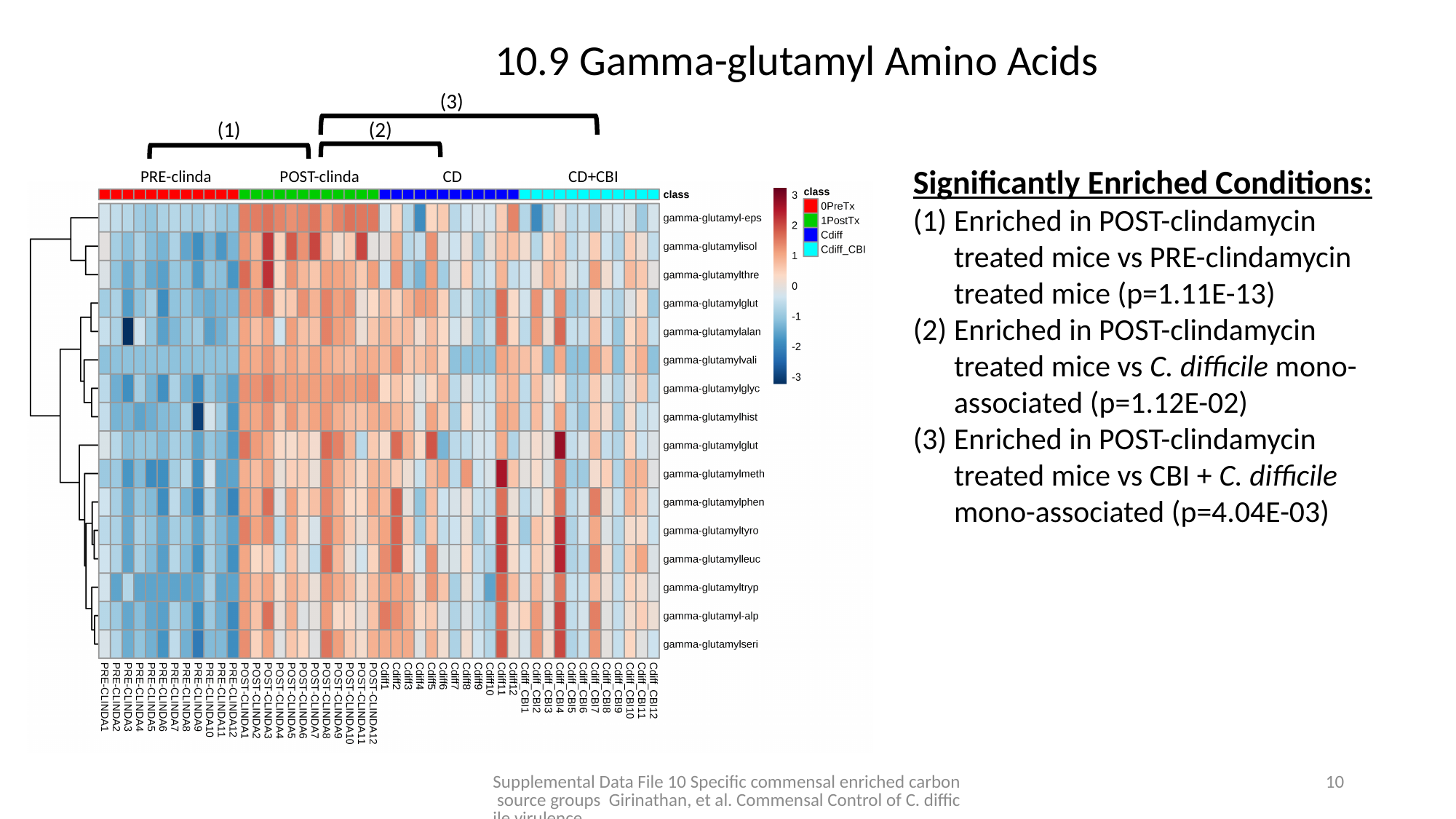

10.9 Gamma-glutamyl Amino Acids
(3)
(1)
(2)
Significantly Enriched Conditions:
Enriched in POST-clindamycin treated mice vs PRE-clindamycin treated mice (p=1.11E-13)
Enriched in POST-clindamycin treated mice vs C. difficile mono-associated (p=1.12E-02)
Enriched in POST-clindamycin treated mice vs CBI + C. difficile mono-associated (p=4.04E-03)
PRE-clinda POST-clinda CD CD+CBI
Supplemental Data File 10 Specific commensal enriched carbon source groups Girinathan, et al. Commensal Control of C. difficile virulence.
9

### Slide 11
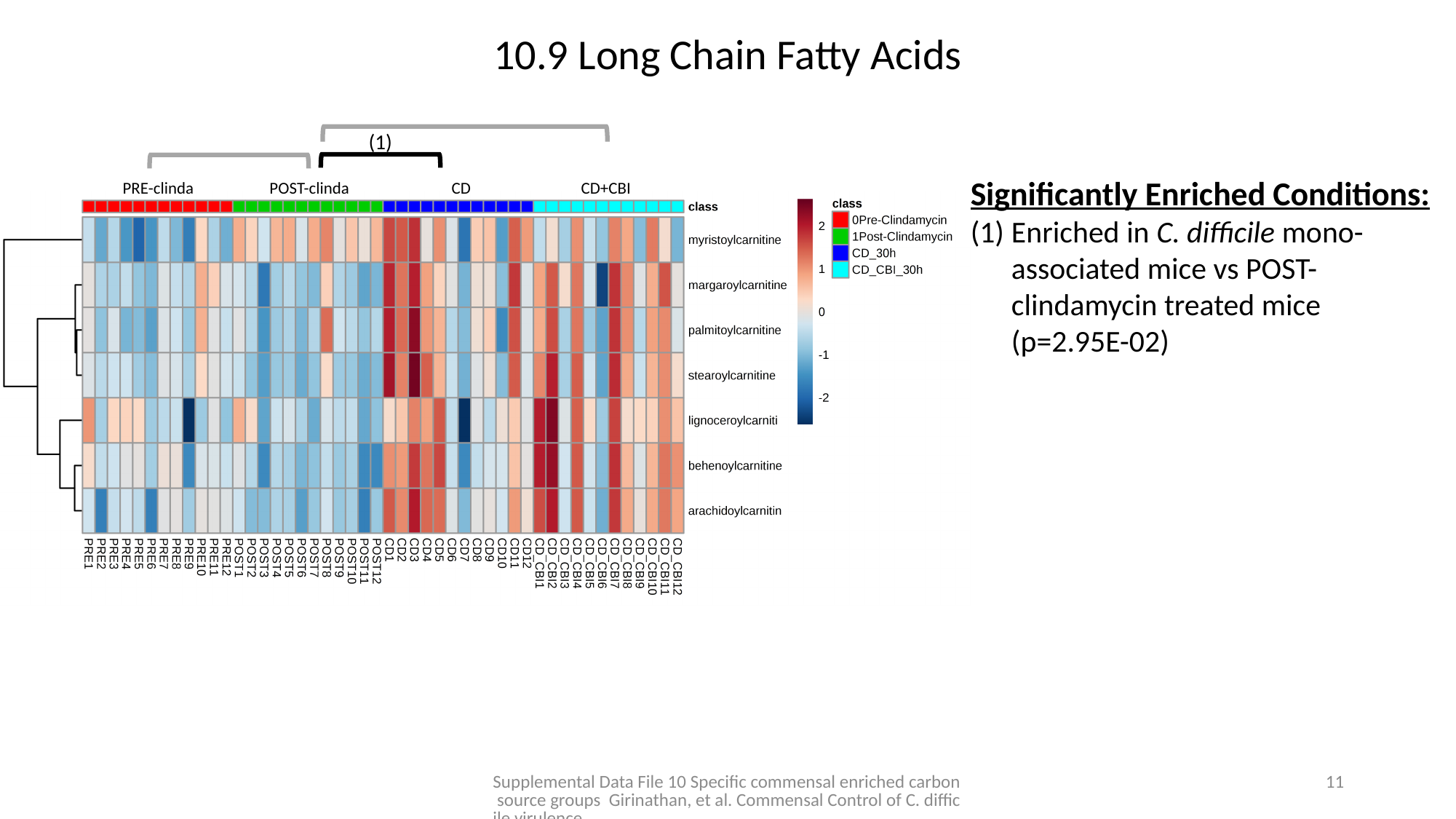

10.9 Long Chain Fatty Acids
(1)
Significantly Enriched Conditions:
Enriched in C. difficile mono-associated mice vs POST-clindamycin treated mice (p=2.95E-02)
PRE-clinda POST-clinda CD CD+CBI
Supplemental Data File 10 Specific commensal enriched carbon source groups Girinathan, et al. Commensal Control of C. difficile virulence.
10

### Slide 12
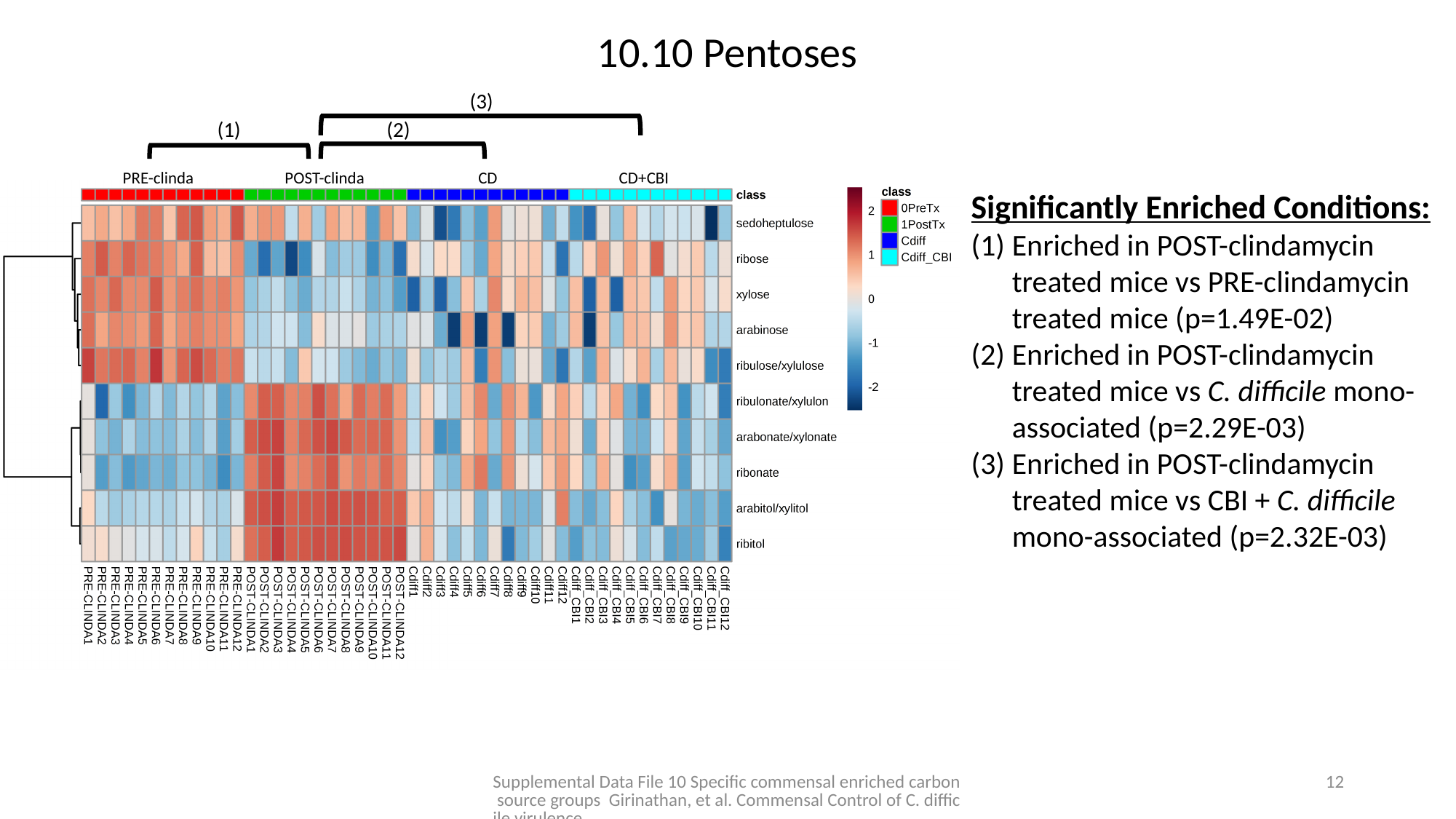

10.10 Pentoses
(3)
(1)
(2)
PRE-clinda POST-clinda CD CD+CBI
Significantly Enriched Conditions:
Enriched in POST-clindamycin treated mice vs PRE-clindamycin treated mice (p=1.49E-02)
Enriched in POST-clindamycin treated mice vs C. difficile mono-associated (p=2.29E-03)
Enriched in POST-clindamycin treated mice vs CBI + C. difficile mono-associated (p=2.32E-03)
Supplemental Data File 10 Specific commensal enriched carbon source groups Girinathan, et al. Commensal Control of C. difficile virulence.
11

### Slide 13
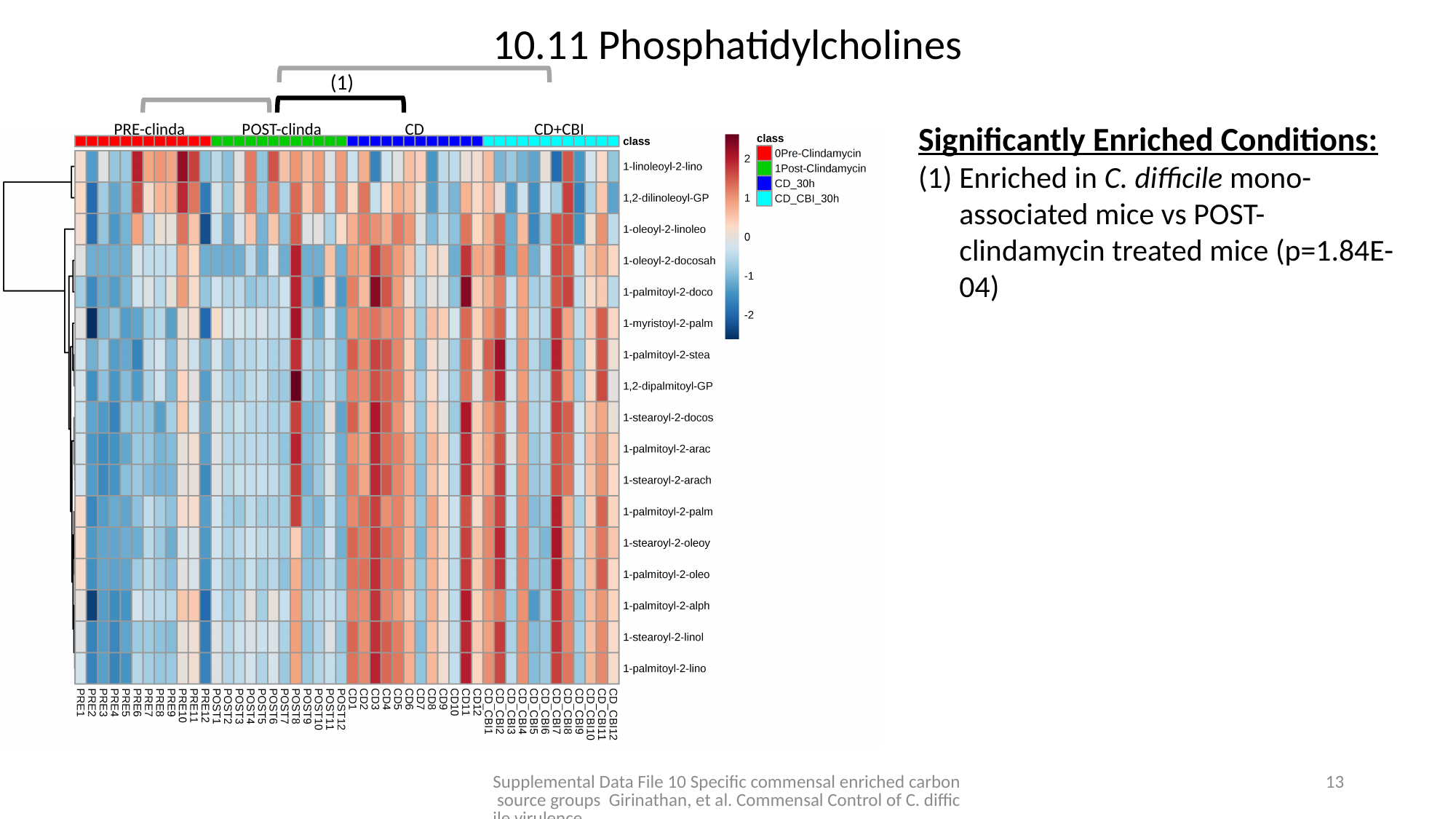

10.11 Phosphatidylcholines
(1)
Significantly Enriched Conditions:
Enriched in C. difficile mono-associated mice vs POST-clindamycin treated mice (p=1.84E-04)
PRE-clinda POST-clinda CD CD+CBI
Supplemental Data File 10 Specific commensal enriched carbon source groups Girinathan, et al. Commensal Control of C. difficile virulence.
12

### Slide 14
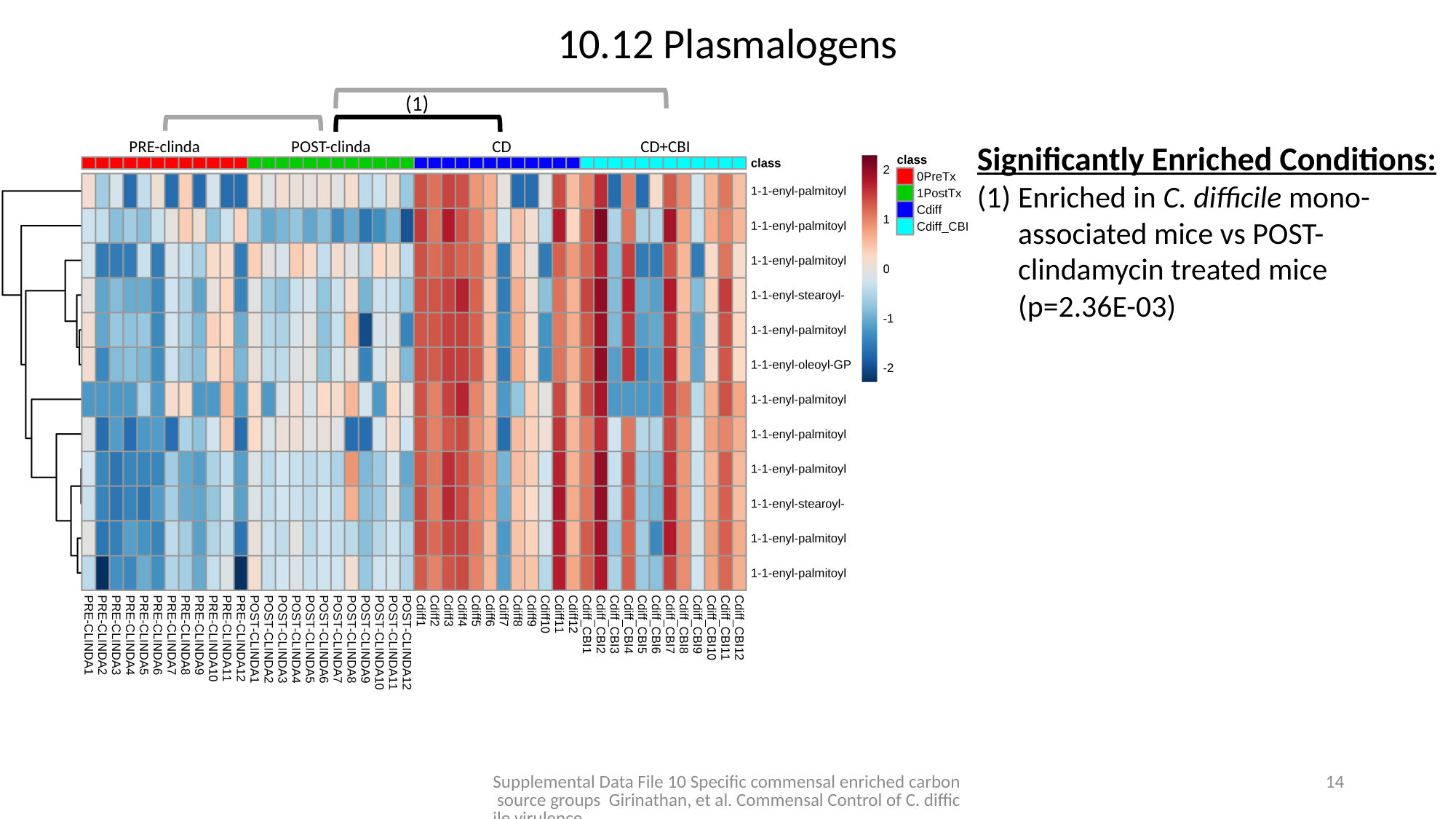

10.12 Plasmalogens
(1)
PRE-clinda POST-clinda CD CD+CBI
Significantly Enriched Conditions:
Enriched in C. difficile mono-associated mice vs POST-clindamycin treated mice (p=2.36E-03)
Supplemental Data File 10 Specific commensal enriched carbon source groups Girinathan, et al. Commensal Control of C. difficile virulence.
13

### Slide 15
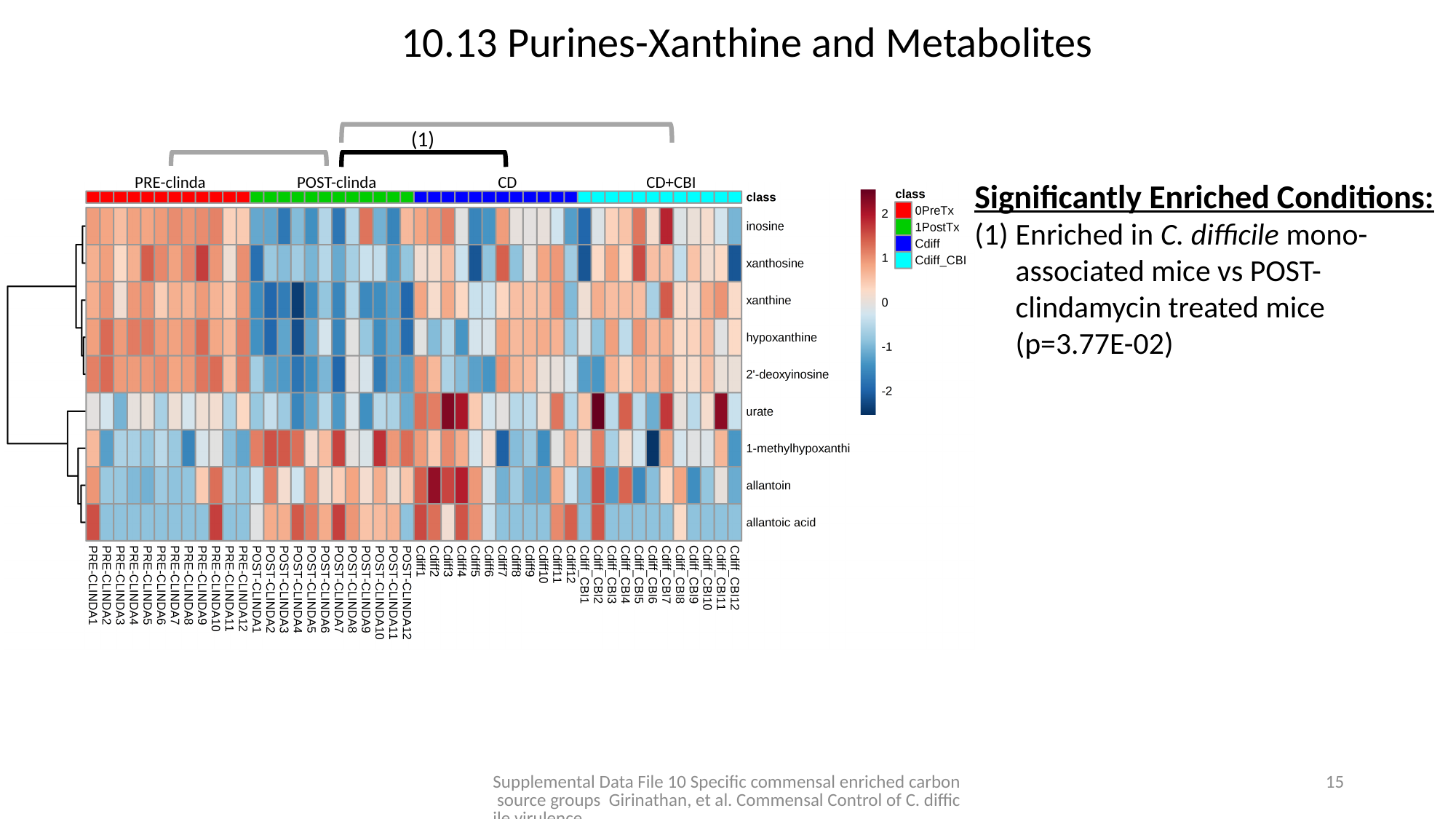

10.13 Purines-Xanthine and Metabolites
(1)
PRE-clinda POST-clinda CD CD+CBI
Significantly Enriched Conditions:
Enriched in C. difficile mono-associated mice vs POST-clindamycin treated mice (p=3.77E-02)
Supplemental Data File 10 Specific commensal enriched carbon source groups Girinathan, et al. Commensal Control of C. difficile virulence.
14

### Slide 16
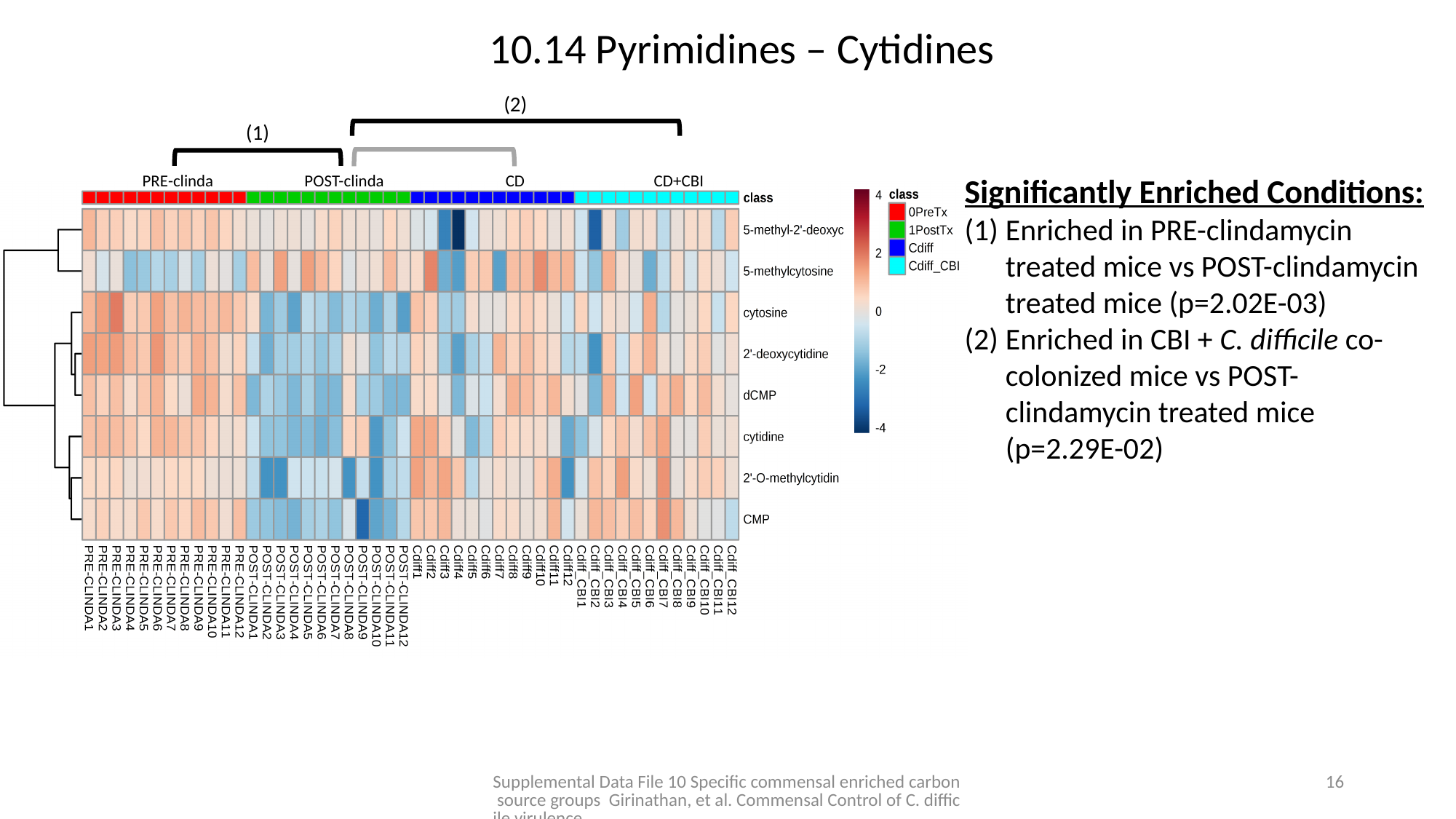

10.14 Pyrimidines – Cytidines
(2)
(1)
PRE-clinda POST-clinda CD CD+CBI
Significantly Enriched Conditions:
Enriched in PRE-clindamycin treated mice vs POST-clindamycin treated mice (p=2.02E-03)
Enriched in CBI + C. difficile co-colonized mice vs POST-clindamycin treated mice (p=2.29E-02)
Supplemental Data File 10 Specific commensal enriched carbon source groups Girinathan, et al. Commensal Control of C. difficile virulence.
15

### Slide 17
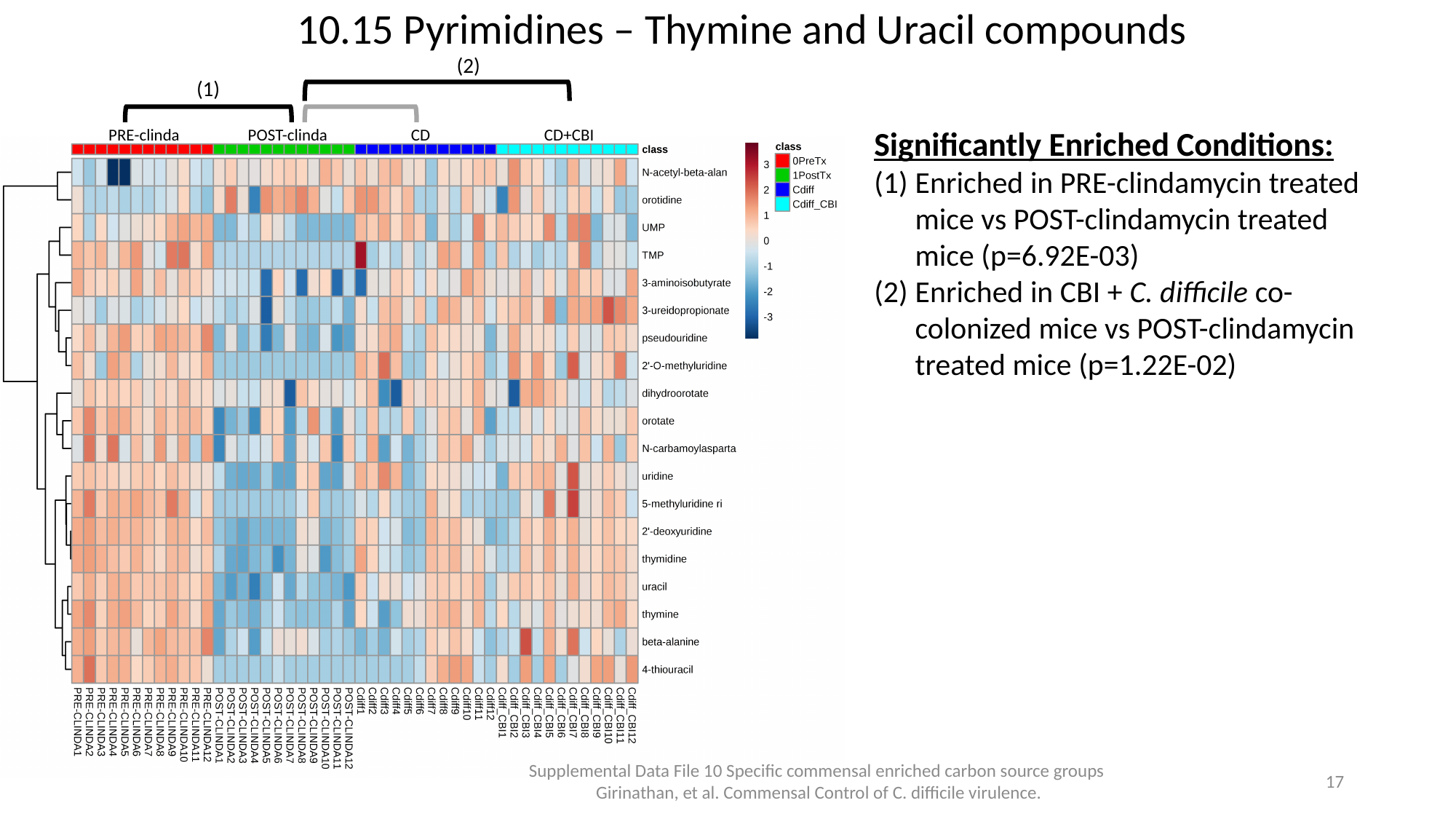

10.15 Pyrimidines – Thymine and Uracil compounds
(2)
(1)
Significantly Enriched Conditions:
Enriched in PRE-clindamycin treated mice vs POST-clindamycin treated mice (p=6.92E-03)
Enriched in CBI + C. difficile co-colonized mice vs POST-clindamycin treated mice (p=1.22E-02)
PRE-clinda POST-clinda CD CD+CBI
Supplemental Data File 10 Specific commensal enriched carbon source groups Girinathan, et al. Commensal Control of C. difficile virulence.
16

### Slide 18
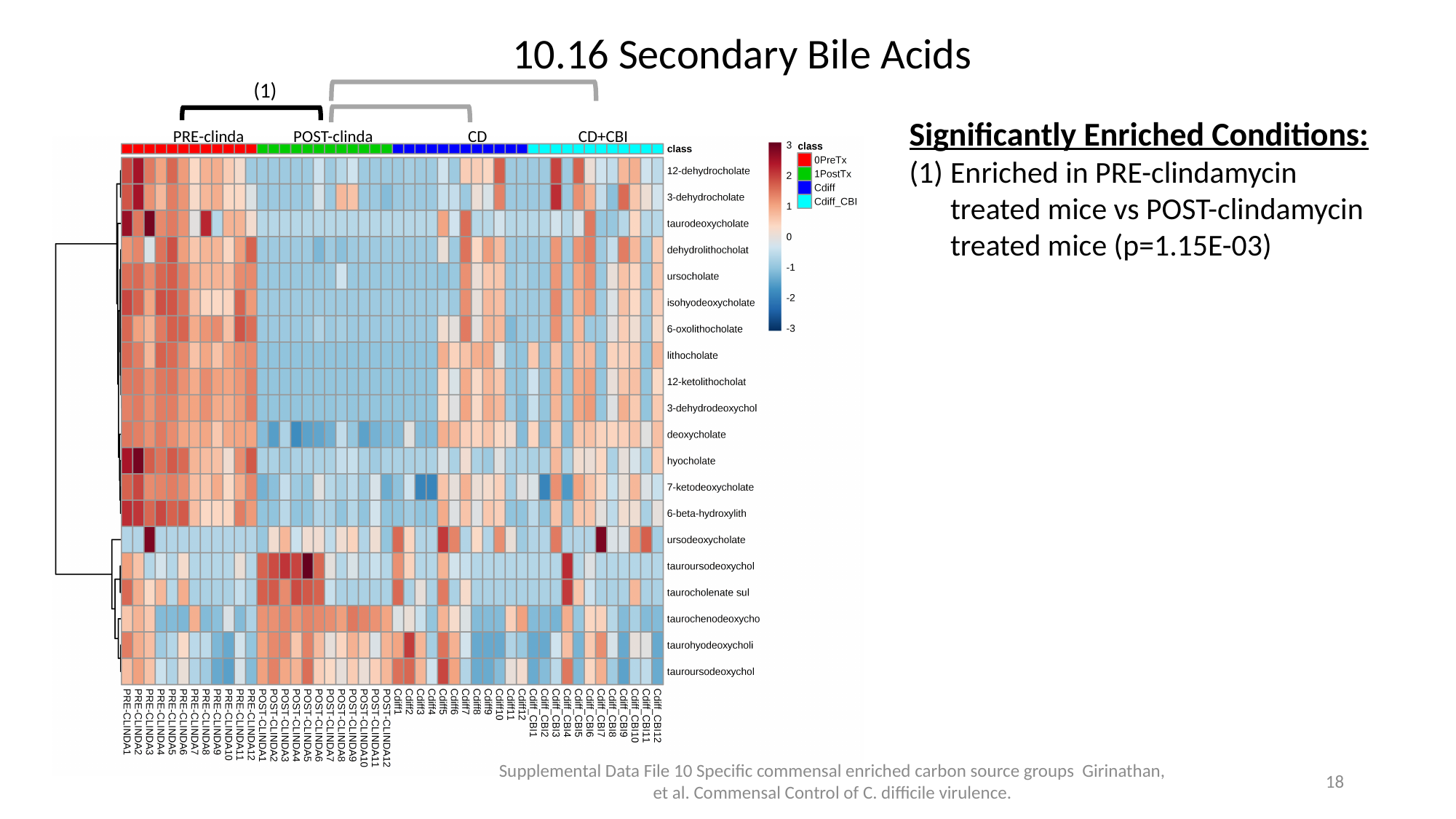

10.16 Secondary Bile Acids
(1)
Significantly Enriched Conditions:
Enriched in PRE-clindamycin treated mice vs POST-clindamycin treated mice (p=1.15E-03)
PRE-clinda POST-clinda CD CD+CBI
Supplemental Data File 10 Specific commensal enriched carbon source groups Girinathan, et al. Commensal Control of C. difficile virulence.
17

### Slide 19
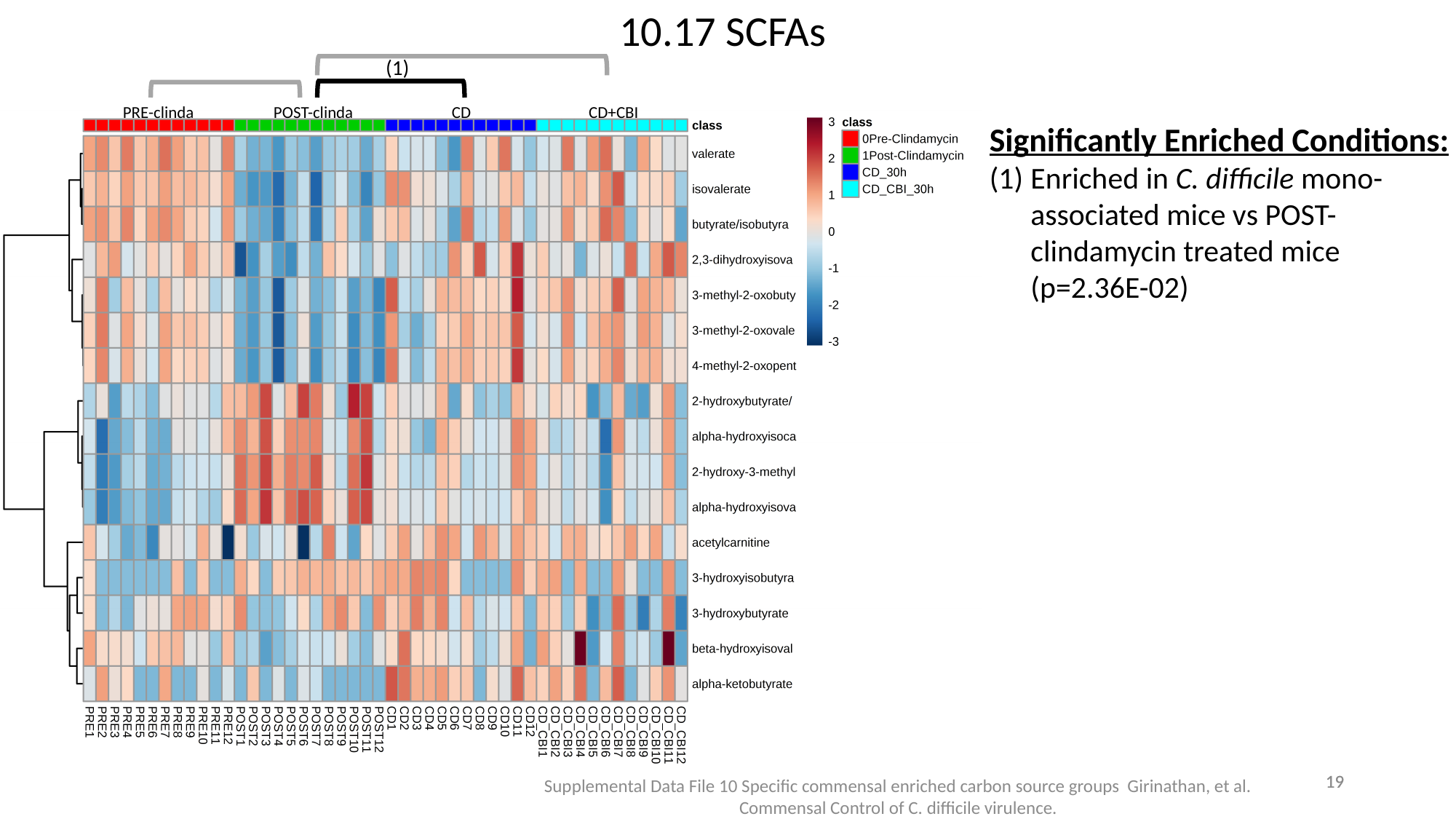

10.17 SCFAs
(1)
PRE-clinda POST-clinda CD CD+CBI
Significantly Enriched Conditions:
Enriched in C. difficile mono-associated mice vs POST-clindamycin treated mice (p=2.36E-02)
18
18
Supplemental Data File 10 Specific commensal enriched carbon source groups Girinathan, et al. Commensal Control of C. difficile virulence.

### Slide 20
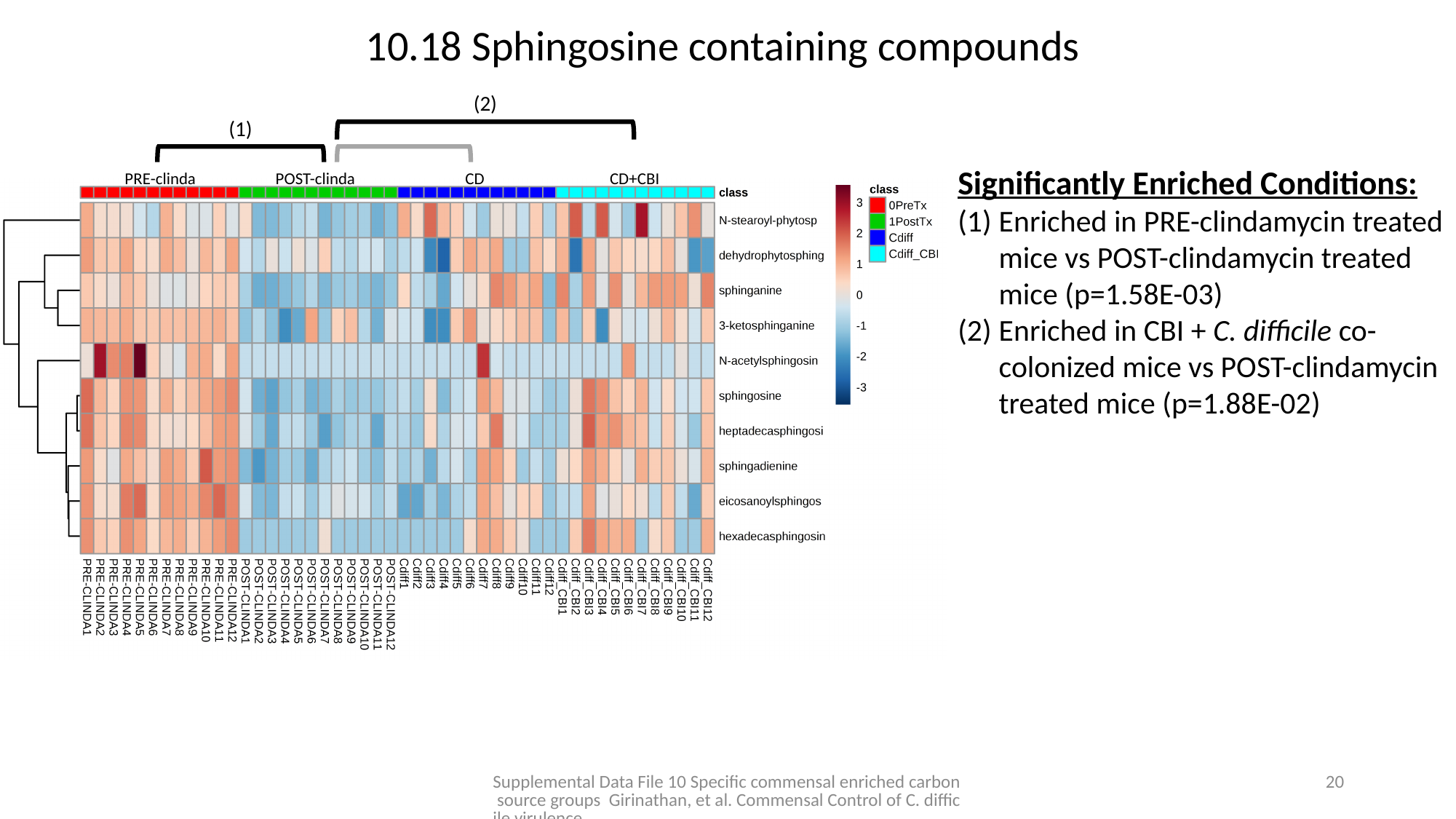

10.18 Sphingosine containing compounds
(2)
(1)
Significantly Enriched Conditions:
Enriched in PRE-clindamycin treated mice vs POST-clindamycin treated mice (p=1.58E-03)
Enriched in CBI + C. difficile co-colonized mice vs POST-clindamycin treated mice (p=1.88E-02)
PRE-clinda POST-clinda CD CD+CBI
Supplemental Data File 10 Specific commensal enriched carbon source groups Girinathan, et al. Commensal Control of C. difficile virulence.
19

### Slide 21
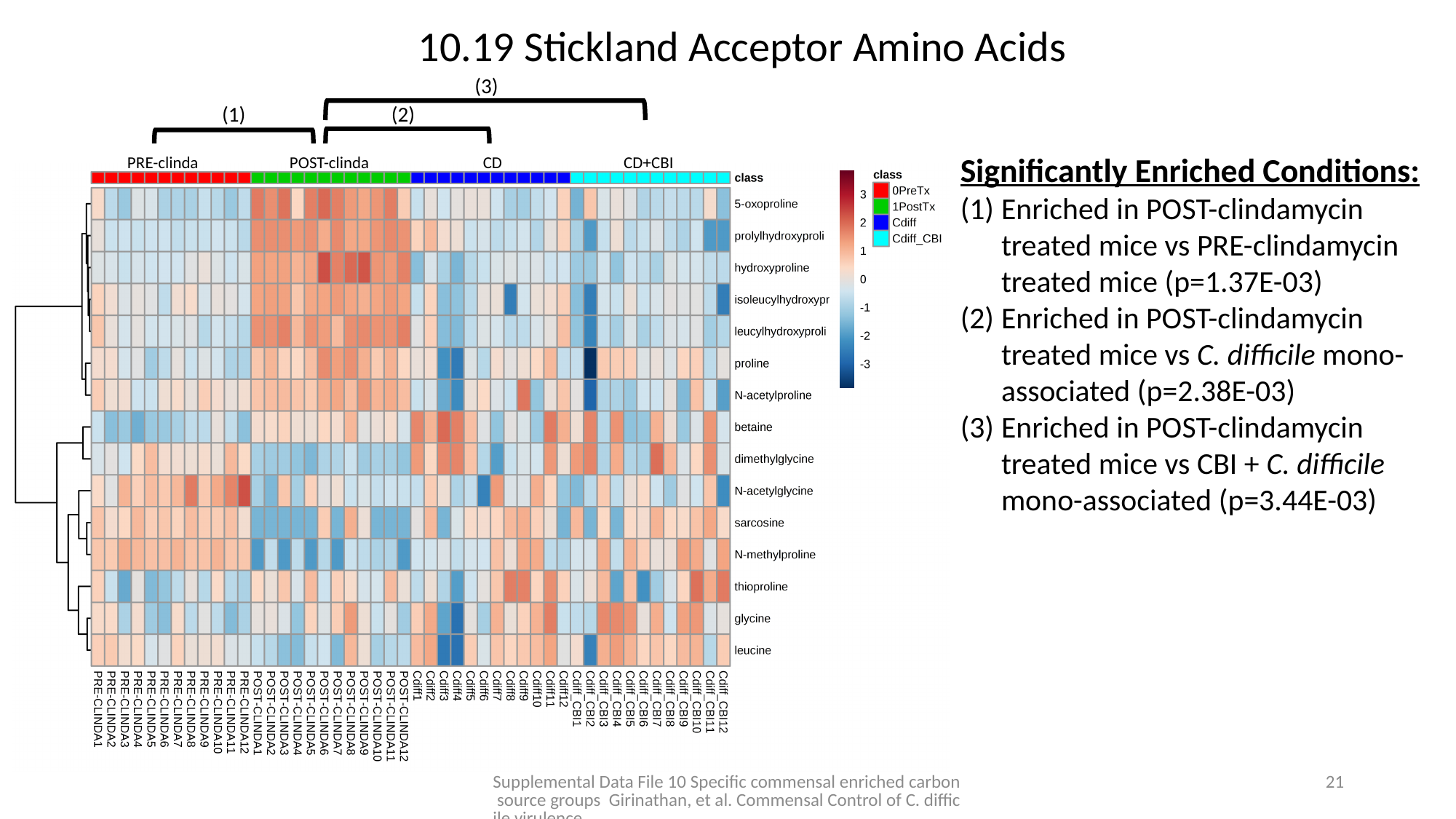

10.19 Stickland Acceptor Amino Acids
(3)
(1)
(2)
Significantly Enriched Conditions:
Enriched in POST-clindamycin treated mice vs PRE-clindamycin treated mice (p=1.37E-03)
Enriched in POST-clindamycin treated mice vs C. difficile mono-associated (p=2.38E-03)
Enriched in POST-clindamycin treated mice vs CBI + C. difficile mono-associated (p=3.44E-03)
PRE-clinda POST-clinda CD CD+CBI
Supplemental Data File 10 Specific commensal enriched carbon source groups Girinathan, et al. Commensal Control of C. difficile virulence.
20

### Slide 22
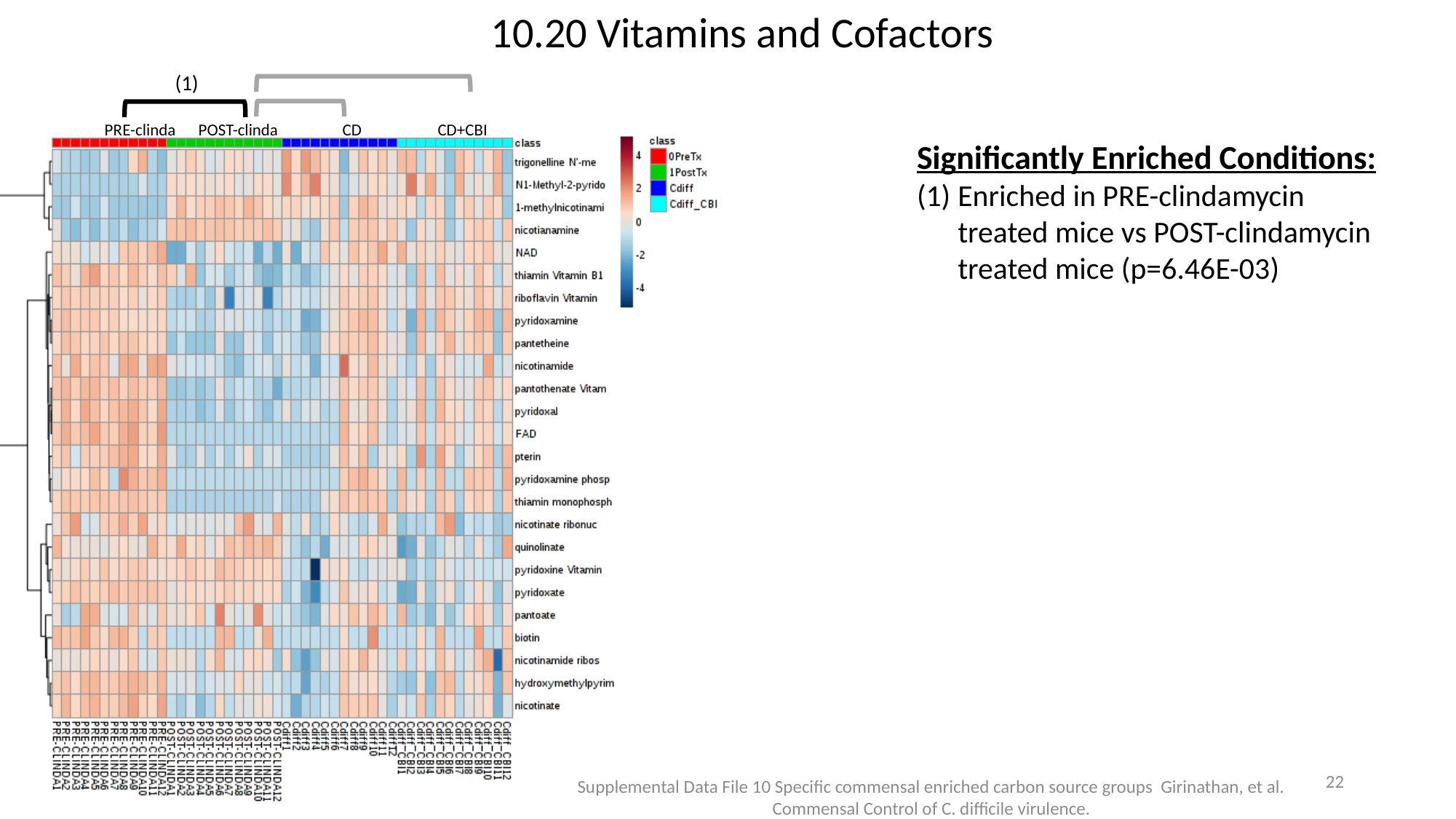

10.20 Vitamins and Cofactors
(1)
PRE-clinda POST-clinda CD CD+CBI
Significantly Enriched Conditions:
Enriched in PRE-clindamycin treated mice vs POST-clindamycin treated mice (p=6.46E-03)
21
Supplemental Data File 10 Specific commensal enriched carbon source groups Girinathan, et al. Commensal Control of C. difficile virulence.
